# Genetic and phenotypic characterization of global *Lupinus albus* genetic resources for the development of a CORE collection

**DOI:** 10.1101/2024.09.25.614894

**Authors:** Umesh Kumar Tanwar, Magdalena Tomaszewska, Katarzyna Czepiel, Mohamed Neji, Humaira Jamil, Lorenzo Rocchetti, Alice Pieri, Elena Bitocchi, Elisa Bellucci, Barbara Pipan, Vladimir Meglič, Magdalena Kroc, Roberto Papa, Karolina Susek

## Abstract

*Lupinus albus* is a food grain legume recognized for its high levels of seed protein (30–40%) and oil (6–13%), and its adaptability to different climatic and soil conditions. To develop the next generation of *L. albus* cultivars, we need access to well-characterized, genetically and phenotypically diverse germplasm. Here we evaluated more than 2000 *L. albus* accessions with passport data based on 35 agro-morphological traits to develop Intelligent CORE Collections. The reference CORE (R-CORE), representing global diversity, exemplified the genotypic variation of cultivars, breeding/research materials, landraces and wild relatives. A subset of 300 R-CORE accessions was selected as a training CORE (T-CORE), representing the diversity in the entire collection. We divided the *L. albus* R-CORE into four phenotypic groups (A1, A2, A3 and B) based on principal component analysis, with groups A3 and B distinguished by pod shattering and seed ornamentation, respectively. The coefficient of additive genetic variation differed across morphological traits, phenotypic groups, geographic regions, and according to biological status. These CORE collections will facilitate agricultural research by identifying the genes responsible for desirable traits in crop improvement programs, and by shedding light on the use of orphan genetic resources for origin and domestication studies in *L. albus*. Understanding the variation in these genetic resources will allow us to develop sustainable tools and technologies that address global challenges such as providing healthy and sustainable diets for all, and contrasting the current climate change crisis.

## 1. INTRODUCTION

Lupins belong to the legume family (Fabaceae) and grow all around the world, having adapted to various climate zones (Gladstones et al., 1998). They are cultivated as grain legumes, offering an alternative and healthy source of plant-based food (Bellucci et al., 2021, Nartea et al., 2023, Kroc et al., 2021). The seeds are rich in protein, oil, carbohydrates and fiber (Bulut et al., 2023, Duranti et al., 2008, Kroc et al., 2021) and have been proposed as traceable protein sources in Europe (de Visser et al., 2014). The cultivated species are *Lupinus angustifolius* (blue lupin)*, L. luteus* (yellow lupin)*, L. mutabilis* (pearl lupin) and *L. albus* (white lupin), the latter particularly notable due to its 30–40% seed protein content (Musco et al., 2017) and adaptability to different climate and soil conditions (Atnaf et al., 2017), resulting in its addition to the EU Novel Food status Catalogue (European Commission, 2024). The additional proposed dietary benefits of lupin seeds include the prevention of diabetes and cardiovascular disease, and the maintenance of a healthy digestive system due to the high fiber content (Gresta et al., 2017, Nartea et al., 2023).

*Lupinus albus* shows broad phenotypic diversity for several traits, many of which vary among accessions from the same geographical region or country of origin, based on passport data (Atnaf et al., 2017, Mulugeta et al., 2015, Berger et al., 2017, Zafeiriou et al., 2021). For example, among *L. albus* ecotypes from 17 countries, the pod wall to whole pod weight ratio was highest for Greek and Italian ecotypes and lowest for Egyptian ecotypes (Lagunes- Espinoza et al., 2000). On a smaller geographical scale, 45 local Greek landraces showed a 12- fold variation in seed weight and a three-fold variation in seed size (Zafeiriou et al., 2021). However, among 35 spring-sown Spanish landraces, only three of 50 quantitative parameters (related to leaf morphology) showed a variation coefficient ≥ 19% and only four of 51 qualitative descriptors (related to flower color) showed variation between accessions (Gonzalez-Andres et al., 2007). Variations have also been observed in nutritional components and minerals such as Ca (709–1284 µg/g), P (671.3–2490.2 µg/g), Fe (77.9–92.8 µg/g) and Mg (1739–2159 µg/g) in the seeds of Ethiopian cultivars (Tizazu and Emire, 2010, Zelalem and Chandravanshi, 2014). The seed protein content of 25 Ethiopian accessions was found to vary in the range 28.55–35.81% (Beyene, 2020).

Genetic resources for *L. albus* comprise 5564 accessions (only 915 of which are available for distribution) collected in gene banks (https://www.genesys-pgr.org/a/overview/v293JdOPwZG, accessed on 21.08.2024) but few studies thus far have considered the diversity of these collections (Ehab et al., 2016, Hibstu, 2016, Mulugeta et al., 2015). The utilization of plant genetic resources (PGRs) is hindered by multiple factors (Bellucci et al., 2021), preventing their integration into programs for sustainable agriculture and for crop improvementnt (McCouch et al., 2020). The characterization of PGRs is therefore necessary to protect agrobiodiversity and to promote the utilization of PGR towards the development of climate-resilient crops and agroecosystems (McCouch et al., 2020).

The CORE collection is a useful concept for the characterization of extensive diversity (Brown, 1989, Frankel, 1984, Van Hintum et al., 2000). A CORE collection is smaller than a comprehensive collection while maximizing diversity, and can therefore be used as an efficient kick-off point to enhance genetic gains, including the use of genomics, phenomics and molecular phenotyping (Bellucci et al., 2021, Bellucci et al., 2023, Lauterberg et al., 2023, Jurado et al., 2023). A new approach to conserve, manage and characterize the genetic resources of food legumes has been structured to develop intelligent collections representing the entire diversity of each crop, and by adopting an exploitation strategy based on the decentralized conservation of PGRs (Bellucci et al., 2021). Such intelligent collections feature a set of nested core collections of different sizes: (i) a reference core (R-CORE) comprising thousands of single-seed descent (SSD) lines representing the genetic resources of the species, (ii) a training core (T-CORE) based on a subset of the R-CORE and featuring 300–500 SSD lines, and (iii) a hyper-core (H-CORE) of 40–50 genotypes sampled on an evolutionary transect (Bellucci et al., 2021).

CORE collections for legumes are already available for soybean (Oliveira et al., 2010), common bean (Blair et al., 2009, García-Fernández et al., 2022, Bellucci et al., 2023), cowpea (Mahalakshmi et al., 2007), pigeon pea (Reddy et al., 2005, Upadhyaya et al., 2007), groundnut (Upadhyaya et al., 2003), chickpea (Upadhyaya et al., 2001), lentil (Tripathi et al., 2022, Tullu et al., 2001, Simon and Hannan, 1995) and lablab bean (Vaijayanthi et al., 2015). Nested intelligent core collections have been developed based on common bean SSD lines (Cortinovis et al., 2021). A large collection of ∼10,000 common bean accessions and a subsample of 500 accessions (Pv_core1) have been characterized using the genotyping-by-sequencing approach. A further subsample of 220 accessions has been genotyped by whole-genome sequencing (Bellucci et al., 2023, Cortinovis et al., 2021). Similarly, a CORE collection of 480 chickpea SSD genotypes has been phenotypically evaluated (Rocchetti et al., 2020). The only CORE collection available for *L. albus* comprises 34 lines based on the molecular characterization of 212 Ethiopian accessions using microsatellite markers (Atnaf et al., 2017).

Here, we developed the first nested core collections of *L. albus* global genetic resources with maximum representativeness and diversity. The phenotypic variation in this collection demonstrates the value of the conservation approach, allowing the sustainable utilization of these PGRs to accelerate innovative *L. albus* breeding programs aiming to improve food security, nutritional security and agricultural sustainability.

## 2. MATERIALS AND METHODS

### 2.1 Plant material and development of CORE collections

Gene banks, INCREASE project partners and stakeholders, and research institutes provided 2288 *L. albus* accessions (Supplementary Table S1). Seed material was collected as a starting point to develop SSD lines, which were derived from one randomly chosen seed from each accession. The SSD lines constituted the R-CORE (Bellucci et al., 2021, Kroc et al., 2021). The SSD lines were grown in the greenhouse at the Institute of Plant Genetics, Polish Academy of Sciences (IPG-PAN), Poznan, Poland (52°26′50.5′′N, 16°54′13.3′′E) with a temperature of 20–35 °C, a 16-h photoperiod, and a relative humidity of 60–65%. At the harvesting stage, images of qualified seeds from each accession were captured for processing (Kroc et al., 2021), and the seeds were packed in paper bags for storage.

T-CORE plants (300 accessions selected from the R-CORE) were grown at the IPG-PAN as above and also in Slovenia (Kmetijski Inštitut Slovenije, KIS), with two plantings in the spring season (March–August 2021 at IPG-PAS and KIS) and one in winter (October 2020 to January 2021, IPG-PAS). The average relative humidity and temperature were 77.9% and 18.6 °C, respectively, with a 16-h photoperiod.

### 2.2 Phenotyping of agro-morphological traits and data analysis

We used 35 agro-morphological traits for phenotypic characterization (Kroc et al., 2021): 11 qualitative (Table 1) and 24 quantitative (Table 2). Relationships between accessions were determined by multivariate analysis of the quantitative and qualitative traits using R-language v4.3.1. First, qualitative traits in the R-CORE collection were characterized to understand the overall variation in the species. Multiple correspondence analysis (MCA), which is principal component analysis (PCA) using binary data, was then applied using the *PCA ()* function in the “FactoMineR” package (Lê et al., 2008) to retain the most meaningful components. The clustering tendency in the dataset was tested with Hopkins statistics “H”, using the *hopkins ()* function in the “clustertend” package (Yilan and Rutong, 2015). Hierarchical clustering of the retained principal components (HCPC) was achieved using the *HCPC ()* function in the “FactoMineR” package. The graphical outputs were visualized using the function *fviz_cluster ()* in the “factoextra” package (Alboukadel and Mundt, 2017). To characterize the phenotypic diversity using 24 quantitative traits, descriptive statistics (minimum, maximum, mean, standard deviation, coefficient of variation, and frequency distribution) were computed and the normality was tested using Kolmogorov–Smirnov (KS) and Shapiro–Wilk normality tests. The quantitative traits were assessed by PCA using the “FactoMineR” package and one-way analysis of variance (ANOVA) followed by Tukey’s multiple comparisons test (P ≤ 0.01) to explore differences in the accessions representing different phenotypic groups, geographical regions and biological statuses. Pearson’s or Spearman’s correlation tests (as appropriate) were applied to find significant relationships between qualitative and quantitative traits using the “corrplot” and “GGally” packages. We also used packages such as “ggplot2” (graphic presentation), “sf”, “rworldxtra” and “rnaturalearth” (for mapping and geo-referencing).

**Table 1:** Qualitative traits used for the phenotypic characterization of *Lupinus albus* (Kroc et al., 2021).

| No. | Descriptors | Code | Descriptor state |
| --- | --- | --- | --- |
| 1 | Color of wing petals at full flowering on the main stem | F6 | Blue, pink, violet, white, white/blue, white/pink, white/violet |
| 2 | Color of standard petal at full flowering on the main stem | F7 | Blue, pink, violet, white, white/blue, white/pink, white/violet |
| 3 | Color of the keel edge at full flowering on the main stem | F8 | Brownish, not visible, transparent, white, white/blue |
| 4 | Purple pigmentation of the main stem after two months of emergence | PP | Medium intensity, no pigment +purple internodes, none, slightly, very intense |
| 5 | Pod shattering on the main stem | P4_m | Complete, moderate, none, slight |
| 6 | Pod shattering on the lateral branches | P4_l | Moderate, none, slight |
| 7 | Seed shape | S5 | Cuboid, flattened oval, flattened cuboid, flattened spherical |
| 8 | Seed ornamentation | S6 | Marbled, none, other |
| 9 | Seed ornamentation color | S7 | Black, dark brown, light brown, none, other |
| 10 | Density of seed ornamentation | S8 | Dense, medium, none, very dense |
| 11 | Growth habit | GH | Determinate, indeterminate, semi-determinate |

**Table 2:** Quantitative traits used for the phenotypic characterization of *Lupinus albus* (Kroc et al., 2021).

|  | No. | Descriptors | Code | Unit |
| --- | --- | --- | --- | --- |
| **Emergence** | 1 | Days to emergence | Emer | Days |
| **Flowering** | 2 | Days to flowering (length of vegetative phase) | F1_m | Days |
|  | 3 | Days to first petal occurrence on the main stem | F2_m | Days |
|  | 4 | Days to first flower occurrence on the main stem | F3_m | Days |
|  | 5 | Days to the full flowering on the main stem | F4_m | Days |
|  | 6 | Days to the first flower withering on the main stem | F5_m | Days |
|  | 7 | Length of the generative phase on the main stem I | Gen | Days |
| **Pod development and physiological maturity** | 8 | Days to the first pod set on the main stem | P1_m | Days |
|  | 9 | Days to the first pod set on the lateral branches | P1_l | Days |
|  | 10 | Days to the first pod color changes on the main stem | P2_m | Days |
|  | 11 | Days to the first pod color changes on the lateral branches | P2_l | Days |
|  | 12 | Length of the pod and seed growth phase on the main stem I | Mat1_m | Days |
|  | 13 | Length of the pod and seed growth phase on the main stem II | Mat1_l | Days |
|  | 14 | Length of the pod and seed growth phase on the lateral branches II | Mat2_m | Days |
|  | 15 | Length of the pod and seed growth phase on the lateral branches II | Mat2_l | Days |
| **Pods and seeds production** | 16 | Number of pods per plant on the main stem | P3_m | Numbers |
|  | 17 | Number of pods per plant on the lateral branches | P3_l | Numbers |
|  | 18 | Number of qualified seeds on the main stem | S1_m | Numbers |
|  | 19 | Number of qualified seeds on the lateral branches | S1_1 | Numbers |
|  | 20 | Total number of qualified seeds | S2 | Numbers |
|  | 21 | Total mass of the qualified seeds [g] | S3 | g |
|  | 22 | 1000 seed weight [g] | S4 | g |
| **Plant vigor** | 23 | Plant height of the main stem [cm] | H1 | cm |
|  | 24 | Total plant height [cm] | H2 | cm |

To partition the phenotypic variance (V_p_) into additive genetic (V_a_) and residual (V_r_) components, we used an REML-linear mixed model (LMM) for each trait in the package “lme4” (Bates et al., 2014). We used the equation *Yi = µ + Ai + εi*, where *Yi* is the response variable of accession *i*, *µ* is the trait mean, *Ai* is the accession’s additive genetic value, and *εi* is the residual value. For comparison across traits, phenotypic groups, geographical regions and biological statuses, we calculated the coefficient of additive genetic variation (CV_a_) according to Garcia-Gonzalez et al. (2012) using the equation *CV_a_* = √*V*_a_/*µ*, where *V*_a_ is the additive genetic variance and *µ* is the mean of the trait. CVa, in contrast to heritability, is a measure of additive genetic variation that is standardized to the trait mean, so is independent of other sources of variance (Gioia et al., 2015, Pieri et al., 2024) .

### 2.3 Characterization of the T-CORE

The T-CORE collection was established to maximize the diversity present in the R-CORE depending on the availability of the seeds, as described in the INCREASE project (Bellucci et al., 2021). The phenotypic data were standardized to eliminate scale differences, and different combinations of variables were used to assess the T-CORE according to qualitative traits, quantitative traits, a combination of quantitative and qualitative traits, and a combination of quantitative and qualitative traits and passport data.

To ensure the T-CORE collection was representative of the R-CORE collection, we applied ‘EvaluateCore’ v0.1.2 (Aravind et al., 2022) to determine the frequency distributions for geographical regions, countries or origin, and biological status, as well as qualitative traits, using a Chi-squared (χ^2^) test for homogeneity. Coverage of the T-CORE qualitative traits was estimated using the equation for class coverage: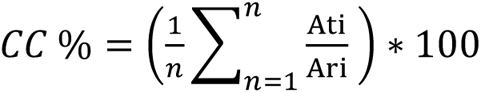 (Kim et al., 2007), where *At_i_* and *Ar_i_* are the number of categories for the *i*^th^ trait observed in the T-CORE and R-CORE collections, respectively, and *n* is the number of qualitative traits. We applied the Shannon–Weaver diversity index (Shannon and Weaver, 1949) to measure phenotypic diversity for each qualitative trait in both collections using the formula H’ = -Ʃ*_i_^n^ p_i_* log_2_ *p_i_* where *p_i_* is the frequency of accessions from the *i*^th^ class and *n* is the number of classes for the designated trait. Furthermore, we computed Pearson’s coefficient (*r*) between quantitative traits and Cramer’s V coefficient (*v*) between qualitative traits in the R-CORE and T-CORE using packages ‘corrplot’ (Wei et al., 2017) and ‘vcd’ (Meyer et al., 2023), respectively, to assess the extent of trait associations captured in the T-CORE collection.

## 3. RESULTS

### 3.1 Development and characterization of the L. albus R-CORE collection

The *L. albus* R-CORE collection contained 2288 accessions and originated from 43 countries distributed over nine geographical regions (Figure 1A, Table 3). Most of the accessions (76.39%) were from Europe, mainly Southern Europe (1336 accessions, 58.39%), followed by Western Europe (209, 9.11%) and Eastern Europe (203, 8.87%). Outside Europe, most accessions originated from Northern Africa (208, 9.09%), followed by the Middle East (47, 2.05%), America (46, 2.01%) and Eastern Africa (40, 1.74%). Australia and New Zealand (11, 0.48%) and Southern Asia (1, 0.04%) were the least representative regions. Another 187 accessions (8.17%) were described as “unknown origin”. In the Southern Europe category, most accessions originated from Spain (615, 26.88%) and Portugal (439, 19.18%).

**Figure 1:**
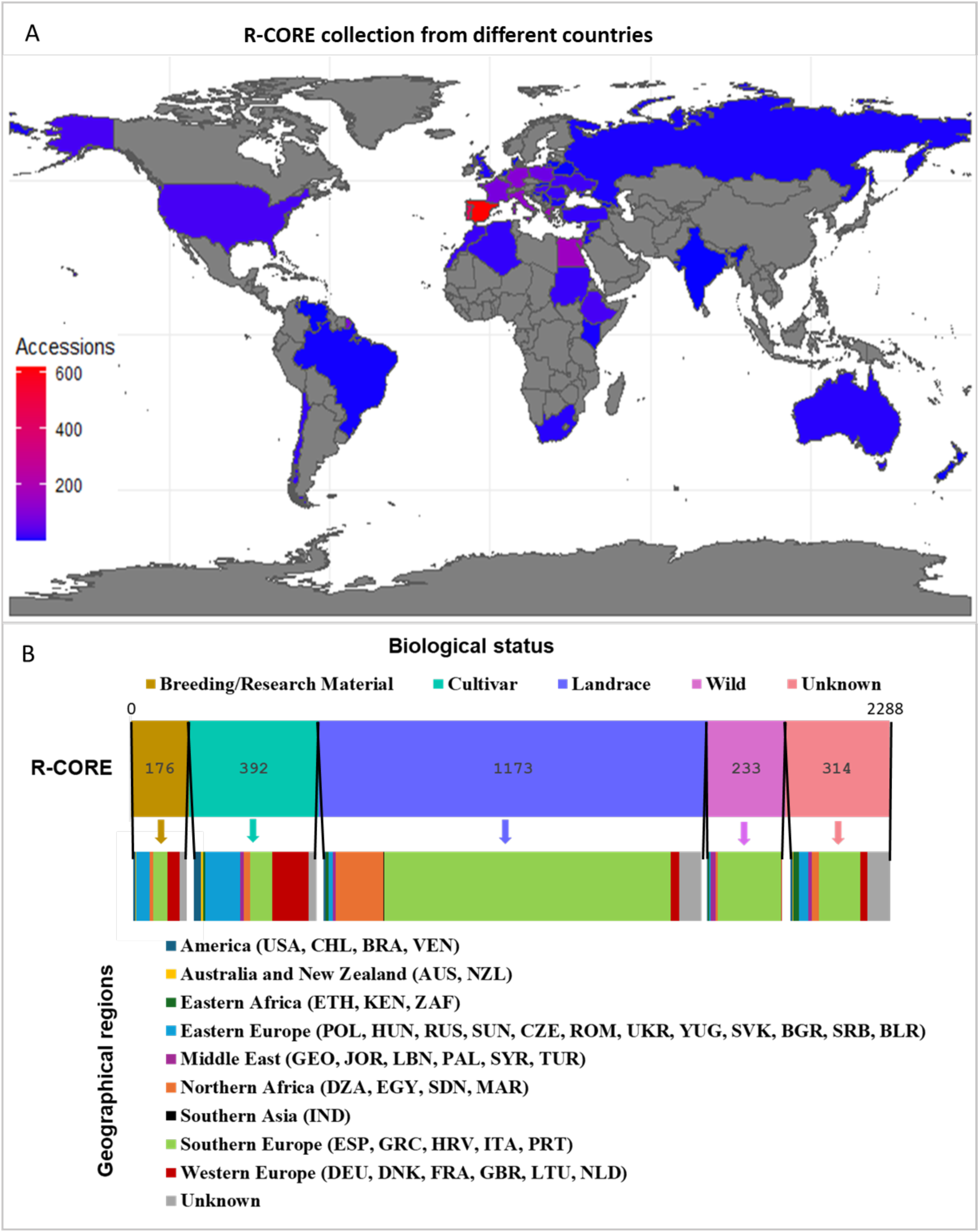
Characterization of *Lupinus albus* accessions in the R-CORE collection based on (A) country of origin and (B) biological status, subdivided by geographical region.

**Table 3:** Number (N) and percentage of accessions from different geographical regions (subdivided by county^#^) in the *L. albus* R-CORE and T-CORE collections (ns = nonsignificant).

|  | **R-CORE**<br>(N = 2288) |  | **T-CORE**<br>(N = 300) |  |  |  |
| --- | --- | --- | --- | --- | --- | --- |
| **Geographical region** | **N** | **%** | **N** | **%** | **$\chi^2$** | **p-value** |
| **Southern Europe** | 1336 | 58.4 | 173 | 57.7 | 0 | ns |
| Spain (ESP) | 615 | 26.8 | 96 | 32 | 4 | 0.049 * |
| Greece (GRC) | 152 | 6.6 | 24 | 8 | 0.8 | ns |
| Croatia (HRV) | 2 | 0.1 | 0 | 0 | 0.3 | ns |
| Italy (ITA) | 128 | 5.6 | 40 | 13.3 | 28.4 | <0.001** |
| Portugal (PRT) | 439 | 19.2 | 13 | 4.3 | 40.5 | <0.001** |
| **Western Europe** | 209 | 9.1 | 33 | 10.3 | 0.5 | ns |
| Germany (DEU) | 110 | 4.8 | 24 | 7.7 | 5 | <0.05* |
| Denmark (DNK) | 1 | 0 | 0 | 0 | 0.1 | ns |
| France (FRA) | 86 | 3.8 | 6 | 1.7 | 3.7 | ns |
| United Kingdom (GBR) | 7 | 0.3 | 1 | 0.3 | 0 | ns |
| Lithuania (LTU) | 1 | 0 | 1 | 0.3 | 2.9 | ns |
| Netherlands (NLD) | 4 | 0.2 | 1 | 0.3 | 0.4 | ns |
| **Northern Africa** | 208 | 9.1 | 21 | 7 | 1.4 | ns |
| Algeria (DZA) | 14 | 0.6 | 4 | 1.3 | 2.1 | ns |
| Egypt (EGY) | 159 | 6.9 | 12 | 4 | 3.7 | ns |
| Sudan (SDN) | 17 | 0.7 | 3 | 1 | 0.2 | ns |
| Morocco (MAR) | 18 | 0.8 | 2 | 0.7 | 0.1 | ns |
| **Eastern Europe** | 203 | 8.9 | 25 | 8.3 | 0.1 | ns |
| Poland (POL) | 67 | 2.9 | 10 | 3.3 | 0.2 | ns |
| Hungary (HUN) | 28 | 1.2 | 5 | 1.7 | 0.4 | ns |
| Russia (RUS) | 5 | 0.2 | 1 | 0.3 | 0.2 | ns |
| USSR (SUN) | 34 | 1.5 | 4 | 1.3 | 0 | ns |
| Czech Republic (CZE) | 9 | 0.4 | 1 | 0.3 | 0 | ns |
| Romania (ROM) | 18 | 0.8 | 0 | 0 | 2.5 | ns |
| Ukraine (UKR) | 22 | 1 | 2 | 0.7 | 0.3 | ns |
| Serbia and Montenegro (YUG) | 9 | 0.4 | 2 | 0.7 | 0.5 | ns |
| Slovakia (SVK) | 1 | 0 | 0 | 0 | 0.1 | ns |
| Bulgaria (BGR) | 5 | 0.2 | 0 | 0 | 0.7 | ns |
| Serbia (SRB) | 4 | 0.2 | 0 | 0 | 0.5 | ns |
| Belarus (BLR) | 1 | 0 | 0 | 0 | 0.1 | ns |
| **Middle East** | 47 | 2.1 | 17 | 5.7 | 14.4 | <0.001** |
| Georgia (GEO) | 2 | 0.1 | 1 | 0.3 | 13.9 | ns |
| Jordan (JOR) | 5 | 0.2 | 2 | 0.7 | 2 | ns |
| Lebanon (LBN) | 2 | 0.1 | 0 | 0 | 0.3 | ns |
| Palestine (PAL) | 10 | 0.4 | 6 | 2 | 10.6 | <0.001 |
| Syria (SYR) | 14 | 0.6 | 2 | 0.7 | 0 | ns |
| Turkey (TUR) | 14 | 0.6 | 6 | 2 | 6.7 | <0.05* |
| **America** | 46 | 2 | 3 | 1 | 1.5 | ns |
| Chile (CHL) | 15 | 0.7 | 2 | 0.7 | 0 | ns |
| Brazil (BRA) | 3 | 0.1 | 0 | 0 | 0.4 | ns |
| Venezuela (VEN) | 1 | 0 | 0 | 0 | 0.1 | ns |
| **Eastern Africa** | 40 | 1.7 | 7 | 2.3 | 0.4 | ns |
| Ethiopia (ETH) | 27 | 1.2 | 5 | 1.7 | 0.5 | ns |
| Kenya (KEN) | 4 | 0.2 | 0 | 0 | 0.5 | ns |
| South Africa (ZAF) | 9 | 0.4 | 2 | 0.7 | 0.5 | ns |
| **Australia and New Zealand** | 11 | 0.5 | 1 | 0.3 | 0.1 | ns |
| Australia (AUS) | 9 | 0.4 | 1 | 0.3 | 0 | ns |
| New Zealand (NZL) | 2 | 0.1 | 0 | 0 | 0.3 | ns |
| **Southern Asia** | 1 | 0 | 0 | 0 | 0.1 | ns |
| India (IND) | 1 | 0 | 0 | 0 | 0.1 | ns |
| **Unknown** | 187 | 8.2 | 20 | 6.7 | 0.9 | ns |

Passport data (Figure 1B, Table 4) revealed that the majority of the R-CORE collection was made up of landraces (1173, 51.26%), but also cultivars (392, 17.13%) and wild accessions (233, 10.18%), as well as breeding/research materials (176, 7.69%). However, the biological status of 314 (13.72%) accessions was unknown. The majority of breeding/research materials and cultivars originated from Europe (Eastern, Western and Southern). The landraces were mainly represented by Southern Europe and Northern Africa, whereas the wild accessions mainly originated from Southern Europe and the Middle East (Figure 1B, Figure S1).

**Table 4:** Number, percentage, and χ^2^ value of accessions representing different biological statuses in the *L. albus* R-CORE and T-CORE collections, based on passport data (ns = nonsignificant).

| Biological status | R-CORE | | T-CORE | | $\chi^2$ | p-value |
| --- | --- | --- | --- | --- | --- | --- |
|  | N | % | N | % |  |  |
| Cultivar | 392 | 17.1 | 52 | 17.3 | 0.0 | ns |
| Landrace | 1173 | 51.3 | 89 | 29.7 | 51.0 | <0.001 |
| Breeding/research material | 176 | 7.7 | 42 | 14.0 | 15.2 | <0.001** |
| Wild accession | 233 | 10.2 | 75 | 25.0 | 58.6 | <0.001** |
| Unknown | 314 | 13.7 | 42 | 14.0 | 0.0 | ns |

#### 3.1.1 Characterization of the R-CORE collection based on qualitative traits

The R-CORE collection showed considerable diversity for most of the qualitative traits (Figure 2, Table S2). The flower color traits at the full flowering stage, namely the color of the wing petal (F6) and standard petal (F7), feature seven descriptors. For the wing petals, the most common colors were white/blue and white/violet (1451 and 383 accessions, respectively), whereas white and white/blue were the most common for the standard petals (1103 and 970 accessions, respectively). Moreover, for most accessions, the keel edge (F8) was either non-visible (1594) or transparent (662). In terms of stem pigmentation, most of the accessions (1450) featured purple internodes only, whereas in others the pigmentation was either absent (439) or very intense (49). In terms of seed shape, flattened spherical (1268) and flattened cuboid (846) were the most common. All accessions formed white seeds, in most cases non-ornamented (2237). Ornamented seeds were identified in only 51 accessions, 50 of which were of the marbled type. The growth habit trait was mainly indeterminate or semi-determinate (1785 and 445 accessions, respectively). Statistical analysis for all the qualitative traits revealed maximum diversity for the color of wing petals (F6), the purple pigmentation on the main stem (PP) and the color of the standard petals (F7), with H′ values of 0.51, 0.48 and 0.44, respectively (Table 5). Minimum diversity was observed for pod shattering on the main stem (P4_m) and on lateral branches (P4_l) with H′ values of 0.04 and 0.01, respectively.

**Figure 2:**
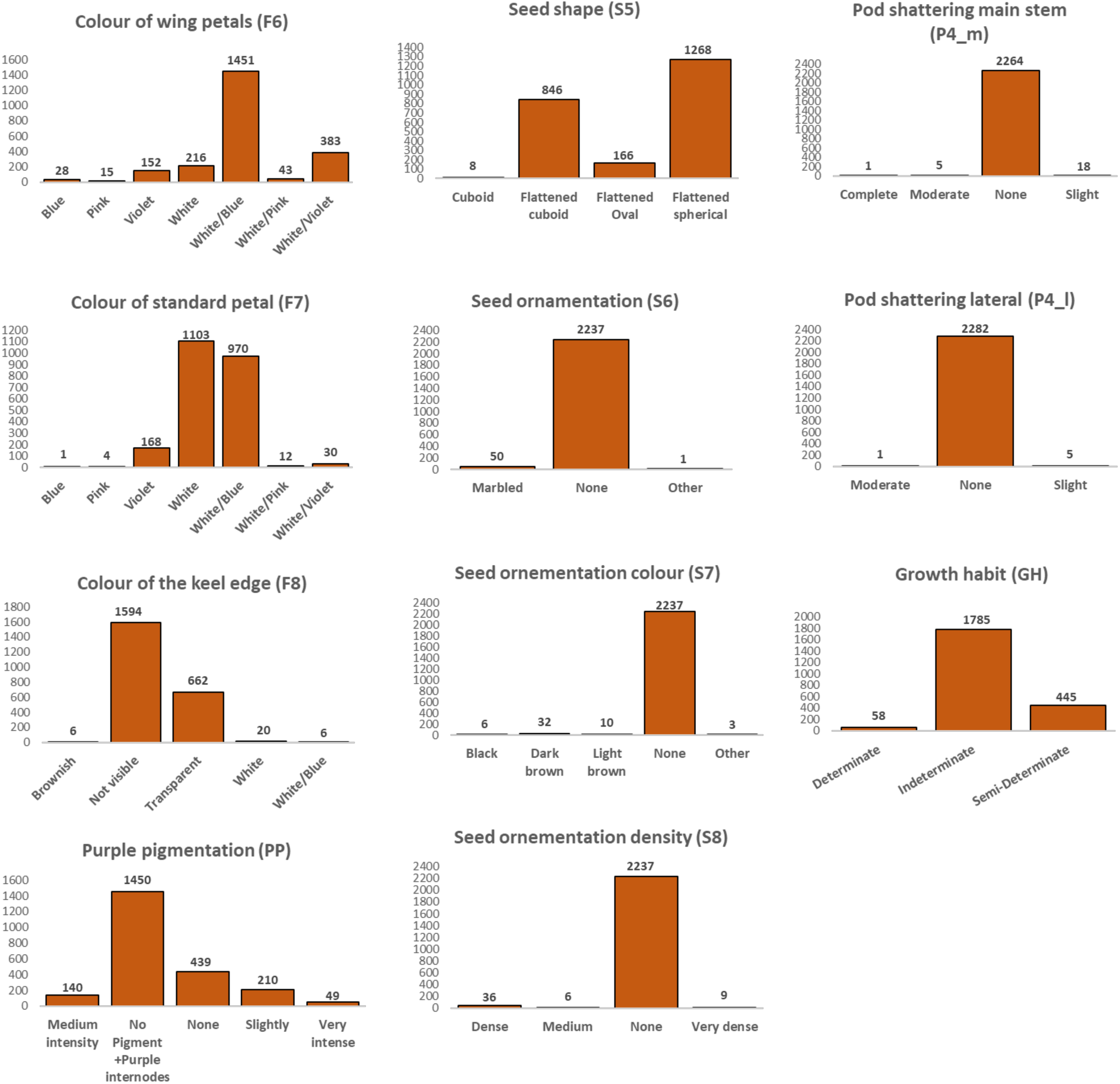
Frequency distribution of qualitative traits in the *L. albus* R-CORE collection.

**Table 5:**
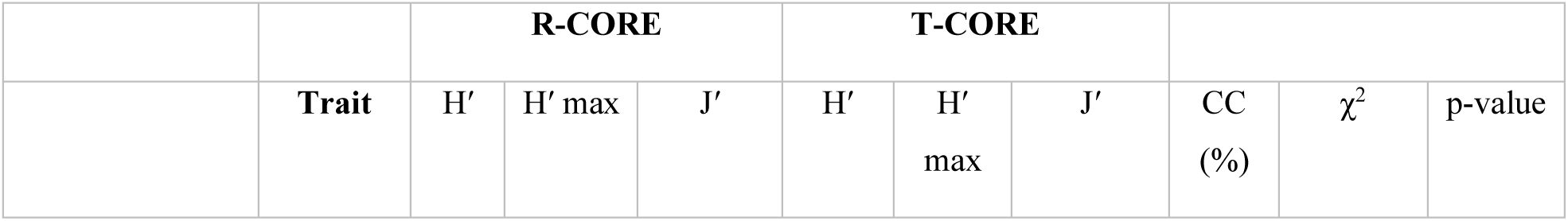
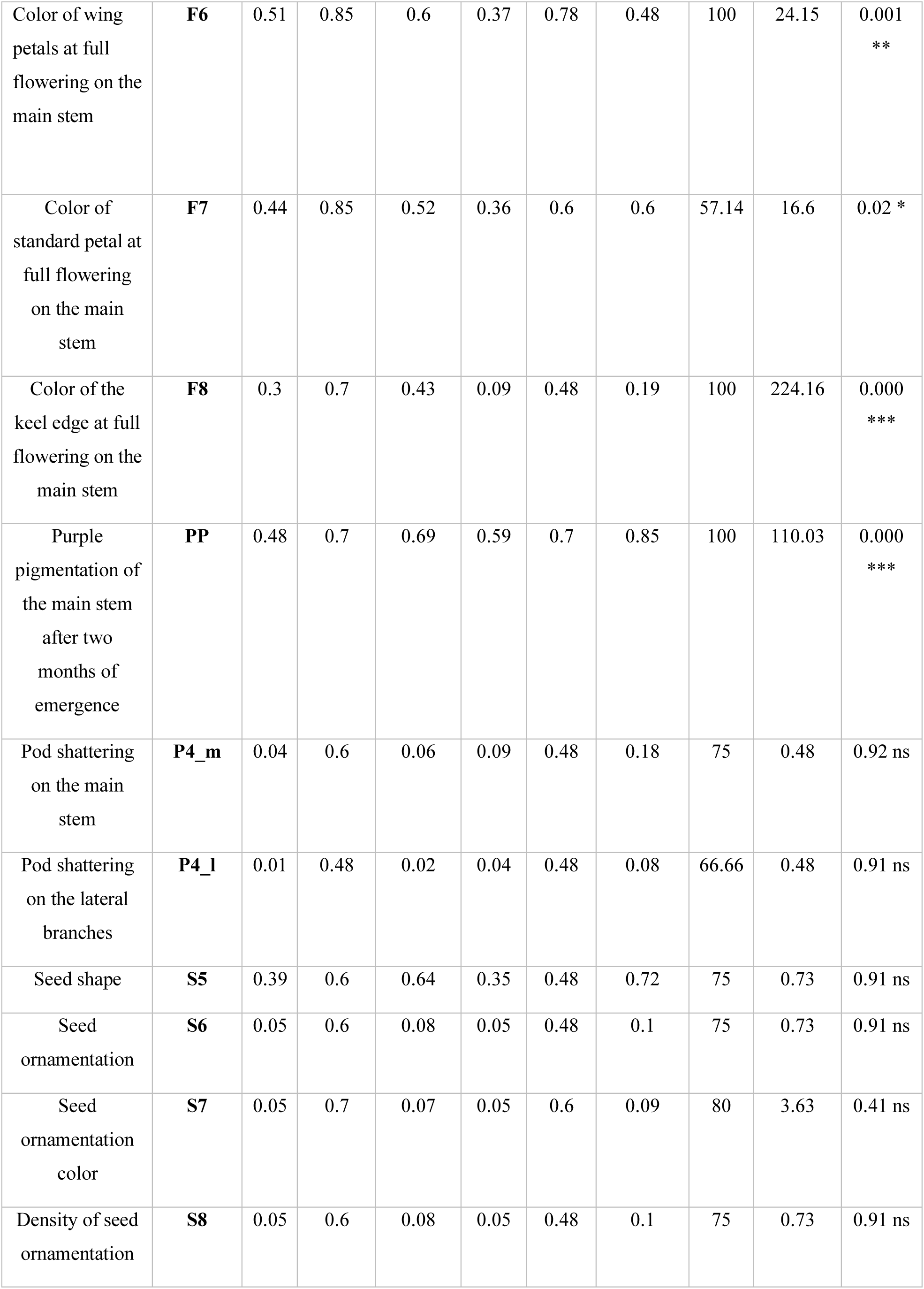
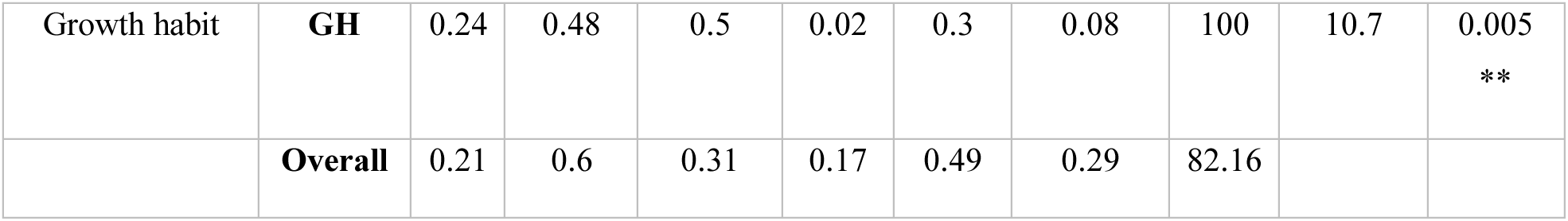
Patterns of variation and χ^2^ values for the frequency distributions of qualitative descriptors in the *L. albus* R-CORE and T-CORE collections. H′, H′_max_ and J′ values indicate the Shannon–Weaver diversity index, maximum diversity and H′-based evenness, respectively (values in bold are significant at p < 0.05; CC (%) = coverage class, ns = nonsignificant).

| Trait | R-CORE |  |  | T-CORE |  |  |  |  |  |
| --- | --- | --- | --- | --- | --- | --- | --- | --- | --- |
| | $H'$ | $H'_{\max}$ | $J'$ | $H'$ | $H'_{\max}$ | $J'$ | CC (%) | $\chi^2$ | p-value |

|  |  |  |  |  |  |  |  |  |  |  |
| --- | --- | --- | --- | --- | --- | --- | --- | --- | --- | --- |
| Color of wing petals at full flowering on the main stem | **F6** | 0.51 | 0.85 | 0.6 | 0.37 | 0.78 | 0.48 | 100 | 24.15 | 0.001<br>** |
| Color of standard petal at full flowering on the main stem | **F7** | 0.44 | 0.85 | 0.52 | 0.36 | 0.6 | 0.6 | 57.14 | 16.6 | 0.02 * |
| Color of the keel edge at full flowering on the main stem | **F8** | 0.3 | 0.7 | 0.43 | 0.09 | 0.48 | 0.19 | 100 | 224.16 | 0.000<br>*** |
| Purple pigmentation of the main stem after two months of emergence | **PP** | 0.48 | 0.7 | 0.69 | 0.59 | 0.7 | 0.85 | 100 | 110.03 | 0.000<br>*** |
| Pod shattering on the main stem | **P4_m** | 0.04 | 0.6 | 0.06 | 0.09 | 0.48 | 0.18 | 75 | 0.48 | 0.92 ns |
| Pod shattering on the lateral branches | **P4_l** | 0.01 | 0.48 | 0.02 | 0.04 | 0.48 | 0.08 | 66.66 | 0.48 | 0.91 ns |
| Seed shape | **S5** | 0.39 | 0.6 | 0.64 | 0.35 | 0.48 | 0.72 | 75 | 0.73 | 0.91 ns |
| Seed ornamentation | **S6** | 0.05 | 0.6 | 0.08 | 0.05 | 0.48 | 0.1 | 75 | 0.73 | 0.91 ns |
| Seed ornamentation color | **S7** | 0.05 | 0.7 | 0.07 | 0.05 | 0.6 | 0.09 | 80 | 3.63 | 0.41 ns |
| Density of seed ornamentation | **S8** | 0.05 | 0.6 | 0.08 | 0.05 | 0.48 | 0.1 | 75 | 0.73 | 0.91 ns |
| Growth habit | **GH** | 0.24 | 0.48 | 0.5 | 0.02 | 0.3 | 0.08 | 100 | 10.7 | 0.005<br>** |
|  | **Overall** | 0.21 | 0.6 | 0.31 | 0.17 | 0.49 | 0.29 | 82.16 |  |  |

#### 3.1.2 Characterization of the R-CORE collection based on quantitative traits

The R-CORE collection also showed considerable variability for most of the quantitative traits, and revealed a non-normal distribution in all cases (Figure 3). Descriptive statistics are presented in Table 6. The 1000 seed weight showed the widest range (code S4; range 0–680 g; mean ± SD = 144.33 ± 2.10 g). This was followed by the length of the pod and seed growth phase on the main stem II (Mat1_l; 3.75–156 days; 115.45 ± 0.83 days), the length of the pod and seed growth phase on the main stem I (Mat1_m; 13.18–136 days; 107.93 ± 0.65 days), and total plant height (H2; 17–134 cm; 70.27 ± 0.38 cm). The lowest variability was observed for the number of pods per plant on the main stem (P3_m; 0–11; 1.90 ± 0.03) and the number of pods per plant on the lateral branches (P3_l; 0–16; 2.88 ± 0.06). The highest coefficient of variation (CV) was observed for the number of seeds on lateral branches (S1_l; CV = 1.50%), on the main stem (S1_m; 1.16%), and in total (S2; 1.15%). The lowest CV of 0.11% was observed for the number of days to full flowering on the main stem (F4_m) and to the first pod set on the main stem (P1_m).

**Figure 3:**
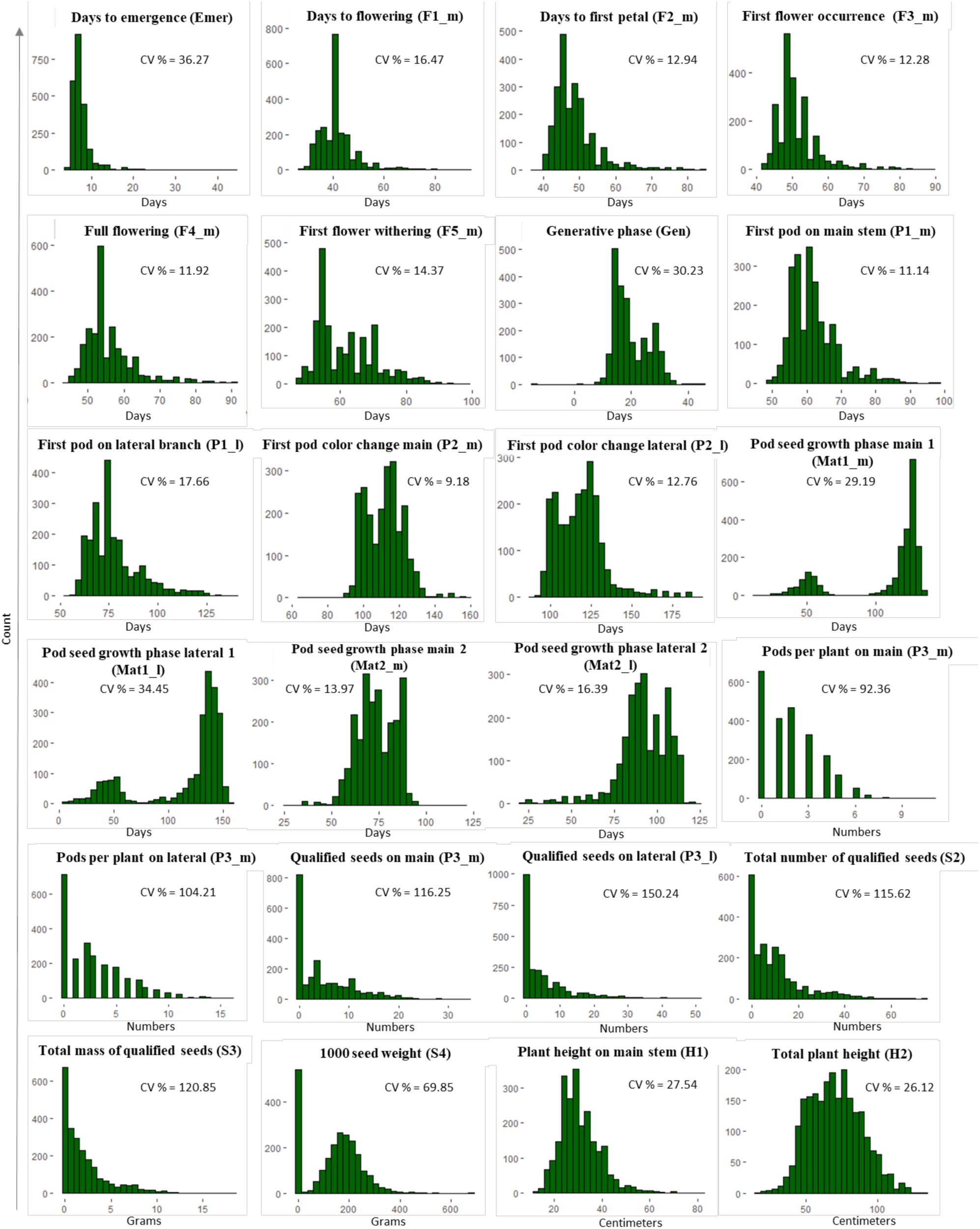
Frequency distribution of 24 quantitative traits in the *L. albus* R-CORE collection (CV = coefficient of variation).

**Table 6:**
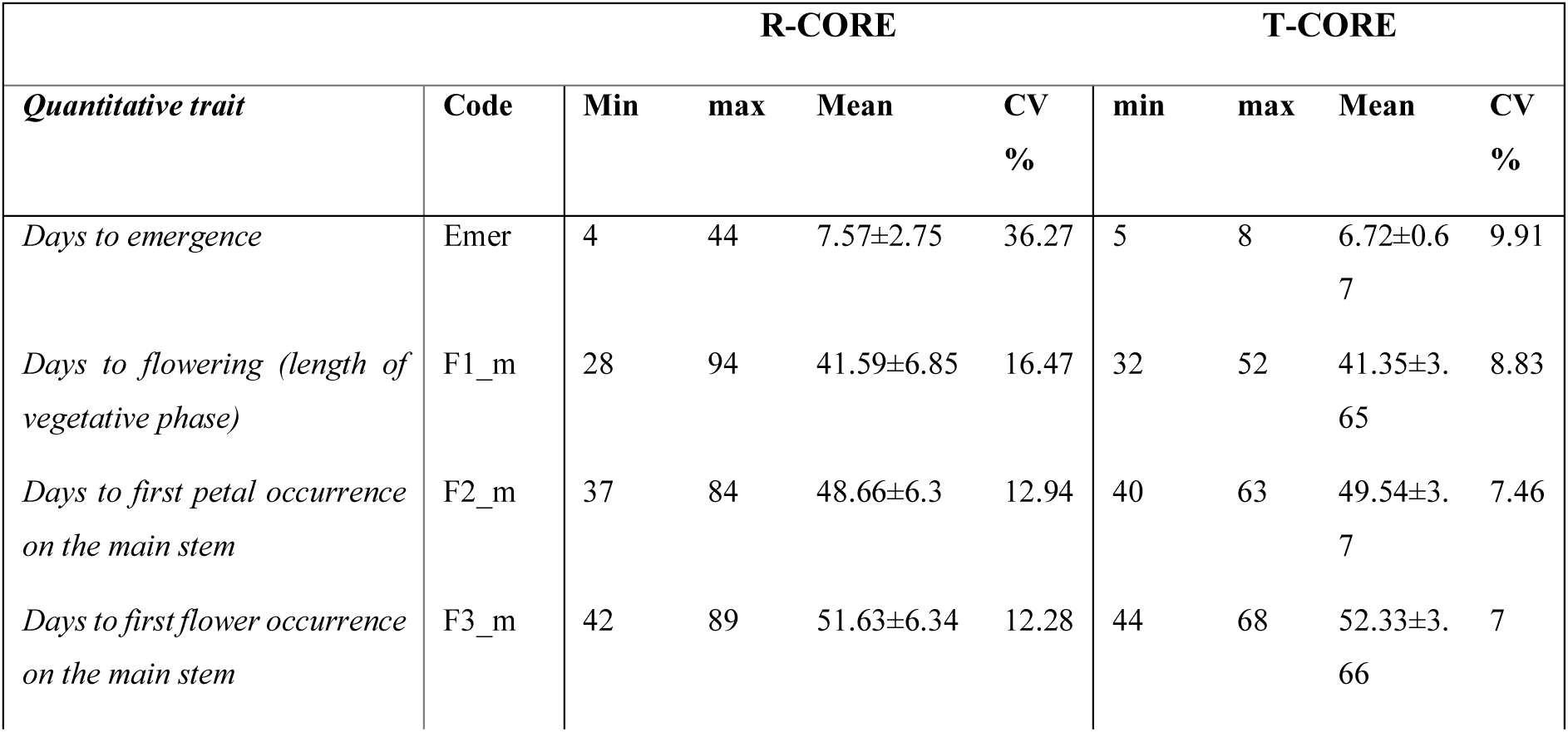
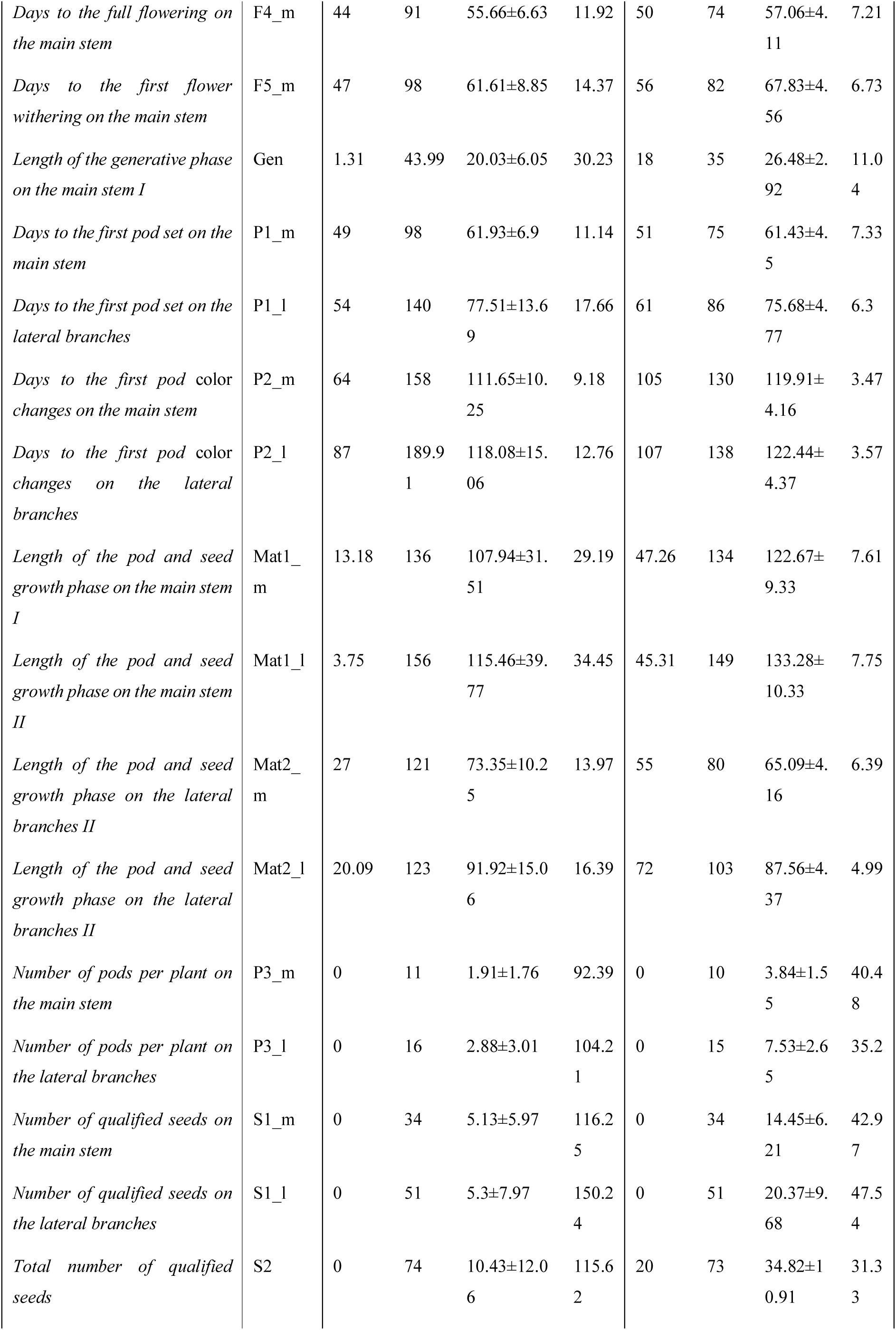
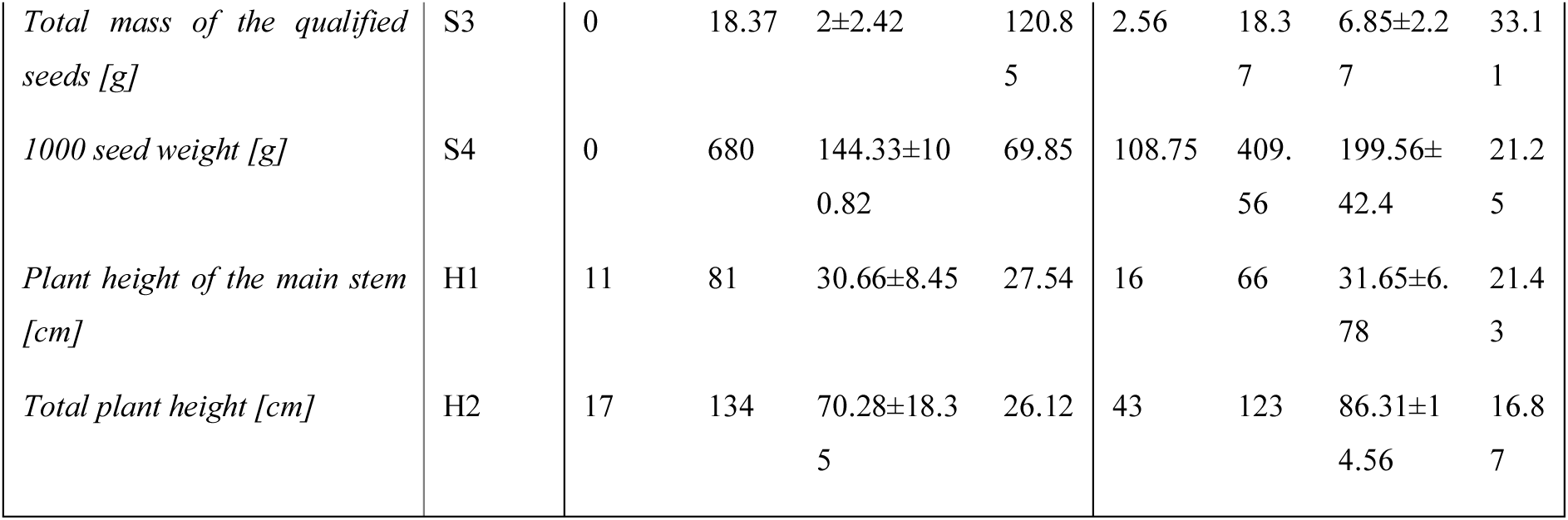
Statistical analysis of the 24 quantitative traits in the R-CORE and T-CORE collections (CV = coefficient of variation).

### 3.2 Development and characterization of the L. albus T-CORE collection

A set of 300 accessions from the *L. albus* R-CORE collection was used to build the T-CORE collection (Table S1). The T-CORE collection comprised 89 landraces (29.7%), 75 wild accessions (25%), 59 cultivars (17.3%) and 42 breeding/research materials (14%) from 30 countries representing eight geographical regions (Table 4, Figure 4A). Another 42 accessions (14%) lacked an assigned biological status and the geographical origin of 20 (6.66%) was unknown. Five accessions lacked both designations. Like the R-CORE, most of the T-CORE accessions (231, 77%) originated from Europe, mainly Southern Europe (173, 57.7%), followed by Western Europe (33, 10.3%) and Eastern Europe (25, 8.3%). The major contributors outside Europe were Northern Africa (21, 7.0%), the Middle East (17, 5.7%) and East Africa (7, 2.3%). America (3, 1%) and Australia and New Zealand (1, 0.3%) were minor contributors. Almost 50% of the T-CORE collection was represented by accessions from Southern Europe, mainly Spain (96, 32%) and Italy (40, 13.33%). The nonsignificant χ^2^ value for all geographical regions and countries indicated that all regions were optimally represented in the T-CORE collection (Table 3).

**Figure 4:**
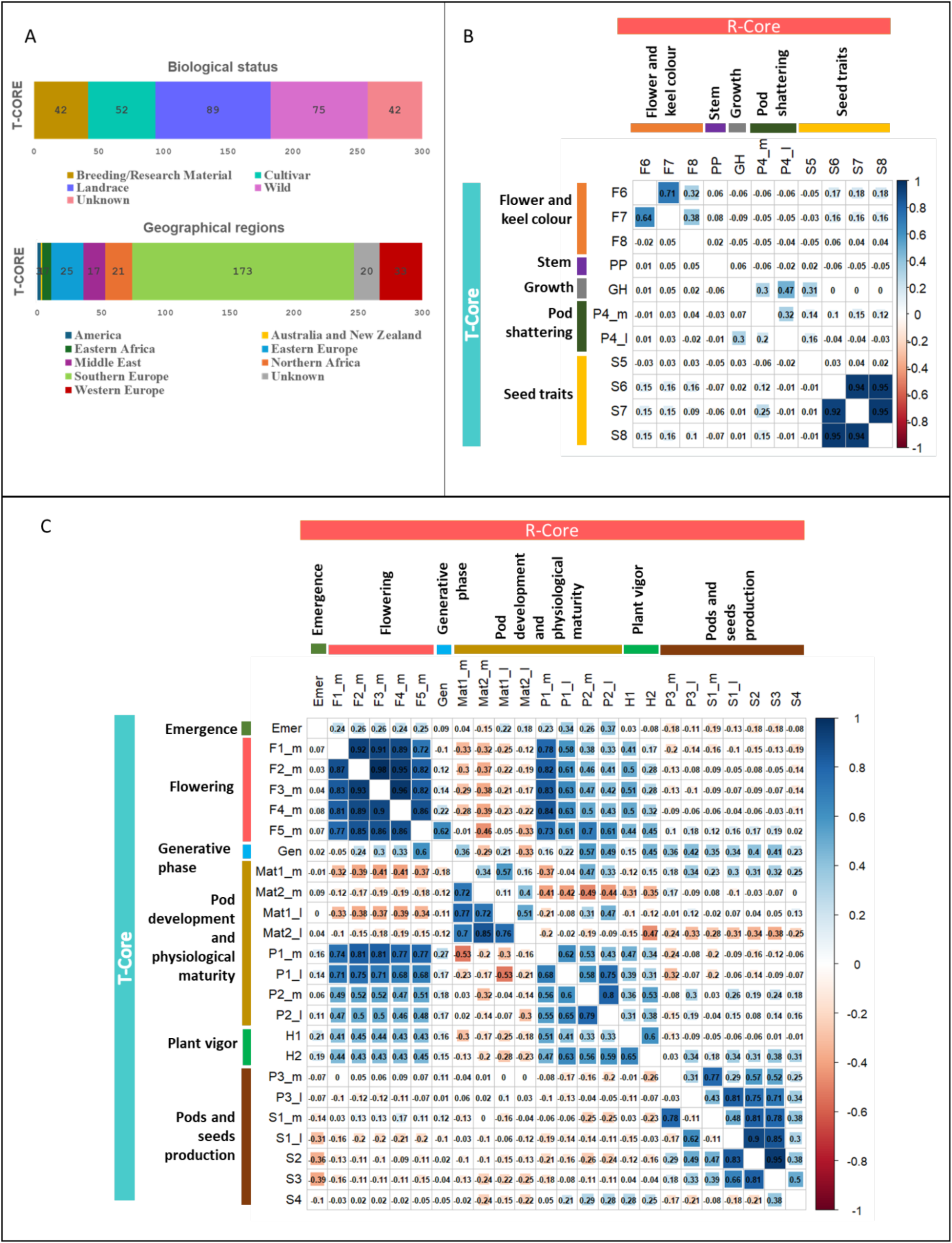
Characterization and comparative phenotypic structures of the *L. albus* R-CORE and T-CORE collections. (A) Biological status and geographical region. (B,C) Bivariate correlation plot of (B) 12 qualitative traits and (C) 24 quantitative traits, with the upper right panel representing the R-CORE and the lower left panel representing the T-CORE (red and blue colors indicate positive and negative correlations, respectively).

Qualitative trait characterization showed that pod shattering on the main stem and lateral branches, seed shape, seed ornamentation, and the color and density of seed ornamentation were homogenously distributed in the R-CORE and T-CORE collections (*p* > 0.05), with a CC of 66.66–100% and mean of 82.16% for all traits (Table 5). Wide variability was observed for most of the quantitative traits and they showed a non-normal distribution, similar to the R-CORE collection. Descriptive statistics of the quantitative traits in the T-CORE collection are presented in Table 6.

#### 3.2.1 Correlation analysis in the T-CORE and R-CORE collections

The correlation patterns for most traits were similar in the R-CORE and T-CORE (Figure 4). For the quantitative traits (Figure 4C, Table S3), we observed strong positive correlations (*r* > 0.70) among the flowering-related traits and relatively strong associations with traits related to pod development (*r* > 0.40). For example, days to first flower on the main stem showed strong positive correlations with days to flowering (length of vegetative phase) (*r* = 0.83) and days to first petal occurrence on the main stem (*r* = 0.93) among the flowering traits, and with days to first pod set on the main stem (*r* = 0.83) and lateral branches (*r* = 0.63) among traits related to pod development. We also observed significant positive correlations among the traits characterizing the physiological maturity on the main stem and lateral branches, but these traits showed significant negative correlations with traits related to flowering. In addition, in both collections, small negative correlations (*r* > –0.21) were observed between flowering-related traits and traits related to the production of pods and seeds. Importantly, the number of pods and qualified seeds on lateral branches showed a strong positive correlation with the total number of qualified seeds. For the qualitative traits, the strongest positive correlations were observed among traits related to seed ornamentation (*r* > 0.9) and between the color of wing petals and standard petals (*r* = 0.71 in the R-CORE and *r* = 0.64 in the T-CORE) (Figure 4B, Table S4). The growth habit showed a moderate positive correlation with pod shattering on the main stem and lateral branches (*r* > 0.30). Furthermore, we observed a small but significant positive correlation (*r* < 0.20) among the traits related to petal color, seed shape, seed ornamentation, and the density of ornamentation.

#### 3.2.2 Phenotypic variation in the T-CORE collection

The results described above confirm the wide variation of quantitative traits in the T-CORE collection, and a Shapiro–Wilk normality test showed that, except for plant height (H2; *p* = 0.198) and the number of qualified seeds on the main stem (S1_m; *p* = 0.096), all traits displayed non-normal distribution (Figure 5). For example, days to first open flower varied from 32 to 52 days, days to full flowering ranged from 50 to 74 days, and days to first pod set ranged from 51 to 75 days on the main stem and from 61 to 86 days on the lateral branches. For the traits related to pods and seeds, the number of pods ranged from 0 to 10 on the main stem and up to 15 on the lateral branches, the number of qualified seeds ranged from 0 to 34 on the main stem and up to 51 on the lateral branches, and the total number of qualified seeds ranged from 20 to 73 (Table S3). Interestingly, the wild accessions INLUP_00272 from Ukraine, INLUP_00131, INLUP_00135 and INLUP_00130 from Spain, INLUP_00233 from Turkey, INLUP_00136 from Portugal, and one landrace (INLUP_00201 from Poland) produced the largest number of seeds (> 62).

**Figure 5:**
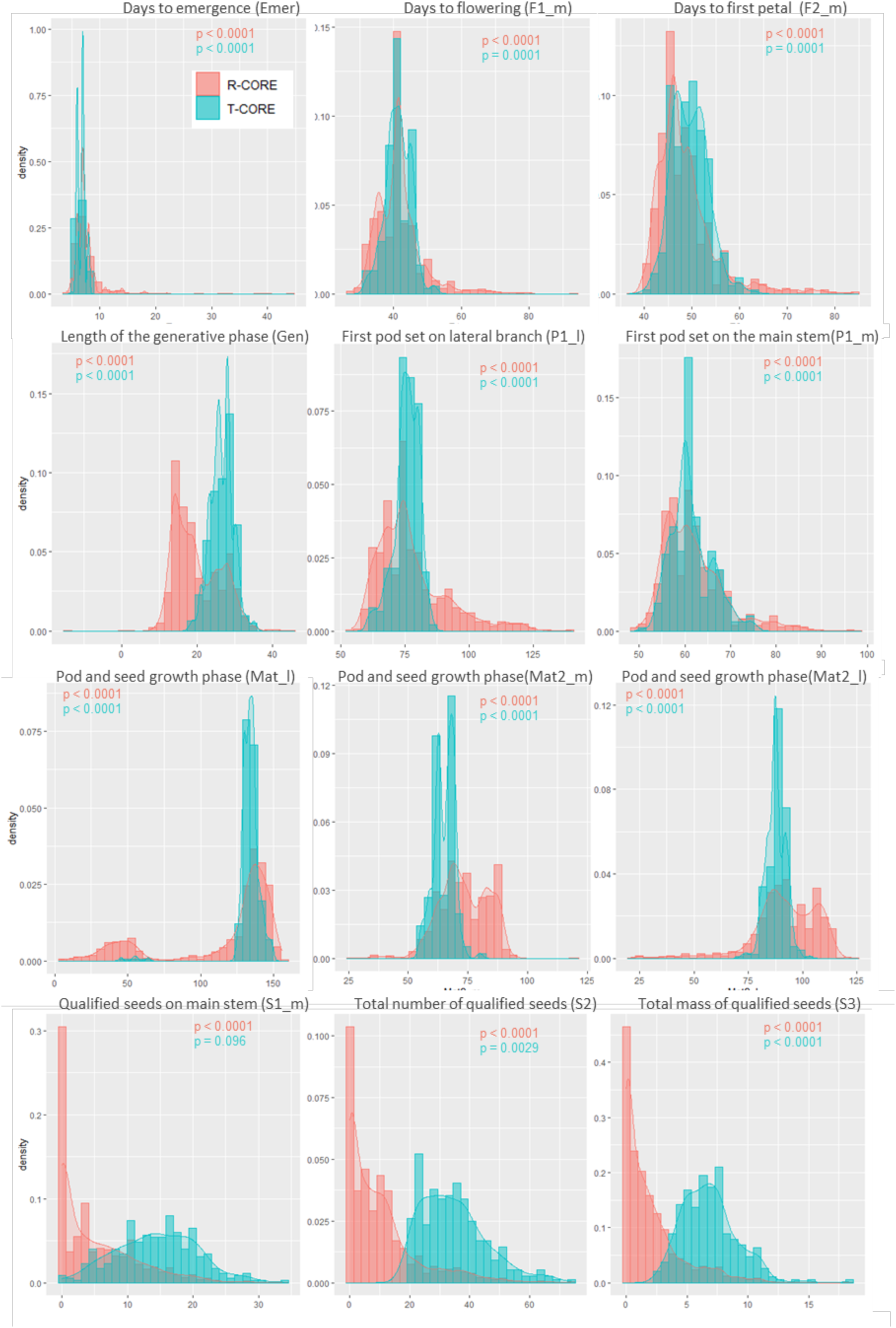
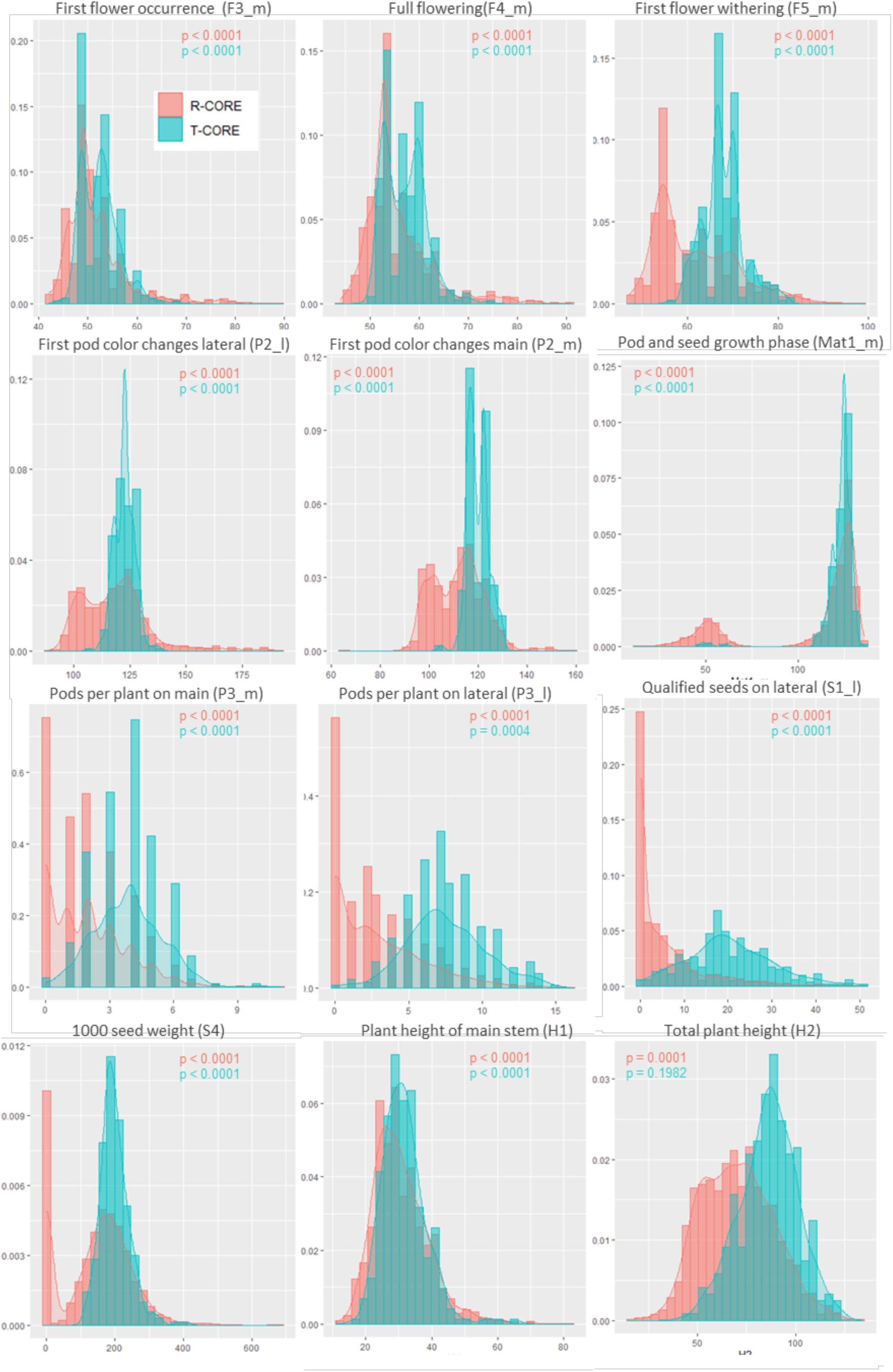
Comparative quantitative variation in the R-CORE (red) and T-CORE (green) collections.

The Shannon diversity index for qualitative traits was similar in the R-CORE and T-CORE collections (Table 5), although the mean value was slightly lower in the T-CORE collection (H′ = 0.17, compared to 0.21 for the R-CORE), indicating that the overall diversity was maintained in the T-CORE. When combined with the frequency distribution of qualitative traits (Table S2), these data show that the color of wing petal (F6; H′ = 0.78) was white/blue in 225 accessions, whereas the standard petal (F7) was white/blue in 178 accessions and white in 110 accessions (H′ = 0.60). The color of the keel edge (F8; H′ = 0.09) was transparent for 284 accessions. Moreover, the purple pigmentation on the main stem (PP; H′ = 0.70) was mainly characterized by slight pigmentation (108 accessions) and pigmentation at inter-nodes (103 accessions). In most T-CORE accessions, pod shattering was totally absent on the main stem (P4_m; 287 accessions) and lateral branches (P4_l; 295 accessions). In addition, the growth habit (GH; H′ = 0.30) was indeterminate in 296 accessions and semi-determinate in four. A determinate growth habit was not observed in any of the T-CORE accessions. Most accessions produced non-ornamented seeds (293 accessions) with flattened spherical (184 accessions) or flattened cuboid (103 accessions) shapes (Table S2).

### 3.3 Genetic diversity and phenotypic structure

MCA applied to the 11 qualitative traits revealed that the two first components explained 19.3% (12.3% and 7%, respectively) of the total variance and divided the R-CORE accessions into two groups: one with non-ornamented seeds (group A, 2237 accessions), and a smaller one with ornamented seeds (group B, 51 accessions) (Figure 6A,B, Table S5). Group A formed three subgroups designated A1, A2 and A3 (Figure 6C,D, Table S6). Group A1 (2032 accessions) mainly featured white or white/blue flowers, transparent keel, purple pigmentation on the stem, and indeterminate growth. It was made up of 176 wild accessions, 1047 landraces, 367 cultivars, 166 breeding/research materials and 276 accessions with unknown biological status, originating from all nine geographical regions (Figure 7A). Group A2 (180 accessions) mainly featured violet flowers, invisible keel, no purple pigmentation on the stem, non-shattering pods and semi-determinate growth habit. It was made up of 26 wild accessions, 114 landraces, 18 cultivars, 6 breeding/research materials and 16 accessions with unknown biological status, again originating from all nine geographical regions. The smallest subgroup (A3, 25 accessions) were mainly distinguished by shattering pods and white/blue flowers (Figure 6C,D, Table S6). It was made up of seven wild accessions, eight landraces, four cultivars, one breeding/research material, and five accessions with unknown biological status, and originated from Southern Europe (Italy and Spain), the Middle East (Palestine), Eastern Europe (Poland) and Northern Africa (Egypt) (Figure 7A,B). Group B (accessions with ornamented seeds) was mainly represented by wild accessions (24), with only four landraces, three cultivars and three breeding/research materials. There were 17 accessions of unknown biological status. The accessions from group B originated from Europe, mostly (33 accessions) from Greece, Southern Europe (Figure 7B, Table S7).

**Figure 6:**
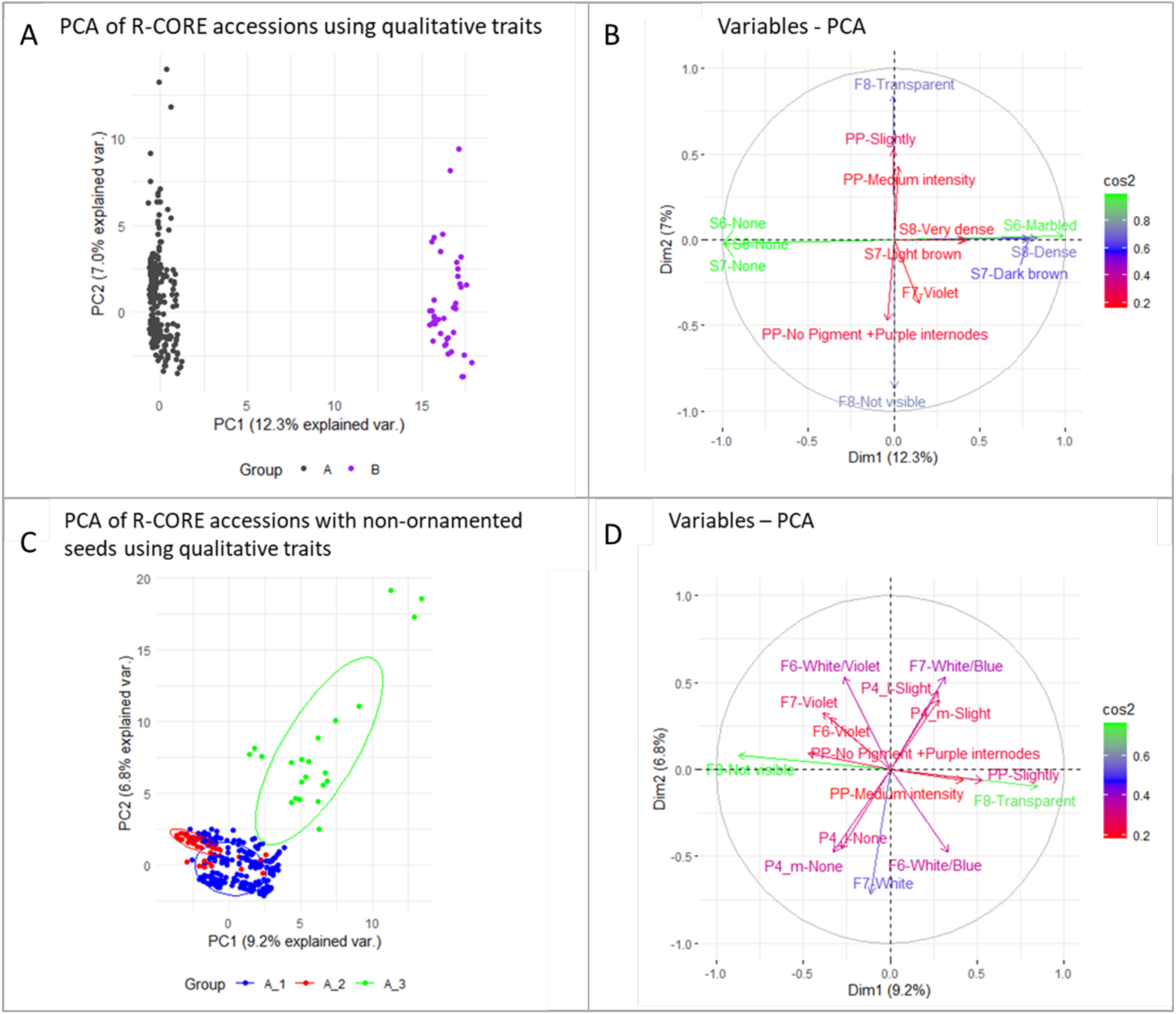
Principal component analysis of the *L. albus* R-CORE accessions. (A) Biplot of all R-CORE accessions. (B) Multiple correspondence analysis (MCA) of qualitative traits. (C) Biplot of R-CORE accessions with non-ornamented seeds. (D) MCA of qualitative traits from R-CORE accessions with non-ornamented seeds.

**Figure 7:**
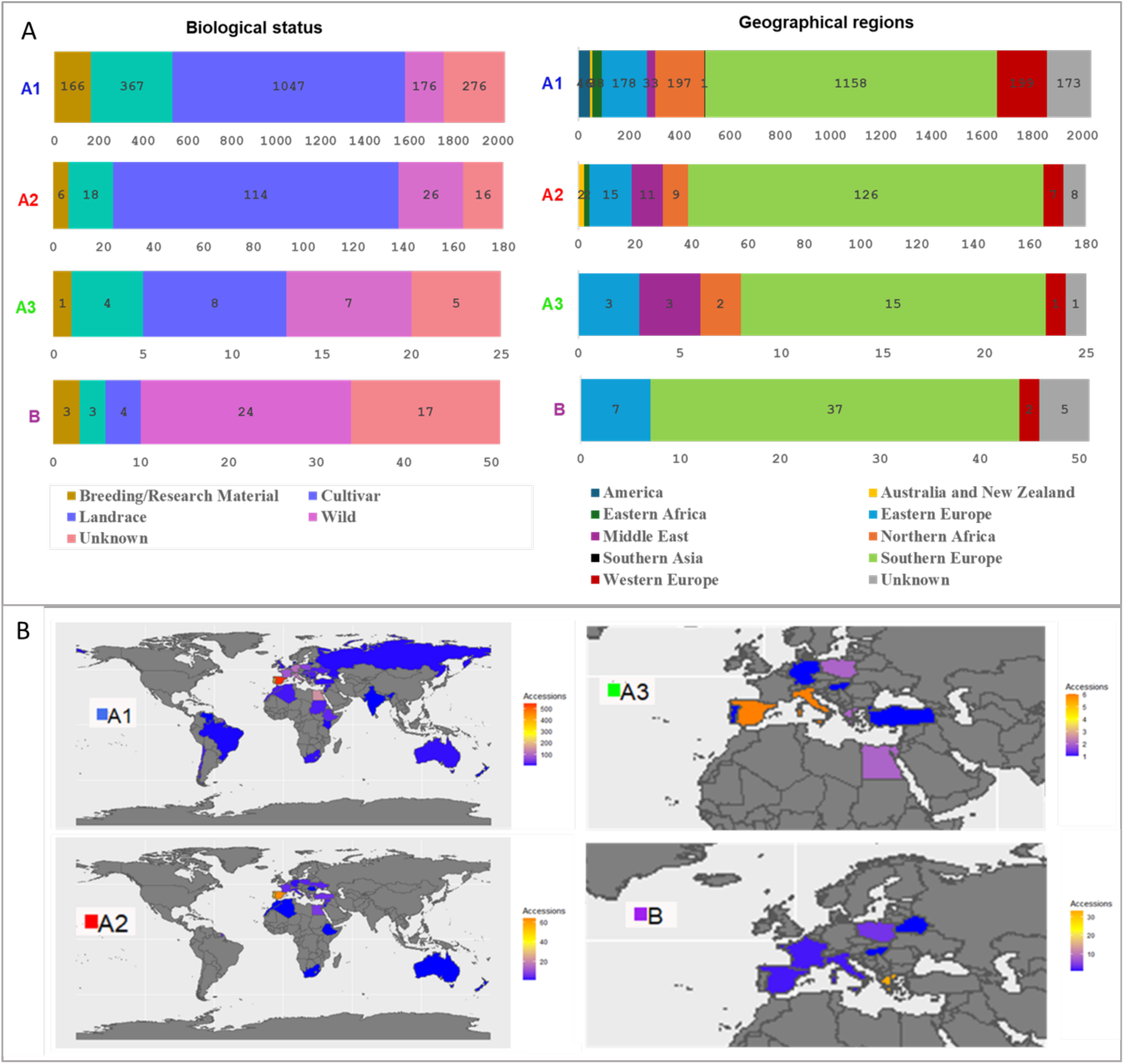
Clustering of the R-CORE accessions in groups A1, A2, A3 and B. (A) Biological status and geographical distribution. (B) Country of origin.

PCA and HCPC were also applied while excluding ornamentation traits and the breeding/research materials to maintain genetic integrity, and to avoid skewed results caused by selection processes (Table S8). PCA excluding ornamentation traits (S6–S8) clustered the accessions in three groups: I (190 accessions), II (2071 accessions), and III (27 accessions) (Figure S2). Cluster I consisted of accessions with violet, white/pink or pink flowers, not visible keel; semi-determinate growth and no pigmentation on the main stem. Cluster II consisted of accessions with white/blue, white or white/violet flowers, non-shattering pods, transparent keel, and purple pigmentation on main stem. Cluster III was differentiated mainly by accessions with shattering pods and a white/transparent keel (Table S8).

The exclusion of breeding/research materials also resulted in three clusters (I, II and III) (Table S9, Figure S3). Cluster I was differentiated by with not visible keel, no pigmentation on the main stem, white/violet, white or violet flowers, non-ornamented seeds, and semi-determinate growth. Cluster II consisted of accessions with transparent/white keel, purple pigmentation on the main stem, indeterminate growth, and (uniquely) shattering pods. Cluster III included the accessions with ornamented seeds, pink or white/pink flowers, and determinate growth. When the ornamentation traits were excluded, PCA again resulted in three groups (I, II and III) (Figure S4). Cluster I featured accessions with violet, white/pink or pink flowers not visible keel and semi-determinate growth. Cluster II featured accessions with white/blue, white or white/violet flowers, non-shattering pods, and a transparent keel. Cluster III was characterized by shattering pods and a white/transparent keel (Table S9).

PCA was also applied to the 24 quantitative traits to identify those differentiating between geographical regions, biological status and defined phenotypic groups. The two first principal components explained 38.3% and 24.5% of the total variance, respectively (Figure S5, Table S10). However, the biplot showed no distinct separation of R-CORE accessions based on phenotypic groups, geographical regions or biological status (Figure S5).

#### 3.3.1 Association of flowering and seed traits with phenotypic groups, geographical origin and biological status

Comparative ANOVA was applied to important domestication traits such as flowering time and seed characteristics, revealing that accessions in group A were early flowering in general, and group A3 showed significantly earlier flowering (40 days) than group B (47 days). Group A3 was also significantly more productive (∼25 qualified seeds) compared to groups A1, A2 and B (∼11, 8 and 11 seeds, respectively) (Figures S6, S7).

All accessions from the Middle East were early flowering and the most productive, with an average of 39 days to first bud occurrence (F6) and an average of 20 qualified seeds. In contrast, accessions from Australia and New Zealand, America and Southern Europe were late flowering (43 days) and the least productive, with an average of 7, 9 and 10 seeds, respectively. In terms of biological status, landraces and wild accessions were later flowering (42 days) than breeding materials and cultivars (∼40 days).

The ornamented accessions (group B) were characterized by a significantly lower mean 1000 seed weight (85.01 g) than group A with non-ornamented accessions (145.71 g). Among the latter, the 1000 seed weight was significantly higher in group A3 (pod-shattering accessions; 203.25 g) than groups A1 and A2 (145.16 and 136.33, respectively). The 1000 seed weight also differed significantly between geographical regions. The highest values were observed in accessions from Northern Africa (196.09 g) and the Middle East (171.95 g) and the lowest in accessions from Australia and New Zealand (86.96 g). There were no significant differences in 1000 seed weight between breeding/research materials (153.02 g), landraces (146.82 g) and cultivars (140.69 g).

In the wild accessions, we distinguished a group of 209 accessions with non-ornamented seeds (Wild_A) and a group of 24 accessions with ornamented seeds (Wild_B) (Figure 8A). Wild_A accessions originated from different regions, whereas Wild_B accessions were mainly from Greece. Notably, the Wild_B group was significantly later flowering (47 days) and produced fewer seeds (11) with a lower 1000 seed weight (85.01 g) than the Wild_A group, which flowered after 40 days and produced an average of 25 seeds with a 1000 seed weight of 185.27 g (Figure 8B).

**Figure 8:**
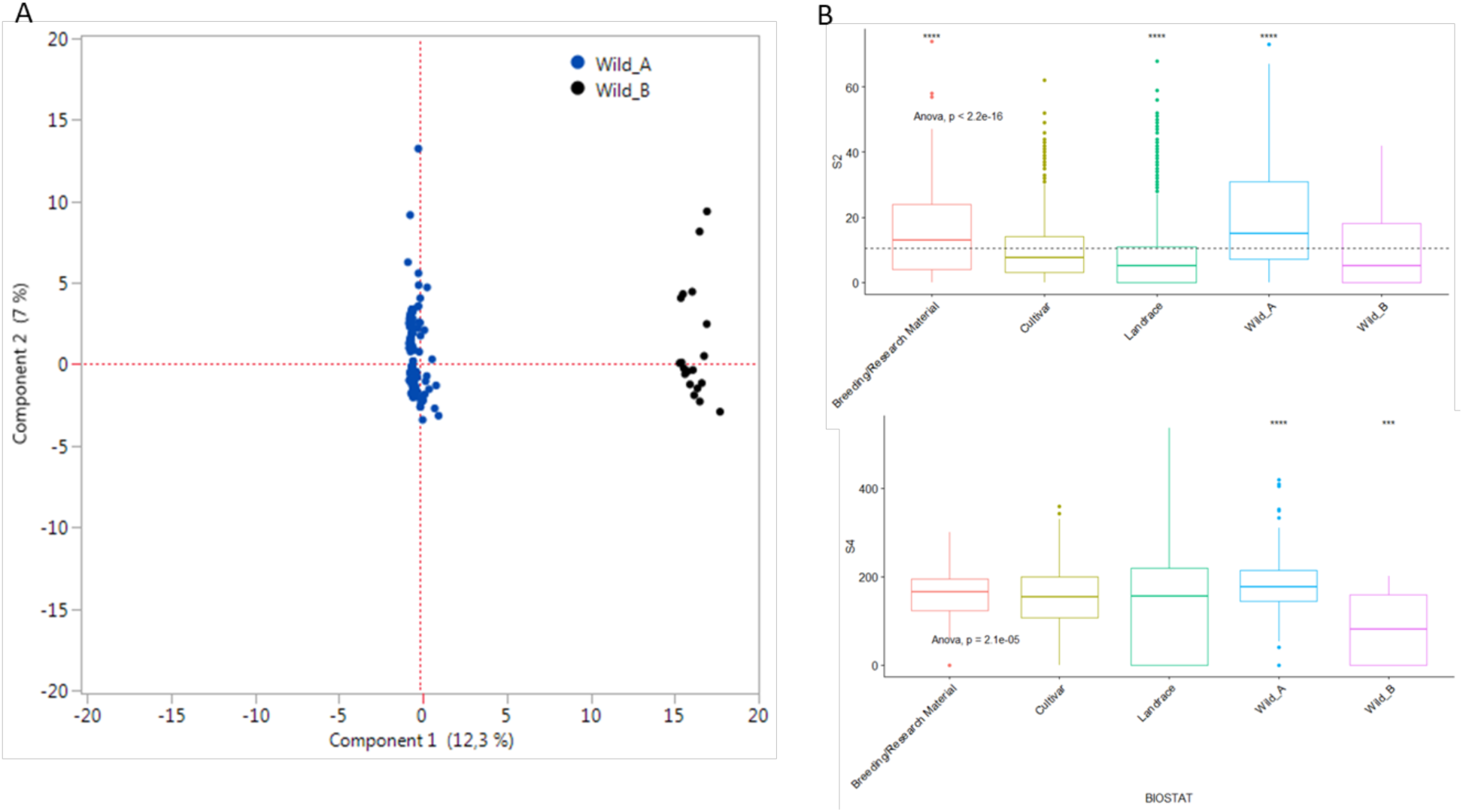
Analysis of wild accessions. (A) Phenotypic structure of wild accessions. (B) Box plots showing differences in the total number of qualified seeds (S2) and 1000 seed weight (S4) compared to accessions with a different biological status. Statistical significance was determined by one-way ANOVA followed by Tukey’s multiple comparisons test (*p ≤ 0.01).

#### 3.3.2 Patterns of additive genetic variation

The coefficient of additive genetic variation (CVa) was highly variable across traits, phenotypic groups, geographical regions, and biological statuses (Table 7, Table S11). Traits related to the production of pods and seeds, such as the number of qualified seeds on the main stem (S1_m), total mass (S3), the total number of qualified seeds (S2), 1000 seed weight (S4), and the number of pods per plant on the main stem (P3_m) had the highest CVa (> 30%) with an average of 34%, whereas the flowering traits had the lowest CVa with an average of 5%. Regarding the defined phenotypic groups, the mean CVa for all traits was highest in the phenotypic group of non-ornamented accessions (group A). Group A3 (with pod-shattering accessions) was the highest of all (CVa = 17%), followed by groups A1 (CVa = 15%) and A2 (CVa = 14%). In terms of biological status, the highest overall CVa was observed for breeding/research materials (19%) and Wild_A (18%), followed by landraces (14%), cultivars (13%) and Wild_B (10%). At the geographical level, the overall additive genetic variation was highest in Eastern Europe (CVa = 23%), Eastern Africa (CVa = 21%) and the Middle East (CVa = 21%), and lowest in Northern Africa and Western Europe (CVa = 12%) (Table S11).

**Table 7.**
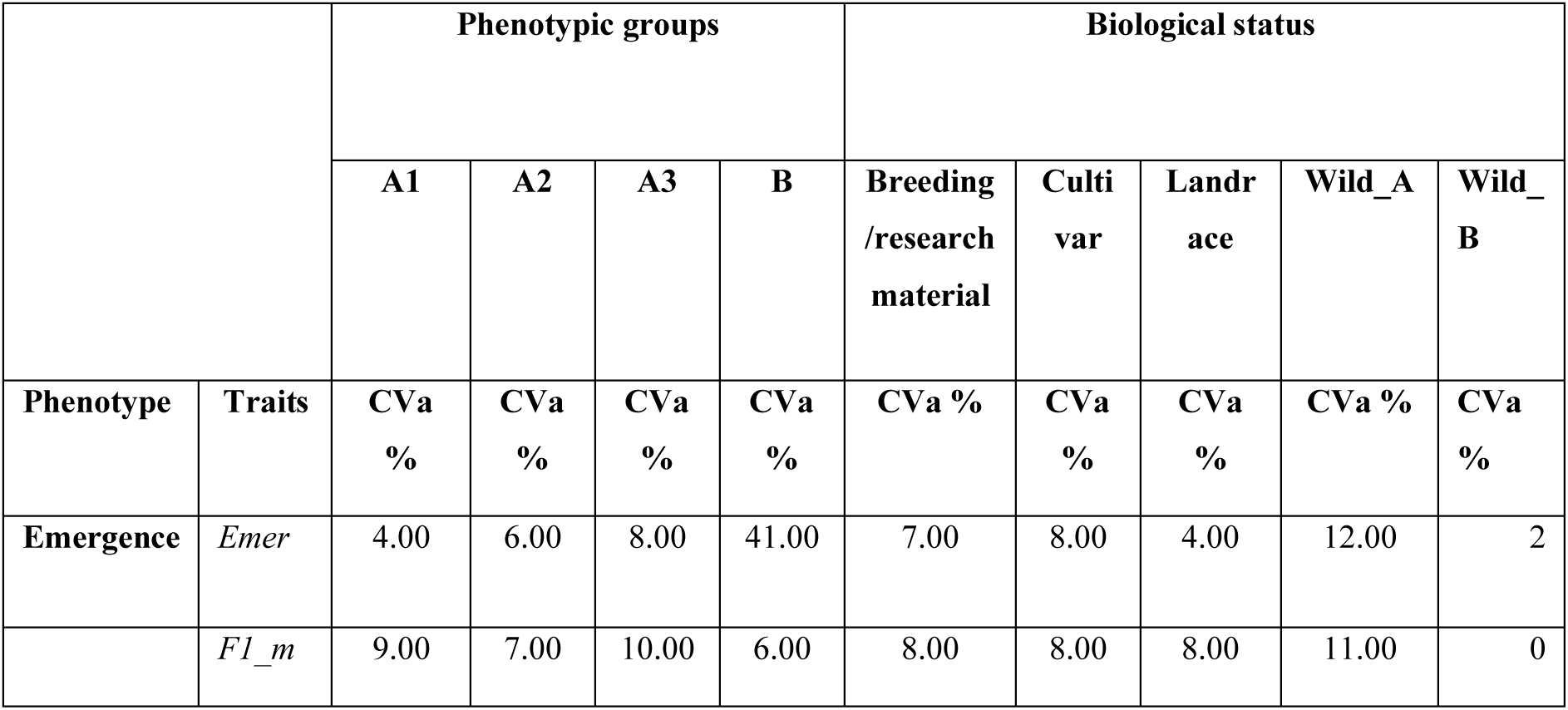
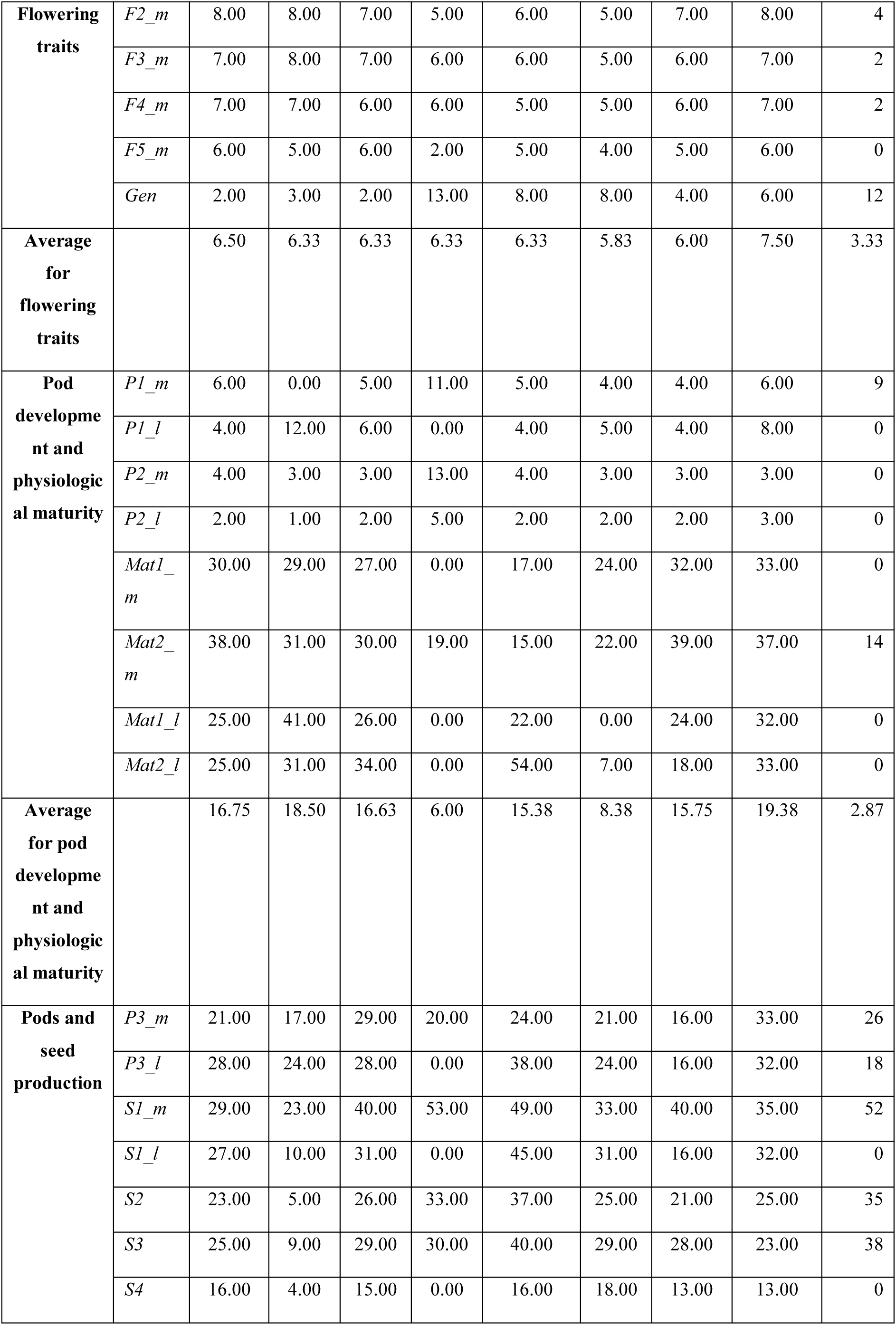
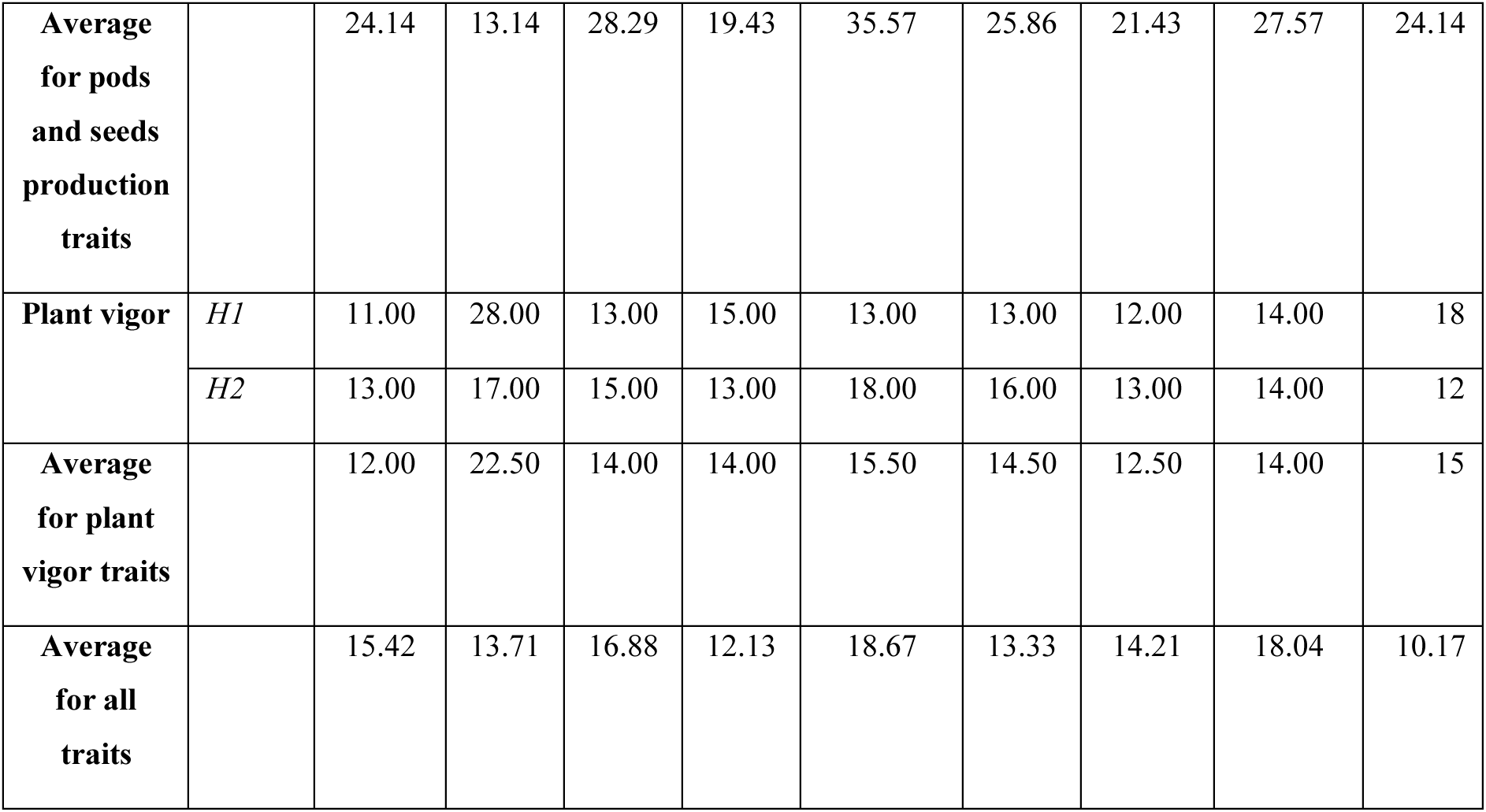
Patterns of additive genetic variation among the 24 quantitative traits in terms of phenotypic groups and the biological status of the accessions.

## 4. DISCUSSION

PGRs must be developed, characterized and maintained to ensure their availability for future crop improvement programs and to protect global agrobiodiversity. Here, for the first time, we have established an extensive collection of *L. albus* germplasm by applying the novel nested core collection approach described in the INCREASE project (Bellucci et al., 2021). Our R-CORE collection is the largest and best-characterized genetic resource for *L. albus* worldwide, and adds to the information from previous studies which relied on smaller collections mostly derived from Southern Europe, mainly Portugal (Vaz et al., 2004), the northwest Iberian plateau (Gonzalez-Andres et al., 2007), Greece (Zafeiriou et al., 2021), and Ethiopia (Beyene, 2020, Atnaf et al., 2017). Our R-CORE collection was developed using SSD lines (homogenous accessions), which facilitate reliable genotype–phenotype association, thus providing access to structural variants and insight into their functions. The conservation of these SSD collections reduces the likelihood of genetic changes during seed multiplication for heterogeneous accessions in gene banks (i.e., genetic drift and/or selection) (Bellucci et al., 2021). The R-CORE collection was developed and characterized using a standard protocol that eliminates data collection and integration bias, thus facilitating exploitation (Kroc et al., 2021). This allowed the more detailed characterization of *L. albus* compared to previous studies, in which a piecemeal approach was used to evaluate agro-morphological traits in germplasm collections (Beyene, 2020, Atnaf et al., 2017, Gonzalez-Andres et al., 2007, Vaz et al., 2004, Zafeiriou et al., 2021). The traits explored in this study can be used to characterize additional *L. albus* germplasm in the future.

The assessment of variability in desired traits is the first step in modern crop improvement strategies, facilitating the utilization of such variability in breeding programs. The characterization of the R-CORE, which represents global diversity, provided significant variations in many quantitative and qualitative agro-morphological traits. This was attributable to the diversity of the collection, including wild accessions, landraces, breeding materials and advanced cultivars with different geographical origins (Phogat et al., 2021). Our well-characterized R-CORE could therefore serve as ready-to-use material for further studies including next-generation omics research and future breeding programs.

The utilization and management of germplasm collections is enhanced by the availability of a limited number of genetically diverse accessions representing the diversity of the whole collection. We therefore generated the T-CORE collection as a single multipurpose resource with fewer accessions than the R-CORE but maximum diversity and representativeness (Bellucci et al., 2021, Kroc et al., 2021). The T-CORE contained ∼13% of the R-CORE accessions (2288) in accordance with the neutral allele theory (Brown, 1989). This approach has been developed in other legumes, including lentil (Simon and Hannan, 1995, Tullu et al., 2001), chickpea (Archak et al. 2016), and common bean (Rivera et al., 2018). Our T-CORE can therefore be used as a diversity hub, as evident from the relative distribution of sources and biological statuses of indigenous and exotic germplasm in the R-CORE and T-CORE collections, to meet the present and future challenges in *L. albus* breeding programs.

Following the development of a core collection, it is useful to assess the extent of variation retained from the reference collection (Mahalakshmi et al., 2007). It is also necessary to preserve trait associations to maintain co-adapted genetic complexes and to ensure the efficient utilization of germplasm while defining the core collection from a reference set (Ortiz et al., 1998, Archak et al., 2016, Tripathi et al., 2022). The variation in our T-CORE collection was representative of the R-CORE, confirmed by the conserved patterns of correlations between the collections for all combinations of quantitative and qualitative traits, as reported in previous studies involving crops such as lentil (Tripathi et al., 2022), chickpea (Archak et al., 2016), wheat (Phogat et al., 2021) and eggplant (Gangopadhyay et al., 2010). We assume that the pairwise correlation between traits related to flowering, pod development, physiological maturity and seed productivity reflects the presence of pleiotropic or linked genes. These findings may facilitate molecular research on pleiotropic effects in *L. albus,* as reported in other legumes (He et al., 2023), for example developmental genes influencing nitrogen source capacity, seed protein content and productivity in pea (Burstin et al., 2007). The correlated traits may be useful for the phenotypic characterization of future *L. albus* germplasm, but could hinder crop breeding when targeting traits with negative associations (He et al., 2023).

The morphological diversity of our R-CORE collection included variations in flower color, stem pigmentation, growth habit and seed traits. Interestingly, we observed considerable variation for seed ornamentation traits and accessions with ornamented seeds formed a distinct group (group B), which has not been identified before (Gonzalez-Andres et al., 2007). These accessions could be useful as a starting point for *L. albus* domestication studies because the loss of seed pigmentation is a sign of domestication syndrome (Bohra et al., 2022, Smýkal and Parker, 2023). Seed coat pigmentation also helps plants adapt to changing environments, and can influence root colonization by rhizobia and plant-growth-promoting microbes (reviewed by Jaiswal and Dakora 2024). The group B accessions identified in our study could therefore be useful in different agroecosystems, contributing significantly to overall soil health through symbiotic relationships with *Rhizobium* spp. Furthermore, the subgroup of wild accessions in group B originating from Greece were characterized by fewer seeds and low seed weight, which is typical for wild accessions with adaptations for efficient seed dispersal in diverse habitats. These accessions might be useful for further investigations to confirm that Greece was the origin of *L. albus*.

Seed weight is an important agronomic trait that directly affects crop yield and quality, and also influences the germination, growth and survival of seedlings (Törő-Szijgyártó et al., 2023, Finch-Savage and Bassel, 2016). The mean seed weight of cultivated varieties was higher than that of wild accessions in our collections, but we observed a considerable overlap, as previously reported for *L. albus* and *L. angustifolius* (Berger et al., 2017). Notably, some wild accessions in group A3 with indeterminate growth were characterized by a high seed weight and also shattering pods. Indeterminacy, however undesirable, is a common feature of wild *L. albus* populations (Huyghe, 1997), and indeterminate growth can produce more pods and seeds on lateral branches, as suggested for soybean (Kato et al., 2019). We found that lateral branches were more productive than the main stem in *L. albus* accessions. Breeders usually prefer determinate accessions, which are more suitable for mechanical harvesting, yet the indeterminate accessions could be useful for targeting specific traits such as crop yield. Therefore, group A3 accessions may be of interest for studies on the origin and domestication of *L. albus*, and represent untapped wild material that could be exploited to increase yields. In grain legumes, the presence of non-shattering pods is a primary trait associated with domestication syndrome (Maity et al., 2021, Kroc et al., 2021, Bohra et al., 2022, Smýkal and Parker, 2023, Rau et al., 2019, Murgia et al., 2017, Di Vittori et al., 2019).

With regard to quantitative traits, the *L. albus* accessions did not cluster into defined groups or according to their passport data, suggesting a weak population structure, which is consistent with the earlier reports on *L. albus* (Iqbal et al. 2012; Atnaf et al. 2015; Hufnagel et al. 2021; Alkemade et al. 2022) and other lupin species (Mousavi-Derazmahalleh et al. 2018) (Turner et al. 2018). This is likely to reflect the slow and sporadic domestication of *L. albus* (Hufnagel et al. 2020; Alkemade et al. 2022).

Early flowering, rapid maturation and high productivity are important traits for crop breeding. Ethiopian accessions might be valuable more generally because the mutual utilization of genomic data from major crops, orphan crops and their wild relatives has gained attention recently for the genetic improvement of both orphan and major crops (Ye and Fan, 2021). Wild accessions and landraces of *L. albus* thus offer a valuable resource for breeding programs (Adhikari et al., 2011, Zafeiriou et al., 2021). We found that some wild accessions and landraces were early flowering and more productive, indicating the presence of a gene reservoir suitable for breeding programs. Notably, accessions from Eastern Africa, particularly from Ethiopia, tended to be early flowering and the most productive. Our findings, based on a large number of accessions, therefore confirm the results from earlier studies using smaller collections showing that Ethiopian accessions provide a unique and important gene pool (Atnaf et al., 2017, Beyene, 2020).

Low additive variation was observed for flowering traits, so we assume that many beneficial alleles for flowering traits became fixed during domestication (Adhikari et al., 2011). This would deplete the additive genetic variation available during selection, as discussed for maize (Yang et al. (2019), but further research is needed to test this assumption. In contrast, moderate to high additive genetic variation was observed for traits related to physiological maturity and seed production, indicating these traits remain under substantial genetic control and may respond well to selection (Wisser et al., 2019). This highlights their relevance for domestication in the context of climate change (Hernández-Máximo et al., 2021).

In general, favorable environments are associated with weak selection that increases additive genetic variation (Etterson and Shaw, 2001). We observed this pattern in accessions from Eastern Europe, Eastern Africa and the Middle East, indicating that these regions may be the most favorable for growing *L. albus*. Notably, landraces showed the least additive genetic variation, possibly indicating that the environmental stress caused by climate change depletes the additive genetic variation in landraces, which may affect their evolutionary potential (Mercer and Perales, 2010). Overall, our work confirms the presence of substantial additive genetic variance, and further studies should aim to isolate the alleles responsible for trait variation to better understand the genetic basis of the morphological changes.

This study is the first step toward the development and characterization of an Intelligent Core Collection to conserve the agrobiodiversity of *L. albus* genetic resources for food security. The T-CORE collection will provide insight into the molecular basis of many important quantitative and qualitative traits, such as early flowering and seed production, as well as flower color and seed morphology (shape, size and ornamentation). It could be used for advanced research on the evolution (Garcia et al., 2021, Cortinovis et al., 2024), genomic selection (Norman et al., 2018, Hu et al., 2022) and the genetic architecture of important agronomic traits (Roorkiwal et al., 2018, Weckwerth et al., 2020) such as grain yield (Pang et al., 2020). Importantly, the T- CORE collection could provide genetic material for molecular characterization to identify genes controlling important agronomic traits, track lupin genome evolution and domestication, and allow the further genomic characterization of *L. albus* germplasm for crop improvement programs.

## Supporting information

L.albus-manuscript_SupplementaryTables

L.albus-manuscript_SupplementaryFigures

## ACKNOWLEDGEMENTS

This study was conducted within the INCREASE project, as funded by the European Union Horizon 2020 Research and Innovation Programme under grant agreement No 862862.

## DATA AVAILABILITY

Raw data used in the present study can be obtained from the corresponding author upon reasonable request.

