## Supplementary material for "Genetic and phenotypic characterization of global *Lupinus albus* genetic resources for the development of a CORE collection": L.albus-manuscript_SupplementaryTables

**Table S1: Details of the *L. albus* accessions used in this st**

| INCREASE ID | R-CORE | T-CORE |
|---|---|---|
| INLUP_00001 | R-CORE | |
| INLUP_00002 | R-CORE | |
| INLUP_00003 | R-CORE | |
| INLUP_00004 | R-CORE | |
| INLUP_00005 | R-CORE | |
| INLUP_00006 | R-CORE | |
| INLUP_00007 | R-CORE | |
| INLUP_00008 | R-CORE | |
| INLUP_00009 | R-CORE | |
| INLUP_00010 | R-CORE | |
| INLUP_00011 | R-CORE | |
| INLUP_00012 | R-CORE | |
| INLUP_00013 | R-CORE | |
| INLUP_00014 | R-CORE | |
| INLUP_00015 | R-CORE | |
| INLUP_00016 | R-CORE | |
| INLUP_00017 | R-CORE | |
| INLUP_00018 | R-CORE | |
| INLUP_00019 | R-CORE | |
| INLUP_00020 | R-CORE | |
| INLUP_00021 | R-CORE | |
| INLUP_00022 | R-CORE | |
| INLUP_00023 | R-CORE | |
| INLUP_00024 | R-CORE | |
| INLUP_00025 | R-CORE | |
| INLUP_00026 | R-CORE | |
| INLUP_00027 | R-CORE | |
| INLUP_00028 | R-CORE | |
| INLUP_00029 | R-CORE | |
| INLUP_00030 | R-CORE | |
| INLUP_00031 | R-CORE | |
| INLUP_00032 | R-CORE | |
| INLUP_00033 | R-CORE | |
| INLUP_00034 | R-CORE | |
| INLUP_00035 | R-CORE | |
| INLUP_00036 | R-CORE | |
| INLUP_00037 | R-CORE | |
| INLUP_00038 | R-CORE | |
| INLUP_00039 | R-CORE | |
| INLUP_00040 | R-CORE | |
| INLUP_00041 | R-CORE | |
| INLUP_00042 | R-CORE | |
| INLUP_00043 | R-CORE | |
| INLUP_00044 | R-CORE | |
| INLUP_00045 | R-CORE | |
| INLUP_00046 | R-CORE | |
| INLUP_00047 | R-CORE | |
| INLUP_00048 | R-CORE | |
| INLUP_00049 | R-CORE | |
| INLUP_00050 | R-CORE | |

| | |
|---|---|
| INLUP_00051 | R-CORE |
| INLUP_00052 | R-CORE |
| INLUP_00053 | R-CORE |
| INLUP_00054 | R-CORE |
| INLUP_00055 | R-CORE |
| INLUP_00056 | R-CORE |
| INLUP_00057 | R-CORE |
| INLUP_00058 | R-CORE |
| INLUP_00059 | R-CORE |
| INLUP_00060 | R-CORE |
| INLUP_00061 | R-CORE |
| INLUP_00062 | R-CORE |
| INLUP_00063 | R-CORE |
| INLUP_00064 | R-CORE |
| INLUP_00065 | R-CORE |
| INLUP_00066 | R-CORE |
| INLUP_00067 | R-CORE |
| INLUP_00068 | R-CORE |
| INLUP_00069 | R-CORE |
| INLUP_00070 | R-CORE |
| INLUP_00071 | R-CORE |
| INLUP_00072 | R-CORE |
| INLUP_00073 | R-CORE |
| INLUP_00074 | R-CORE |
| INLUP_00075 | R-CORE |
| INLUP_00076 | R-CORE |
| INLUP_00077 | R-CORE |
| INLUP_00078 | R-CORE |
| INLUP_00079 | R-CORE |
| INLUP_00080 | R-CORE |
| INLUP_00081 | R-CORE |
| INLUP_00082 | R-CORE |
| INLUP_00083 | R-CORE |
| INLUP_00084 | R-CORE |
| INLUP_00085 | R-CORE |
| INLUP_00086 | R-CORE |
| INLUP_00087 | R-CORE |
| INLUP_00088 | R-CORE |
| INLUP_00089 | R-CORE |
| INLUP_00090 | R-CORE |
| INLUP_00091 | R-CORE |
| INLUP_00092 | R-CORE |
| INLUP_00093 | R-CORE |
| INLUP_00094 | R-CORE |
| INLUP_00095 | R-CORE |
| INLUP_00096 | R-CORE |
| INLUP_00097 | R-CORE |
| INLUP_00098 | R-CORE |
| INLUP_00099 | R-CORE |
| INLUP_00100 | R-CORE |
| INLUP_00101 | R-CORE |
| INLUP_00102 | R-CORE |

| | | |
|---|---|---|
| INLUP_00103 | R-CORE | |
| INLUP_00104 | R-CORE | |
| INLUP_00105 | R-CORE | |
| INLUP_00106 | R-CORE | |
| INLUP_00107 | R-CORE | |
| INLUP_00108 | R-CORE | |
| INLUP_00109 | R-CORE | |
| INLUP_00110 | R-CORE | |
| INLUP_00111 | R-CORE | |
| INLUP_00112 | R-CORE | |
| INLUP_00113 | R-CORE | |
| INLUP_00114 | R-CORE | |
| INLUP_00115 | R-CORE | |
| INLUP_00116 | R-CORE | |
| INLUP_00117 | R-CORE | |
| INLUP_00118 | R-CORE | |
| INLUP_00119 | R-CORE | |
| INLUP_00120 | R-CORE | |
| INLUP_00121 | R-CORE | |
| INLUP_00122 | R-CORE | |
| INLUP_00123 | R-CORE | |
| INLUP_00124 | R-CORE | |
| INLUP_00125 | R-CORE | |
| INLUP_00126 | R-CORE | |
| INLUP_00127 | R-CORE | |
| INLUP_00128 | R-CORE | |
| INLUP_00129 | R-CORE | |
| INLUP_00130 | R-CORE | T-CORE |
| INLUP_00131 | R-CORE | T-CORE |
| INLUP_00132 | R-CORE | T-CORE |
| INLUP_00133 | R-CORE | T-CORE |
| INLUP_00134 | R-CORE | T-CORE |
| INLUP_00135 | R-CORE | T-CORE |
| INLUP_00136 | R-CORE | T-CORE |
| INLUP_00137 | R-CORE | T-CORE |
| INLUP_00138 | R-CORE | T-CORE |
| INLUP_00139 | R-CORE | T-CORE |
| INLUP_00140 | R-CORE | T-CORE |
| INLUP_00141 | R-CORE | T-CORE |
| INLUP_00142 | R-CORE | T-CORE |
| INLUP_00143 | R-CORE | T-CORE |
| INLUP_00144 | R-CORE | T-CORE |
| INLUP_00145 | R-CORE | T-CORE |
| INLUP_00146 | R-CORE | T-CORE |
| INLUP_00147 | R-CORE | T-CORE |
| INLUP_00148 | R-CORE | |
| INLUP_00149 | R-CORE | |
| INLUP_00150 | R-CORE | |
| INLUP_00151 | R-CORE | |
| INLUP_00152 | R-CORE | T-CORE |
| INLUP_00153 | R-CORE | T-CORE |
| INLUP_00154 | R-CORE | |

| | | |
|---|---|---|
| INLUP_00155 | R-CORE | T-CORE |
| INLUP_00156 | R-CORE | T-CORE |
| INLUP_00157 | R-CORE | T-CORE |
| INLUP_00158 | R-CORE | T-CORE |
| INLUP_00159 | R-CORE | T-CORE |
| INLUP_00160 | R-CORE | T-CORE |
| INLUP_00161 | R-CORE | T-CORE |
| INLUP_00162 | R-CORE | T-CORE |
| INLUP_00163 | R-CORE | |
| INLUP_00164 | R-CORE | T-CORE |
| INLUP_00165 | R-CORE | T-CORE |
| INLUP_00166 | R-CORE | T-CORE |
| INLUP_00167 | R-CORE | T-CORE |
| INLUP_00168 | R-CORE | |
| INLUP_00169 | R-CORE | T-CORE |
| INLUP_00170 | R-CORE | T-CORE |
| INLUP_00171 | R-CORE | T-CORE |
| INLUP_00172 | R-CORE | T-CORE |
| INLUP_00173 | R-CORE | T-CORE |
| INLUP_00174 | R-CORE | T-CORE |
| INLUP_00175 | R-CORE | |
| INLUP_00176 | R-CORE | |
| INLUP_00177 | R-CORE | T-CORE |
| INLUP_00178 | R-CORE | T-CORE |
| INLUP_00179 | R-CORE | T-CORE |
| INLUP_00180 | R-CORE | T-CORE |
| INLUP_00181 | R-CORE | |
| INLUP_00182 | R-CORE | T-CORE |
| INLUP_00183 | R-CORE | |
| INLUP_00184 | R-CORE | T-CORE |
| INLUP_00185 | R-CORE | T-CORE |
| INLUP_00186 | R-CORE | |
| INLUP_00187 | R-CORE | T-CORE |
| INLUP_00188 | R-CORE | T-CORE |
| INLUP_00189 | R-CORE | T-CORE |
| INLUP_00190 | R-CORE | T-CORE |
| INLUP_00191 | R-CORE | T-CORE |
| INLUP_00192 | R-CORE | T-CORE |
| INLUP_00193 | R-CORE | T-CORE |
| INLUP_00194 | R-CORE | T-CORE |
| INLUP_00195 | R-CORE | T-CORE |
| INLUP_00196 | R-CORE | T-CORE |
| INLUP_00197 | R-CORE | |
| INLUP_00198 | R-CORE | T-CORE |
| INLUP_00199 | R-CORE | T-CORE |
| INLUP_00200 | R-CORE | T-CORE |
| INLUP_00201 | R-CORE | T-CORE |
| INLUP_00202 | R-CORE | |
| INLUP_00203 | R-CORE | |
| INLUP_00204 | R-CORE | |
| INLUP_00205 | R-CORE | |
| INLUP_00206 | R-CORE | T-CORE |

| | | |
|---|---|---|
| INLUP_00207 | R-CORE | T-CORE |
| INLUP_00208 | R-CORE | T-CORE |
| INLUP_00209 | R-CORE | T-CORE |
| INLUP_00210 | R-CORE | T-CORE |
| INLUP_00211 | R-CORE | T-CORE |
| INLUP_00212 | R-CORE | T-CORE |
| INLUP_00213 | R-CORE | T-CORE |
| INLUP_00214 | R-CORE | T-CORE |
| INLUP_00215 | R-CORE | T-CORE |
| INLUP_00216 | R-CORE | T-CORE |
| INLUP_00217 | R-CORE | T-CORE |
| INLUP_00218 | R-CORE | T-CORE |
| INLUP_00219 | R-CORE | T-CORE |
| INLUP_00220 | R-CORE | T-CORE |
| INLUP_00221 | R-CORE | |
| INLUP_00222 | R-CORE | |
| INLUP_00223 | R-CORE | T-CORE |
| INLUP_00224 | R-CORE | T-CORE |
| INLUP_00225 | R-CORE | T-CORE |
| INLUP_00226 | R-CORE | |
| INLUP_00227 | R-CORE | T-CORE |
| INLUP_00228 | R-CORE | T-CORE |
| INLUP_00229 | R-CORE | T-CORE |
| INLUP_00230 | R-CORE | T-CORE |
| INLUP_00231 | R-CORE | T-CORE |
| INLUP_00232 | R-CORE | T-CORE |
| INLUP_00233 | R-CORE | T-CORE |
| INLUP_00234 | R-CORE | T-CORE |
| INLUP_00235 | R-CORE | T-CORE |
| INLUP_00236 | R-CORE | T-CORE |
| INLUP_00237 | R-CORE | |
| INLUP_00238 | R-CORE | T-CORE |
| INLUP_00239 | R-CORE | T-CORE |
| INLUP_00240 | R-CORE | T-CORE |
| INLUP_00241 | R-CORE | |
| INLUP_00242 | R-CORE | T-CORE |
| INLUP_00243 | R-CORE | T-CORE |
| INLUP_00244 | R-CORE | T-CORE |
| INLUP_00245 | R-CORE | T-CORE |
| INLUP_00246 | R-CORE | T-CORE |
| INLUP_00247 | R-CORE | |
| INLUP_00248 | R-CORE | T-CORE |
| INLUP_00249 | R-CORE | |
| INLUP_00250 | R-CORE | T-CORE |
| INLUP_00251 | R-CORE | |
| INLUP_00252 | R-CORE | T-CORE |
| INLUP_00253 | R-CORE | T-CORE |
| INLUP_00254 | R-CORE | T-CORE |
| INLUP_00255 | R-CORE | T-CORE |
| INLUP_00256 | R-CORE | |
| INLUP_00257 | R-CORE | |
| INLUP_00258 | R-CORE | T-CORE |

| | | |
|---|---|---|
| INLUP_00259 | R-CORE | T-CORE |
| INLUP_00260 | R-CORE | |
| INLUP_00261 | R-CORE | |
| INLUP_00262 | R-CORE | T-CORE |
| INLUP_00263 | R-CORE | |
| INLUP_00264 | R-CORE | T-CORE |
| INLUP_00265 | R-CORE | |
| INLUP_00266 | R-CORE | T-CORE |
| INLUP_00267 | R-CORE | |
| INLUP_00268 | R-CORE | T-CORE |
| INLUP_00269 | R-CORE | |
| INLUP_00270 | R-CORE | |
| INLUP_00271 | R-CORE | T-CORE |
| INLUP_00272 | R-CORE | T-CORE |
| INLUP_00273 | R-CORE | T-CORE |
| INLUP_00274 | R-CORE | T-CORE |
| INLUP_00275 | R-CORE | T-CORE |
| INLUP_00276 | R-CORE | T-CORE |
| INLUP_00277 | R-CORE | T-CORE |
| INLUP_00278 | R-CORE | |
| INLUP_00279 | R-CORE | T-CORE |
| INLUP_00280 | R-CORE | T-CORE |
| INLUP_00281 | R-CORE | T-CORE |
| INLUP_00282 | R-CORE | |
| INLUP_00283 | R-CORE | |
| INLUP_00284 | R-CORE | T-CORE |
| INLUP_00285 | R-CORE | T-CORE |
| INLUP_00286 | R-CORE | T-CORE |
| INLUP_00287 | R-CORE | T-CORE |
| INLUP_00288 | R-CORE | T-CORE |
| INLUP_00289 | R-CORE | T-CORE |
| INLUP_00290 | R-CORE | |
| INLUP_00291 | R-CORE | T-CORE |
| INLUP_00292 | R-CORE | |
| INLUP_00293 | R-CORE | T-CORE |
| INLUP_00294 | R-CORE | |
| INLUP_00295 | R-CORE | T-CORE |
| INLUP_00296 | R-CORE | |
| INLUP_00297 | R-CORE | T-CORE |
| INLUP_00298 | R-CORE | T-CORE |
| INLUP_00299 | R-CORE | T-CORE |
| INLUP_00300 | R-CORE | |
| INLUP_00301 | R-CORE | |
| INLUP_00302 | R-CORE | T-CORE |
| INLUP_00303 | R-CORE | T-CORE |
| INLUP_00304 | R-CORE | T-CORE |
| INLUP_00305 | R-CORE | T-CORE |
| INLUP_00306 | R-CORE | T-CORE |
| INLUP_00307 | R-CORE | T-CORE |
| INLUP_00308 | R-CORE | T-CORE |
| INLUP_00309 | R-CORE | T-CORE |
| INLUP_00310 | R-CORE | T-CORE |

| | | |
|---|---|---|
| INLUP_00311 | R-CORE | T-CORE |
| INLUP_00312 | R-CORE | T-CORE |
| INLUP_00313 | R-CORE | T-CORE |
| INLUP_00314 | R-CORE | T-CORE |
| INLUP_00315 | R-CORE | T-CORE |
| INLUP_00316 | R-CORE | |
| INLUP_00317 | R-CORE | T-CORE |
| INLUP_00318 | R-CORE | T-CORE |
| INLUP_00319 | R-CORE | T-CORE |
| INLUP_00320 | R-CORE | T-CORE |
| INLUP_00321 | R-CORE | |
| INLUP_00322 | R-CORE | |
| INLUP_00323 | R-CORE | T-CORE |
| INLUP_00324 | R-CORE | T-CORE |
| INLUP_00325 | R-CORE | T-CORE |
| INLUP_00326 | R-CORE | |
| INLUP_00327 | R-CORE | T-CORE |
| INLUP_00328 | R-CORE | T-CORE |
| INLUP_00329 | R-CORE | T-CORE |
| INLUP_00330 | R-CORE | T-CORE |
| INLUP_00331 | R-CORE | T-CORE |
| INLUP_00332 | R-CORE | T-CORE |
| INLUP_00333 | R-CORE | T-CORE |
| INLUP_00334 | R-CORE | T-CORE |
| INLUP_00335 | R-CORE | T-CORE |
| INLUP_00336 | R-CORE | T-CORE |
| INLUP_00337 | R-CORE | |
| INLUP_00338 | R-CORE | T-CORE |
| INLUP_00339 | R-CORE | |
| INLUP_00340 | R-CORE | T-CORE |
| INLUP_00341 | R-CORE | |
| INLUP_00342 | R-CORE | T-CORE |
| INLUP_00343 | R-CORE | |
| INLUP_00344 | R-CORE | T-CORE |
| INLUP_00345 | R-CORE | T-CORE |
| INLUP_00346 | R-CORE | T-CORE |
| INLUP_00347 | R-CORE | T-CORE |
| INLUP_00348 | R-CORE | T-CORE |
| INLUP_00349 | R-CORE | |
| INLUP_00350 | R-CORE | |
| INLUP_00351 | R-CORE | T-CORE |
| INLUP_00352 | R-CORE | |
| INLUP_00353 | R-CORE | T-CORE |
| INLUP_00354 | R-CORE | T-CORE |
| INLUP_00355 | R-CORE | |
| INLUP_00356 | R-CORE | T-CORE |
| INLUP_00357 | R-CORE | |
| INLUP_00358 | R-CORE | |
| INLUP_00359 | R-CORE | T-CORE |
| INLUP_00360 | R-CORE | |
| INLUP_00361 | R-CORE | T-CORE |
| INLUP_00362 | R-CORE | |

| | | |
|---|---|---|
| INLUP_00363 | R-CORE | |
| INLUP_00364 | R-CORE | |
| INLUP_00365 | R-CORE | |
| INLUP_00366 | R-CORE | T-CORE |
| INLUP_00367 | R-CORE | |
| INLUP_00368 | R-CORE | T-CORE |
| INLUP_00369 | R-CORE | |
| INLUP_00370 | R-CORE | |
| INLUP_00371 | R-CORE | T-CORE |
| INLUP_00372 | R-CORE | |
| INLUP_00373 | R-CORE | T-CORE |
| INLUP_00374 | R-CORE | |
| INLUP_00375 | R-CORE | T-CORE |
| INLUP_00376 | R-CORE | |
| INLUP_00377 | R-CORE | |
| INLUP_00378 | R-CORE | |
| INLUP_00379 | R-CORE | T-CORE |
| INLUP_00380 | R-CORE | T-CORE |
| INLUP_00381 | R-CORE | T-CORE |
| INLUP_00382 | R-CORE | |
| INLUP_00383 | R-CORE | |
| INLUP_00384 | R-CORE | T-CORE |
| INLUP_00385 | R-CORE | T-CORE |
| INLUP_00386 | R-CORE | |
| INLUP_00387 | R-CORE | T-CORE |
| INLUP_00388 | R-CORE | T-CORE |
| INLUP_00389 | R-CORE | T-CORE |
| INLUP_00390 | R-CORE | |
| INLUP_00391 | R-CORE | |
| INLUP_00392 | R-CORE | T-CORE |
| INLUP_00393 | R-CORE | T-CORE |
| INLUP_00394 | R-CORE | |
| INLUP_00395 | R-CORE | |
| INLUP_00396 | R-CORE | |
| INLUP_00397 | R-CORE | |
| INLUP_00398 | R-CORE | T-CORE |
| INLUP_00399 | R-CORE | T-CORE |
| INLUP_00400 | R-CORE | T-CORE |
| INLUP_00401 | R-CORE | T-CORE |
| INLUP_00402 | R-CORE | T-CORE |
| INLUP_00403 | R-CORE | |
| INLUP_00404 | R-CORE | T-CORE |
| INLUP_00405 | R-CORE | T-CORE |
| INLUP_00406 | R-CORE | |
| INLUP_00407 | R-CORE | |
| INLUP_00408 | R-CORE | T-CORE |
| INLUP_00409 | R-CORE | |
| INLUP_00410 | R-CORE | |
| INLUP_00411 | R-CORE | T-CORE |
| INLUP_00412 | R-CORE | |
| INLUP_00413 | R-CORE | T-CORE |
| INLUP_00414 | R-CORE | T-CORE |

| | | |
|---|---|---|
| INLUP_00415 | R-CORE | T-CORE |
| INLUP_00416 | R-CORE | T-CORE |
| INLUP_00417 | R-CORE | T-CORE |
| INLUP_00418 | R-CORE | T-CORE |
| INLUP_00419 | R-CORE | T-CORE |
| INLUP_00420 | R-CORE | T-CORE |
| INLUP_00421 | R-CORE | |
| INLUP_00422 | R-CORE | |
| INLUP_00423 | R-CORE | |
| INLUP_00424 | R-CORE | T-CORE |
| INLUP_00425 | R-CORE | T-CORE |
| INLUP_00426 | R-CORE | |
| INLUP_00427 | R-CORE | |
| INLUP_00428 | R-CORE | T-CORE |
| INLUP_00429 | R-CORE | T-CORE |
| INLUP_00430 | R-CORE | |
| INLUP_00431 | R-CORE | T-CORE |
| INLUP_00432 | R-CORE | T-CORE |
| INLUP_00433 | R-CORE | T-CORE |
| INLUP_00434 | R-CORE | |
| INLUP_00435 | R-CORE | T-CORE |
| INLUP_00436 | R-CORE | T-CORE |
| INLUP_00437 | R-CORE | |
| INLUP_00438 | R-CORE | |
| INLUP_00439 | R-CORE | |
| INLUP_00440 | R-CORE | T-CORE |
| INLUP_00441 | R-CORE | |
| INLUP_00442 | R-CORE | T-CORE |
| INLUP_00443 | R-CORE | T-CORE |
| INLUP_00444 | R-CORE | |
| INLUP_00445 | R-CORE | |
| INLUP_00446 | R-CORE | T-CORE |
| INLUP_00447 | R-CORE | |
| INLUP_00448 | R-CORE | T-CORE |
| INLUP_00449 | R-CORE | |
| INLUP_00450 | R-CORE | T-CORE |
| INLUP_00451 | R-CORE | T-CORE |
| INLUP_00452 | R-CORE | T-CORE |
| INLUP_00453 | R-CORE | |
| INLUP_00454 | R-CORE | T-CORE |
| INLUP_00455 | R-CORE | T-CORE |
| INLUP_00456 | R-CORE | T-CORE |
| INLUP_00457 | R-CORE | T-CORE |
| INLUP_00458 | R-CORE | T-CORE |
| INLUP_00459 | R-CORE | T-CORE |
| INLUP_00460 | R-CORE | |
| INLUP_00461 | R-CORE | |
| INLUP_00462 | R-CORE | |
| INLUP_00463 | R-CORE | |
| INLUP_00464 | R-CORE | |
| INLUP_00465 | R-CORE | |
| INLUP_00466 | R-CORE | |

| | | |
|---|---|---|
| INLUP_00467 | R-CORE | |
| INLUP_00468 | R-CORE | |
| INLUP_00469 | R-CORE | |
| INLUP_00470 | R-CORE | |
| INLUP_00471 | R-CORE | |
| INLUP_00472 | R-CORE | |
| INLUP_00473 | R-CORE | |
| INLUP_00474 | R-CORE | |
| INLUP_00475 | R-CORE | |
| INLUP_00476 | R-CORE | |
| INLUP_00477 | R-CORE | |
| INLUP_00478 | R-CORE | |
| INLUP_00479 | R-CORE | |
| INLUP_00480 | R-CORE | |
| INLUP_00481 | R-CORE | |
| INLUP_00482 | R-CORE | |
| INLUP_00483 | R-CORE | |
| INLUP_00484 | R-CORE | |
| INLUP_00485 | R-CORE | |
| INLUP_00486 | R-CORE | |
| INLUP_00487 | R-CORE | T-CORE |
| INLUP_00488 | R-CORE | |
| INLUP_00489 | R-CORE | T-CORE |
| INLUP_00490 | R-CORE | |
| INLUP_00491 | R-CORE | |
| INLUP_00492 | R-CORE | T-CORE |
| INLUP_00493 | R-CORE | T-CORE |
| INLUP_00494 | R-CORE | T-CORE |
| INLUP_00495 | R-CORE | T-CORE |
| INLUP_00496 | R-CORE | T-CORE |
| INLUP_00497 | R-CORE | |
| INLUP_00498 | R-CORE | |
| INLUP_00499 | R-CORE | |
| INLUP_00500 | R-CORE | |
| INLUP_00501 | R-CORE | |
| INLUP_00502 | R-CORE | T-CORE |
| INLUP_00503 | R-CORE | |
| INLUP_00504 | R-CORE | |
| INLUP_00505 | R-CORE | T-CORE |
| INLUP_00506 | R-CORE | |
| INLUP_00507 | R-CORE | |
| INLUP_00508 | R-CORE | |
| INLUP_00509 | R-CORE | T-CORE |
| INLUP_00510 | R-CORE | T-CORE |
| INLUP_00511 | R-CORE | |
| INLUP_00512 | R-CORE | T-CORE |
| INLUP_00513 | R-CORE | |
| INLUP_00514 | R-CORE | |
| INLUP_00515 | R-CORE | |
| INLUP_00516 | R-CORE | |
| INLUP_00517 | R-CORE | |
| INLUP_00518 | R-CORE | |

| | | |
|---|---|---|
| INLUP_00519 | R-CORE | |
| INLUP_00520 | R-CORE | |
| INLUP_00521 | R-CORE | |
| INLUP_00522 | R-CORE | |
| INLUP_00523 | R-CORE | |
| INLUP_00524 | R-CORE | |
| INLUP_00525 | R-CORE | |
| INLUP_00526 | R-CORE | T-CORE |
| INLUP_00527 | R-CORE | |
| INLUP_00528 | R-CORE | T-CORE |
| INLUP_00529 | R-CORE | |
| INLUP_00530 | R-CORE | |
| INLUP_00531 | R-CORE | |
| INLUP_00532 | R-CORE | T-CORE |
| INLUP_00533 | R-CORE | |
| INLUP_00534 | R-CORE | T-CORE |
| INLUP_00535 | R-CORE | T-CORE |
| INLUP_00536 | R-CORE | |
| INLUP_00537 | R-CORE | |
| INLUP_00538 | R-CORE | T-CORE |
| INLUP_00539 | R-CORE | |
| INLUP_00540 | R-CORE | T-CORE |
| INLUP_00541 | R-CORE | T-CORE |
| INLUP_00542 | R-CORE | T-CORE |
| INLUP_00543 | R-CORE | |
| INLUP_00544 | R-CORE | T-CORE |
| INLUP_00545 | R-CORE | T-CORE |
| INLUP_00546 | R-CORE | |
| INLUP_00547 | R-CORE | T-CORE |
| INLUP_00548 | R-CORE | T-CORE |
| INLUP_00549 | R-CORE | |
| INLUP_00550 | R-CORE | |
| INLUP_00551 | R-CORE | |
| INLUP_00552 | R-CORE | T-CORE |
| INLUP_00553 | R-CORE | T-CORE |
| INLUP_00554 | R-CORE | T-CORE |
| INLUP_00555 | R-CORE | |
| INLUP_00556 | R-CORE | T-CORE |
| INLUP_00557 | R-CORE | T-CORE |
| INLUP_00558 | R-CORE | T-CORE |
| INLUP_00559 | R-CORE | |
| INLUP_00560 | R-CORE | T-CORE |
| INLUP_00561 | R-CORE | |
| INLUP_00562 | R-CORE | T-CORE |
| INLUP_00563 | R-CORE | |
| INLUP_00564 | R-CORE | |
| INLUP_00565 | R-CORE | T-CORE |
| INLUP_00566 | R-CORE | T-CORE |
| INLUP_00567 | R-CORE | T-CORE |
| INLUP_00568 | R-CORE | |
| INLUP_00569 | R-CORE | |
| INLUP_00570 | R-CORE | |

| | | |
|---|---|---|
| INLUP_00571 | R-CORE | |
| INLUP_00572 | R-CORE | |
| INLUP_00573 | R-CORE | |
| INLUP_00574 | R-CORE | T-CORE |
| INLUP_00575 | R-CORE | T-CORE |
| INLUP_00576 | R-CORE | T-CORE |
| INLUP_00577 | R-CORE | T-CORE |
| INLUP_00578 | R-CORE | |
| INLUP_00579 | R-CORE | |
| INLUP_00580 | R-CORE | |
| INLUP_00581 | R-CORE | |
| INLUP_00582 | R-CORE | |
| INLUP_00583 | R-CORE | |
| INLUP_00584 | R-CORE | T-CORE |
| INLUP_00585 | R-CORE | T-CORE |
| INLUP_00586 | R-CORE | |
| INLUP_00587 | R-CORE | T-CORE |
| INLUP_00588 | R-CORE | T-CORE |
| INLUP_00589 | R-CORE | T-CORE |
| INLUP_00590 | R-CORE | T-CORE |
| INLUP_00591 | R-CORE | |
| INLUP_00592 | R-CORE | T-CORE |
| INLUP_00593 | R-CORE | |
| INLUP_00594 | R-CORE | T-CORE |
| INLUP_00595 | R-CORE | |
| INLUP_00596 | R-CORE | |
| INLUP_00597 | R-CORE | T-CORE |
| INLUP_00598 | R-CORE | T-CORE |
| INLUP_00599 | R-CORE | |
| INLUP_00600 | R-CORE | |
| INLUP_00601 | R-CORE | |
| INLUP_00602 | R-CORE | |
| INLUP_00603 | R-CORE | |
| INLUP_00604 | R-CORE | |
| INLUP_00605 | R-CORE | |
| INLUP_00606 | R-CORE | |
| INLUP_00607 | R-CORE | |
| INLUP_00608 | R-CORE | T-CORE |
| INLUP_00609 | R-CORE | T-CORE |
| INLUP_00610 | R-CORE | |
| INLUP_00611 | R-CORE | |
| INLUP_00612 | R-CORE | |
| INLUP_00613 | R-CORE | |
| INLUP_00614 | R-CORE | |
| INLUP_00615 | R-CORE | T-CORE |
| INLUP_00616 | R-CORE | T-CORE |
| INLUP_00617 | R-CORE | |
| INLUP_00618 | R-CORE | |
| INLUP_00619 | R-CORE | |
| INLUP_00620 | R-CORE | |
| INLUP_00621 | R-CORE | T-CORE |
| INLUP_00622 | R-CORE | |

| | | |
|---|---|---|
| INLUP_00623 | R-CORE | |
| INLUP_00624 | R-CORE | |
| INLUP_00625 | R-CORE | |
| INLUP_00626 | R-CORE | |
| INLUP_00627 | R-CORE | |
| INLUP_00628 | R-CORE | |
| INLUP_00629 | R-CORE | T-CORE |
| INLUP_00630 | R-CORE | |
| INLUP_00631 | R-CORE | |
| INLUP_00632 | R-CORE | |
| INLUP_00633 | R-CORE | T-CORE |
| INLUP_00634 | R-CORE | |
| INLUP_00635 | R-CORE | |
| INLUP_00636 | R-CORE | |
| INLUP_00637 | R-CORE | |
| INLUP_00638 | R-CORE | |
| INLUP_00639 | R-CORE | |
| INLUP_00640 | R-CORE | |
| INLUP_00641 | R-CORE | |
| INLUP_00642 | R-CORE | |
| INLUP_00643 | R-CORE | |
| INLUP_00644 | R-CORE | |
| INLUP_00645 | R-CORE | |
| INLUP_00646 | R-CORE | |
| INLUP_00647 | R-CORE | |
| INLUP_00648 | R-CORE | |
| INLUP_00649 | R-CORE | |
| INLUP_00650 | R-CORE | |
| INLUP_00651 | R-CORE | T-CORE |
| INLUP_00652 | R-CORE | T-CORE |
| INLUP_00653 | R-CORE | |
| INLUP_00654 | R-CORE | |
| INLUP_00655 | R-CORE | |
| INLUP_00656 | R-CORE | |
| INLUP_00657 | R-CORE | |
| INLUP_00658 | R-CORE | T-CORE |
| INLUP_00659 | R-CORE | |
| INLUP_00660 | R-CORE | |
| INLUP_00661 | R-CORE | |
| INLUP_00662 | R-CORE | |
| INLUP_00663 | R-CORE | |
| INLUP_00664 | R-CORE | |
| INLUP_00665 | R-CORE | |
| INLUP_00666 | R-CORE | |
| INLUP_00667 | R-CORE | |
| INLUP_00668 | R-CORE | |
| INLUP_00670 | R-CORE | |
| INLUP_00671 | R-CORE | |
| INLUP_00672 | R-CORE | |
| INLUP_00673 | R-CORE | |
| INLUP_00674 | R-CORE | |
| INLUP_00675 | R-CORE | |

| | | |
|---|---|---|
| INLUP_00676 | R-CORE | |
| INLUP_00677 | R-CORE | |
| INLUP_00678 | R-CORE | |
| INLUP_00679 | R-CORE | |
| INLUP_00680 | R-CORE | T-CORE |
| INLUP_00681 | R-CORE | |
| INLUP_00682 | R-CORE | |
| INLUP_00683 | R-CORE | |
| INLUP_00684 | R-CORE | |
| INLUP_00685 | R-CORE | T-CORE |
| INLUP_00686 | R-CORE | T-CORE |
| INLUP_00687 | R-CORE | |
| INLUP_00688 | R-CORE | |
| INLUP_00689 | R-CORE | |
| INLUP_00690 | R-CORE | |
| INLUP_00691 | R-CORE | T-CORE |
| INLUP_00692 | R-CORE | |
| INLUP_00693 | R-CORE | |
| INLUP_00694 | R-CORE | |
| INLUP_00695 | R-CORE | |
| INLUP_00696 | R-CORE | |
| INLUP_00697 | R-CORE | |
| INLUP_00698 | R-CORE | |
| INLUP_00699 | R-CORE | |
| INLUP_00700 | R-CORE | |
| INLUP_00701 | R-CORE | T-CORE |
| INLUP_00702 | R-CORE | |
| INLUP_00703 | R-CORE | |
| INLUP_00704 | R-CORE | T-CORE |
| INLUP_00705 | R-CORE | |
| INLUP_00706 | R-CORE | |
| INLUP_00707 | R-CORE | |
| INLUP_00708 | R-CORE | |
| INLUP_00709 | R-CORE | |
| INLUP_00710 | R-CORE | |
| INLUP_00711 | R-CORE | T-CORE |
| INLUP_00712 | R-CORE | |
| INLUP_00713 | R-CORE | |
| INLUP_00714 | R-CORE | |
| INLUP_00715 | R-CORE | |
| INLUP_00716 | R-CORE | |
| INLUP_00717 | R-CORE | |
| INLUP_00718 | R-CORE | |
| INLUP_00719 | R-CORE | |
| INLUP_00720 | R-CORE | |
| INLUP_00721 | R-CORE | |
| INLUP_00722 | R-CORE | |
| INLUP_00723 | R-CORE | |
| INLUP_00724 | R-CORE | |
| INLUP_00725 | R-CORE | |
| INLUP_00726 | R-CORE | |
| INLUP_00727 | R-CORE | |

| | | |
|---|---|---|
| INLUP_00728 | R-CORE | |
| INLUP_00729 | R-CORE | |
| INLUP_00730 | R-CORE | |
| INLUP_00731 | R-CORE | |
| INLUP_00732 | R-CORE | |
| INLUP_00733 | R-CORE | |
| INLUP_00734 | R-CORE | |
| INLUP_00735 | R-CORE | |
| INLUP_00736 | R-CORE | |
| INLUP_00737 | R-CORE | |
| INLUP_00738 | R-CORE | |
| INLUP_00739 | R-CORE | T-CORE |
| INLUP_00740 | R-CORE | |
| INLUP_00741 | R-CORE | |
| INLUP_00742 | R-CORE | |
| INLUP_00743 | R-CORE | |
| INLUP_00744 | R-CORE | |
| INLUP_00745 | R-CORE | |
| INLUP_00746 | R-CORE | |
| INLUP_00747 | R-CORE | |
| INLUP_00748 | R-CORE | |
| INLUP_00749 | R-CORE | |
| INLUP_00750 | R-CORE | |
| INLUP_00751 | R-CORE | |
| INLUP_00752 | R-CORE | |
| INLUP_00753 | R-CORE | |
| INLUP_00754 | R-CORE | |
| INLUP_00755 | R-CORE | |
| INLUP_00756 | R-CORE | |
| INLUP_00757 | R-CORE | |
| INLUP_00758 | R-CORE | |
| INLUP_00759 | R-CORE | T-CORE |
| INLUP_00760 | R-CORE | |
| INLUP_00761 | R-CORE | |
| INLUP_00762 | R-CORE | |
| INLUP_00763 | R-CORE | |
| INLUP_00764 | R-CORE | |
| INLUP_00765 | R-CORE | |
| INLUP_00766 | R-CORE | |
| INLUP_00767 | R-CORE | |
| INLUP_00768 | R-CORE | |
| INLUP_00769 | R-CORE | |
| INLUP_00770 | R-CORE | |
| INLUP_00771 | R-CORE | |
| INLUP_00772 | R-CORE | |
| INLUP_00773 | R-CORE | |
| INLUP_00774 | R-CORE | |
| INLUP_00775 | R-CORE | |
| INLUP_00776 | R-CORE | |
| INLUP_00777 | R-CORE | |
| INLUP_00778 | R-CORE | |
| INLUP_00779 | R-CORE | |

| | | |
|---|---|---|
| INLUP_00780 | R-CORE | |
| INLUP_00781 | R-CORE | |
| INLUP_00782 | R-CORE | |
| INLUP_00783 | R-CORE | |
| INLUP_00784 | R-CORE | |
| INLUP_00785 | R-CORE | |
| INLUP_00786 | R-CORE | |
| INLUP_00787 | R-CORE | |
| INLUP_00788 | R-CORE | |
| INLUP_00789 | R-CORE | |
| INLUP_00790 | R-CORE | |
| INLUP_00791 | R-CORE | |
| INLUP_00792 | R-CORE | |
| INLUP_00793 | R-CORE | |
| INLUP_00794 | R-CORE | |
| INLUP_00795 | R-CORE | |
| INLUP_00796 | R-CORE | |
| INLUP_00797 | R-CORE | |
| INLUP_00798 | R-CORE | |
| INLUP_00799 | R-CORE | |
| INLUP_00800 | R-CORE | |
| INLUP_00801 | R-CORE | |
| INLUP_00802 | R-CORE | |
| INLUP_00803 | R-CORE | |
| INLUP_00804 | R-CORE | |
| INLUP_00805 | R-CORE | |
| INLUP_00806 | R-CORE | |
| INLUP_00807 | R-CORE | |
| INLUP_00808 | R-CORE | |
| INLUP_00809 | R-CORE | |
| INLUP_00810 | R-CORE | |
| INLUP_00811 | R-CORE | |
| INLUP_00812 | R-CORE | |
| INLUP_00813 | R-CORE | |
| INLUP_00814 | R-CORE | |
| INLUP_00815 | R-CORE | |
| INLUP_00816 | R-CORE | |
| INLUP_00817 | R-CORE | |
| INLUP_00818 | R-CORE | |
| INLUP_00819 | R-CORE | |
| INLUP_00820 | R-CORE | |
| INLUP_00821 | R-CORE | |
| INLUP_00822 | R-CORE | |
| INLUP_00823 | R-CORE | |
| INLUP_00824 | R-CORE | |
| INLUP_00825 | R-CORE | T-CORE |
| INLUP_00826 | R-CORE | T-CORE |
| INLUP_00827 | R-CORE | |
| INLUP_00828 | R-CORE | |
| INLUP_00829 | R-CORE | |
| INLUP_00830 | R-CORE | |
| INLUP_00831 | R-CORE | |

| | |
|---|---|
| INLUP_00832 | R-CORE |
| INLUP_00833 | R-CORE |
| INLUP_00834 | R-CORE |
| INLUP_00836 | R-CORE |
| INLUP_00837 | R-CORE |
| INLUP_00838 | R-CORE |
| INLUP_00839 | R-CORE |
| INLUP_00840 | R-CORE |
| INLUP_00841 | R-CORE |
| INLUP_00842 | R-CORE |
| INLUP_00843 | R-CORE |
| INLUP_00844 | R-CORE |
| INLUP_00845 | R-CORE |
| INLUP_00846 | R-CORE |
| INLUP_00847 | R-CORE |
| INLUP_00848 | R-CORE |
| INLUP_00849 | R-CORE |
| INLUP_00850 | R-CORE |
| INLUP_00851 | R-CORE |
| INLUP_00852 | R-CORE |
| INLUP_00853 | R-CORE |
| INLUP_00854 | R-CORE |
| INLUP_00855 | R-CORE |
| INLUP_00856 | R-CORE |
| INLUP_00857 | R-CORE |
| INLUP_00858 | R-CORE |
| INLUP_00859 | R-CORE |
| INLUP_00860 | R-CORE |
| INLUP_00861 | R-CORE |
| INLUP_00862 | R-CORE |
| INLUP_00863 | R-CORE |
| INLUP_00864 | R-CORE |
| INLUP_00865 | R-CORE |
| INLUP_00866 | R-CORE |
| INLUP_00867 | R-CORE |
| INLUP_00868 | R-CORE |
| INLUP_00869 | R-CORE |
| INLUP_00870 | R-CORE |
| INLUP_00871 | R-CORE |
| INLUP_00872 | R-CORE |
| INLUP_00873 | R-CORE |
| INLUP_00874 | R-CORE |
| INLUP_00875 | R-CORE |
| INLUP_00876 | R-CORE |
| INLUP_00877 | R-CORE |
| INLUP_00878 | R-CORE |
| INLUP_00879 | R-CORE |
| INLUP_00880 | R-CORE |
| INLUP_00881 | R-CORE |
| INLUP_00882 | R-CORE |
| INLUP_00883 | R-CORE |
| INLUP_00884 | R-CORE |

| | |
|---|---|
| INLUP_00885 | R-CORE |
| INLUP_00886 | R-CORE |
| INLUP_00887 | R-CORE |
| INLUP_00888 | R-CORE |
| INLUP_00889 | R-CORE |
| INLUP_00890 | R-CORE |
| INLUP_00891 | R-CORE |
| INLUP_00892 | R-CORE |
| INLUP_00893 | R-CORE |
| INLUP_00894 | R-CORE |
| INLUP_00895 | R-CORE |
| INLUP_00896 | R-CORE |
| INLUP_00897 | R-CORE |
| INLUP_00898 | R-CORE |
| INLUP_00899 | R-CORE |
| INLUP_00900 | R-CORE |
| INLUP_00901 | R-CORE |
| INLUP_00902 | R-CORE |
| INLUP_00903 | R-CORE |
| INLUP_00904 | R-CORE |
| INLUP_00905 | R-CORE |
| INLUP_00906 | R-CORE |
| INLUP_00907 | R-CORE |
| INLUP_00908 | R-CORE |
| INLUP_00909 | R-CORE |
| INLUP_00910 | R-CORE |
| INLUP_00911 | R-CORE |
| INLUP_00912 | R-CORE |
| INLUP_00913 | R-CORE |
| INLUP_00914 | R-CORE |
| INLUP_00915 | R-CORE |
| INLUP_00916 | R-CORE |
| INLUP_00917 | R-CORE |
| INLUP_00918 | R-CORE |
| INLUP_00919 | R-CORE |
| INLUP_00920 | R-CORE |
| INLUP_00921 | R-CORE |
| INLUP_00922 | R-CORE |
| INLUP_00923 | R-CORE |
| INLUP_00924 | R-CORE |
| INLUP_00925 | R-CORE |
| INLUP_00926 | R-CORE |
| INLUP_00927 | R-CORE |
| INLUP_00928 | R-CORE |
| INLUP_00929 | R-CORE |
| INLUP_00930 | R-CORE |
| INLUP_00931 | R-CORE |
| INLUP_00932 | R-CORE |
| INLUP_00933 | R-CORE |
| INLUP_00934 | R-CORE |
| INLUP_00935 | R-CORE |
| INLUP_00936 | R-CORE |

| | |
|---|---|
| INLUP_00937 | R-CORE |
| INLUP_00938 | R-CORE |
| INLUP_00939 | R-CORE |
| INLUP_00940 | R-CORE |
| INLUP_00941 | R-CORE |
| INLUP_00942 | R-CORE |
| INLUP_00943 | R-CORE |
| INLUP_00944 | R-CORE |
| INLUP_00945 | R-CORE |
| INLUP_00946 | R-CORE |
| INLUP_00947 | R-CORE |
| INLUP_00948 | R-CORE |
| INLUP_00949 | R-CORE |
| INLUP_00950 | R-CORE |
| INLUP_00951 | R-CORE |
| INLUP_00952 | R-CORE |
| INLUP_00953 | R-CORE |
| INLUP_00954 | R-CORE |
| INLUP_00955 | R-CORE |
| INLUP_00956 | R-CORE |
| INLUP_00957 | R-CORE |
| INLUP_00958 | R-CORE |
| INLUP_00959 | R-CORE |
| INLUP_00960 | R-CORE |
| INLUP_00961 | R-CORE |
| INLUP_00962 | R-CORE |
| INLUP_00963 | R-CORE |
| INLUP_00964 | R-CORE |
| INLUP_00965 | R-CORE |
| INLUP_00966 | R-CORE |
| INLUP_00967 | R-CORE |
| INLUP_00968 | R-CORE |
| INLUP_00969 | R-CORE |
| INLUP_00970 | R-CORE |
| INLUP_00971 | R-CORE |
| INLUP_00972 | R-CORE |
| INLUP_00973 | R-CORE |
| INLUP_00974 | R-CORE |
| INLUP_00975 | R-CORE |
| INLUP_00976 | R-CORE |
| INLUP_00977 | R-CORE |
| INLUP_00978 | R-CORE |
| INLUP_00979 | R-CORE |
| INLUP_00980 | R-CORE |
| INLUP_00981 | R-CORE |
| INLUP_00982 | R-CORE |
| INLUP_00983 | R-CORE |
| INLUP_00984 | R-CORE |
| INLUP_00985 | R-CORE |
| INLUP_00986 | R-CORE |
| INLUP_00987 | R-CORE |
| INLUP_00988 | R-CORE |

| | |
|---|---|
| INLUP_00989 | R-CORE |
| INLUP_00990 | R-CORE |
| INLUP_00991 | R-CORE |
| INLUP_00992 | R-CORE |
| INLUP_00993 | R-CORE |
| INLUP_00994 | R-CORE |
| INLUP_00995 | R-CORE |
| INLUP_00996 | R-CORE |
| INLUP_00997 | R-CORE |
| INLUP_00998 | R-CORE |
| INLUP_00999 | R-CORE |
| INLUP_01000 | R-CORE |
| INLUP_01001 | R-CORE |
| INLUP_01002 | R-CORE |
| INLUP_01003 | R-CORE |
| INLUP_01004 | R-CORE |
| INLUP_01005 | R-CORE |
| INLUP_01006 | R-CORE |
| INLUP_01007 | R-CORE |
| INLUP_01008 | R-CORE |
| INLUP_01009 | R-CORE |
| INLUP_01010 | R-CORE |
| INLUP_01011 | R-CORE |
| INLUP_01012 | R-CORE |
| INLUP_01013 | R-CORE |
| INLUP_01014 | R-CORE |
| INLUP_01015 | R-CORE |
| INLUP_01016 | R-CORE |
| INLUP_01017 | R-CORE |
| INLUP_01018 | R-CORE |
| INLUP_01019 | R-CORE |
| INLUP_01020 | R-CORE |
| INLUP_01021 | R-CORE |
| INLUP_01022 | R-CORE |
| INLUP_01023 | R-CORE |
| INLUP_01024 | R-CORE |
| INLUP_01025 | R-CORE |
| INLUP_01026 | R-CORE |
| INLUP_01027 | R-CORE |
| INLUP_01028 | R-CORE |
| INLUP_01029 | R-CORE |
| INLUP_01030 | R-CORE |
| INLUP_01031 | R-CORE |
| INLUP_01032 | R-CORE |
| INLUP_01033 | R-CORE |
| INLUP_01034 | R-CORE |
| INLUP_01035 | R-CORE |
| INLUP_01036 | R-CORE |
| INLUP_01037 | R-CORE |
| INLUP_01038 | R-CORE |
| INLUP_01039 | R-CORE |
| INLUP_01040 | R-CORE |

| | |
|---|---|
| INLUP_01041 | R-CORE |
| INLUP_01042 | R-CORE |
| INLUP_01043 | R-CORE |
| INLUP_01044 | R-CORE |
| INLUP_01045 | R-CORE |
| INLUP_01046 | R-CORE |
| INLUP_01047 | R-CORE |
| INLUP_01048 | R-CORE |
| INLUP_01049 | R-CORE |
| INLUP_01050 | R-CORE |
| INLUP_01051 | R-CORE |
| INLUP_01052 | R-CORE |
| INLUP_01053 | R-CORE |
| INLUP_01054 | R-CORE |
| INLUP_01055 | R-CORE |
| INLUP_01056 | R-CORE |
| INLUP_01057 | R-CORE |
| INLUP_01058 | R-CORE |
| INLUP_01059 | R-CORE |
| INLUP_01060 | R-CORE |
| INLUP_01061 | R-CORE |
| INLUP_01062 | R-CORE |
| INLUP_01063 | R-CORE |
| INLUP_01064 | R-CORE |
| INLUP_01065 | R-CORE |
| INLUP_01066 | R-CORE |
| INLUP_01067 | R-CORE |
| INLUP_01068 | R-CORE |
| INLUP_01069 | R-CORE |
| INLUP_01070 | R-CORE |
| INLUP_01071 | R-CORE |
| INLUP_01072 | R-CORE |
| INLUP_01073 | R-CORE |
| INLUP_01074 | R-CORE |
| INLUP_01075 | R-CORE |
| INLUP_01076 | R-CORE |
| INLUP_01077 | R-CORE |
| INLUP_01078 | R-CORE |
| INLUP_01079 | R-CORE |
| INLUP_01080 | R-CORE |
| INLUP_01081 | R-CORE |
| INLUP_01082 | R-CORE |
| INLUP_01083 | R-CORE |
| INLUP_01084 | R-CORE |
| INLUP_01085 | R-CORE |
| INLUP_01086 | R-CORE |
| INLUP_01087 | R-CORE |
| INLUP_01088 | R-CORE |
| INLUP_01089 | R-CORE |
| INLUP_01090 | R-CORE |
| INLUP_01091 | R-CORE |
| INLUP_01092 | R-CORE |

| | |
|---|---|
| INLUP_01093 | R-CORE |
| INLUP_01094 | R-CORE |
| INLUP_01095 | R-CORE |
| INLUP_01096 | R-CORE |
| INLUP_01097 | R-CORE |
| INLUP_01098 | R-CORE |
| INLUP_01099 | R-CORE |
| INLUP_01100 | R-CORE |
| INLUP_01101 | R-CORE |
| INLUP_01102 | R-CORE |
| INLUP_01103 | R-CORE |
| INLUP_01104 | R-CORE |
| INLUP_01105 | R-CORE |
| INLUP_01106 | R-CORE |
| INLUP_01107 | R-CORE |
| INLUP_01108 | R-CORE |
| INLUP_01109 | R-CORE |
| INLUP_01110 | R-CORE |
| INLUP_01111 | R-CORE |
| INLUP_01112 | R-CORE |
| INLUP_01113 | R-CORE |
| INLUP_01114 | R-CORE |
| INLUP_01115 | R-CORE |
| INLUP_01116 | R-CORE |
| INLUP_01117 | R-CORE |
| INLUP_01118 | R-CORE |
| INLUP_01119 | R-CORE |
| INLUP_01120 | R-CORE |
| INLUP_01121 | R-CORE |
| INLUP_01122 | R-CORE |
| INLUP_01123 | R-CORE |
| INLUP_01124 | R-CORE |
| INLUP_01125 | R-CORE |
| INLUP_01126 | R-CORE |
| INLUP_01127 | R-CORE |
| INLUP_01128 | R-CORE |
| INLUP_01129 | R-CORE |
| INLUP_01130 | R-CORE |
| INLUP_01131 | R-CORE |
| INLUP_01132 | R-CORE |
| INLUP_01133 | R-CORE |
| INLUP_01134 | R-CORE |
| INLUP_01135 | R-CORE |
| INLUP_01136 | R-CORE |
| INLUP_01137 | R-CORE |
| INLUP_01138 | R-CORE |
| INLUP_01139 | R-CORE |
| INLUP_01140 | R-CORE |
| INLUP_01141 | R-CORE |
| INLUP_01142 | R-CORE |
| INLUP_01143 | R-CORE |
| INLUP_01144 | R-CORE |

| | |
|---|---|
| INLUP_01145 | R-CORE |
| INLUP_01146 | R-CORE |
| INLUP_01147 | R-CORE |
| INLUP_01148 | R-CORE |
| INLUP_01149 | R-CORE |
| INLUP_01150 | R-CORE |
| INLUP_01151 | R-CORE |
| INLUP_01152 | R-CORE |
| INLUP_01153 | R-CORE |
| INLUP_01154 | R-CORE |
| INLUP_01155 | R-CORE |
| INLUP_01156 | R-CORE |
| INLUP_01157 | R-CORE |
| INLUP_01158 | R-CORE |
| INLUP_01159 | R-CORE |
| INLUP_01160 | R-CORE |
| INLUP_01161 | R-CORE |
| INLUP_01162 | R-CORE |
| INLUP_01163 | R-CORE |
| INLUP_01164 | R-CORE |
| INLUP_01165 | R-CORE |
| INLUP_01166 | R-CORE |
| INLUP_01167 | R-CORE |
| INLUP_01168 | R-CORE |
| INLUP_01169 | R-CORE |
| INLUP_01170 | R-CORE |
| INLUP_01171 | R-CORE |
| INLUP_01172 | R-CORE |
| INLUP_01173 | R-CORE |
| INLUP_01174 | R-CORE |
| INLUP_01175 | R-CORE |
| INLUP_01176 | R-CORE |
| INLUP_01177 | R-CORE |
| INLUP_01178 | R-CORE |
| INLUP_01179 | R-CORE |
| INLUP_01180 | R-CORE |
| INLUP_01181 | R-CORE |
| INLUP_01182 | R-CORE |
| INLUP_01183 | R-CORE |
| INLUP_01184 | R-CORE |
| INLUP_01185 | R-CORE |
| INLUP_01186 | R-CORE |
| INLUP_01187 | R-CORE |
| INLUP_01188 | R-CORE |
| INLUP_01189 | R-CORE |
| INLUP_01190 | R-CORE |
| INLUP_01191 | R-CORE |
| INLUP_01192 | R-CORE |
| INLUP_01193 | R-CORE |
| INLUP_01194 | R-CORE |
| INLUP_01195 | R-CORE |
| INLUP_01196 | R-CORE |

| | |
|---|---|
| INLUP_01197 | R-CORE |
| INLUP_01198 | R-CORE |
| INLUP_01199 | R-CORE |
| INLUP_01200 | R-CORE |
| INLUP_01201 | R-CORE |
| INLUP_01202 | R-CORE |
| INLUP_01203 | R-CORE |
| INLUP_01204 | R-CORE |
| INLUP_01205 | R-CORE |
| INLUP_01206 | R-CORE |
| INLUP_01207 | R-CORE |
| INLUP_01208 | R-CORE |
| INLUP_01209 | R-CORE |
| INLUP_01210 | R-CORE |
| INLUP_01211 | R-CORE |
| INLUP_01212 | R-CORE |
| INLUP_01213 | R-CORE |
| INLUP_01214 | R-CORE |
| INLUP_01215 | R-CORE |
| INLUP_01216 | R-CORE |
| INLUP_01217 | R-CORE |
| INLUP_01218 | R-CORE |
| INLUP_01219 | R-CORE |
| INLUP_01220 | R-CORE |
| INLUP_01221 | R-CORE |
| INLUP_01222 | R-CORE |
| INLUP_01223 | R-CORE |
| INLUP_01224 | R-CORE |
| INLUP_01225 | R-CORE |
| INLUP_01226 | R-CORE |
| INLUP_01227 | R-CORE |
| INLUP_01228 | R-CORE |
| INLUP_01229 | R-CORE |
| INLUP_01230 | R-CORE |
| INLUP_01231 | R-CORE |
| INLUP_01232 | R-CORE |
| INLUP_01233 | R-CORE |
| INLUP_01234 | R-CORE |
| INLUP_01235 | R-CORE |
| INLUP_01236 | R-CORE |
| INLUP_01237 | R-CORE |
| INLUP_01238 | R-CORE |
| INLUP_01239 | R-CORE |
| INLUP_01240 | R-CORE |
| INLUP_01241 | R-CORE |
| INLUP_01242 | R-CORE |
| INLUP_01243 | R-CORE |
| INLUP_01244 | R-CORE |
| INLUP_01245 | R-CORE |
| INLUP_01246 | R-CORE |
| INLUP_01247 | R-CORE |
| INLUP_01248 | R-CORE |

| | |
|---|---|
| INLUP_01249 | R-CORE |
| INLUP_01250 | R-CORE |
| INLUP_01251 | R-CORE |
| INLUP_01252 | R-CORE |
| INLUP_01253 | R-CORE |
| INLUP_01254 | R-CORE |
| INLUP_01255 | R-CORE |
| INLUP_01256 | R-CORE |
| INLUP_01257 | R-CORE |
| INLUP_01258 | R-CORE |
| INLUP_01259 | R-CORE |
| INLUP_01260 | R-CORE |
| INLUP_01261 | R-CORE |
| INLUP_01262 | R-CORE |
| INLUP_01263 | R-CORE |
| INLUP_01264 | R-CORE |
| INLUP_01265 | R-CORE |
| INLUP_01266 | R-CORE |
| INLUP_01267 | R-CORE |
| INLUP_01268 | R-CORE |
| INLUP_01269 | R-CORE |
| INLUP_01270 | R-CORE |
| INLUP_01271 | R-CORE |
| INLUP_01272 | R-CORE |
| INLUP_01273 | R-CORE |
| INLUP_01274 | R-CORE |
| INLUP_01275 | R-CORE |
| INLUP_01276 | R-CORE |
| INLUP_01277 | R-CORE |
| INLUP_01278 | R-CORE |
| INLUP_01279 | R-CORE |
| INLUP_01280 | R-CORE |
| INLUP_01281 | R-CORE |
| INLUP_01282 | R-CORE |
| INLUP_01283 | R-CORE |
| INLUP_01284 | R-CORE |
| INLUP_01285 | R-CORE |
| INLUP_01286 | R-CORE |
| INLUP_01287 | R-CORE |
| INLUP_01288 | R-CORE |
| INLUP_01289 | R-CORE |
| INLUP_01290 | R-CORE |
| INLUP_01291 | R-CORE |
| INLUP_01292 | R-CORE |
| INLUP_01293 | R-CORE |
| INLUP_01294 | R-CORE |
| INLUP_01295 | R-CORE |
| INLUP_01296 | R-CORE |
| INLUP_01297 | R-CORE |
| INLUP_01298 | R-CORE |
| INLUP_01299 | R-CORE |
| INLUP_01300 | R-CORE |

| | |
|---|---|
| INLUP_01301 | R-CORE |
| INLUP_01302 | R-CORE |
| INLUP_01303 | R-CORE |
| INLUP_01304 | R-CORE |
| INLUP_01305 | R-CORE |
| INLUP_01306 | R-CORE |
| INLUP_01307 | R-CORE |
| INLUP_01308 | R-CORE |
| INLUP_01309 | R-CORE |
| INLUP_01310 | R-CORE |
| INLUP_01311 | R-CORE |
| INLUP_01312 | R-CORE |
| INLUP_01313 | R-CORE |
| INLUP_01314 | R-CORE |
| INLUP_01316 | R-CORE |
| INLUP_01317 | R-CORE |
| INLUP_01318 | R-CORE |
| INLUP_01319 | R-CORE |
| INLUP_01320 | R-CORE |
| INLUP_01321 | R-CORE |
| INLUP_01322 | R-CORE |
| INLUP_01323 | R-CORE |
| INLUP_01324 | R-CORE |
| INLUP_01325 | R-CORE |
| INLUP_01326 | R-CORE |
| INLUP_01327 | R-CORE |
| INLUP_01328 | R-CORE |
| INLUP_01329 | R-CORE |
| INLUP_01330 | R-CORE |
| INLUP_01331 | R-CORE |
| INLUP_01332 | R-CORE |
| INLUP_01333 | R-CORE |
| INLUP_01334 | R-CORE |
| INLUP_01335 | R-CORE |
| INLUP_01336 | R-CORE |
| INLUP_01337 | R-CORE |
| INLUP_01338 | R-CORE |
| INLUP_01339 | R-CORE |
| INLUP_01340 | R-CORE |
| INLUP_01341 | R-CORE |
| INLUP_01342 | R-CORE |
| INLUP_01343 | R-CORE |
| INLUP_01344 | R-CORE |
| INLUP_01345 | R-CORE |
| INLUP_01346 | R-CORE |
| INLUP_01347 | R-CORE |
| INLUP_01348 | R-CORE |
| INLUP_01349 | R-CORE |
| INLUP_01350 | R-CORE |
| INLUP_01351 | R-CORE |
| INLUP_01352 | R-CORE |
| INLUP_01353 | R-CORE |

| | |
|---|---|
| INLUP_01354 | R-CORE |
| INLUP_01355 | R-CORE |
| INLUP_01356 | R-CORE |
| INLUP_01357 | R-CORE |
| INLUP_01358 | R-CORE |
| INLUP_01359 | R-CORE |
| INLUP_01360 | R-CORE |
| INLUP_01361 | R-CORE |
| INLUP_01362 | R-CORE |
| INLUP_01363 | R-CORE |
| INLUP_01364 | R-CORE |
| INLUP_01365 | R-CORE |
| INLUP_01366 | R-CORE |
| INLUP_01367 | R-CORE |
| INLUP_01368 | R-CORE |
| INLUP_01369 | R-CORE |
| INLUP_01370 | R-CORE |
| INLUP_01371 | R-CORE |
| INLUP_01372 | R-CORE |
| INLUP_01373 | R-CORE |
| INLUP_01374 | R-CORE |
| INLUP_01375 | R-CORE |
| INLUP_01376 | R-CORE |
| INLUP_01377 | R-CORE |
| INLUP_01378 | R-CORE |
| INLUP_01379 | R-CORE |
| INLUP_01380 | R-CORE |
| INLUP_01381 | R-CORE |
| INLUP_01382 | R-CORE |
| INLUP_01383 | R-CORE |
| INLUP_01384 | R-CORE |
| INLUP_01385 | R-CORE |
| INLUP_01386 | R-CORE |
| INLUP_01387 | R-CORE |
| INLUP_01388 | R-CORE |
| INLUP_01389 | R-CORE |
| INLUP_01390 | R-CORE |
| INLUP_01391 | R-CORE |
| INLUP_01392 | R-CORE |
| INLUP_01393 | R-CORE |
| INLUP_01394 | R-CORE |
| INLUP_01395 | R-CORE |
| INLUP_01396 | R-CORE |
| INLUP_01397 | R-CORE |
| INLUP_01398 | R-CORE |
| INLUP_01399 | R-CORE |
| INLUP_01400 | R-CORE |
| INLUP_01401 | R-CORE |
| INLUP_01402 | R-CORE |
| INLUP_01403 | R-CORE |
| INLUP_01404 | R-CORE |
| INLUP_01405 | R-CORE |

| | |
|---|---|
| INLUP_01406 | R-CORE |
| INLUP_01407 | R-CORE |
| INLUP_01408 | R-CORE |
| INLUP_01409 | R-CORE |
| INLUP_01410 | R-CORE |
| INLUP_01411 | R-CORE |
| INLUP_01412 | R-CORE |
| INLUP_01413 | R-CORE |
| INLUP_01414 | R-CORE |
| INLUP_01415 | R-CORE |
| INLUP_01416 | R-CORE |
| INLUP_01417 | R-CORE |
| INLUP_01418 | R-CORE |
| INLUP_01419 | R-CORE |
| INLUP_01420 | R-CORE |
| INLUP_01421 | R-CORE |
| INLUP_01422 | R-CORE |
| INLUP_01423 | R-CORE |
| INLUP_01424 | R-CORE |
| INLUP_01425 | R-CORE |
| INLUP_01426 | R-CORE |
| INLUP_01427 | R-CORE |
| INLUP_01428 | R-CORE |
| INLUP_01429 | R-CORE |
| INLUP_01430 | R-CORE |
| INLUP_01431 | R-CORE |
| INLUP_01432 | R-CORE |
| INLUP_01433 | R-CORE |
| INLUP_01434 | R-CORE |
| INLUP_01435 | R-CORE |
| INLUP_01436 | R-CORE |
| INLUP_01437 | R-CORE |
| INLUP_01438 | R-CORE |
| INLUP_01439 | R-CORE |
| INLUP_01440 | R-CORE |
| INLUP_01441 | R-CORE |
| INLUP_01442 | R-CORE |
| INLUP_01443 | R-CORE |
| INLUP_01444 | R-CORE |
| INLUP_01445 | R-CORE |
| INLUP_01446 | R-CORE |
| INLUP_01447 | R-CORE |
| INLUP_01448 | R-CORE |
| INLUP_01449 | R-CORE |
| INLUP_01450 | R-CORE |
| INLUP_01451 | R-CORE |
| INLUP_01452 | R-CORE |
| INLUP_01453 | R-CORE |
| INLUP_01454 | R-CORE |
| INLUP_01455 | R-CORE |
| INLUP_01456 | R-CORE |
| INLUP_01457 | R-CORE |

| | |
|---|---|
| INLUP_01458 | R-CORE |
| INLUP_01459 | R-CORE |
| INLUP_01460 | R-CORE |
| INLUP_01461 | R-CORE |
| INLUP_01462 | R-CORE |
| INLUP_01463 | R-CORE |
| INLUP_01464 | R-CORE |
| INLUP_01465 | R-CORE |
| INLUP_01466 | R-CORE |
| INLUP_01467 | R-CORE |
| INLUP_01468 | R-CORE |
| INLUP_01469 | R-CORE |
| INLUP_01470 | R-CORE |
| INLUP_01471 | R-CORE |
| INLUP_01472 | R-CORE |
| INLUP_01473 | R-CORE |
| INLUP_01474 | R-CORE |
| INLUP_01475 | R-CORE |
| INLUP_01476 | R-CORE |
| INLUP_01477 | R-CORE |
| INLUP_01478 | R-CORE |
| INLUP_01479 | R-CORE |
| INLUP_01480 | R-CORE |
| INLUP_01481 | R-CORE |
| INLUP_01482 | R-CORE |
| INLUP_01483 | R-CORE |
| INLUP_01484 | R-CORE |
| INLUP_01485 | R-CORE |
| INLUP_01486 | R-CORE |
| INLUP_01487 | R-CORE |
| INLUP_01488 | R-CORE |
| INLUP_01489 | R-CORE |
| INLUP_01490 | R-CORE |
| INLUP_01491 | R-CORE |
| INLUP_01492 | R-CORE |
| INLUP_01493 | R-CORE |
| INLUP_01494 | R-CORE |
| INLUP_01495 | R-CORE |
| INLUP_01496 | R-CORE |
| INLUP_01497 | R-CORE |
| INLUP_01498 | R-CORE |
| INLUP_01499 | R-CORE |
| INLUP_01500 | R-CORE |
| INLUP_01501 | R-CORE |
| INLUP_01502 | R-CORE |
| INLUP_01503 | R-CORE |
| INLUP_01504 | R-CORE |
| INLUP_01505 | R-CORE |
| INLUP_01506 | R-CORE |
| INLUP_01507 | R-CORE |
| INLUP_01508 | R-CORE |
| INLUP_01509 | R-CORE |

| | |
|---|---|
| INLUP_01510 | R-CORE |
| INLUP_01511 | R-CORE |
| INLUP_01512 | R-CORE |
| INLUP_01513 | R-CORE |
| INLUP_01514 | R-CORE |
| INLUP_01515 | R-CORE |
| INLUP_01516 | R-CORE |
| INLUP_01517 | R-CORE |
| INLUP_01518 | R-CORE |
| INLUP_01519 | R-CORE |
| INLUP_01520 | R-CORE |
| INLUP_01521 | R-CORE |
| INLUP_01522 | R-CORE |
| INLUP_01523 | R-CORE |
| INLUP_01524 | R-CORE |
| INLUP_01525 | R-CORE |
| INLUP_01526 | R-CORE |
| INLUP_01527 | R-CORE |
| INLUP_01528 | R-CORE |
| INLUP_01529 | R-CORE |
| INLUP_01530 | R-CORE |
| INLUP_01531 | R-CORE |
| INLUP_01532 | R-CORE |
| INLUP_01533 | R-CORE |
| INLUP_01534 | R-CORE |
| INLUP_01535 | R-CORE |
| INLUP_01536 | R-CORE |
| INLUP_01537 | R-CORE |
| INLUP_01538 | R-CORE |
| INLUP_01539 | R-CORE |
| INLUP_01540 | R-CORE |
| INLUP_01541 | R-CORE |
| INLUP_01542 | R-CORE |
| INLUP_01543 | R-CORE |
| INLUP_01544 | R-CORE |
| INLUP_01545 | R-CORE |
| INLUP_01546 | R-CORE |
| INLUP_01547 | R-CORE |
| INLUP_01548 | R-CORE |
| INLUP_01549 | R-CORE |
| INLUP_01550 | R-CORE |
| INLUP_01551 | R-CORE |
| INLUP_01552 | R-CORE |
| INLUP_01553 | R-CORE |
| INLUP_01554 | R-CORE |
| INLUP_01555 | R-CORE |
| INLUP_01556 | R-CORE |
| INLUP_01557 | R-CORE |
| INLUP_01558 | R-CORE |
| INLUP_01559 | R-CORE |
| INLUP_01560 | R-CORE |
| INLUP_01561 | R-CORE |

| | |
|---|---|
| INLUP_01562 | R-CORE |
| INLUP_01563 | R-CORE |
| INLUP_01564 | R-CORE |
| INLUP_01565 | R-CORE |
| INLUP_01566 | R-CORE |
| INLUP_01567 | R-CORE |
| INLUP_01568 | R-CORE |
| INLUP_01569 | R-CORE |
| INLUP_01570 | R-CORE |
| INLUP_01571 | R-CORE |
| INLUP_01572 | R-CORE |
| INLUP_01573 | R-CORE |
| INLUP_01574 | R-CORE |
| INLUP_01575 | R-CORE |
| INLUP_01576 | R-CORE |
| INLUP_01577 | R-CORE |
| INLUP_01578 | R-CORE |
| INLUP_01579 | R-CORE |
| INLUP_01580 | R-CORE |
| INLUP_01581 | R-CORE |
| INLUP_01582 | R-CORE |
| INLUP_01583 | R-CORE |
| INLUP_01584 | R-CORE |
| INLUP_01585 | R-CORE |
| INLUP_01586 | R-CORE |
| INLUP_01587 | R-CORE |
| INLUP_01588 | R-CORE |
| INLUP_01589 | R-CORE |
| INLUP_01590 | R-CORE |
| INLUP_01591 | R-CORE |
| INLUP_01592 | R-CORE |
| INLUP_01593 | R-CORE |
| INLUP_01594 | R-CORE |
| INLUP_01595 | R-CORE |
| INLUP_01596 | R-CORE |
| INLUP_01597 | R-CORE |
| INLUP_01598 | R-CORE |
| INLUP_01599 | R-CORE |
| INLUP_01600 | R-CORE |
| INLUP_01601 | R-CORE |
| INLUP_01602 | R-CORE |
| INLUP_01603 | R-CORE |
| INLUP_01604 | R-CORE |
| INLUP_01605 | R-CORE |
| INLUP_01606 | R-CORE |
| INLUP_01607 | R-CORE |
| INLUP_01608 | R-CORE |
| INLUP_01609 | R-CORE |
| INLUP_01610 | R-CORE |
| INLUP_01611 | R-CORE |
| INLUP_01612 | R-CORE |
| INLUP_01613 | R-CORE |

| | |
|---|---|
| INLUP_01614 | R-CORE |
| INLUP_01615 | R-CORE |
| INLUP_01616 | R-CORE |
| INLUP_01617 | R-CORE |
| INLUP_01618 | R-CORE |
| INLUP_01619 | R-CORE |
| INLUP_01620 | R-CORE |
| INLUP_01621 | R-CORE |
| INLUP_01622 | R-CORE |
| INLUP_01623 | R-CORE |
| INLUP_01624 | R-CORE |
| INLUP_01625 | R-CORE |
| INLUP_01626 | R-CORE |
| INLUP_01627 | R-CORE |
| INLUP_01628 | R-CORE |
| INLUP_01629 | R-CORE |
| INLUP_01630 | R-CORE |
| INLUP_01631 | R-CORE |
| INLUP_01632 | R-CORE |
| INLUP_01633 | R-CORE |
| INLUP_01634 | R-CORE |
| INLUP_01635 | R-CORE |
| INLUP_01636 | R-CORE |
| INLUP_01637 | R-CORE |
| INLUP_01638 | R-CORE |
| INLUP_01639 | R-CORE |
| INLUP_01640 | R-CORE |
| INLUP_01641 | R-CORE |
| INLUP_01642 | R-CORE |
| INLUP_01643 | R-CORE |
| INLUP_01644 | R-CORE |
| INLUP_01645 | R-CORE |
| INLUP_01646 | R-CORE |
| INLUP_01647 | R-CORE |
| INLUP_01648 | R-CORE |
| INLUP_01649 | R-CORE |
| INLUP_01650 | R-CORE |
| INLUP_01651 | R-CORE |
| INLUP_01652 | R-CORE |
| INLUP_01653 | R-CORE |
| INLUP_01654 | R-CORE |
| INLUP_01655 | R-CORE |
| INLUP_01656 | R-CORE |
| INLUP_01657 | R-CORE |
| INLUP_01658 | R-CORE |
| INLUP_01659 | R-CORE |
| INLUP_01660 | R-CORE |
| INLUP_01661 | R-CORE |
| INLUP_01662 | R-CORE |
| INLUP_01663 | R-CORE |
| INLUP_01664 | R-CORE |
| INLUP_01665 | R-CORE |

| | |
|---|---|
| INLUP_01666 | R-CORE |
| INLUP_01667 | R-CORE |
| INLUP_01668 | R-CORE |
| INLUP_01669 | R-CORE |
| INLUP_01670 | R-CORE |
| INLUP_01671 | R-CORE |
| INLUP_01672 | R-CORE |
| INLUP_01673 | R-CORE |
| INLUP_01674 | R-CORE |
| INLUP_01675 | R-CORE |
| INLUP_01676 | R-CORE |
| INLUP_01677 | R-CORE |
| INLUP_01678 | R-CORE |
| INLUP_01679 | R-CORE |
| INLUP_01680 | R-CORE |
| INLUP_01681 | R-CORE |
| INLUP_01682 | R-CORE |
| INLUP_01683 | R-CORE |
| INLUP_01684 | R-CORE |
| INLUP_01685 | R-CORE |
| INLUP_01686 | R-CORE |
| INLUP_01687 | R-CORE |
| INLUP_01688 | R-CORE |
| INLUP_01689 | R-CORE |
| INLUP_01690 | R-CORE |
| INLUP_01691 | R-CORE |
| INLUP_01692 | R-CORE |
| INLUP_01693 | R-CORE |
| INLUP_01694 | R-CORE |
| INLUP_01695 | R-CORE |
| INLUP_01696 | R-CORE |
| INLUP_01697 | R-CORE |
| INLUP_01698 | R-CORE |
| INLUP_01699 | R-CORE |
| INLUP_01700 | R-CORE |
| INLUP_01701 | R-CORE |
| INLUP_01702 | R-CORE |
| INLUP_01703 | R-CORE |
| INLUP_01704 | R-CORE |
| INLUP_01705 | R-CORE |
| INLUP_01706 | R-CORE |
| INLUP_01707 | R-CORE |
| INLUP_01708 | R-CORE |
| INLUP_01709 | R-CORE |
| INLUP_01710 | R-CORE |
| INLUP_01711 | R-CORE |
| INLUP_01712 | R-CORE |
| INLUP_01713 | R-CORE |
| INLUP_01714 | R-CORE |
| INLUP_01715 | R-CORE |
| INLUP_01716 | R-CORE |
| INLUP_01717 | R-CORE |

| | |
|---|---|
| INLUP_01718 | R-CORE |
| INLUP_01719 | R-CORE |
| INLUP_01720 | R-CORE |
| INLUP_01721 | R-CORE |
| INLUP_01722 | R-CORE |
| INLUP_01723 | R-CORE |
| INLUP_01724 | R-CORE |
| INLUP_01725 | R-CORE |
| INLUP_01726 | R-CORE |
| INLUP_01727 | R-CORE |
| INLUP_01728 | R-CORE |
| INLUP_01729 | R-CORE |
| INLUP_01730 | R-CORE |
| INLUP_01731 | R-CORE |
| INLUP_01732 | R-CORE |
| INLUP_01733 | R-CORE |
| INLUP_01734 | R-CORE |
| INLUP_01735 | R-CORE |
| INLUP_01736 | R-CORE |
| INLUP_01737 | R-CORE |
| INLUP_01738 | R-CORE |
| INLUP_01739 | R-CORE |
| INLUP_01740 | R-CORE |
| INLUP_01741 | R-CORE |
| INLUP_01742 | R-CORE |
| INLUP_01743 | R-CORE |
| INLUP_01744 | R-CORE |
| INLUP_01745 | R-CORE |
| INLUP_01746 | R-CORE |
| INLUP_01747 | R-CORE |
| INLUP_01748 | R-CORE |
| INLUP_01749 | R-CORE |
| INLUP_01750 | R-CORE |
| INLUP_01751 | R-CORE |
| INLUP_01752 | R-CORE |
| INLUP_01753 | R-CORE |
| INLUP_01754 | R-CORE |
| INLUP_01755 | R-CORE |
| INLUP_01756 | R-CORE |
| INLUP_01757 | R-CORE |
| INLUP_01758 | R-CORE |
| INLUP_01759 | R-CORE |
| INLUP_01760 | R-CORE |
| INLUP_01761 | R-CORE |
| INLUP_01762 | R-CORE |
| INLUP_01763 | R-CORE |
| INLUP_01764 | R-CORE |
| INLUP_01765 | R-CORE |
| INLUP_01766 | R-CORE |
| INLUP_01767 | R-CORE |
| INLUP_01768 | R-CORE |
| INLUP_01769 | R-CORE |

| | |
|---|---|
| INLUP_01770 | R-CORE |
| INLUP_01771 | R-CORE |
| INLUP_01772 | R-CORE |
| INLUP_01773 | R-CORE |
| INLUP_01774 | R-CORE |
| INLUP_01775 | R-CORE |
| INLUP_01776 | R-CORE |
| INLUP_01777 | R-CORE |
| INLUP_01778 | R-CORE |
| INLUP_01779 | R-CORE |
| INLUP_01780 | R-CORE |
| INLUP_01781 | R-CORE |
| INLUP_01782 | R-CORE |
| INLUP_01783 | R-CORE |
| INLUP_01784 | R-CORE |
| INLUP_01785 | R-CORE |
| INLUP_01786 | R-CORE |
| INLUP_01787 | R-CORE |
| INLUP_01788 | R-CORE |
| INLUP_01789 | R-CORE |
| INLUP_01790 | R-CORE |
| INLUP_01791 | R-CORE |
| INLUP_01792 | R-CORE |
| INLUP_01793 | R-CORE |
| INLUP_01794 | R-CORE |
| INLUP_01795 | R-CORE |
| INLUP_01796 | R-CORE |
| INLUP_01797 | R-CORE |
| INLUP_01798 | R-CORE |
| INLUP_01799 | R-CORE |
| INLUP_01800 | R-CORE |
| INLUP_01801 | R-CORE |
| INLUP_01802 | R-CORE |
| INLUP_01803 | R-CORE |
| INLUP_01804 | R-CORE |
| INLUP_01805 | R-CORE |
| INLUP_01806 | R-CORE |
| INLUP_01807 | R-CORE |
| INLUP_01808 | R-CORE |
| INLUP_01809 | R-CORE |
| INLUP_01810 | R-CORE |
| INLUP_01811 | R-CORE |
| INLUP_01812 | R-CORE |
| INLUP_01813 | R-CORE |
| INLUP_01814 | R-CORE |
| INLUP_01815 | R-CORE |
| INLUP_01816 | R-CORE |
| INLUP_01817 | R-CORE |
| INLUP_01818 | R-CORE |
| INLUP_01819 | R-CORE |
| INLUP_01820 | R-CORE |
| INLUP_01821 | R-CORE |

| | |
|---|---|
| INLUP_01822 | R-CORE |
| INLUP_01823 | R-CORE |
| INLUP_01824 | R-CORE |
| INLUP_01825 | R-CORE |
| INLUP_01826 | R-CORE |
| INLUP_01827 | R-CORE |
| INLUP_01828 | R-CORE |
| INLUP_01829 | R-CORE |
| INLUP_01830 | R-CORE |
| INLUP_01831 | R-CORE |
| INLUP_01832 | R-CORE |
| INLUP_01833 | R-CORE |
| INLUP_01834 | R-CORE |
| INLUP_01835 | R-CORE |
| INLUP_01836 | R-CORE |
| INLUP_01837 | R-CORE |
| INLUP_01838 | R-CORE |
| INLUP_01839 | R-CORE |
| INLUP_01840 | R-CORE |
| INLUP_01841 | R-CORE |
| INLUP_01842 | R-CORE |
| INLUP_01843 | R-CORE |
| INLUP_01844 | R-CORE |
| INLUP_01845 | R-CORE |
| INLUP_01846 | R-CORE |
| INLUP_01847 | R-CORE |
| INLUP_01848 | R-CORE |
| INLUP_01849 | R-CORE |
| INLUP_01850 | R-CORE |
| INLUP_01851 | R-CORE |
| INLUP_01852 | R-CORE |
| INLUP_01853 | R-CORE |
| INLUP_01854 | R-CORE |
| INLUP_01855 | R-CORE |
| INLUP_01856 | R-CORE |
| INLUP_01857 | R-CORE |
| INLUP_01858 | R-CORE |
| INLUP_01859 | R-CORE |
| INLUP_01860 | R-CORE |
| INLUP_01861 | R-CORE |
| INLUP_01862 | R-CORE |
| INLUP_01863 | R-CORE |
| INLUP_01864 | R-CORE |
| INLUP_01865 | R-CORE |
| INLUP_01866 | R-CORE |
| INLUP_01867 | R-CORE |
| INLUP_01868 | R-CORE |
| INLUP_01869 | R-CORE |
| INLUP_01870 | R-CORE |
| INLUP_01871 | R-CORE |
| INLUP_01872 | R-CORE |
| INLUP_01873 | R-CORE |

| | |
|---|---|
| INLUP_01874 | R-CORE |
| INLUP_01875 | R-CORE |
| INLUP_01876 | R-CORE |
| INLUP_01877 | R-CORE |
| INLUP_01878 | R-CORE |
| INLUP_01879 | R-CORE |
| INLUP_01880 | R-CORE |
| INLUP_01881 | R-CORE |
| INLUP_01882 | R-CORE |
| INLUP_01883 | R-CORE |
| INLUP_01884 | R-CORE |
| INLUP_01885 | R-CORE |
| INLUP_01886 | R-CORE |
| INLUP_01887 | R-CORE |
| INLUP_01888 | R-CORE |
| INLUP_01889 | R-CORE |
| INLUP_01890 | R-CORE |
| INLUP_01891 | R-CORE |
| INLUP_01892 | R-CORE |
| INLUP_01893 | R-CORE |
| INLUP_01894 | R-CORE |
| INLUP_01895 | R-CORE |
| INLUP_01896 | R-CORE |
| INLUP_01897 | R-CORE |
| INLUP_01898 | R-CORE |
| INLUP_01899 | R-CORE |
| INLUP_01900 | R-CORE |
| INLUP_01901 | R-CORE |
| INLUP_01902 | R-CORE |
| INLUP_01903 | R-CORE |
| INLUP_01904 | R-CORE |
| INLUP_01905 | R-CORE |
| INLUP_01906 | R-CORE |
| INLUP_01907 | R-CORE |
| INLUP_01908 | R-CORE |
| INLUP_01909 | R-CORE |
| INLUP_01910 | R-CORE |
| INLUP_01911 | R-CORE |
| INLUP_01912 | R-CORE |
| INLUP_01913 | R-CORE |
| INLUP_01914 | R-CORE |
| INLUP_01915 | R-CORE |
| INLUP_01916 | R-CORE |
| INLUP_01917 | R-CORE |
| INLUP_01918 | R-CORE |
| INLUP_01919 | R-CORE |
| INLUP_01920 | R-CORE |
| INLUP_01921 | R-CORE |
| INLUP_01922 | R-CORE |
| INLUP_01923 | R-CORE |
| INLUP_01924 | R-CORE |
| INLUP_01925 | R-CORE |

| | |
|---|---|
| INLUP_01926 | R-CORE |
| INLUP_01927 | R-CORE |
| INLUP_01928 | R-CORE |
| INLUP_01929 | R-CORE |
| INLUP_01930 | R-CORE |
| INLUP_01931 | R-CORE |
| INLUP_01932 | R-CORE |
| INLUP_01933 | R-CORE |
| INLUP_01934 | R-CORE |
| INLUP_01935 | R-CORE |
| INLUP_01936 | R-CORE |
| INLUP_01937 | R-CORE |
| INLUP_01938 | R-CORE |
| INLUP_01939 | R-CORE |
| INLUP_01940 | R-CORE |
| INLUP_01941 | R-CORE |
| INLUP_01942 | R-CORE |
| INLUP_01943 | R-CORE |
| INLUP_01944 | R-CORE |
| INLUP_01945 | R-CORE |
| INLUP_01946 | R-CORE |
| INLUP_01947 | R-CORE |
| INLUP_01948 | R-CORE |
| INLUP_01949 | R-CORE |
| INLUP_01950 | R-CORE |
| INLUP_01951 | R-CORE |
| INLUP_01952 | R-CORE |
| INLUP_01953 | R-CORE |
| INLUP_01954 | R-CORE |
| INLUP_01955 | R-CORE |
| INLUP_01956 | R-CORE |
| INLUP_01957 | R-CORE |
| INLUP_01958 | R-CORE |
| INLUP_01959 | R-CORE |
| INLUP_01960 | R-CORE |
| INLUP_01961 | R-CORE |
| INLUP_01962 | R-CORE |
| INLUP_01963 | R-CORE |
| INLUP_01964 | R-CORE |
| INLUP_01965 | R-CORE |
| INLUP_01966 | R-CORE |
| INLUP_01967 | R-CORE |
| INLUP_01968 | R-CORE |
| INLUP_01969 | R-CORE |
| INLUP_01970 | R-CORE |
| INLUP_01971 | R-CORE |
| INLUP_01972 | R-CORE |
| INLUP_01973 | R-CORE |
| INLUP_01974 | R-CORE |
| INLUP_01975 | R-CORE |
| INLUP_01976 | R-CORE |
| INLUP_01977 | R-CORE |

| | |
|---|---|
| INLUP_01978 | R-CORE |
| INLUP_01979 | R-CORE |
| INLUP_01980 | R-CORE |
| INLUP_01981 | R-CORE |
| INLUP_01982 | R-CORE |
| INLUP_01983 | R-CORE |
| INLUP_01984 | R-CORE |
| INLUP_01985 | R-CORE |
| INLUP_01986 | R-CORE |
| INLUP_01987 | R-CORE |
| INLUP_01988 | R-CORE |
| INLUP_01989 | R-CORE |
| INLUP_01990 | R-CORE |
| INLUP_01991 | R-CORE |
| INLUP_01992 | R-CORE |
| INLUP_01993 | R-CORE |
| INLUP_01994 | R-CORE |
| INLUP_01995 | R-CORE |
| INLUP_01996 | R-CORE |
| INLUP_01997 | R-CORE |
| INLUP_01998 | R-CORE |
| INLUP_01999 | R-CORE |
| INLUP_02000 | R-CORE |
| INLUP_02001 | R-CORE |
| INLUP_02002 | R-CORE |
| INLUP_02003 | R-CORE |
| INLUP_02004 | R-CORE |
| INLUP_02005 | R-CORE |
| INLUP_02006 | R-CORE |
| INLUP_02007 | R-CORE |
| INLUP_02008 | R-CORE |
| INLUP_02009 | R-CORE |
| INLUP_02010 | R-CORE |
| INLUP_02011 | R-CORE |
| INLUP_02012 | R-CORE |
| INLUP_02013 | R-CORE |
| INLUP_02014 | R-CORE |
| INLUP_02015 | R-CORE |
| INLUP_02016 | R-CORE |
| INLUP_02017 | R-CORE |
| INLUP_02018 | R-CORE |
| INLUP_02019 | R-CORE |
| INLUP_02020 | R-CORE |
| INLUP_02021 | R-CORE |
| INLUP_02022 | R-CORE |
| INLUP_02023 | R-CORE |
| INLUP_02024 | R-CORE |
| INLUP_02025 | R-CORE |
| INLUP_02026 | R-CORE |
| INLUP_02027 | R-CORE |
| INLUP_02028 | R-CORE |
| INLUP_02029 | R-CORE |

| | |
|---|---|
| INLUP_02030 | R-CORE |
| INLUP_02031 | R-CORE |
| INLUP_02032 | R-CORE |
| INLUP_02033 | R-CORE |
| INLUP_02034 | R-CORE |
| INLUP_02035 | R-CORE |
| INLUP_02036 | R-CORE |
| INLUP_02037 | R-CORE |
| INLUP_02038 | R-CORE |
| INLUP_02039 | R-CORE |
| INLUP_02040 | R-CORE |
| INLUP_02041 | R-CORE |
| INLUP_02042 | R-CORE |
| INLUP_02043 | R-CORE |
| INLUP_02044 | R-CORE |
| INLUP_02045 | R-CORE |
| INLUP_02046 | R-CORE |
| INLUP_02047 | R-CORE |
| INLUP_02048 | R-CORE |
| INLUP_02049 | R-CORE |
| INLUP_02050 | R-CORE |
| INLUP_02051 | R-CORE |
| INLUP_02052 | R-CORE |
| INLUP_02053 | R-CORE |
| INLUP_02054 | R-CORE |
| INLUP_02055 | R-CORE |
| INLUP_02056 | R-CORE |
| INLUP_02057 | R-CORE |
| INLUP_02058 | R-CORE |
| INLUP_02059 | R-CORE |
| INLUP_02060 | R-CORE |
| INLUP_02061 | R-CORE |
| INLUP_02062 | R-CORE |
| INLUP_02063 | R-CORE |
| INLUP_02064 | R-CORE |
| INLUP_02065 | R-CORE |
| INLUP_02066 | R-CORE |
| INLUP_02067 | R-CORE |
| INLUP_02068 | R-CORE |
| INLUP_02069 | R-CORE |
| INLUP_02070 | R-CORE |
| INLUP_02071 | R-CORE |
| INLUP_02072 | R-CORE |
| INLUP_02073 | R-CORE |
| INLUP_02074 | R-CORE |
| INLUP_02075 | R-CORE |
| INLUP_02076 | R-CORE |
| INLUP_02077 | R-CORE |
| INLUP_02078 | R-CORE |
| INLUP_02079 | R-CORE |
| INLUP_02080 | R-CORE |
| INLUP_02081 | R-CORE |

| | |
|---|---|
| INLUP_02082 | R-CORE |
| INLUP_02083 | R-CORE |
| INLUP_02084 | R-CORE |
| INLUP_02085 | R-CORE |
| INLUP_02086 | R-CORE |
| INLUP_02087 | R-CORE |
| INLUP_02088 | R-CORE |
| INLUP_02089 | R-CORE |
| INLUP_02090 | R-CORE |
| INLUP_02091 | R-CORE |
| INLUP_02092 | R-CORE |
| INLUP_02093 | R-CORE |
| INLUP_02094 | R-CORE |
| INLUP_02095 | R-CORE |
| INLUP_02096 | R-CORE |
| INLUP_02097 | R-CORE |
| INLUP_02098 | R-CORE |
| INLUP_02099 | R-CORE |
| INLUP_02100 | R-CORE |
| INLUP_02101 | R-CORE |
| INLUP_02102 | R-CORE |
| INLUP_02103 | R-CORE |
| INLUP_02104 | R-CORE |
| INLUP_02105 | R-CORE |
| INLUP_02106 | R-CORE |
| INLUP_02107 | R-CORE |
| INLUP_02108 | R-CORE |
| INLUP_02109 | R-CORE |
| INLUP_02110 | R-CORE |
| INLUP_02111 | R-CORE |
| INLUP_02112 | R-CORE |
| INLUP_02113 | R-CORE |
| INLUP_02114 | R-CORE |
| INLUP_02115 | R-CORE |
| INLUP_02116 | R-CORE |
| INLUP_02117 | R-CORE |
| INLUP_02118 | R-CORE |
| INLUP_02119 | R-CORE |
| INLUP_02120 | R-CORE |
| INLUP_02121 | R-CORE |
| INLUP_02122 | R-CORE |
| INLUP_02123 | R-CORE |
| INLUP_02124 | R-CORE |
| INLUP_02125 | R-CORE |
| INLUP_02126 | R-CORE |
| INLUP_02127 | R-CORE |
| INLUP_02128 | R-CORE |
| INLUP_02129 | R-CORE |
| INLUP_02130 | R-CORE |
| INLUP_02131 | R-CORE |
| INLUP_02132 | R-CORE |
| INLUP_02133 | R-CORE |

| | |
|---|---|
| INLUP_02134 | R-CORE |
| INLUP_02135 | R-CORE |
| INLUP_02136 | R-CORE |
| INLUP_02137 | R-CORE |
| INLUP_02138 | R-CORE |
| INLUP_02139 | R-CORE |
| INLUP_02140 | R-CORE |
| INLUP_02141 | R-CORE |
| INLUP_02142 | R-CORE |
| INLUP_02143 | R-CORE |
| INLUP_02144 | R-CORE |
| INLUP_02145 | R-CORE |
| INLUP_02146 | R-CORE |
| INLUP_02147 | R-CORE |
| INLUP_02148 | R-CORE |
| INLUP_02149 | R-CORE |
| INLUP_02150 | R-CORE |
| INLUP_02151 | R-CORE |
| INLUP_02152 | R-CORE |
| INLUP_02153 | R-CORE |
| INLUP_02154 | R-CORE |
| INLUP_02155 | R-CORE |
| INLUP_02156 | R-CORE |
| INLUP_02157 | R-CORE |
| INLUP_02158 | R-CORE |
| INLUP_02159 | R-CORE |
| INLUP_02160 | R-CORE |
| INLUP_02161 | R-CORE |
| INLUP_02162 | R-CORE |
| INLUP_02163 | R-CORE |
| INLUP_02164 | R-CORE |
| INLUP_02165 | R-CORE |
| INLUP_02166 | R-CORE |
| INLUP_02167 | R-CORE |
| INLUP_02168 | R-CORE |
| INLUP_02169 | R-CORE |
| INLUP_02170 | R-CORE |
| INLUP_02171 | R-CORE |
| INLUP_02172 | R-CORE |
| INLUP_02173 | R-CORE |
| INLUP_02174 | R-CORE |
| INLUP_02175 | R-CORE |
| INLUP_02176 | R-CORE |
| INLUP_02177 | R-CORE |
| INLUP_02178 | R-CORE |
| INLUP_02179 | R-CORE |
| INLUP_02180 | R-CORE |
| INLUP_02181 | R-CORE |
| INLUP_02182 | R-CORE |
| INLUP_02183 | R-CORE |
| INLUP_02184 | R-CORE |
| INLUP_02185 | R-CORE |

| | |
|---|---|
| INLUP_02186 | R-CORE |
| INLUP_02187 | R-CORE |
| INLUP_02188 | R-CORE |
| INLUP_02189 | R-CORE |
| INLUP_02190 | R-CORE |
| INLUP_02191 | R-CORE |
| INLUP_02192 | R-CORE |
| INLUP_02193 | R-CORE |
| INLUP_02194 | R-CORE |
| INLUP_02195 | R-CORE |
| INLUP_02196 | R-CORE |
| INLUP_02197 | R-CORE |
| INLUP_02198 | R-CORE |
| INLUP_02199 | R-CORE |
| INLUP_02200 | R-CORE |
| INLUP_02201 | R-CORE |
| INLUP_02202 | R-CORE |
| INLUP_02203 | R-CORE |
| INLUP_02204 | R-CORE |
| INLUP_02205 | R-CORE |
| INLUP_02206 | R-CORE |
| INLUP_02207 | R-CORE |
| INLUP_02208 | R-CORE |
| INLUP_02209 | R-CORE |
| INLUP_02210 | R-CORE |
| INLUP_02211 | R-CORE |
| INLUP_02212 | R-CORE |
| INLUP_02213 | R-CORE |
| INLUP_02214 | R-CORE |
| INLUP_02215 | R-CORE |
| INLUP_02216 | R-CORE |
| INLUP_02217 | R-CORE |
| INLUP_02218 | R-CORE |
| INLUP_02219 | R-CORE |
| INLUP_02220 | R-CORE |
| INLUP_02221 | R-CORE |
| INLUP_02222 | R-CORE |
| INLUP_02223 | R-CORE |
| INLUP_02224 | R-CORE |
| INLUP_02225 | R-CORE |
| INLUP_02226 | R-CORE |
| INLUP_02227 | R-CORE |
| INLUP_02228 | R-CORE |
| INLUP_02229 | R-CORE |
| INLUP_02230 | R-CORE |
| INLUP_02231 | R-CORE |
| INLUP_02232 | R-CORE |
| INLUP_02233 | R-CORE |
| INLUP_02234 | R-CORE |
| INLUP_02235 | R-CORE |
| INLUP_02236 | R-CORE |
| INLUP_02237 | R-CORE |

| | |
|---|---|
| INLUP_02238 | R-CORE |
| INLUP_02239 | R-CORE |
| INLUP_02240 | R-CORE |
| INLUP_02241 | R-CORE |
| INLUP_02242 | R-CORE |
| INLUP_02243 | R-CORE |
| INLUP_02244 | R-CORE |
| INLUP_02245 | R-CORE |
| INLUP_02246 | R-CORE |
| INLUP_02247 | R-CORE |
| INLUP_02248 | R-CORE |
| INLUP_02249 | R-CORE |
| INLUP_02250 | R-CORE |
| INLUP_02251 | R-CORE |
| INLUP_02252 | R-CORE |
| INLUP_02253 | R-CORE |
| INLUP_02254 | R-CORE |
| INLUP_02255 | R-CORE |
| INLUP_02256 | R-CORE |
| INLUP_02257 | R-CORE |
| INLUP_02258 | R-CORE |
| INLUP_02259 | R-CORE |
| INLUP_02260 | R-CORE |
| INLUP_02261 | R-CORE |
| INLUP_02262 | R-CORE |
| INLUP_02263 | R-CORE |
| INLUP_02264 | R-CORE |
| INLUP_02265 | R-CORE |
| INLUP_02266 | R-CORE |
| INLUP_02267 | R-CORE |
| INLUP_02268 | R-CORE |
| INLUP_02269 | R-CORE |
| INLUP_02270 | R-CORE |
| INLUP_02271 | R-CORE |
| INLUP_02272 | R-CORE |
| INLUP_02273 | R-CORE |
| INLUP_02274 | R-CORE |
| INLUP_02275 | R-CORE |
| INLUP_02276 | R-CORE |
| INLUP_02277 | R-CORE |
| INLUP_02278 | R-CORE |
| INLUP_02279 | R-CORE |
| INLUP_02280 | R-CORE |
| INLUP_02281 | R-CORE |
| INLUP_02282 | R-CORE |
| INLUP_02283 | R-CORE |
| INLUP_02284 | R-CORE |
| INLUP_02285 | R-CORE |
| INLUP_02286 | R-CORE |
| INLUP_02287 | R-CORE |
| INLUP_02288 | R-CORE |
| INLUP_02289 | R-CORE |

| | |
|---|---|
| INLUP_02290 | R-CORE |
| INLUP_02291 | R-CORE |

udy

| Common Name/ cultivar name | Accession/ line code | COUNTRY |
|---|---|---|
| HINOJOSO DE DUERO-1 | PL95001 | ESP |
| MULTO LUPA | PL95002 | ESP |
| SOUTH SPAIN-1 | PL95003 | ESP |
| SOLDANA | PL95006 | ESP |
| MANSILLA DE LOS MULAS | PL95007 | ESP |
| ALAMANZA | PL95008 | ESP |
| SIERRA DE FRANCIA | PL95009 | ESP |
| CASTILIA | PL95010 | ESP |
| MAGARRAZ-2 | PL95012 | ESP |
| MAGARRAZ-3 | PL95013 | ESP |
| MAGARRAZ-4 | PL95014 | ESP |
| BANOBAREZ | PL95016 | ESP |
| OEIRAS-551/5 | PL95019 | PRT |
| OEIRAS-802/15 | PL95020 | PRT |
| OEIRAS-804/4 | PL95021 | PRT |
| OEIRAS-930/3 | PL95023 | PRT |
| KAIRO-1 | PL95024 | EGY |
| KAIRO-2 | PL95025 | EGY |
| KAIRO-3 | PL95026 | EGY |
| KAIRO-4 | PL95027 | EGY |
| MADRID-1 | PL95028 | ESP |
| POPULATION-8002 | PL95029 | ESP |
| POPULATION-8004 | PL95031 | ESP |
| POPULATION-8005 | PL95032 | ESP |
| POPULATION-8006 | PL95033 | ESP |
| POPULATION-8022 | PL95042 | ESP |
| POPULATION-8023 | PL95043 | ESP |
| POPULATION-8030 | PL95047 | ESP |
| POPULATION-8032 | PL95049 | ESP |
| POPULATION-8038 | PL95053 | ESP |
| POPULATION-8062 | PL95064 | ESP |
| POPULATION-8067 | PL95068 | ESP |
| POPULATION-8068 | PL95069 | ESP |
| POPULATION B-265/79 | PL95070 | ESP |
| POPULATION B-266/79 | PL95071 | ESP |
| POPULATION B-267/79 | PL95072 | ESP |
| POPULATION B-273/79 | PL95073 | ESP |
| POPULATION B-275/79 | PL95074 | ESP |
| POPULATION B-295/79 | PL95075 | ESP |
| POPULATION B-296/76 | PL95076 | ESP |
| POPULATION B-297/79 | PL95077 | ESP |
| POPULATION B-300/79 | PL95078 | ESP |
| POPULATION B-2005/80 | PL95079 | ESP |
| POPULATION B-2007/80 | PL95080 | ESP |
| POPULATION B-2012/80 | PL95082 | ESP |
| POPULATION B-2013/83 | PL95083 | ESP |
| EL HARRACH-1 | PL95084 | DZA |
| EL HARRACH-2 | PL95085 | DZA |
| EL HARRACH-3 | PL95086 | DZA |
| EL HARRACH-4 | PL95087 | DZA |

| | | |
|---|---|---|
| HOFMANN KAIRO | PL95088 | EGY |
| POPULATION-1 | PL95089 | EGY |
| KRETA | PL95090 | GRC |
| PELOPONNES | PL95091 | GRC |
| WUHOLIES KRETA | PL95092 | GRC |
| ASTRA | PL95093 | ESP |
| RIMPAUS FRÜHE | PL95094 | DEU |
| AZORSKI | PL95095 | PRT |
| | PL95097 | PRT |
| SATMAREAN | PL95098 | PRT |
| SACAVEM | PL95099 | PRT |
| CALABRIA | PL95101 | ITA |
| LA-39 | PL95104 | ESP |
| LA-108 | PL95113 | FRA |
| POPULATION-775 | PL95122 | GRC |
| POPULATION-780 | PL95124 | GRC |
| POPULATION-84094 | PL95129 | ESP |
| VIR LENINGR.-490 | PL95133 | ETH |
| POPULATION-180 | PL95136 | ESP |
| AL-18 | PL95145 | ESP |
| AL-25 | PL95150 | ESP |
| R-61 | PL95156 | ESP |
| R-114 | PL95157 | ESP |
| R-243 | PL95160 | ESP |
| SF-479 | PL95166 | ESP |
| R-491 | PL95168 | ESP |
| R-84141 | PL95175 | ESP |
| [(MALYXWAT)XWAT]XMUTANT SELF | PL95181 | POL |
| BGRC-3907 | PL95183 | ITA |
| BGRC-3915 | PL95191 | ITA |
| BGRC-3916 | PL95192 | ITA |
| BGRC-48656 | PL95202 | FRA |
| POPULACJA Z SYRII | PL95203 | SYR |
| PI-24472 | PL95209 | BRA |
| IFLU-29 | PL95214 | SYR |
| IFLU-33 | PL95215 | SYR |
| L-229-N | PL95216 | ZAF |
| TR-5 | PL95223 | TUR |
| TR-7 | PL95224 | TUR |
| TR-12 | PL95225 | TUR |
| TR-22 | PL95226 | TUR |
| POP.WENEZUELA | PL95229 | VEN |
| L.46-10 | PL95230 | USA |
| POP.Z SYRII | PL95231 | SYR |
| L.GRAECUS X MUT.TOPLESSXHETMAN F | PL95237 | POL |
| BAC | PL95238 | POL |
| POP.2053 | PL95241 | ITA |
| POP.2023 | PL95243 | DZA |
| LUPINI BEAN | PL95245 | GBR |
| POP.2078 | PL95246 | ETH |
| POP.SETUBAL | PL95266 | PRT |
| ASCAR | PL95270 | FRA |

| | | |
|---|---|---|
| POP.TENERIFE 1 | PL95271 | ESP |
| POP.TENERIFE 2 | PL95272 | ESP |
| NÄHRQUELL | PL95278 | DEU |
| SHIENFIELD GARD. | PL95401 | GBR |
| R-6012 | PL95402 | POL |
| R-6020 | PL95404 | POL |
| R-933 | PL95406 | POL |
| TREMOCO DOCE | PL95407 | PRT |
| BLANKA | PL95408 | DEU |
| KIEVSKIJ SKOROSPELIJ | PL95412 | UKR |
| KIJEVSKIJ MUTANT | PL95413 | UKR |
| BIALORUS-1 | PL95414 | UKR |
| BIALORUS-3 | PL95416 | UKR |
| KALINA | PL95419 | POL |
| HETMAN | PL95420 | POL |
| START | PL95422 | SUN |
| AMIGA | PL95423 | CHL |
| DNIEPR | PL95433 | SUN |
| HETMANXSTART | PL95440 | POL |
| WAT X L.GRAECUS BOISS. | PL95441 | POL |
| LUBLANC | PL95443 | FRA |
| HAMBURG | PL95450 | ZAF |
| NÄHRQUELL LUP 235/82 | PL95458 | DEU |
| AZURNYJ | PL95468 | BLR |
| R6902 X EL HARRACH 2 | PL95470 | POL |
| KATON | PL95514 | POL |
| FELI | PL95531 | SVK |
| SOUTH SPAIN-2 | PL95004 | ESP |
| HINOJOSO DE DUERO-2 | PL95005 | ESP |
| MAGARRAZ-1 | PL95011 | ESP |
| SAN FELICES | PL95015 | ESP |
| SAN FELICES-2 | PL95017 | ESP |
| DUERO DE ARABA | PL95018 | ESP |
| OEIRAS-893/7 | PL95022 | PRT |
| POPULATION-8009 | PL95034 | ESP |
| POPULATION-8010 | PL95035 | ESP |
| POPULATION-8011 | PL95036 | ESP |
| POPULATION-8018 | PL95038 | ESP |
| POPULATION-8020 | PL95040 | ESP |
| POPULATION-8021 | PL95041 | ESP |
| POPULATION-8027 | PL95044 | ESP |
| POPULATION-8031 | PL95048 | ESP |
| POPULATION-8035 | PL95050 | ESP |
| POPULATION-8036 | PL95051 | ESP |
| POPULATION-8037 | PL95052 | ESP |
| POPULATION-8039 | PL95054 | ESP |
| POPULATION-8049 | PL95057 | ESP |
| POPULATION-8053 | PL95059 | ESP |
| POPULATION-8054 | PL95060 | ESP |
| POPULATION-8056 | PL95061 | ESP |
| POPULATION-8057 | PL95062 | ESP |
| POPULATION-8063 | PL95065 | ESP |

| | | |
|---|---|---|
| POPULATION-8065 | PL95066 | ESP |
| POPULATION-8066 | PL95067 | ESP |
| POPULATION B-2010/80 | PL95081 | ESP |
| SHARKIA-13 | PL95096 | PRT |
| ODORE MARINA | PL95100 | ITA |
| LA-31 SAN FELICES | PL95102 | ESP |
| LA-20 | PL95103 | ESP |
| LA-40 | PL95105 | ESP |
| LA-46 | PL95106 | ESP |
| POPULATION-800 | PL95107 | GRC |
| POPULATION-84002 | PL95108 | ESP |
| POPULATION-84003 | PL95109 | ESP |
| POPULATION-84050 | PL95110 | ESP |
| LA-109 | PL95114 | FRA |
| LA-01 BANOBAREZ | PL95115 | ESP |
| MUTANT CIENKOŚCIENNY | PL95116 | ESP |
| POPULATION-733 | PL95118 | GRC |
| POPULATION-755 | PL95119 | GRC |
| POPULATION-769 | PL95120 | GRC |
| POPULATION-757 | PL95123 | GRC |
| POPULATION-877 | PL95125 | GRC |
| POPULATION-84053 | PL95126 | ESP |
| POPULATION-84075 | PL95128 | ESP |
| POPULATION-84152 | PL95130 | ESP |
| TREMOCO GORZKI | PL95132 | PRT |
| POPULATION-182 | PL95137 | ESP |
| POPULATION-187 | PL95138 | ESP |
| POPULATION-188 | PL95139 | ESP |
| POPULATION-189 | PL95140 | ESP |
| POPULATION-197 | PL95142 | ESP |
| POPULATION-199 | PL95143 | ESP |
| M.MUT.Z HISZP.(TOPLESS) | PL95144 | ESP |
| AL-19 | PL95146 | ESP |
| AL-20 | PL95147 | ESP |
| AL-21 | PL95148 | ESP |
| AL-22 | PL95149 | ESP |
| AL-26 | PL95151 | ESP |
| AL-27 | PL95152 | ESP |
| AL-28 | PL95153 | ESP |
| R-32 | PL95155 | ESP |
| R-243 | PL95161 | ESP |
| SF-386 | PL95162 | ESP |
| SF-387 | PL95163 | ESP |
| SF-479 | PL95167 | ESP |
| WTD 6020XL.GRAECUS BOISS | PL95173 | POL |
| R-84057 | PL95174 | ESP |
| WTD 6020 | PL95176 | POL |
| MUTANT SELF COMPLETING | PL95177 | POL |
| MUTANT SELF COMPLETINGXL.GRA | PL95179 | POL |
| MUTANT SELF COMPLETINGXL.VAVILOVII | PL95180 | POL |
| BGRC-3908 | PL95184 | ITA |
| BGRC-3909 | PL95185 | ITA |

| | | |
|---|---|---|
| BGRC-3910 | PL95186 | ITA |
| BGRC-3911 | PL95187 | ITA |
| BGRC-3912 | PL95188 | ITA |
| BGRC-3913 | PL95189 | ITA |
| BGRC-3914 | PL95190 | ITA |
| BGRC-3917 | PL95193 | ITA |
| BGRC-3918 | PL95194 | ITA |
| BGRC-3919 | PL95195 | ITA |
| BGRC-3920 | PL95196 | ITA |
| BGRC-3921 | PL95197 | ITA |
| BGRC-3922 | PL95198 | ITA |
| BGRC-3923 | PL95199 | ITA |
| BGRC-3924 | PL95200 | ITA |
| BGRC-3927 | PL95201 | DZA |
| FAO NR-69946 | PL95204 | ITA |
| FAO NR-69947 | PL95205 | ITA |
| FAO NR-69948 | PL95206 | ITA |
| FAO NR-69948/GROSSA AMINA/ | PL95207 | ITA |
| ASTRA | PL95208 | CHL |
| LA-231 | PL95210 | FRA |
| TURMUS BALADY GADIM,DRUZE L.R. | PL95211 | PAL |
| TURMUS BALADY GADIM,L.R.M.-2738 | PL95212 | JOR |
| TURMUS BALADY L.R.M.-2675 | PL95213 | JOR |
| BUSH TYPE PI 250094 | PL95218 | EGY |
| GIZA-2 | PL95219 | EGY |
| FAM 120 | PL95220 | TUR |
| TR-9 | PL95222 | TUR |
| L.46-4 | PL95227 | USA |
| L.413 | PL95232 | LTU |
| M-28 NOWY | PL95233 | Unknown |
| P.21525 | PL95234 | CHL |
| TURMUS | PL95236 | TUR |
| ALTRAMUZ CHOCHO 1 | PL95239 | ESP |
| ALTRAMUZ CHOCHO 2 | PL95240 | ESP |
| POP.2057 | PL95242 | ITA |
| POP.2027 | PL95244 | DZA |
| SIEBACHER RED | PL95247 | ZAF |
| NÄHRQUELL | PL95248 | DEU |
| POP.2738 | PL95249 | PAL |
| POP.2579 | PL95251 | PAL |
| POP.2579B | PL95252 | PAL |
| EGIPT-5 | PL95253 | EGY |
| ARKA-10 | PL95268 | USA |
| BG-9787 | PL95269 | PAL |
| BG-18829 | PL95273 | GRC |
| BGR 6303 | PL95274 | GRC |
| BGR 6305 | PL95275 | PAL |
| SIEBACHER RED | PL95276 | PRT |
| BGR 6325 | PL95277 | ITA |
| WAT | PL95403 | POL |
| KALI | PL95405 | POL |
| NEUTRA | PL95409 | DEU |

| | | |
|---|---|---|
| KRAFTQUELL | PL95410 | DEU |
| SWEET WHITE | PL95411 | ZAF |
| BIALORUS-2 | PL95415 | UKR |
| BIALORUS-4 | PL95417 | UKR |
| BIALORUS-5 | PL95418 | UKR |
| R-6001 | PL95421 | POL |
| KALI X LAJU BIAŁY-1 | PL95424 | POL |
| SEL.ECOT. X LA-109/2 | PL95426 | ESP |
| SEL.ECOT. X LA-109/1 | PL95427 | Unknown |
| PFLUGS GELA | PL95428 | DEU |
| NÄHRQUELL | PL95429 | DEU |
| RIMPAUS FRÜHE SÜSSE WEISSLUP | PL95430 | DEU |
| VOLODIA | PL95431 | DEU |
| PRIMORSKIJ | PL95432 | UKR |
| LOTOS | PL95434 | SUN |
| SILOSNYJ | PL95439 | SUN |
| LUTOP | PL95445 | FRA |
| ESTORIL | PL95447 | PRT |
| MUSTAL | PL95448 | PRT |
| BUTTERCUP | PL95449 | ZAF |
| DETTY | PL95451 | HUN |
| WTD-286 | PL95452 | POL |
| BRGC-48655 | PL95453 | FRA |
| TOMBOWSKIJ SKOROSPIELYJ | PL95454 | SUN |
| MUTANT-28 | PL95455 | SUN |
| GORIZONT | PL95456 | SUN |
| NÄHRQUELL II-5-8 | PL95457 | DEU |
| BEST | PL95460 | PAL |
| CULTIVATED LUPINES | PL95461 | SYR |
| MULTOLUPA NOWA | PL95467 | CHL |
| WTD 393 PIKADOR | PL95469 | POL |
| ADAM | PL95471 | FRA |
| ALBAN | PL95472 | FRA |
| AMIGA | PL95475 | FRA |
| ATHOS | PL95476 | FRA |
| KIJEVSKIJ MUTANT | PL95479 | SUN |
| NELLY | PL95480 | HUN |
| BARDO | PL95481 | POL |
| PIKADOR | PL95482 | POL |
| WAT-I 0,5 SA | PL95494 | POL |
| WAT-I 1,5 SA | PL95495 | POL |
| WAT-I 2,0 SA | PL95503 | POL |
| WAT-I 2,0 SA | PL95506 | POL |
| HADMERSLEBENER EPIGON | PL95508 | DEU |
| NÄHRQUELL | PL95509 | DEU |
| HANSA | PL95510 | DEU |
| TERRE | PL95511 | ZAF |
| MJS208-1 | PL95515 | ESP |
| MJS208-2 | PL95516 | ESP |
| DAMASCUS | PL95517 | SYR |
| ILCA13665 | PL95518 | ETH |
| ILCA13673 | PL95519 | ETH |

| | | |
|---|---|---|
| ILCA13674 | PL95520 | ETH |
| ILCA13677 | PL95521 | ETH |
| FRA6706B | PL95522 | TUR |
| FRA6708B | PL95523 | TUR |
| FRA6712B | PL95524 | TUR |
| FRA6713B | PL95525 | TUR |
| LA359 | PL95526 | FRA |
| LA382 | PL95527 | EGY |
| LA384 | PL95528 | EGY |
| LA423 | PL95529 | SDN |
| BOROS | PL95530 | POL |
| POPULATION-1 | PL95601 | GRC |
| POPULATION-775 | PL95602 | GRC |
| POPULATION-743 | PL95605 | GRC |
| POPULATION-776 | PL95606 | GRC |
| POPULATION-757 | PL95607 | GRC |
| POPULATION-804 | PL95608 | GRC |
| POPULATION-1 | PL95631 | Unknown |
| POPULATION 2 | PL95632 | Unknown |
| POPULATION-774 | PL95673 | GRC |
| POPULATION-807 | PL95121 | YUG |
| | LUP 203 | GRC |
| | LUP 204 | GRC |
| | LUP 206 | GRC |
| | LUP 207 | GRC |
| | LUP 208 | GRC |
| | LUP 209 | GRC |
| | LUP 210 | GRC |
| | LUP 211 | GRC |
| | LUP 212 | GRC |
| | LUP 213 | GRC |
| | LUP 215 | GRC |
| | LUP 216 | GRC |
| | LUP 217 | GRC |
| SÜßLUPINE | LUP 220 | Unknown |
| WEIßE BITTERLUPINE | LUP 221 | Unknown |
| | LUP 222 | ITA |
| | LUP 223 | ITA |
| | LUP 224 | ITA |
| | LUP 225 | ITA |
| | LUP 226 | Unknown |
| KRAFTQUELL | LUP 227 | DEU |
| PETKUSER BITTERE WEISSLUPINE | LUP 230 | DEU |
| KERESKEDELMI | LUP 234 | HUN |
| NÄHRQUELL | LUP 235 | DEU |
| | LUP 236 | PRT |
| ZÁGREBESKÁ | LUP 237 | Unknown |
| NEUTRA | LUP 239 | DEU |
| EGYPTSKÁ | LUP 240 | Unknown |
| GYULATANYAI FEHÉRVIRÁGÚ ÉDES CSILLAGFÜRT | LUP 241 | HUN |
| | LUP 242 | PRT |
| PFLUGS GELA | LUP 243 | Unknown |

| | | |
|---|---|---|
| HANSA | LUP 244 | Unknown |
| KISVÁRDAI 1432/49 | LUP 245 | HUN |
| KISVÁRDAI FEHÉRVIRÁGÚ ÉDES CSILLAGFÜRT | LUP 246 | HUN |
| PRZEBEDOWSKI SREDNIO-WCZESNY | LUP 247 | POL |
| PRZEBEDOWSKI WCZESNY | LUP 248 | POL |
| SIEBACHER RED | LUP 249 | Unknown |
| HADMERSLEBENER ST. 387 | LUP 250 | DEU |
| PFLUGS ULTRA | LUP 251 | DEU |
| | LUP 252 | GEO |
| SWEET WHITE 7731 | LUP 253 | Unknown |
| SHARKIA 13 | LUP 254 | Unknown |
| KIEVSKIJ SKOROSPELIJ | LUP 255 | SUN |
| TURMUS (ARAB.) [VERNACULAR NAME] | LUP 256 | EGY |
| KIJEVSKIJ MUTANT | LUP 257 | UKR |
| | LUP 258 | ETH |
| | LUP 259 | ITA |
| | LUP 260 | EGY |
| | LUP 261 | ESP |
| | LUP 262 | ESP |
| Altramuz [vernacular name] | LUP 263 | ESP |
| Altramuz [vernacular name] | LUP 264 | ESP |
| | LUP 265 | ESP |
| | LUP 266 | ESP |
| | LUP 267 | ESP |
| | LUP 268 | ESP |
| | LUP 269 | ESP |
| Altramuz | LUP 270 | ESP |
| Chocho | LUP 271 | ESP |
| Chocho | LUP 272 | ESP |
| | LUP 273 | ESP |
| Altramuz [vernacular name] | LUP 274 | ESP |
| | LUP 275 | ESP |
| | LUP 276 | ESP |
| Altramuz | LUP 277 | ESP |
| | LUP 278 | ESP |
| Altramuz [vernacular name] | LUP 279 | ESP |
| Chocho [vernacular name] | LUP 280 | ESP |
| Altramuz | LUP 281 | ESP |
| Chocho [vernacular name] | LUP 282 | ESP |
| Altramuz | LUP 283 | ESP |
| Altramuz | LUP 284 | ESP |
| | LUP 285 | ESP |
| Altramuz [vernacular name] | LUP 286 | ESP |
| Altramuz [vernacular name] | LUP 287 | ESP |
| Altramuz, Chocho [vernacular name] | LUP 288 | ESP |
| Altramuz | LUP 289 | ESP |
| | LUP 290 | ESP |
| Altramuz [vernacular name] | LUP 291 | ESP |
| Altramuz | LUP 292 | ESP |
| Altramuz | LUP 293 | ESP |
| | LUP 294 | ESP |
| | LUP 295 | ESP |

| | | |
|---|---|---|
| | LUP 296 | ESP |
| | LUP 297 | ESP |
| | LUP 298 | ESP |
| | LUP 299 | ESP |
| | LUP 300 | ESP |
| | LUP 494 | ITA |
| | LUP 521 | ITA |
| | LUP 522 | ITA |
| Altramuz | LUP 2001 | ESP |
| Altramuz | LUP 2002 | ESP |
| Altramuz | LUP 2003 | ESP |
| Altramuz | LUP 2004 | ESP |
| Altramuz | LUP 2005 | ESP |
| Altramuz | LUP 2006 | ESP |
| Altramuz | LUP 2007 | ESP |
| Altramuz | LUP 2008 | ESP |
| Altramuz | LUP 2009 | ESP |
| Altramuz | LUP 2010 | ESP |
| Altramuz | LUP 2011 | ESP |
| Altramuz | LUP 2012 | ESP |
| Altramuz | LUP 2013 | ESP |
| Altramuz | LUP 2014 | ESP |
| Altramuz | LUP 2015 | ESP |
| Altramuz | LUP 2016 | ESP |
| Altramuz | LUP 2017 | ESP |
| Altramuz | LUP 2018 | ESP |
| | LUP 2019 | Unknown |
| SWEET WHITE | LUP 2020 | Unknown |
| | LUP 2022 | Unknown |
| | LUP 2023 | Unknown |
| | LUP 2024 | Unknown |
| | LUP 2025 | Unknown |
| | LUP 2026 | Unknown |
| | LUP 2027 | Unknown |
| | LUP 2028 | ITA |
| | LUP 2029 | Unknown |
| | LUP 2030 | MAR |
| | LUP 2031 | ITA |
| | LUP 2034 | ITA |
| | LUP 2035 | ESP |
| | LUP 2036 | ESP |
| | LUP 2037 | ESP |
| | LUP 2038 | ESP |
| | LUP 2039 | ESP |
| | LUP 2040 | ESP |
| | LUP 2041 | ESP |
| | LUP 2042 | ESP |
| SSK-79 | LUP 2043 | Unknown |
| LUPINI BEAN | LUP 2044 | Unknown |
| | LUP 2045 | SDN |
| DOCE | LUP 2046 | Unknown |
| | LUP 2047 | Unknown |

| | | |
|---|---|---|
| | LUP 2048 | ITA |
| | LUP 2049 | ITA |
| SATMAREAN | LUP 2051 | ROM |
| | LUP 2052 | ITA |
| | LUP 2053 | ITA |
| | LUP 2054 | ITA |
| | LUP 2055 | ITA |
| | LUP 2057 | ITA |
| | LUP 2059 | ITA |
| | LUP 2062 | ITA |
| | LUP 2063 | ITA |
| | LUP 2064 | ITA |
| | LUP 2065 | ITA |
| | LUP 2066 | ITA |
| | LUP 2067 | ITA |
| | LUP 2068 | ITA |
| | LUP 2069 | ITA |
| | LUP 2070 | ITA |
| | LUP 2071 | ITA |
| | LUP 2072 | ITA |
| | LUP 2073 | ESP |
| HETMAN | LUP 2074 | Unknown |
| | LUP 2075 | PRT |
| | LUP 2076 | ITA |
| | LUP 2077 | PAL |
| | LUP 2078 | ETH |
| | LUP 2079 | NLD |
| | LUP 2081 | GBR |
| START | LUP 2082 | RUS |
| | LUP 2083 | ITA |
| | LUP 2084 | ITA |
| | LUP 2085 | ITA |
| LUPPINO [VERNACULAR NAME] | LUP 2086 | ITA |
| | LUP 2087 | ITA |
| WOLODJA | LUP 2088 | SUN |
| ULTRA | LUP 2089 | Unknown |
| WAT | LUP 2090 | Unknown |
| | LUP 2091 | ESP |
| C 17/8 | LUP 2093 | DEU |
| | LUP 2094 | PRT |
| | LUP 2095 | ITA |
| | LUP 2096 | ITA |
| | LUP 2097 | ITA |
| | LUP 2098 | ITA |
| | LUP 2099 | ITA |
| | LUP 2100 | ITA |
| | LUP 2101 | ITA |
| MUTANTE "PLENUS" AUS NEUTRA | LUP 2104 | Unknown |
| HADM. 51901/63 (KOBIN. MUT. "PLENUS" X "ELATUS" A | LUP 2105 | Unknown |
| HADM. 38444/62 (MUT. "ELATUS" AUS NEUTRA X GÜLZ | LUP 2106 | Unknown |
| HADM. 16535/57 (HADM. 2635/50 X NEUTRA) | LUP 2107 | Unknown |
| HADM. 28791/59 (HADM. 2635/50 X NÄHRQUELL) | LUP 2108 | Unknown |

| | | |
|---|---|---|
| GÜLZOW ST. 91 | LUP 2112 | Unknown |
| KISVÁRDAI 1432/49 | LUP 5002 | HUN |
| | LUP 5004 | Unknown |
| MUENCHEBERGER STAMM 246 | LUP 5005 | DEU |
| | LUP 5006 | Unknown |
| PERAGIS STAMM 786/41 (SUESS) | LUP 5007 | DEU |
| GUELZOWER WEISSE SUESSLUPINE | LUP 5008 | DEU |
| GUELZOWER STAMM 883 | LUP 5009 | DEU |
| PERAGIS STAMM 786/41 | LUP 5011 | DEU |
| MATTIS BITTERE WEISSLUPINE | LUP 5012 | DEU |
| FIRLBECK 293 | LUP 5013 | DEU |
| | LUP 5015 | PRT |
| PFLUGS GELA | LUP 5016 | DEU |
| BLANCA | LUP 5017 | DEU |
| KISVARDAI FEHERVIRAGU | LUP 5018 | HUN |
| KRAFTQUELL | LUP 5020 | DEU |
| | LUP 5021 | YUG |
| MUENCHEBERGER STAMM 019 | LUP 5023 | DEU |
| GUELZOWER STAMM 884 | LUP 5024 | DEU |
| LANGE 11/57 | LUP 5025 | DEU |
| GYULATANYAI FEHERVIRAGU | LUP 5026 | HUN |
| PERAGIS STAMM 749/40 | LUP 5027 | DEU |
| | LUP 5028 | Unknown |
| KNEHDENER WEISSLUPINE | LUP 6410 | DEU |
| KALINA | LUP 6411 | Unknown |
| | LUP 6412 | ITA |
| KRUSE 186/59 | LUP 6413 | DEU |
| KIEVSKIJ SKOROSPELIJ | LUP 6414 | Unknown |
| KAIRO-1 | LUP 6415 | Unknown |
| EL HARRACH-2 | LUP 6417 | Unknown |
| GIBRIDSTART X KIEVSKIJSCEDRYJ | LUP 6419 | Unknown |
| BIATORUS-5 | LUP 6421 | Unknown |
| WUHOLIES KRETA | LUP 6423 | GRC |
| VOLODIA | LUP 6424 | SUN |
| | LUP 6425 | ITA |
| | LUP 6426 | ITA |
| | LUP 6427 | ITA |
| | LUP 6428 | DZA |
| | LUP 6429 | ITA |
| RENSCHER STAMM 60 | LUP 6430 | DEU |
| WEISSE TELLERLUPINE | LUP 6431 | Unknown |
| | LUP 6432 | ITA |
| LUCKY | LUP 6433 | FRA |
| | LUP 6434 | ITA |
| LUBLANC | LUP 6435 | FRA |
| | LUP 6436 | GRC |
| | LUP 6437 | ITA |
| D 9-7 | LUP 6438 | Unknown |
| EL HARRACH-1 | LUP 6440 | Unknown |
| KAIRO-4 | LUP 6443 | Unknown |
| | LUP 6444 | Unknown |
| MUTANT-28 | LUP 6445 | Unknown |

| | | |
|---|---|---|
| NEULAND | LUP 6446 | Unknown |
| | LUP 6447 | GRC |
| HETMAN | LUP 6449 | Unknown |
| LA 71 | LUP 6455 | DZA |
| | LUP 6457 | ITA |
| | LUP 6458 | ITA |
| | LUP 6459 | ITA |
| | LUP 6460 | ITA |
| | LUP 6461 | ITA |
| MULTO LUPA | LUP 6462 | DEU |
| | LUP 6463 | EGY |
| | LUP 6464 | ITA |
| | LUP 6465 | ITA |
| BLONDIN | LUP 6466 | DEU |
| | LUP 6467 | ITA |
| MUTANT-5 | LUP 6469 | Unknown |
| PETKUSER BITTERE WEISSLUPINE | LUP 6470 | DEU |
| | LUP 6471 | EGY |
| LA 106 | LUP 6476 | ITA |
| | LUP 6477 | GRC |
| | LUP 6481 | EGY |
| | LUP 6483 | EGY |
| | LUP 6485 | EGY |
| | LUP 6489 | GRC |
| 2884/49 | LUP 6490 | DEU |
| ZÁGREBESKÁ | LUP 6493 | YUG |
| | LUP 6494 | EGY |
| | LUP 6495 | ESP |
| IDA | LUP 6496 | DEU |
| | LUP 6497 | EGY |
| | LUP 6498 | EGY |
| | LUP 6499 | ITA |
| | LUP 6500 | ITA |
| LUTOP | LUP 6501 | FRA |
| | LUP 6502 | EGY |
| | LUP 6503 | USA |
| RENSCHER STAMM 23 | LUP 6507 | DEU |
| | LUP 6511 | EGY |
| | LUP 6513 | EGY |
| LUTOP | LUP 6515 | FRA |
| WEIBIT | LUP 6981 | DEU |
| | LUP 6985 | SUN |
| | LUP 6988 | ITA |
| BARDO | LUP 6993 | DEU |
| | LUP 7008 | ITA |
| NELLY | LUP 7014 | HUN |
| FORTUNA | LUP 7018 | DEU |
| | LUP 7021 | ITA |
| N91/50 | CGN10101 | ITA |
| N92/50 | CGN10102 | ITA |
| N96/51 | CGN10103 | ITA |
| N105/50 | CGN10104 | ITA |

| | | |
|---|---|---|
| N106/50 | CGN10105 | ITA |
| N107/50 | CGN10106 | ITA |
| N112/50 | CGN10107 | ITA |
| N121/50 | CGN10108 | ITA |
| N122/50 | CGN10109 | ITA |
| RENSCHER STAMM 60 | CGN10110 | DEU |
| PETKUSER BITTERE WEISS LUPINE | CGN10111 | DEU |
| PRZEBEDOWSKI WCZESNY | CGN10112 | POL |
| KISORDAI FEHEROIRAGU | CGN10113 | HUN |
| BILA LANGSBERSKA A | 05L0700002 | CZE |
| BILA LANGSBERSKA B | 05L0700003 | CZE |
| BILA LANGSBERSKA C | 05L0700004 | CZE |
| BILA ZAGREPSKA | 05L0700006 | YUG |
| NSL. LUPINA BILA | 05L0700014 | CZE |
| KIJEVSKIJ MUTANT | 05L0700028 | SUN |
| R-933 | 05L0700029 | POL |
| WAT | 05L0700030 | POL |
| KALINA | 05L0700031 | POL |
| KIJEVSKIJ MUTANT | 05L0700038 | CZE |
| WOLODJA | 05L0700040 | SUN |
| WTD | 05L0700041 | POL |
| POP I | 05L0700042 | POL |
| GORIZONT | 05L0700043 | SUN |
| SOLNECNYJ | 05L0700044 | SUN |
| START | 05L0700046 | SUN |
| MARIBANEZ | 05L0700047 | ESP |
| LOS PALACIOS | 05L0700048 | ESP |
| GUADAJINA | 05L0700049 | ESP |
| ELVAS | 05L0700050 | PRT |
| LABON | 05L0700051 | ESP |
| ALMENDRALEJO | 05L0700052 | ESP |
| MARCHENA | 05L0700053 | ESP |
| DRUZBA | 05L0700054 | SUN |
| ULTRA | 05L0700055 | AUS |
| BAC | 05L0700057 | POL |
| KALI | 05L0700058 | POL |
| HETMAN | 05L0700059 | POL |
| SATMAREAN | 05L0700060 | ROM |
| GOLF | 05L0700061 | DEU |
| LUBLANC | 05L0700062 | FRA |
| LUCKY | 05L0700063 | FRA |
| LUTOP | 05L0700064 | FRA |
| AMIGA | 05L0700069 | FRA |
| BARDO | 05L0700071 | POL |
| SINIJ PARUS | 05L0700073 | UKR |
| OLEZKA | 05L0700074 | UKR |
| DIETA | 05L0700077 | DNK |
| TERMIS | 250094 | EGY |
| Population-775 | 95602 | GRC |
| | 91-0037D | Unknown |
| | M20 | Unknown |
| P.25785 | 95611 | AUS |

| | | |
|---|---|---|
| POPULATION - 774 | 95673 | GRC |
| Population-1 | 95601 | GRC |
| EGIPT-7 | 95254 | EGY |
| EGIPT-8 | 95255 | EGY |
| POP.36415 | 95256 | MAR |
| POP.52892 | 95257 | ETH |
| MULTITALIA | 95258 | ITA |
| ALTRAMUZ SEWILLA | 95260 | ESP |
| TREMOCO FARO | 95262 | PRT |
| TREMOCO PORTALEGRO | 95263 | PRT |
| BG-5527 | 95265 | ITA |
| LOBBI BARCELONA | 95267 | ESP |
| EGYPTSKA | LR-0001 | EGY |
| TERRE | LR-0002 | ZAF |
| LA57-1 | LR-0003 | FRA |
| LUP2171 | LR-0004 | GRC |
| NAVAL_VILLAR_DE_IBOR-1 | LR-0005 | ESP |
| ALCANICES_ZAMORA | LR-0006 | ESP |
| GYULATANYA | LR-0118 | HUN |
| HANSA | LR-0119 | DEU |
| KALINA | LR-0123 | POL |
| NÄHRQUELL | LR-0129 | DEU |
| SHINFIELD | LR-0135 | GBR |
| | LR-0137 | PRT |
| ZÁGREBESKÁ | LR-0141 | YUG |
| TAPIOSZELE_II-85-1 | LR-0143 | HUN |
| LITHUANIAN662 | LR-0145 | SUN |
| JENA | LR-0150 | DEU |
| VIR 1930 | LR-0153 | SDN |
| PRIMORSKIJ | LR-0155 | UKR |
| VIR 490 | LR-0156 | ETH |
| NEUTRA | LR-0165 | DEU |
| LUCKY | LR-0168 | FRA |
| L-799-76 | LR-0175 | CHL |
| VIR1642 | LR-0176 | SDN |
| VIR 1643 | LR-0177 | SDN |
| POPULATION_AZORSKI | LR-0183 | PRT |
| SALAMANCA | LR-0200 | ESP |
| PORTO_VILLA_REAL | LR-0201 | PRT |
| BENELUP_DE_SIDONIA_ | LR-0202 | ESP |
| VILLA_DEL_RIO | LR-0203 | ESP |
| ESPIEL | LR-0204 | ESP |
| BELACAZAR | LR-0205 | ESP |
| ALCARACEJOS | LR-0206 | ESP |
| CAZELLA_DE_LA_SIERRA | LR-0207 | ESP |
| ALCUESCAR | LR-0208 | ESP |
| CACERES | LR-0209 | ESP |
| BURGUILLOS_DEL_CERRO | LR-0210 | ESP |
| SOTIEL_CORONADA | LR-0211 | ESP |
| RIO_PIEDRAS | LR-0212 | ESP |
| VENDAS_DE_RONCAO | LR-0213 | PRT |
| CERCAL | LR-0214 | PRT |

| | | |
|---|---|---|
| BENSAFRIM | LR-0215 | PRT |
| ODEMIRA | LR-0216 | PRT |
| PORTO_DO_LAGOS | LR-0217 | PRT |
| RIO_DE_TALARES | LR-0218 | PRT |
| CORTE_CARCIA | LR-0219 | PRT |
| AMEIXAL | LR-0220 | PRT |
| ALJUSTREL | LR-0221 | PRT |
| MIMOSA | LR-0222 | PRT |
| MONTIJO | LR-0223 | PRT |
| CORUCHE | LR-0224 | PRT |
| ALMEIRIM | LR-0225 | PRT |
| MONTOMORO-NOVO | LR-0226 | PRT |
| EVORA | LR-0227 | PRT |
| PORTEL | LR-0228 | PRT |
| REGUENCOS_DE_MONSARAZ | LR-0229 | PRT |
| VILA_VIGOSA | LR-0230 | PRT |
| REDONDO | LR-0231 | PRT |
| SINTRA | LR-0232 | PRT |
| PALHOCA | LR-0233 | PRT |
| ESPARRAGALEJO | LR-0234 | ESP |
| NAVA_DE_SANTAIAGO | LR-0235 | ESP |
| ZARZA | LR-0236 | ESP |
| VILLAR_DEL_REY | LR-0237 | ESP |
| ALBUQUERQUE | LR-0238 | ESP |
| BADAJOZ | LR-0239 | ESP |
| VILLA_MESIIAS | LR-0240 | ESP |
| NAVAL_VILLAR_DE_IBOR-2 | LR-0241 | ESP |
| BAHONA_DE_IBOR | LR-0242 | ESP |
| LA_FRAGENDA | LR-0243 | ESP |
| SALDANA | LR-0244 | ESP |
| SANT_CELONI | LR-0245 | ESP |
| MEALHADA | LR-0246 | PRT |
| KESSARIAN | LR-0247 | GRC |
| KISSOS | LR-0248 | GRC |
| IRAKLION | LR-0250 | GRC |
| NOMOS_RETHIMNO | LR-0256 | GRC |
| FLORIA | LR-0260 | GRC |
| GAVRION | LR-0261 | GRC |
| BEODEMIADOS | LR-0264 | GRC |
| LAKKOPETRA | LR-0265 | GRC |
| ARAXOS | LR-0267 | GRC |
| SIMOPOULON | LR-0268 | GRC |
| | LR-0269 | GRC |
| LIVADAKI | LR-0270 | GRC |
| KALITHEA | LR-0271 | GRC |
| ALLAGI | LR-0273 | GRC |
| VASILIKON | LR-0274 | GRC |
| CHORA | LR-0276 | GRC |
| THOURIA | LR-0277 | GRC |
| GEORGITSI | LR-0279 | GRC |
| KARAVAS | LR-0282 | GRC |
| ARTEMISIA | LR-0285 | GRC |

| | | |
|---|---|---|
| AGIOS_NIKON | LR-0287 | GRC |
| GITHION | LR-0289 | GRC |
| MULINE-1 | LR-0290 | HRV |
| LUKORAN | LR-0291 | HRV |
| MONTE_FIASCONE | LR-0327 | ITA |
| VITERBO-1 | LR-0328 | ITA |
| VITERBO-2 | LR-0329 | ITA |
| PICO_FROSINONE | LR-0331 | ITA |
| GROTTAMINORDA | LR-0332 | ITA |
| POUTIGNANO | LR-0333 | ITA |
| ALBEROBELLO | LR-0334 | ITA |
| LOCOROTONDO | LR-0335 | ITA |
| MARTINA-1 | LR-0336 | ITA |
| MARTINA-2 | LR-0337 | ITA |
| SAN_PANCRAZIO | LR-0338 | ITA |
| MESAGNE | LR-0339 | ITA |
| SAN_VITO | LR-0340 | ITA |
| ETH013_6286 | LR-0402 | ETH |
| ETH013_13661 | LR-0403 | ETH |
| ETH013_13662 | LR-0404 | ETH |
| ETH013_13664 | LR-0405 | ETH |
| ETH013_13666 | LR-0406 | ETH |
| ETH013_13667 | LR-0407 | ETH |
| VIGGIANO | LR-0443 | ITA |
| LUP2057 | LR-0444 | ITA |
| LUP2064 | LR-0447 | ITA |
| START | LR-0451 | RUS |
| LUP6447 | LR-0463 | GRC |
| N3507 | LR-0467 | PAL |
| GM141 | LR-0469 | MAR |
| MJS355-1 | LR-0471 | GRC |
| MJS355-2 | LR-0472 | GRC |
| GRC5659B | LR-0476 | GRC |
| DAMASCUS | LR-0477 | SYR |
| KEN5274B | LR-0478 | KEN |
| CMO103 | LR-0479 | MAR |
| KEN5143B | LR-0482 | KEN |
| ILCA13665 | LR-0486 | ETH |
| EGY6484B | LR-0503 | EGY |
| SYR6258B | LR-0511 | SYR |
| EGY6517B | LR-0516 | EGY |
| GRC5262B | LR-0517 | GRC |
| ALB.01 | LR-0526 | DZA |
| EGY6435B | LR-0533 | EGY |
| LA37-ROUND_ | LR-0536 | FRA |
| MURRINGO | LR-0539 | AUS |
| MEGALOPOLIS | LR-0549 | GRC |
| ARKANSAS_34 | LR-0552 | USA |
| VILLAX | LR-0555 | MAR |
| BALADI | LR-0557 | JOR |
| KONYA | LR-0558 | TUR |
| ALGERIA_P14-A4 | LR-0560 | DZA |

| | | |
|---|---|---|
| LUCKY | LR-0562 | FRA |
| AMIGA | LR-0567 | CHL |
| LD037_JB | LR-0574 | HUN |
| TERMS - CAIRO SUPERMARKET | AGG20010LUPN1 | Unknown |
| P20903 | AGG20156LUPN1 | PRT |
| P20909 | AGG20159LUPN1 | PRT |
| P20910 | AGG20160LUPN1 | PRT |
| P20918 | AGG20166LUPN1 | DEU |
| P20931 | AGG20176LUPN1 | DEU |
| P23037 | AGG20533LUPN1 | GRC |
| MJS016 | AGG20728LUPN1 | ESP |
| MJS025 | AGG20730LUPN1 | ESP |
| MJS029 | AGG20731LUPN1 | ESP |
| MJS034 | AGG20732LUPN1 | ESP |
| MJS044 | AGG20733LUPN1 | ESP |
| MJS053 | AGG20734LUPN1 | ESP |
| MJS091 | AGG20736LUPN1 | PRT |
| MJS100 | AGG20737LUPN1 | PRT |
| MJS112 | AGG20738LUPN1 | PRT |
| MJS138 | AGG20740LUPN1 | PRT |
| MJS144 | AGG20741LUPN1 | PRT |
| MJS178 | AGG20744LUPN1 | ESP |
| MJS180 | AGG20745LUPN1 | ESP |
| MJS208 | AGG20746LUPN1 | ESP |
| MJS305 | AGG20749LUPN1 | GRC |
| MJS343 | AGG20752LUPN1 | GRC |
| MJS344 | AGG20753LUPN1 | GRC |
| MJS352 | AGG20754LUPN1 | GRC |
| MJS362 | AGG20756LUPN1 | GRC |
| MJS374 | AGG20758LUPN1 | GRC |
| MJS377 | AGG20759LUPN1 | GRC |
| MJS380 | AGG20760LUPN1 | GRC |
| MJS383 | AGG20761LUPN1 | GRC |
| MJS053 | AGG20864LUPN1 | ESP |
| MJS053 | AGG20865LUPN1 | ESP |
| MJS138 | AGG20866LUPN1 | PRT |
| MJS138 | AGG20867LUPN1 | PRT |
| MJS178 | AGG20869LUPN1 | ESP |
| MJS303 | AGG20872LUPN1 | GRC |
| MJS303 | AGG20873LUPN1 | GRC |
| MJS304 | AGG20875LUPN1 | GRC |
| MJS305 | AGG20876LUPN1 | GRC |
| MJS305 | AGG20877LUPN1 | GRC |
| MJS343 | AGG20880LUPN1 | GRC |
| MJS343 | AGG20881LUPN1 | GRC |
| MJS366 | AGG20882LUPN1 | GRC |
| MJS366 | AGG20883LUPN1 | GRC |
| MJS366 | AGG20884LUPN1 | GRC |
| MJS377 | AGG20885LUPN1 | GRC |
| MJS377 | AGG20886LUPN1 | GRC |
| MJS380 | AGG20887LUPN1 | GRC |
| MJS380 | AGG20888LUPN1 | GRC |

| | | |
|---|---|---|
| MJS323 | AGG20893LUPN1 | GRC |
| MJS355 | AGG20896LUPN1 | GRC |
| MJS362 | AGG20897LUPN1 | GRC |
| MJS362 | AGG20898LUPN1 | GRC |
| LUBLANC | AGG20992LUPN1 | FRA |
| LUCKY | AGG20993LUPN1 | FRA |
| LUTOP | AGG20994LUPN1 | FRA |
| NYIRSEGI EDES | AGG20995LUPN1 | HUN |
| NYIRSEGI KESERU | AGG20996LUPN1 | HUN |
| START | AGG20997LUPN1 | SUN |
| WB2 | AGG21000LUPN1 | Unknown |
| INRA6089 | AGG21001LUPN1 | MAR |
| INRA6090 | AGG21002LUPN1 | MAR |
| INRA6210 | AGG21003LUPN1 | MAR |
| K2 | AGG21005LUPN1 | USA |
| K3 | AGG21006LUPN1 | USA |
| K4 | AGG21007LUPN1 | USA |
| PISCEVOJ P25947? | AGG21009LUPN1 | UKR |
| 16564 CPI16564? | AGG21295LUPN1 | PRT |
| 16569 CPI16569? | AGG21296LUPN1 | PRT |
| LANGSBERGSKA (ST B) | AGG21297LUPN1 | CZE |
| 23300 | AGG21298LUPN1 | HUN |
| 31620 | AGG21299LUPN1 | GRC |
| 31621 | AGG21300LUPN1 | BGR |
| 53828 | AGG21301LUPN1 | NLD |
| MJS462 | AGG21303LUPN1 | ESP |
| MJS468 | AGG21304LUPN1 | ESP |
| MJS074 | AGG21305LUPN1 | ESP |
| MJS074 | AGG21306LUPN1 | ESP |
| MJS074 | AGG21307LUPN1 | ESP |
| MJS094 | AGG21308LUPN1 | PRT |
| MJS094 | AGG21309LUPN1 | PRT |
| MJS094 | AGG21310LUPN1 | PRT |
| MJS119 | AGG21311LUPN1 | PRT |
| MJS554 | AGG21312LUPN1 | ESP |
| MJS148 | AGG21313LUPN1 | PRT |
| MJS162 | AGG21314LUPN1 | PRT |
| MJS162 | AGG21315LUPN1 | PRT |
| MJS162 | AGG21316LUPN1 | PRT |
| MJS189 | AGG21317LUPN1 | ESP |
| MJS189 | AGG21318LUPN1 | ESP |
| MJS189 | AGG21319LUPN1 | ESP |
| MJS234 | AGG21320LUPN1 | ESP |
| MJS253 | AGG21321LUPN1 | PRT |
| MJS253 | AGG21322LUPN1 | PRT |
| MJS253 | AGG21323LUPN1 | PRT |
| MJS261 | AGG21324LUPN1 | PRT |
| MJS276 | AGG21325LUPN1 | PRT |
| MJS282 | AGG21326LUPN1 | PRT |
| MJS337 | AGG21327LUPN1 | GRC |
| MJS337 | AGG21328LUPN1 | GRC |
| MJS337 | AGG21329LUPN1 | GRC |

| | | |
|---|---|---|
| MJS337 | AGG21330LUPN1 | GRC |
| MJS337 | AGG21331LUPN1 | GRC |
| MJS374 | AGG21332LUPN1 | GRC |
| MJS374 | AGG21333LUPN1 | GRC |
| GRC5032B | AGG21334LUPN1 | GRC |
| GRC5032B | AGG21335LUPN1 | GRC |
| GRC5032B | AGG21336LUPN1 | GRC |
| GRC5032B | AGG21337LUPN1 | GRC |
| GRC5038B | AGG21338LUPN1 | GRC |
| GRC5659B | AGG21339LUPN1 | GRC |
| GRC5659B | AGG21340LUPN1 | GRC |
| GRC5659B | AGG21341LUPN1 | GRC |
| GRC5661B | AGG21342LUPN1 | GRC |
| GRC5663B | AGG21343LUPN1 | GRC |
| GRC5668B | AGG21344LUPN1 | GRC |
| GRC5669B | AGG21345LUPN1 | GRC |
| GRC5670B | AGG21346LUPN1 | GRC |
| CMO102 | AGG21348LUPN1 | MAR |
| CMO102 | AGG21349LUPN1 | MAR |
| DAMASCUS | AGG21350LUPN1 | SYR |
| GMJ14B | AGG21351LUPN1 | GRC |
| KEN5274B | AGG21352LUPN1 | KEN |
| KEN5141B | AGG21353LUPN1 | KEN |
| CFP70 | AGG21495LUPN1 | PRT |
| GIZA 2 | AGG21496LUPN1 | EGY |
| HAMBURG | AGG21497LUPN1 | DEU |
| K1904 | AGG21498LUPN1 | SUN |
| FRA6635B | AGG21925LUPN1 | PRT |
| FRA6645B | AGG21926LUPN1 | PRT |
| FRA6643B | AGG21927LUPN1 | PRT |
| FRA6650B | AGG21928LUPN1 | PRT |
| FRA6649B | AGG21929LUPN1 | PRT |
| L/23 | AGG21930LUPN1 | CHL |
| NYIRSEGI | AGG21933LUPN1 | HUN |
| 100E60 | AGG21935LUPN1 | GBR |
| 114E63 | AGG21936LUPN1 | GBR |
| 78E55 | AGG21938LUPN1 | GBR |
| HETMAN | AGG21940LUPN1 | POL |
| BAC | AGG21941LUPN1 | POL |
| POL6284B | AGG21977LUPN1 | POL |
| POL6282B | AGG21978LUPN1 | POL |
| POL6272B | AGG21979LUPN1 | POL |
| USA6307B | AGG21981LUPN1 | USA |
| USA6308B | AGG21982LUPN1 | USA |
| USA6311B | AGG21983LUPN1 | USA |
| USA6309B | AGG21984LUPN1 | USA |
| USA6310B | AGG21985LUPN1 | USA |
| FRA6559B | AGG21986LUPN1 | PRT |
| FRA6582B | AGG21988LUPN1 | PRT |
| EGY6423B | AGG21989LUPN1 | EGY |
| EGY6420B | AGG21990LUPN1 | EGY |
| EGY6428B | AGG21991LUPN1 | EGY |

| | | |
|---|---|---|
| EGY6427B | AGG21992LUPN1 | EGY |
| EGY6424B | AGG21993LUPN1 | EGY |
| EGY6430B | AGG21994LUPN1 | EGY |
| EGY6431B | AGG21995LUPN1 | EGY |
| EGY6429B | AGG21996LUPN1 | EGY |
| EGY6437B | AGG21998LUPN1 | EGY |
| EGY6434B | AGG21999LUPN1 | EGY |
| EGY6440B | AGG22000LUPN1 | EGY |
| EGY6442B | AGG22001LUPN1 | EGY |
| EGY6447B | AGG22002LUPN1 | EGY |
| EGY6444B | AGG22003LUPN1 | EGY |
| EGY6453B | AGG22004LUPN1 | EGY |
| EGY6451B | AGG22005LUPN1 | EGY |
| EGY6458B | AGG22006LUPN1 | EGY |
| EGY6457B | AGG22007LUPN1 | EGY |
| EGY6456B | AGG22008LUPN1 | EGY |
| EGY6455B | AGG22009LUPN1 | EGY |
| EGY6454B | AGG22010LUPN1 | EGY |
| EGY6463B | AGG22011LUPN1 | EGY |
| EGY6462B | AGG22012LUPN1 | EGY |
| EGY6460B | AGG22013LUPN1 | EGY |
| EGY6467B | AGG22014LUPN1 | EGY |
| EGY6468B | AGG22015LUPN1 | EGY |
| EGY6471B | AGG22016LUPN1 | EGY |
| EGY6470B | AGG22017LUPN1 | EGY |
| EGY6478B | AGG22018LUPN1 | EGY |
| EGY6477B | AGG22019LUPN1 | EGY |
| EGY6483B | AGG22020LUPN1 | EGY |
| EGY6482B | AGG22021LUPN1 | EGY |
| EGY6480B | AGG22022LUPN1 | EGY |
| EGY6479B | AGG22023LUPN1 | EGY |
| EGY6487B | AGG22026LUPN1 | EGY |
| EGY6486B | AGG22027LUPN1 | EGY |
| EGY6492B | AGG22028LUPN1 | EGY |
| EGY6490B | AGG22029LUPN1 | EGY |
| EGY6501B | AGG22030LUPN1 | EGY |
| EGY6502B | AGG22031LUPN1 | EGY |
| EGY6507B | AGG22032LUPN1 | EGY |
| EGY6508B | AGG22033LUPN1 | EGY |
| EGY6506B | AGG22034LUPN1 | EGY |
| EGY6505B | AGG22035LUPN1 | EGY |
| EGY6504B | AGG22036LUPN1 | EGY |
| EGY6511B | AGG22037LUPN1 | EGY |
| EGY6512B | AGG22038LUPN1 | EGY |
| EGY6513B | AGG22039LUPN1 | EGY |
| EGY6509B | AGG22040LUPN1 | EGY |
| EGY6515B | AGG22041LUPN1 | EGY |
| EGY6514B | AGG22042LUPN1 | EGY |
| EGY6525B | AGG22043LUPN1 | EGY |
| EGY6529B | AGG22044LUPN1 | EGY |
| AMIGA | AGG22047LUPN1 | DEU |
| IDA | AGG22048LUPN1 | DEU |

| | | |
|---|---|---|
| DDR6336B | AGG22049LUPN1 | DEU |
| EGY6413B | AGG22050LUPN1 | EGY |
| EGY6414B | AGG22051LUPN1 | EGY |
| EGY6415B | AGG22052LUPN1 | EGY |
| EGY6412B | AGG22053LUPN1 | EGY |
| EGY6418B | AGG22054LUPN1 | EGY |
| EGY6417B | AGG22055LUPN1 | EGY |
| ESP6064B.1 | AGG22057LUPN1 | ESP |
| ESTA.1 | AGG22065LUPN1 | ZAF |
| EGY6460B | AGG22117LUPN1 | EGY |
| 131E9-2 | AGG22118LUPN1 | Unknown |
| FRA6628B | AGG22122LUPN1 | FRA |
| FRA6639B | AGG22123LUPN1 | FRA |
| FRA6640B | AGG22124LUPN1 | FRA |
| FRA6641B | AGG22125LUPN1 | FRA |
| FRA6642B | AGG22126LUPN1 | FRA |
| FRA6644B | AGG22127LUPN1 | FRA |
| FRA6655B | AGG22129LUPN1 | FRA |
| FRA6660B | AGG22131LUPN1 | FRA |
| FRA6663B | AGG22132LUPN1 | FRA |
| FRA6665B | AGG22133LUPN1 | FRA |
| FRA6666B | AGG22134LUPN1 | FRA |
| FRA6670B | AGG22135LUPN1 | FRA |
| FRA6673B | AGG22136LUPN1 | FRA |
| FRA6713B | AGG22143LUPN1 | FRA |
| FRA6715B | AGG22144LUPN1 | FRA |
| FRA6719B | AGG22145LUPN1 | FRA |
| FRA6720B | AGG22146LUPN1 | FRA |
| FRA6724B | AGG22147LUPN1 | FRA |
| EGY6419B | AGG22148LUPN1 | EGY |
| EGY6421B | AGG22149LUPN1 | EGY |
| EGY6443B | AGG22150LUPN1 | EGY |
| EGY6445B | AGG22151LUPN1 | EGY |
| EGY6448B | AGG22152LUPN1 | EGY |
| EGY6450B | AGG22153LUPN1 | EGY |
| EGY6475B | AGG22155LUPN1 | EGY |
| EGY6489B | AGG22156LUPN1 | EGY |
| EGY6491B | AGG22157LUPN1 | EGY |
| EGY6493B | AGG22158LUPN1 | EGY |
| EGY6495B | AGG22159LUPN1 | EGY |
| EGY6500B | AGG22160LUPN1 | EGY |
| EGY6518B | AGG22162LUPN1 | EGY |
| EGY6520B | AGG22163LUPN1 | EGY |
| EGY6521B | AGG22164LUPN1 | EGY |
| EGY6522B | AGG22165LUPN1 | EGY |
| EGY6526B | AGG22166LUPN1 | EGY |
| SUN6332B | AGG22167LUPN1 | SUN |
| SYR6728B | AGG22168LUPN1 | SYR |
| DDR6793B | AGG22169LUPN1 | DEU |
| DDR6794B | AGG22170LUPN1 | DEU |
| DDR6795B | AGG22171LUPN1 | DEU |
| DDR6796B | AGG22172LUPN1 | DEU |

| | | |
|---|---|---|
| MINORI | AGG22173LUPN1 | Unknown |
| 131E9-2.1 | AGG22174LUPN1 | Unknown |
| MJS356.1 | AGG22175LUPN1 | Unknown |
| MJS356.1.1 | AGG22176LUPN1 | Unknown |
| MJS356.1.1.1 | AGG22177LUPN1 | Unknown |
| MJS356.1.1.2 | AGG22178LUPN1 | Unknown |
| FRA6627B.1 | AGG22180LUPN1 | FRA |
| FRA6628B.1 | AGG22181LUPN1 | FRA |
| FRA6715B.1 | AGG22182LUPN1 | FRA |
| FRA6719B.1 | AGG22183LUPN1 | FRA |
| FRA6720B.1 | AGG22184LUPN1 | FRA |
| GRC5262B | AGG22339LUPN1 | GRC |
| FRA6646B | AGG22350LUPN1 | PRT |
| FRA6647B | AGG22351LUPN1 | PRT |
| FRA6648B | AGG22352LUPN1 | PRT |
| FRA6667B | AGG22353LUPN1 | PRT |
| FRA6668B | AGG22354LUPN1 | PRT |
| FRA6723B | AGG22360LUPN1 | FRA |
| EGY6439B | AGG22361LUPN1 | EGY |
| EGY6498B | AGG22364LUPN1 | EGY |
| EGY6527B | AGG22365LUPN1 | EGY |
| GRC6331B | AGG22366LUPN1 | GRC |
| DDR6791B | AGG22367LUPN1 | DEU |
| DDR6792B | AGG22368LUPN1 | DEU |
| EGY6448B.1 | AGG22380LUPN1 | EGY |
| FRA6656B.1 | AGG22390LUPN1 | PRT |
| FRA6669B.1 | AGG22391LUPN1 | PRT |
| LUCYANE | AGG22401LUPN1 | FRA |
| LUDET | AGG22402LUPN1 | FRA |
| SHINFIELD | AGG22409LUPN2 | Unknown |
| FRA6611B | AGG22411LUPN2 | Unknown |
| CW77 | AGG22506LUPN2 | AUS |
| FRA6561B | AGG22518LUPN2 | PRT |
| FRA6562B | AGG22519LUPN2 | PRT |
| FRA6563B | AGG22520LUPN2 | PRT |
| FRA6565B | AGG22521LUPN2 | PRT |
| FRA6566B | AGG22522LUPN2 | PRT |
| FRA6567B | AGG22523LUPN2 | PRT |
| FRA6570B | AGG22524LUPN2 | PRT |
| FRA6571B | AGG22525LUPN2 | PRT |
| FRA6572B | AGG22526LUPN2 | PRT |
| FRA6573B | AGG22527LUPN2 | PRT |
| FRA6575B | AGG22529LUPN2 | PRT |
| FRA6576B | AGG22530LUPN2 | PRT |
| FRA6577B | AGG22531LUPN2 | PRT |
| FRA6583B | AGG22532LUPN2 | PRT |
| FRA6586B | AGG22533LUPN2 | PRT |
| FRA6587B | AGG22534LUPN2 | PRT |
| FRA6588B | AGG22535LUPN2 | PRT |
| FRA6589B | AGG22536LUPN2 | PRT |
| FRA6590B | AGG22537LUPN2 | PRT |
| FRA6591B | AGG22538LUPN2 | PRT |

| | | |
|---|---|---|
| FRA6592B | AGG22539LUPN2 | PRT |
| FRA6593B | AGG22540LUPN2 | PRT |
| FRA6598B | AGG22543LUPN2 | PRT |
| FRA6599B | AGG22544LUPN2 | PRT |
| FRA6600B | AGG22545LUPN2 | PRT |
| FRA6601B | AGG22546LUPN2 | PRT |
| FRA6602B | AGG22547LUPN2 | PRT |
| FRA6603B | AGG22548LUPN2 | PRT |
| FRA6604B | AGG22549LUPN2 | PRT |
| FRA6605B | AGG22550LUPN2 | PRT |
| FRA6606B | AGG22551LUPN2 | PRT |
| FRA6613B | AGG22552LUPN2 | PRT |
| FRA6614B | AGG22553LUPN2 | PRT |
| FRA6615B | AGG22554LUPN2 | PRT |
| FRA6616B | AGG22555LUPN2 | PRT |
| FRA6617B | AGG22556LUPN2 | PRT |
| FRA6618B | AGG22557LUPN2 | PRT |
| FRA6619B | AGG22558LUPN2 | PRT |
| FRA6620B | AGG22559LUPN2 | PRT |
| FRA6621B | AGG22560LUPN2 | PRT |
| FRA6622B | AGG22561LUPN2 | PRT |
| FRA6623B | AGG22562LUPN2 | PRT |
| FRA6624B | AGG22563LUPN2 | PRT |
| FRA6625B | AGG22564LUPN2 | PRT |
| FRA6637B | AGG22565LUPN2 | PRT |
| FRA6638B | AGG22566LUPN2 | PRT |
| FRA6651B | AGG22567LUPN2 | PRT |
| FRA6671B | AGG22568LUPN2 | PRT |
| FRA6672B | AGG22569LUPN2 | PRT |
| FRA6674B | AGG22570LUPN2 | PRT |
| FRA6675B | AGG22571LUPN2 | PRT |
| FRA6676B | AGG22572LUPN2 | PRT |
| FRA6677B | AGG22573LUPN2 | PRT |
| FRA6678B | AGG22574LUPN2 | PRT |
| FRA6679B | AGG22575LUPN2 | PRT |
| FRA6680B | AGG22576LUPN2 | PRT |
| FRA6681B | AGG22577LUPN2 | PRT |
| FRA6727B | AGG22579LUPN2 | PRT |
| FRA6641B | AGG22678LUPN2 | PRT |
| FRA6667B | AGG22680LUPN2 | PRT |
| LA173 | AGG22683LUPN2 | ETH |
| LA294 | AGG22684LUPN2 | EGY |
| LA295 | AGG22685LUPN2 | EGY |
| LA338 | AGG22686LUPN2 | SDN |
| LA339 | AGG22687LUPN2 | SDN |
| LA340 | AGG22688LUPN2 | SDN |
| LA341 | AGG22689LUPN2 | SDN |
| LA343 | AGG22690LUPN2 | EGY |
| LA344 | AGG22691LUPN2 | EGY |
| LA345 | AGG22692LUPN2 | EGY |
| LA347 | AGG22693LUPN2 | EGY |
| LA348 | AGG22694LUPN2 | EGY |

| | | |
|---|---|---|
| LA349 | AGG22695LUPN2 | EGY |
| LA350 | AGG22696LUPN2 | EGY |
| LA351 | AGG22697LUPN2 | EGY |
| LA352 | AGG22698LUPN2 | EGY |
| LA353 | AGG22699LUPN2 | EGY |
| LA354 | AGG22700LUPN2 | EGY |
| LA355 | AGG22701LUPN2 | FRA |
| LA356 | AGG22702LUPN2 | FRA |
| LA357 | AGG22703LUPN2 | FRA |
| LA358 | AGG22704LUPN2 | FRA |
| LA359 | AGG22705LUPN2 | FRA |
| LA360 | AGG22706LUPN2 | FRA |
| LA376 | AGG22707LUPN2 | EGY |
| LA377 | AGG22708LUPN2 | EGY |
| LA378 | AGG22709LUPN2 | EGY |
| LA379 | AGG22710LUPN2 | EGY |
| LA380 | AGG22711LUPN2 | EGY |
| LA381 | AGG22712LUPN2 | EGY |
| LA382 | AGG22713LUPN2 | EGY |
| LA383 | AGG22714LUPN2 | EGY |
| LA384 | AGG22715LUPN2 | EGY |
| LA385 | AGG22716LUPN2 | EGY |
| LA386 | AGG22717LUPN2 | EGY |
| LA387 | AGG22718LUPN2 | EGY |
| LA388 | AGG22719LUPN2 | EGY |
| LA389 | AGG22720LUPN2 | EGY |
| LA391 | AGG22721LUPN2 | EGY |
| LA393 | AGG22722LUPN2 | EGY |
| LA394 | AGG22723LUPN2 | EGY |
| LA395 | AGG22724LUPN2 | EGY |
| LA396 | AGG22725LUPN2 | EGY |
| LA397 | AGG22726LUPN2 | EGY |
| LA398 | AGG22727LUPN2 | EGY |
| LA401 | AGG22729LUPN2 | ETH |
| C407 | AGG22730LUPN2 | FRA |
| LA420 | AGG22731LUPN2 | SDN |
| LA421 | AGG22732LUPN2 | SDN |
| LA422 | AGG22733LUPN2 | SDN |
| LA423 | AGG22734LUPN2 | SDN |
| LA424 | AGG22735LUPN2 | SDN |
| LA433 | AGG22736LUPN2 | EGY |
| LA441 | AGG22737LUPN2 | EGY |
| LA558 | AGG22738LUPN2 | EGY |
| LA559 | AGG22739LUPN2 | EGY |
| LA629 | AGG22740LUPN2 | ETH |
| LA674 | AGG22741LUPN2 | FRA |
| 71/328 | AGG22742LUPN2 | GRC |
| 34/154 | AGG22743LUPN2 | GRC |
| 38/180 | AGG22744LUPN2 | GRC |
| 66/289 | AGG22745LUPN2 | GRC |
| FRA6634B | AGG22891LUPN1 | PRT |
| ETH188 | AGG23146LUPN1 | ETH |

| | | |
|---|---|---|
| ETH208 | AGG23151LUPN1 | ETH |
| ETH225 | AGG23164LUPN1 | ETH |
| ETH226 | AGG23165LUPN1 | ETH |
| ETH228 | AGG23167LUPN1 | ETH |
| KALI | AGG23187LUPN1 | POL |
| P26777 | AGG23312LUPN1 | Unknown |
| P27448 | AGG23313LUPN1 | Unknown |
| P 28655 | AGG23314LUPN1 | Unknown |
| P 28699 | AGG23315LUPN1 | Unknown |
| P 28700 | AGG23316LUPN1 | Unknown |
| P 28701 | AGG23317LUPN1 | Unknown |
| P 28702 | AGG23318LUPN1 | Unknown |
| P 28703 | AGG23319LUPN1 | Unknown |
| P 28704 | AGG23320LUPN1 | Unknown |
| P 28705 | AGG23321LUPN1 | Unknown |
| P 28752 | AGG23322LUPN1 | Unknown |
| P 28755 | AGG23325LUPN1 | Unknown |
| P 28757 | AGG23326LUPN1 | Unknown |
| P 28762 | AGG23330LUPN1 | Unknown |
| P 28764 | AGG23332LUPN1 | Unknown |
| P 28765 | AGG23333LUPN1 | Unknown |
| P 28766 | AGG23334LUPN1 | Unknown |
| P 28767 | AGG23335LUPN1 | Unknown |
| P28985 | AGG23479LUPN1 | Unknown |
| P28986 | AGG23480LUPN1 | Unknown |
| P28988 | AGG23481LUPN1 | Unknown |
| P28989 | AGG23482LUPN1 | Unknown |
| P28990 | AGG23483LUPN1 | Unknown |
| P28991 | AGG23484LUPN1 | Unknown |
| P28992 | AGG23485LUPN1 | Unknown |
| P28993 | AGG23486LUPN1 | Unknown |
| P28996 | AGG23489LUPN1 | Unknown |
| P28997 | AGG23490LUPN1 | Unknown |
| P29000 | AGG23493LUPN1 | Unknown |
| P29002 | AGG23495LUPN1 | Unknown |
| P29003 | AGG23496LUPN1 | Unknown |
| P29004 | AGG23497LUPN1 | Unknown |
| P29005 | AGG23498LUPN1 | Unknown |
| P29006 | AGG23499LUPN1 | Unknown |
| P29013 | AGG23503LUPN1 | Unknown |
| P29021 | AGG23504LUPN1 | Unknown |
| P29028 | AGG23509LUPN1 | Unknown |
| P29030 | AGG23511LUPN1 | Unknown |
| MURRINGO | AGG23531LUPN1 | AUS |
| | SVGB-12129 | POL |
| | SVGB-12130 | ROM |
| | SVGB-12131 | FRA |
| | SVGB-12135 | DEU |
| | SVGB-12143 | HUN |
| | SVGB-12150 | UKR |
| | SVGB-12151 | DEU |
| | SVGB-12153 | DEU |

| | | |
|---|---|---|
| | SVGB-12154 | DEU |
| | SVGB-12156 | POL |
| | SVGB-12158 | ROM |
| | SVGB-12159 | ROM |
| | SVGB-12160 | ROM |
| | SVGB-12161 | FRA |
| | SVGB-12163 | ROM |
| | SVGB-12165 | DEU |
| Blanca | SVGB-12167 | DEU |
| | SVGB-12174 | DEU |
| | SVGB-12175 | DEU |
| | SVGB-12560 | ROM |
| | SVGB-13837 | ROM |
| | SVGB-13839 | FRA |
| | SVGB-19914 | ROM |
| | TENNIS | Unknown |
| | MULTITALIA | ITA |
| | IS-LUP19-1 | Unknown |
| | SCBC 2851 | ROM |
| | SCBC 2853 | ROM |
| | SCBC 2854 | ROM |
| | SCBC 2855 | ROM |
| | SCBC 2856 | ROM |
| | SCBC 2857 | ROM |
| | SCBC 2859 | ROM |
| | SCBC 2860 | ROM |
| | NC001518 | ESP |
| | NC070931 | ESP |
| | NC001520 | ESP |
| | NC070933 | ESP |
| | NC001522 | ESP |
| | NC001523 | ESP |
| | NC001524 | ESP |
| | NC001525 | ESP |
| | NC001526 | ESP |
| | NC001527 | ESP |
| | NC070940 | ESP |
| | NC070941 | ESP |
| | NC001530 | ESP |
| | NC070943 | ESP |
| | NC001532 | ESP |
| | NC001533 | ESP |
| | NC070946 | ESP |
| PFLUGS GELA | NC001539 | HUN |
| NÄHRQUELL | NC001540 | HUN |
| KRAFTQUELL | NC001541 | HUN |
| ASCAR | NC001542 | FRA |
| ITALIEN | NC001543 | DEU |
| PELOJANNES | NC001544 | DEU |
| WEISE BITTERLUPINE | NC001545 | DEU |
| KALIRIA | NC001546 | POL |
| KALI | NC001547 | POL |

| | | |
|---|---|---|
| KIEVSKIJ | NC000648 | SUN |
| PALESTINE | NC001549 | SUN |
| | NC001551 | FRA |
| | NC001552 | GRC |
| | NC001553 | GRC |
| | NC001554 | DZA |
| GELA | NC001555 | DEU |
| KIEVSKIJ SKOROSPELIJ | NC001556 | SUN |
| ULTRA (WB2) | NC001558 | DEU |
| HAMBURG (WB1) | NC001559 | DEU |
| KIJEVSKIJ MUTANT | NC001560 | SUN |
| | NC070967 | SDN |
| | NC070968 | ESP |
| MULTO LUPA | NC001709 | CHL |
| ASTRA | NC001710 | CHL |
| L-799-76 | NC001714 | CHL |
| KALINA | NC001716 | POL |
| LUCKY | NC001717 | FRA |
| INVERNAL BLANCA | NC001720 | Unknown |
| TIFTWHITE-78 | NC001721 | USA |
| ARKA-10 | NC001756 | USA |
| | NC070977 | PRT |
| | NC070978 | PRT |
| | NC070979 | Unknown |
| | NC070980 | ESP |
| | NC070981 | ESP |
| TOLUPA-0 | NC070982 | CHL |
| TOLUPA-7 | NC070983 | CHL |
| TOLUPA-14 | NC070984 | CHL |
| WAT | NC070985 | POL |
| S-60-17-2 | NC070986 | YUG |
| S-63-81 | NC070987 | YUG |
| S-63-276 | NC070988 | YUG |
| BLANCA | NC070989 | DEU |
| BOSNA | NC070990 | YUG |
| KIJEVSKIJ MUTANT | NC070991 | SUN |
| | NC070992 | ESP |
| | NC070993 | ESP |
| | NC070994 | ESP |
| | NC070995 | ESP |
| | NC070996 | ESP |
| | NC070997 | ESP |
| | NC070998 | ESP |
| | NC070999 | ESP |
| | NC071000 | Unknown |
| | NC071001 | ESP |
| | NC071002 | ESP |
| | NC071003 | ESP |
| | NC071004 | ESP |
| | NC071005 | ESP |
| | NC071006 | ESP |
| | NC071007 | ESP |

| | |
|---|---|
| NC071008 | ESP |
| NC071009 | ESP |
| NC071010 | ESP |
| NC071011 | ESP |
| NC071012 | ESP |
| NC071013 | ESP |
| NC071014 | ESP |
| NC071015 | ESP |
| NC071016 | ESP |
| NC071017 | ESP |
| NC071018 | ESP |
| NC071019 | ESP |
| NC071020 | ESP |
| NC071021 | ESP |
| NC071022 | ESP |
| NC071023 | ESP |
| NC071024 | ESP |
| NC001519 | ESP |
| NC071026 | ESP |
| NC071027 | BRA |
| NC071028 | MAR |
| NC071029 | CZE |
| NC071030 | IND |
| NC071031 | ESP |
| NC071032 | ESP |
| NC071033 | ESP |
| NC071034 | ESP |
| NC071035 | ESP |
| NC071036 | ESP |
| NC071037 | USA |
| NC071038 | ESP |
| NC071039 | ESP |
| NC071040 | ESP |
| NC071041 | ESP |
| NC071042 | ESP |
| NC071043 | ESP |
| NC071044 | ESP |
| NC071045 | ESP |
| NC071046 | ESP |
| NC071047 | ESP |
| NC071048 | ESP |
| NC071049 | ESP |
| NC071050 | ESP |
| NC071051 | ESP |
| NC071052 | ESP |
| NC071053 | ESP |
| NC071054 | ESP |
| NC071055 | ESP |
| NC071056 | ESP |
| NC071057 | ESP |
| NC071058 | ESP |
| NC071059 | ESP |

| | | |
|---|---|---|
| | NC071060 | ESP |
| | NC071061 | ESP |
| | NC071062 | ESP |
| | NC071063 | PRT |
| | NC071064 | ESP |
| | NC071065 | ESP |
| | NC004546 | ESP |
| | NC004595 | ESP |
| | NC006450 | ESP |
| | NC006459 | ESP |
| | NC006947 | ESP |
| | NC006952 | ESP |
| | NC006954 | ESP |
| | NC006963 | ESP |
| | NC006966 | ESP |
| | NC006978 | ESP |
| | NC006982 | ESP |
| | NC006985 | ESP |
| | NC006986 | ESP |
| | NC006998 | ESP |
| Chocho | NC002584 | ESP |
| | NC008262 | ESP |
| | NC008264 | ESP |
| | NC008265 | ESP |
| | NC008288 | ESP |
| | NC008294 | ESP |
| | NC008308 | ESP |
| | NC008311 | ESP |
| | NC008312 | ESP |
| | NC008316 | ESP |
| | NC008324 | ESP |
| | NC008329 | ESP |
| | NC008360 | ESP |
| | NC008361 | ESP |
| | NC008364 | ESP |
| | NC008368 | ESP |
| | NC008377 | ESP |
| ENTREMOSO | NC011071 | ESP |
| | NC011073 | ESP |
| | NC011074 | ESP |
| | NC011076 | ESP |
| | NC011083 | ESP |
| | NC071066 | ESP |
| | NC071067 | ESP |
| WTD 6018 | NC071068 | POL |
| | NC004463 | ESP |
| | NC004525 | ESP |
| | NC004561 | ESP |
| | NC004562 | ESP |
| | NC006311 | ESP |
| | NC006458 | ESP |
| | NC006965 | ESP |

| | | |
|---|---|---|
| | NC006971 | ESP |
| | NC006976 | ESP |
| | NC006979 | ESP |
| | NC006991 | ESP |
| | NC006993 | ESP |
| | NC008304 | ESP |
| TOCHO | NC011072 | ESP |
| | NC011077 | ESP |
| | NC011080 | ESP |
| | NC011081 | ESP |
| | NC011085 | ESP |
| | NC011086 | ESP |
| | NC011418 | PRT |
| | NC011422 | PRT |
| | NC011424 | PRT |
| | NC011425 | PRT |
| | NC011426 | PRT |
| | NC011427 | PRT |
| | NC011428 | PRT |
| | NC011429 | PRT |
| | NC011432 | PRT |
| | NC011433 | PRT |
| | NC011434 | PRT |
| | NC011437 | PRT |
| | NC011438 | PRT |
| | NC011441 | PRT |
| | NC011443 | PRT |
| | NC011446 | PRT |
| | NC011448 | PRT |
| | NC011449 | PRT |
| | NC011450 | PRT |
| | NC011451 | PRT |
| | NC011452 | PRT |
| | NC011453 | PRT |
| | NC011454 | PRT |
| | NC011455 | PRT |
| | NC011457 | PRT |
| | NC011459 | PRT |
| | NC011460 | PRT |
| | NC011463 | PRT |
| | NC011464 | PRT |
| | NC011466 | PRT |
| | NC011469 | PRT |
| | NC011471 | PRT |
| | NC011472 | PRT |
| | NC011474 | PRT |
| | NC011475 | PRT |
| | NC011476 | PRT |
| | NC011477 | PRT |
| | NC011478 | PRT |
| | NC011483 | PRT |
| | NC011495 | PRT |

| | |
|---|---|
| NC011507 | PRT |
| NC011509 | PRT |
| NC011511 | PRT |
| NC011512 | PRT |
| NC011514 | PRT |
| NC011515 | PRT |
| NC011800 | PRT |
| NC011801 | PRT |
| NC011891 | PRT |
| NC011892 | PRT |
| NC011893 | PRT |
| NC004508 | ESP |
| NC004509 | ESP |
| NC004563 | ESP |
| NC006467 | ESP |
| NC006950 | ESP |
| NC006953 | ESP |
| NC006960 | ESP |
| NC006973 | ESP |
| NC006988 | ESP |
| NC006997 | ESP |
| NC004446 | ESP |
| NC008285 | ESP |
| NC008295 | ESP |
| NC008296 | ESP |
| NC008344 | ESP |
| NC008367 | ESP |
| NC008369 | ESP |
| NC008397 | ESP |
| NC008635 | ESP |
| NC008636 | ESP |
| NC011075 | ESP |
| NC011078 | ESP |
| NC011079 | ESP |
| NC011082 | ESP |
| NC011084 | ESP |
| NC011419 | PRT |
| NC011420 | PRT |
| NC011421 | PRT |
| NC011423 | PRT |
| NC011430 | PRT |
| NC011431 | PRT |
| NC011435 | PRT |
| NC011436 | PRT |
| NC011439 | PRT |
| NC011440 | PRT |
| NC011442 | PRT |
| NC011444 | PRT |
| NC011445 | PRT |
| NC011447 | PRT |
| NC011456 | PRT |
| NC011458 | PRT |

| | |
|---|---|
| NC011461 | PRT |
| NC011462 | PRT |
| NC011465 | PRT |
| NC011467 | PRT |
| NC011468 | PRT |
| NC011470 | PRT |
| NC011473 | PRT |
| NC011479 | PRT |
| NC011480 | PRT |
| NC011481 | PRT |
| NC011482 | PRT |
| NC011484 | PRT |
| NC011485 | PRT |
| NC011486 | PRT |
| NC011487 | PRT |
| NC011488 | PRT |
| NC011489 | PRT |
| NC011490 | PRT |
| NC011491 | PRT |
| NC011493 | PRT |
| NC011494 | PRT |
| NC011496 | PRT |
| NC011497 | PRT |
| NC011498 | PRT |
| NC011499 | PRT |
| NC011500 | PRT |
| NC011501 | PRT |
| NC011502 | PRT |
| NC011503 | PRT |
| NC011504 | PRT |
| NC011505 | PRT |
| NC011506 | PRT |
| NC011508 | PRT |
| NC011510 | PRT |
| NC011513 | PRT |
| NC011516 | PRT |
| NC011517 | PRT |
| NC011518 | PRT |
| NC011519 | PRT |
| NC011520 | PRT |
| NC011521 | PRT |
| NC011522 | PRT |
| NC011799 | PRT |
| NC011802 | PRT |
| NC011803 | PRT |
| NC011804 | PRT |
| NC011805 | PRT |
| NC011806 | PRT |
| NC011807 | PRT |
| NC011808 | PRT |
| NC011809 | PRT |
| NC011810 | PRT |

| | | |
|---|---|---|
| | NC011887 | PRT |
| | NC011888 | PRT |
| | NC011889 | PRT |
| | NC011890 | PRT |
| | NC012396 | PRT |
| | NC012397 | PRT |
| | NC012398 | PRT |
| | NC071069 | ESP |
| | NC071070 | ESP |
| LUBLANC | NC000653 | FRA |
| | NC071072 | PRT |
| NIRSEGNY | NC071073 | FRA |
| | NC071074 | ESP |
| C.8 | NC071075 | FRA |
| C.9 | NC071076 | FRA |
| C.32 | NC071077 | FRA |
| | NC071078 | ESP |
| | NC006373 | ESP |
| | NC013104 | ESP |
| | NC013105 | ESP |
| TOCHO | NC013107 | ESP |
| ENTREMOÇO | NC013205 | ESP |
| | NC013853 | PRT |
| | NC013854 | PRT |
| ENTREMOÇO | NC013856 | PRT |
| | NC013857 | PRT |
| | NC013858 | PRT |
| | NC013859 | PRT |
| | NC013860 | PRT |
| | NC013861 | PRT |
| | NC013862 | PRT |
| | NC013863 | PRT |
| | NC013864 | PRT |
| | NC013865 | PRT |
| | NC013867 | PRT |
| | NC013868 | PRT |
| | NC013869 | PRT |
| | NC013892 | PRT |
| | NC013893 | PRT |
| | NC013896 | PRT |
| | NC013897 | PRT |
| | NC013899 | PRT |
| | NC013900 | PRT |
| | NC013905 | PRT |
| | NC013908 | PRT |
| | NC013914 | PRT |
| | NC013923 | PRT |
| | NC013932 | PRT |
| | NC013942 | PRT |
| | NC013946 | PRT |
| | NC013949 | PRT |
| | NC013951 | PRT |

| | | |
|---|---|---|
| | NC013988 | PRT |
| | NC013989 | PRT |
| | NC013990 | PRT |
| | NC013992 | PRT |
| | NC013993 | PRT |
| | NC013994 | PRT |
| | NC013995 | PRT |
| | NC013996 | PRT |
| | NC013997 | PRT |
| | NC013998 | PRT |
| | NC013999 | PRT |
| | NC014000 | PRT |
| | NC009671 | PRT |
| | NC009672 | PRT |
| | NC009673 | PRT |
| | NC009674 | PRT |
| | NC009675 | PRT |
| | NC009676 | PRT |
| | NC009677 | PRT |
| | NC009678 | PRT |
| | NC009679 | PRT |
| ENTREMOÇO | NC010006 | PRT |
| | NC010008 | PRT |
| | NC010117 | Unknown |
| | NC010118 | Unknown |
| | NC010119 | Unknown |
| | NC010120 | Unknown |
| | NC010125 | Unknown |
| | NC010130 | Unknown |
| | NC010131 | Unknown |
| | NC010132 | Unknown |
| | NC010133 | Unknown |
| | NC010134 | Unknown |
| | NC010135 | Unknown |
| | NC010136 | Unknown |
| | NC010137 | Unknown |
| | NC010138 | Unknown |
| | NC010139 | Unknown |
| | NC010140 | Unknown |
| | NC010141 | Unknown |
| | NC010142 | Unknown |
| | NC010143 | Unknown |
| | NC010144 | Unknown |
| | NC010145 | Unknown |
| | NC010146 | Unknown |
| | NC010147 | Unknown |
| | NC010148 | Unknown |
| | NC010149 | Unknown |
| | NC010150 | Unknown |
| | NC010151 | Unknown |
| | NC010153 | Unknown |
| | NC010154 | Unknown |

| | | |
|---|---|---|
| | NC010155 | Unknown |
| | NC010156 | Unknown |
| | NC010157 | Unknown |
| | NC010158 | Unknown |
| | NC010159 | Unknown |
| | NC010160 | Unknown |
| | NC010161 | Unknown |
| | NC010162 | Unknown |
| | NC010163 | Unknown |
| | NC010164 | Unknown |
| | NC010165 | Unknown |
| | NC010166 | Unknown |
| | NC010167 | Unknown |
| | NC010168 | Unknown |
| | NC010169 | Unknown |
| | NC010170 | Unknown |
| | NC010171 | Unknown |
| | NC010172 | Unknown |
| | NC010247 | ESP |
| | NC010248 | PRT |
| | NC010250 | PRT |
| | NC010251 | ESP |
| | NC010253 | ESP |
| | NC010254 | ESP |
| | NC010255 | ESP |
| | NC010256 | ESP |
| | NC010257 | ESP |
| | NC010258 | ESP |
| | NC010259 | ESP |
| | NC010260 | ESP |
| | NC010261 | ESP |
| | NC010262 | ESP |
| Altramuz | NC010263 | ESP |
| Altramuz | NC010264 | ESP |
| Altramuz | NC010265 | ESP |
| Altramuz | NC010266 | ESP |
| Altramuz | NC010267 | ESP |
| Altramuz | NC010268 | ESP |
| Altramuz | NC010269 | ESP |
| Altramuz | NC010270 | ESP |
| Altramuz | NC010272 | ESP |
| Altramuz | NC010273 | ESP |
| Altramuz | NC010274 | ESP |
| Chocho | NC010275 | ESP |
| Chocho | NC010276 | ESP |
| Chocho | NC010277 | ESP |
| Chocho | NC010278 | ESP |
| Chocho | NC010279 | ESP |
| Chocho | NC010280 | ESP |
| Chocho | NC010281 | ESP |
| | NC010282 | PRT |
| | NC010283 | PRT |

| | | |
|---|---|---|
| | NC010284 | PRT |
| | NC010285 | PRT |
| | NC010286 | PRT |
| | NC010287 | PRT |
| | NC010126 | Unknown |
| | NC010152 | Unknown |
| LUTOP | NC071079 | FRA |
| LUCROP | NC071080 | FRA |
| Altramuz | NC071081 | ESP |
| TOCHO | NC013106 | ESP |
| | NC013108 | ESP |
| ENTREMOÇO | NC013855 | PRT |
| ESTORIL | NC071082 | PRT |
| MURTAL | NC071083 | PRT |
| | NC010288 | PRT |
| | NC010289 | PRT |
| | NC010290 | PRT |
| | NC010291 | PRT |
| | NC010292 | PRT |
| | NC010293 | PRT |
| | NC010294 | PRT |
| | NC010295 | PRT |
| | NC010296 | PRT |
| | NC010297 | PRT |
| | NC010298 | PRT |
| | NC010299 | PRT |
| | NC010300 | PRT |
| | NC010301 | PRT |
| | NC010303 | PRT |
| | NC010304 | PRT |
| | NC010305 | PRT |
| | NC010306 | PRT |
| | NC010307 | PRT |
| | NC010308 | PRT |
| | NC010309 | PRT |
| | NC010310 | PRT |
| | NC010311 | PRT |
| | NC010312 | PRT |
| | NC010313 | PRT |
| | NC010314 | PRT |
| | NC010315 | PRT |
| | NC010316 | PRT |
| | NC010317 | PRT |
| | NC010318 | PRT |
| | NC010319 | PRT |
| | NC010320 | PRT |
| | NC010321 | PRT |
| | NC010322 | PRT |
| | NC010323 | PRT |
| | NC010324 | PRT |
| | NC010325 | PRT |
| | NC010326 | PRT |

| | | |
|---|---|---|
| | NC010327 | PRT |
| | NC010328 | PRT |
| | NC010330 | PRT |
| | NC010331 | PRT |
| | NC010332 | PRT |
| | NC010333 | PRT |
| | NC010334 | PRT |
| | NC010335 | PRT |
| | NC010336 | PRT |
| | NC010337 | PRT |
| | NC010338 | PRT |
| | NC010339 | PRT |
| | NC010340 | PRT |
| | NC010341 | PRT |
| | NC010342 | PRT |
| | NC010343 | PRT |
| | NC010344 | PRT |
| | NC010345 | PRT |
| | NC010346 | PRT |
| | NC010347 | PRT |
| | NC010348 | PRT |
| | NC010349 | PRT |
| | NC010350 | PRT |
| | NC010351 | PRT |
| | NC010352 | PRT |
| | NC010353 | PRT |
| | NC010354 | PRT |
| | NC010355 | PRT |
| Altramuz | NC010356 | ESP |
| Altramuz | NC010357 | ESP |
| Altramuz | NC010358 | ESP |
| Altramuz | NC010359 | ESP |
| Altramuz | NC010360 | ESP |
| Altramuz | NC010361 | ESP |
| Altramuz | NC010362 | ESP |
| Altramuz | NC010363 | ESP |
| Altramuz | NC010364 | ESP |
| Altramuz | NC010365 | ESP |
| Altramuz | NC010366 | ESP |
| Altramuz | NC010367 | ESP |
| Altramuz | NC010368 | ESP |
| Altramuz | NC010369 | ESP |
| Altramuz | NC010370 | ESP |
| Altramuz | NC010371 | ESP |
| Altramuz | NC010372 | ESP |
| Altramuz | NC010373 | ESP |
| Altramuz | NC010374 | ESP |
| Altramuz | NC010375 | ESP |
| Altramuz | NC010376 | ESP |
| Altramuz | NC010377 | ESP |
| Altramuz | NC010378 | ESP |
| Altramuz | NC010379 | ESP |

| | | |
|---|---|---|
| Altramuz | NC010380 | ESP |
| Altramuz | NC010381 | ESP |
| Altramuz | NC010382 | ESP |
| Altramuz | NC010383 | ESP |
| Altramuz | NC010384 | ESP |
| Altramuz | NC010385 | ESP |
| Altramuz | NC010386 | ESP |
| Altramuz | NC010387 | ESP |
| LLOBI | NC010388 | ESP |
| LLOBI | NC010391 | ESP |
| LLOBI | NC010392 | ESP |
| Chocho | NC010393 | ESP |
| Chocho | NC010394 | ESP |
| | NC018159 | PRT |
| | NC018160 | PRT |
| | NC018161 | PRT |
| | NC018162 | PRT |
| | NC018163 | PRT |
| | NC018164 | PRT |
| | NC018165 | PRT |
| | NC018166 | PRT |
| | NC018167 | PRT |
| Altramuz | NC004566 | ESP |
| Chocho | NC008880 | ESP |
| Chocho | NC008881 | ESP |
| Chocho | NC011031 | ESP |
| TOCHO | NC011032 | ESP |
| | NC010112 | Unknown |
| | NC010113 | Unknown |
| | NC010115 | Unknown |
| | NC010116 | Unknown |
| | NC010122 | Unknown |
| | NC010123 | Unknown |
| | NC018168 | PRT |
| | NC018169 | PRT |
| | NC018170 | PRT |
| | NC018171 | PRT |
| | NC018172 | PRT |
| | NC018173 | PRT |
| | NC018174 | PRT |
| | NC018175 | PRT |
| | NC018176 | PRT |
| | NC018177 | PRT |
| | NC018178 | PRT |
| | NC018179 | PRT |
| | NC018180 | PRT |
| | NC018181 | PRT |
| Chocho | NC018663 | ESP |
| Chocho | NC018664 | ESP |
| Chocho | NC018665 | ESP |
| Chocho | NC018666 | ESP |
| RIO-MAIOR | NC071084 | PRT |

| | | |
|---|---|---|
| | NC071085 | PRT |
| B-101 | NC071086 | SUN |
| | NC071087 | ESP |
| | NC071088 | ESP |
| | NC071089 | ESP |
| | NC071090 | ESP |
| | NC071091 | ESP |
| | NC071092 | ESP |
| | NC071093 | ESP |
| | NC071094 | ESP |
| | NC071095 | ESP |
| | NC071096 | ESP |
| | NC071097 | ESP |
| | NC071098 | ESP |
| | NC071099 | ESP |
| | NC071100 | ESP |
| | NC071101 | ESP |
| | NC071102 | ESP |
| 1131 | NC071103 | SUN |
| | NC071104 | ESP |
| | NC001550 | FRA |
| WTD 6020 | NC071106 | POL |
| | NC071107 | ESP |
| TOCHO | NC011033 | ESP |
| | NC015970 | GRC |
| | NC015974 | GRC |
| | NC010121 | Unknown |
| | NC010127 | Unknown |
| | NC010129 | Unknown |
| | NC019649 | ESP |
| Chocho | NC019717 | ESP |
| Altramuz amargo | NC020239 | ESP |
| Chocho | NC026130 | ESP |
| TOCHO | NC019628 | ESP |
| Altramuz | NC020285 | ESP |
| Chocho | NC035661 | ESP |
| Chocho | NC050273 | ESP |
| | NC050468 | ESP |
| Altramuz | NC052301 | ESP |
| Chocho | NC052354 | ESP |
| Altramuz | NC052454 | ESP |
| Altramuz | NC052524 | ESP |
| Altramuz | NC052548 | ESP |
| | NC015969 | GRC |
| Chocho | NC052727 | ESP |
| Chocho | NC035663 | ESP |
| | NC015966 | GRC |
| | NC015968 | GRC |
| | NC015971 | GRC |
| | NC015972 | GRC |
| | NC015973 | GRC |
| | NC015975 | GRC |

| | | |
|---|---|---|
| | NC015976 | GRC |
| | NC015977 | GRC |
| Chocho | NC015978 | ESP |
| Chocho | NC027245 | ESP |
| Altramuz | NC000615 | ESP |
| | NC044876 | ITA |
| | NC044229 | ESP |
| Altramuz | NC054628 | ESP |
| Chocho | NC068892 | ESP |
| Chocho | NC068905 | ESP |
| Chocho | NC069104 | ESP |
| Chocho | NC069124 | ESP |
| TOCHO | NC069356 | ESP |
| Altramuz | NC069482 | ESP |
| | NC010114 | Unknown |
| | NC010124 | Unknown |
| Chocho | NC008879 | ESP |
| | NC010128 | Unknown |
| Altramuz | NC054629 | ESP |
| Chocho | NC076208 | ESP |
| Chocho | NC076357 | ESP |
| Chocho | NC076363 | ESP |
| | NC076446 | ESP |
| Altramuz amargoso | NC077606 | ESP |
| Altramuz | NC077764 | ESP |
| | NC077827 | ESP |
| | NC078076 | ESP |
| NELLY | NC084535 | HUN |
| | NC082340 | ESP |
| | NC080642 | ESP |
| | NC080572 | ESP |
| | NC082389 | ESP |
| | NC082394 | ESP |
| ALTRAMUZ DULCE | NC052296 | ESP |
| | NC094443 | ESP |
| | NC094450 | ESP |
| | NC094460 | PRT |
| | NC097900 | PRT |
| | NC097901 | ESP |
| MARTA | NC110722 | Unknown |
| XA100 | W6 15551 | FRA |
| RUMBO BAER | W6 19548 | CHL |
| NELLY | W6 24907 | HUN |
| LUP 2044 | W6 39797 | DEU |
| LUP 2046 | W6 39798 | DEU |
| LUP 2079 | W6 39803 | NLD |
| LUP 2081 | W6 39805 | DEU |
| LUP 2082 | W6 39806 | RUS |
| LUP 2086 | W6 39807 | ITA |
| LUP 229 | W6 39811 | DEU |
| LUP 235 | W6 39815 | DEU |
| LUP 252 | W6 39818 | GEO |

| | | |
|---|---|---|
| LUP 253 | W6 39819 | DEU |
| LUP 257 | W6 39821 | UKR |
| LUP 6462 | W6 39909 | DEU |
| | PI 168891 | NLD |
| ME 115 | PI 170528 | TUR |
| | PI 179361 | TUR |
| GYULATANYAI EDES | PI 232924 | HUN |
| PFLUG-MANSA | PI 237719 | DEU |
| | PI 243335 | ZAF |
| MN 3 | PI 244572 | Unknown |
| TERMIS | PI 250094 | EGY |
| | PI 250572 | EGY |
| MN 181 | PI 251559 | SRB |
| | PI 255375 | SRB |
| | PI 255471 | SRB |
| MN 48 | PI 287241 | DEU |
| KRAFTQUELL | PI 289160 | HUN |
| SAATGUT | PI 316610 | DEU |
| | PI 337089 | BRA |
| 22840 | PI 368911 | CZE |
| 31620 | PI 368914 | BGR |
| 31621 | PI 368915 | BGR |
| KIJEVSKIJ MUTANT | PI 381322 | SUN |
| KALI | PI 386098 | POL |
| KALI | PI 434855 | NZL |
| ULTRA | PI 434856 | NZL |
| GR 1 | PI 457921 | GRC |
| GR 3 | PI 457923 | GRC |
| GR 5 | PI 457924 | GRC |
| GR 7 | PI 457926 | GRC |
| GR 8 | PI 457927 | GRC |
| GR 9 | PI 457928 | GRC |
| GR 10 | PI 457929 | GRC |
| GR 11 | PI 457930 | GRC |
| GR 12 | PI 457931 | GRC |
| GR 13 | PI 457932 | GRC |
| LA 25 | PI 457933 | FRA |
| LA 26 | PI 457934 | FRA |
| LA 27 | PI 457935 | FRA |
| LA 56 | PI 457936 | GRC |
| LA 57 | PI 457937 | GRC |
| 9483 | PI 457938 | MAR |
| LA 70 | PI 457939 | ESP |
| LA 71 | PI 457940 | DZA |
| 1082 | PI 457941 | ESP |
| 1107 | PI 457942 | ESP |
| 1186 | PI 457943 | ESP |
| 1134 | PI 457944 | ESP |
| 1190 | PI 457945 | ESP |
| 1585 | PI 457946 | ESP |
| 1586 | PI 457947 | ESP |
| 1587 | PI 457948 | ESP |

| | | |
|---|---|---|
| 1588 | PI 457949 | ESP |
| 1589 | PI 457950 | ESP |
| 1590 | PI 457951 | ESP |
| 1591 | PI 457952 | ESP |
| 1592 | PI 457953 | ESP |
| 1593 | PI 457954 | ESP |
| 1594 | PI 457955 | ESP |
| VIR 1423 | PI 457956 | RUS |
| LA 105 | PI 457958 | ITA |
| LA 106 | PI 457959 | ITA |
| LUBLANC | PI 467348 | FRA |
| LUCKY | PI 467349 | FRA |
| C 9 | PI 467351 | FRA |
| WAT | PI 468127 | POL |
| KALINA | PI 468128 | POL |
| WTD 180 | PI 468129 | POL |
| HAMBURG | PI 469095 | AUS |
| ULTRA | PI 469096 | AUS |
| VIR 1642 | PI 476370 | SDN |
| SHARKEIA K | PI 476371 | EGY |
| KIEVSKIJ SKOROSPELIJ | PI 476372 | UKR |
| GORIZONT | PI 476373 | UKR |
| KALINA | PI 476374 | POL |
| KALI | PI 476375 | POL |
| PRIMORSKIJ | PI 476376 | UKR |
| M-2738 | PI 480483 | LBN |
| VIR 494 | PI 481545 | ETH |
| VIR 1437 | PI 481546 | GRC |
| VIR 1504 | PI 481547 | HUN |
| BIALY POZNY | PI 481548 | POL |
| VIR 1550 | PI 481549 | BGR |
| VIR 1600 | PI 481550 | ITA |
| EGYPTICA BALADY | PI 481551 | EGY |
| VIR 1644 | PI 481552 | SDN |
| HADMERSLEBENER NAHRQUELL | PI 481553 | DEU |
| PETKUSER BITTERE WEISLUPINE | PI 481554 | DEU |
| WEISE BITTERLUPINE | PI 481555 | DEU |
| VIR 2005 | PI 481556 | MAR |
| VIR 1820 | PI 481557 | CZE |
| VIR 2229 | PI 481558 | SYR |
| VIR 2362 | PI 481559 | ESP |
| VIR 2374 | PI 481560 | SRB |
| IFLU 29 | PI 483072 | SYR |
| IFLU 31 | PI 483073 | MAR |
| IFLU 32 | PI 483074 | LBN |
| IFLU 33 | PI 483075 | SYR |
| | PI 487432 | JOR |
| BLANCA | PI 487433 | DEU |
| KIEV EARLY | PI 487434 | SUN |
| KIEV N409 | PI 487435 | SUN |
| CPI 31620 | PI 487437 | BGR |
| | PI 502648 | DEU |

| | | |
|---|---|---|
| ASTRA | PI 502649 | CHL |
| KALI | PI 502650 | POL |
| KIJEVSKIJ MUTANT | PI 502651 | SUN |
| ME 74 | PI 502652 | HUN |
| BIALY 1 | PI 505844 | POL |
| NOSOVSKIJ-3 | PI 505845 | UKR |
| KIJEVSKIJ MUTANT | PI 505846 | UKR |
| LOTOS | PI 505847 | UKR |
| SOLNECNYJ | PI 505848 | SUN |
| START | PI 508101 | RUS |
| GR 327 | PI 516622 | MAR |
| GR 333 | PI 516623 | MAR |
| GR 334 | PI 516624 | ITA |
| GR 337 | PI 516625 | DEU |
| GR 338 | PI 516626 | DEU |
| GR 339 | PI 516627 | ITA |
| GR 342 | PI 516629 | MAR |
| GR 343 | PI 516630 | DEU |
| GR 335 | PI 516639 | MAR |
| NO. 267 | PI 533694 | ESP |
| NO. 269 | PI 533695 | ESP |
| VIR 2603 | PI 533696 | UKR |
| NO. 530 | PI 533697 | ESP |
| NO. 544 | PI 533698 | ESP |
| NO. 556 | PI 533699 | ESP |
| NO. 558 | PI 533700 | ESP |
| NO. 571 | PI 533701 | ESP |
| NO. 576 | PI 533702 | ESP |
| NO. 584 | PI 533703 | ESP |
| R-6002;NORTO 486 | PI 533704 | ESP |
| R-6019;NORTO 484 | PI 533705 | ESP |
| NO. 47 | PI 533706 | ESP |
| NO. 175 | PI 533707 | ESP |
| 870529-02 | PI 533714 | ESP |
| NO. 22 | PI 543013 | ESP |
| NO. 24 | PI 543014 | ESP |
| NO. 47 | PI 543015 | ESP |
| NO. 53 | PI 543016 | ESP |
| NO. 189 | PI 543018 | ESP |
| NO. 272 | PI 543019 | ESP |
| NO. 274 | PI 543020 | ESP |
| NO. 539 | PI 543022 | ESP |
| NO. 561 | PI 543023 | ESP |
| NO. 563 | PI 543024 | ESP |
| NO. 570 | PI 543026 | ESP |
| NO. 572 | PI 543027 | ESP |
| NO. 573 | PI 543028 | ESP |
| LUPINUS ALBUS | PI 587203 | USA |
| HOPE | PI 594916 | USA |
| LINE NO 10 | PI 606481 | USA |
| LUCKY | PI 606482 | ESP |
| MULTULUPA | PI 606483 | ESP |

| | | |
|---|---|---|
| E92-2 | PI 606484 | EGY |
| E92-13 | PI 606485 | EGY |
| LINE NO 2 | PI 615401 | USA |
| LINE NO 3 | PI 615402 | USA |
| LINE NO 5 | PI 615403 | USA |
| LINE NO 6 | PI 615404 | USA |
| LINE NO 7 | PI 615405 | USA |
| LINE NO 8 | PI 615406 | USA |
| TIFWHITE-78 | PI 615409 | USA |
| WJK-3 | PI 660685 | ESP |
| CPI 122632 | PI 660711 | ESP |
| CPI 125778 | PI 660712 | ETH |
| P27435 | PI 660713 | SYR |
| P27715 | PI 660714 | EGY |
| ALBAN | PI 660721 | FRA |
| ADAM | PI 660722 | FRA |
| PMR 182 | PI 660723 | USA |
| SACCHARATUS | PI 660724 | JOR |
| MJS 374(L0477P) | PI 660726 | AUS |
| WJK-2 | PI 666380 | ESP |
| WJK-4 | PI 666381 | ESP |
| WJK-5 | PI 666382 | ESP |
| P2061/GELA | PI 666385 | AUS |
| ALB.01 | PI 666386 | DZA |
| | PI 673381 | DEU |
| | PI 673382 | DEU |
| | PI 673383 | DEU |
| | PI 673384 | DEU |
| | PI 673385 | PRT |
| | PI 673386 | PAL |
| | PI 673387 | DEU |
| | PI 673388 | DEU |
| | PI 673389 | DEU |
| | PI 673390 | DEU |
| | PI 673391 | DEU |
| | PI 673392 | DEU |
| | PI 673393 | POL |
| ASTER | Aster | FRA |
| CLOVIS | Clovis | Unknown |
| ORUS | Orus | Unknown |
| MAGNUS | Magnus | Unknown |
| ULYSSE | Ulysse | Unknown |
| 29-2 | 29-2 | Unknown |
| 433 | 433 | Unknown |
| LUXE | Luxe | Unknown |
| LUMEN | Lumen | Unknown |
| ENERGY | Energy | Unknown |
| FIGARO | Figaro | Unknown |
| SULIMO | Sulimo | Unknown |
| 557-8A | 557-8a | Unknown |
| 476B | 476b | EGY |
| | Lupino Multitalia | Unknown |

| | |
|---|---|
| Lupino Gigante di Vai | Unknown |
| Lupino di Recanati | ITA |

| GEO_REGION | BIOSTAT | Cluster |
|---|---|---|
| Southern Europe | Wild | A_1 |
| Southern Europe | Cultivar | A_1 |
| Southern Europe | Wild | A_2 |
| Southern Europe | Landrace | A_1 |
| Southern Europe | Wild | A_1 |
| Southern Europe | Landrace | A_1 |
| Southern Europe | Wild | A_2 |
| Southern Europe | Wild | A_1 |
| Southern Europe | Landrace | A_1 |
| Southern Europe | Wild | A_1 |
| Southern Europe | Wild | A_2 |
| Southern Europe | Landrace | A_1 |
| Southern Europe | Wild | A_1 |
| Southern Europe | Wild | A_1 |
| Southern Europe | Wild | A_1 |
| Southern Europe | Wild | A_1 |
| Northern Africa | Landrace | A_1 |
| Northern Africa | Landrace | A_1 |
| Northern Africa | Landrace | A_2 |
| Northern Africa | Landrace | A_2 |
| Southern Europe | Landrace | A_1 |
| Southern Europe | Landrace | A_1 |
| Southern Europe | Wild | A_2 |
| Southern Europe | Wild | A_1 |
| Southern Europe | Wild | A_2 |
| Southern Europe | Wild | A_1 |
| Southern Europe | Wild | A_2 |
| Southern Europe | Wild | A_2 |
| Southern Europe | Wild | A_2 |
| Southern Europe | Wild | A_1 |
| Southern Europe | Wild | A_1 |
| Southern Europe | Wild | A_2 |
| Southern Europe | Wild | A_1 |
| Southern Europe | Wild | A_1 |
| Southern Europe | Wild | A_1 |
| Southern Europe | Wild | A_1 |
| Southern Europe | Wild | A_1 |
| Southern Europe | Wild | A_1 |
| Southern Europe | Wild | A_1 |
| Southern Europe | Wild | A_2 |
| Southern Europe | Wild | A_1 |
| Southern Europe | Wild | A_1 |
| Southern Europe | Wild | A_1 |
| Southern Europe | Wild | A_2 |
| Southern Europe | Wild | A_1 |
| Southern Europe | Wild | A_1 |
| Northern Africa | Landrace | A_1 |
| Northern Africa | Landrace | A_1 |
| Northern Africa | Landrace | A_1 |
| Northern Africa | Landrace | A_2 |

| | | |
|---|---|---|
| Northern Africa | Landrace | A_1 |
| Northern Africa | Landrace | A_1 |
| Southern Europe | Wild | A_2 |
| Southern Europe | Wild | A_3 |
| Southern Europe | Wild | A_2 |
| Southern Europe | Cultivar | A_2 |
| Western Europe | Wild | A_1 |
| Southern Europe | Wild | A_1 |
| Southern Europe | Wild | A_2 |
| Southern Europe | Wild | A_1 |
| Southern Europe | Wild | A_1 |
| Southern Europe | Wild | A_1 |
| Southern Europe | Landrace | A_1 |
| Western Europe | Landrace | A_1 |
| Southern Europe | Wild | A_1 |
| Southern Europe | Wild | A_1 |
| Southern Europe | Landrace | A_1 |
| Eastern Africa | Breeding/Research Material | A_1 |
| Southern Europe | Wild | A_1 |
| Southern Europe | Landrace | A_2 |
| Southern Europe | Landrace | A_1 |
| Southern Europe | Landrace | A_2 |
| Southern Europe | Landrace | A_1 |
| Southern Europe | Landrace | A_1 |
| Southern Europe | Landrace | A_1 |
| Southern Europe | Landrace | A_1 |
| Southern Europe | Landrace | A_1 |
| Eastern Europe | Breeding/Research Material | A_1 |
| Southern Europe | Wild | A_1 |
| Southern Europe | Wild | A_1 |
| Southern Europe | Wild | A_2 |
| Western Europe | Wild | A_1 |
| Middle East | Wild | A_1 |
| America | Landrace | A_1 |
| Middle East | Wild | A_1 |
| Middle East | Wild | A_2 |
| Eastern Africa | Wild | A_1 |
| Middle East | Wild | A_1 |
| Middle East | Wild | A_2 |
| Middle East | Wild | A_2 |
| Middle East | Wild | A_2 |
| America | Wild | A_1 |
| America | Wild | A_1 |
| Middle East | Wild | A_1 |
| Eastern Europe | Breeding/Research Material | A_1 |
| Eastern Europe | Cultivar | A_1 |
| Southern Europe | Wild | A_1 |
| Northern Africa | Wild | A_1 |
| Western Europe | Wild | A_1 |
| Eastern Africa | Wild | A_1 |
| Southern Europe | Wild | A_1 |
| Western Europe | Cultivar | A_1 |

| | | |
|---|---|---|
| Southern Europe | Wild | A_1 |
| Southern Europe | Wild | A_1 |
| Western Europe | Cultivar | A_1 |
| Western Europe | Cultivar | A_1 |
| Eastern Europe | Breeding/Research Material | A_1 |
| Eastern Europe | Breeding/Research Material | B |
| Eastern Europe | Breeding/Research Material | A_1 |
| Southern Europe | Landrace | A_1 |
| Western Europe | Cultivar | A_1 |
| Eastern Europe | Cultivar | A_1 |
| Eastern Europe | Cultivar | A_1 |
| Eastern Europe | Breeding/Research Material | A_1 |
| Eastern Europe | Breeding/Research Material | A_2 |
| Eastern Europe | Wild | A_1 |
| Eastern Europe | Cultivar | A_2 |
| Eastern Europe | Cultivar | A_1 |
| America | Cultivar | A_1 |
| Eastern Europe | Cultivar | A_1 |
| Eastern Europe | Breeding/Research Material | A_1 |
| Eastern Europe | Breeding/Research Material | A_1 |
| Western Europe | Advanced/improved cultivar | A_1 |
| Eastern Africa | Cultivar | A_2 |
| Western Europe | Cultivar | A_1 |
| Eastern Europe | Landrace | B |
| Eastern Europe | Breeding/Research Material | A_1 |
| Eastern Europe | Cultivar | A_1 |
| Eastern Europe | Advanced/improved cultivar | A_1 |
| Southern Europe | Wild | A_1 |
| Southern Europe | Wild | A_1 |
| Southern Europe | Wild | A_1 |
| Southern Europe | Wild | A_1 |
| Southern Europe | Wild | A_1 |
| Southern Europe | Wild | A_1 |
| Southern Europe | Wild | A_1 |
| Southern Europe | Wild | A_1 |
| Southern Europe | Wild | A_1 |
| Southern Europe | Landrace | A_1 |
| Southern Europe | Wild | A_1 |
| Southern Europe | Wild | A_1 |
| Southern Europe | Wild | A_1 |
| Southern Europe | Wild | A_1 |
| Southern Europe | Wild | A_1 |
| Southern Europe | Wild | A_1 |
| Southern Europe | Wild | A_1 |
| Southern Europe | Wild | A_1 |
| Southern Europe | Wild | A_1 |
| Southern Europe | Wild | A_1 |
| Southern Europe | Wild | A_1 |
| Southern Europe | Wild | A_1 |
| Southern Europe | Wild | A_1 |
| Southern Europe | Landrace | A_1 |
| Southern Europe | Wild | A_1 |

| | | |
|---|---|---|
| Southern Europe | Wild | A_1 |
| Southern Europe | Wild | A_1 |
| Southern Europe | Wild | A_1 |
| Southern Europe | Wild | A_1 |
| Southern Europe | Wild | A_1 |
| Southern Europe | Wild | A_1 |
| Southern Europe | Landrace | A_1 |
| Southern Europe | Landrace | A_1 |
| Southern Europe | Landrace | A_1 |
| Southern Europe | Wild | A_1 |
| Southern Europe | Wild | A_3 |
| Southern Europe | Wild | A_1 |
| Southern Europe | Wild | A_1 |
| Western Europe | Landrace | A_1 |
| Southern Europe | Landrace | A_1 |
| Southern Europe | Breeding/Research Material | A_1 |
| Southern Europe | Wild | A_1 |
| Southern Europe | Wild | A_1 |
| Southern Europe | Wild | A_1 |
| Southern Europe | Wild | A_1 |
| Southern Europe | Wild | A_1 |
| Southern Europe | Wild | A_1 |
| Southern Europe | Wild | A_1 |
| Southern Europe | Landrace | A_1 |
| Southern Europe | Wild | A_1 |
| Southern Europe | Wild | A_1 |
| Southern Europe | Wild | A_1 |
| Southern Europe | Wild | A_1 |
| Southern Europe | Wild | A_1 |
| Southern Europe | Wild | A_1 |
| Southern Europe | Wild | A_1 |
| Southern Europe | Breeding/Research Material | A_1 |
| Southern Europe | Landrace | A_3 |
| Southern Europe | Landrace | A_1 |
| Southern Europe | Landrace | A_1 |
| Southern Europe | Landrace | A_1 |
| Southern Europe | Landrace | A_1 |
| Southern Europe | Landrace | A_1 |
| Southern Europe | Landrace | A_1 |
| Southern Europe | Landrace | A_1 |
| Southern Europe | Landrace | A_1 |
| Southern Europe | Landrace | A_1 |
| Southern Europe | Landrace | A_1 |
| Southern Europe | Landrace | A_1 |
| Eastern Europe | Landrace | B |
| Southern Europe | Landrace | A_1 |
| Eastern Europe | Landrace | A_1 |
| Eastern Europe | Breeding/Research Material | A_1 |
| Eastern Europe | Breeding/Research Material | A_1 |
| Eastern Europe | Breeding/Research Material | B |
| Southern Europe | Wild | A_1 |
| Southern Europe | Wild | A_1 |

| | | |
|---|---|---|
| Southern Europe | Wild | A_1 |
| Southern Europe | Wild | A_2 |
| Southern Europe | Wild | A_1 |
| Southern Europe | Wild | A_1 |
| Southern Europe | Wild | A_1 |
| Southern Europe | Wild | A_3 |
| Southern Europe | Wild | A_1 |
| Southern Europe | Wild | A_3 |
| Southern Europe | Wild | A_1 |
| Southern Europe | Wild | A_1 |
| Southern Europe | Wild | A_1 |
| Southern Europe | Wild | A_1 |
| Southern Europe | Wild | A_1 |
| Northern Africa | Wild | A_1 |
| Southern Europe | Wild | A_1 |
| Southern Europe | Wild | A_1 |
| Southern Europe | Wild | A_1 |
| Southern Europe | Wild | A_1 |
| America | Cultivar | A_1 |
| Western Europe | Landrace | A_1 |
| Middle East | Landrace | A_1 |
| Middle East | Landrace | A_1 |
| Middle East | Landrace | A_1 |
| Northern Africa | Landrace | A_3 |
| Northern Africa | Landrace | A_1 |
| Middle East | Landrace | A_1 |
| Middle East | Wild | A_2 |
| America | Landrace | A_1 |
| Western Europe | Landrace | A_1 |
| Unknown | Breeding/Research Material | A_1 |
| America | Landrace | A_1 |
| Middle East | Landrace | A_3 |
| Southern Europe | Wild | A_1 |
| Southern Europe | Wild | A_1 |
| Southern Europe | Wild | A_1 |
| Northern Africa | Wild | A_1 |
| Eastern Africa | Wild | A_1 |
| Western Europe | Cultivar | A_1 |
| Middle East | Wild | A_3 |
| Middle East | Wild | A_1 |
| Middle East | Wild | A_1 |
| Northern Africa | Wild | A_1 |
| America | Wild | A_1 |
| Middle East | Wild | A_1 |
| Southern Europe | Wild | A_3 |
| Southern Europe | Cultivar | A_1 |
| Middle East | Cultivar | A_1 |
| Southern Europe | Cultivar | A_1 |
| Southern Europe | Cultivar | A_1 |
| Eastern Europe | Cultivar | A_1 |
| Eastern Europe | Cultivar | A_1 |
| Western Europe | Cultivar | A_3 |

| | | |
|---|---|---|
| Western Europe | Cultivar | A_1 |
| Eastern Africa | Cultivar | A_1 |
| Eastern Europe | Unknown | A_1 |
| Eastern Europe | Breeding/Research Material | A_1 |
| Eastern Europe | Unknown | A_1 |
| Eastern Europe | Breeding/Research Material | A_1 |
| Eastern Europe | Breeding/Research Material | A_1 |
| Southern Europe | Breeding/Research Material | A_1 |
| Unknown | Breeding/Research Material | A_1 |
| Western Europe | Cultivar | A_1 |
| Western Europe | Cultivar | A_1 |
| Western Europe | Cultivar | A_1 |
| Western Europe | Cultivar | A_1 |
| Eastern Europe | Wild | A_1 |
| Eastern Europe | Wild | A_1 |
| Eastern Europe | Cultivar | A_1 |
| Western Europe | Advanced/improved cultivar | A_1 |
| Southern Europe | Cultivar | A_1 |
| Southern Europe | Cultivar | A_1 |
| Eastern Africa | Cultivar | A_1 |
| Eastern Europe | Cultivar | A_3 |
| Eastern Europe | Cultivar | A_3 |
| Western Europe | Cultivar | A_1 |
| Eastern Europe | Cultivar | A_1 |
| Eastern Europe | Cultivar | A_1 |
| Eastern Europe | Cultivar | A_2 |
| Western Europe | Cultivar | A_1 |
| Middle East | Landrace | A_1 |
| Middle East | Landrace | A_1 |
| America | Cultivar | A_1 |
| Eastern Europe | Breeding/Research Material | A_1 |
| Western Europe | Cultivar | A_1 |
| Western Europe | Cultivar | A_1 |
| Western Europe | Cultivar | A_1 |
| Western Europe | Cultivar | A_1 |
| Eastern Europe | Wild | A_1 |
| Eastern Europe | Cultivar | A_1 |
| Eastern Europe | Cultivar | A_1 |
| Eastern Europe | Cultivar | A_1 |
| Eastern Europe | Breeding/Research Material | A_3 |
| Eastern Europe | Breeding/Research Material | A_1 |
| Eastern Europe | Breeding/Research Material | A_1 |
| Eastern Europe | Breeding/Research Material | A_1 |
| Western Europe | Breeding/Research Material | A_1 |
| Western Europe | Cultivar | A_1 |
| Western Europe | Cultivar | A_1 |
| Eastern Africa | Cultivar | A_1 |
| Southern Europe | Unknown | A_1 |
| Southern Europe | Unknown | A_1 |
| Middle East | Unknown | A_2 |
| Eastern Africa | Unknown | A_1 |
| Eastern Africa | Unknown | A_1 |

| | | |
|---|---|---|
| Eastern Africa | Unknown | A_1 |
| Eastern Africa | Unknown | A_1 |
| Middle East | Unknown | A_1 |
| Middle East | Unknown | A_2 |
| Middle East | Unknown | A_2 |
| Middle East | Unknown | A_1 |
| Western Europe | Unknown | A_1 |
| Northern Africa | Unknown | A_1 |
| Northern Africa | Unknown | A_1 |
| Northern Africa | Unknown | A_1 |
| Eastern Europe | Advanced/improved cultivar | A_1 |
| Southern Europe | Wild | B |
| Southern Europe | Wild | B |
| Southern Europe | Wild | B |
| Southern Europe | Wild | B |
| Southern Europe | Wild | B |
| Southern Europe | Wild | B |
| Unknown | Landrace | A_1 |
| Unknown | Landrace | A_1 |
| Southern Europe | Wild | B |
| Eastern Europe | Wild | A_1 |
| Southern Europe | Traditional cultivar/Landrace | A_1 |
| Southern Europe | Advanced/improved cultivar | A_2 |
| Southern Europe | Traditional cultivar/Landrace | A_2 |
| Southern Europe | Advanced/improved cultivar | A_1 |
| Southern Europe | Advanced/improved cultivar | A_1 |
| Southern Europe | Traditional cultivar/Landrace | A_1 |
| Southern Europe | Traditional cultivar/Landrace | A_1 |
| Southern Europe | Advanced/improved cultivar | A_1 |
| Southern Europe | Traditional cultivar/Landrace | A_1 |
| Southern Europe | Traditional cultivar/Landrace | A_1 |
| Southern Europe | Traditional cultivar/Landrace | A_1 |
| Southern Europe | Traditional cultivar/Landrace | A_1 |
| Southern Europe | Traditional cultivar/Landrace | A_1 |
| Unknown | Advanced/improved cultivar | A_1 |
| Unknown | Advanced/improved cultivar | A_1 |
| Southern Europe | Traditional cultivar/Landrace | A_1 |
| Southern Europe | Traditional cultivar/Landrace | A_2 |
| Southern Europe | Traditional cultivar/Landrace | A_1 |
| Southern Europe | Traditional cultivar/Landrace | A_1 |
| Unknown | Unknown | A_1 |
| Western Europe | Advanced/improved cultivar | A_1 |
| Western Europe | Advanced/improved cultivar | A_1 |
| Eastern Europe | Advanced/improved cultivar | A_1 |
| Western Europe | Advanced/improved cultivar | A_1 |
| Southern Europe | Advanced/improved cultivar | A_1 |
| Unknown | Advanced/improved cultivar | A_1 |
| Western Europe | Advanced/improved cultivar | A_1 |
| Unknown | Advanced/improved cultivar | A_1 |
| Eastern Europe | Advanced/improved cultivar | A_1 |
| Southern Europe | Unknown | A_3 |
| Unknown | Advanced/improved cultivar | A_1 |

| | | |
|---|---|---|
| Unknown | Advanced/improved cultivar | A_1 |
| Eastern Europe | Breeding/Research Material | A_1 |
| Eastern Europe | Advanced/improved cultivar | A_1 |
| Eastern Europe | Advanced/improved cultivar | A_1 |
| Eastern Europe | Advanced/improved cultivar | A_1 |
| Unknown | Advanced/improved cultivar | A_1 |
| Western Europe | Breeding/Research Material | A_1 |
| Western Europe | Advanced/improved cultivar | A_1 |
| Middle East | Unknown | A_1 |
| Unknown | Advanced/improved cultivar | A_1 |
| Unknown | Breeding/Research Material | A_1 |
| Eastern Europe | Advanced/improved cultivar | A_1 |
| Northern Africa | Traditional cultivar/Landrace | A_1 |
| Eastern Europe | Breeding/Research Material | A_1 |
| Eastern Africa | Unknown | A_1 |
| Southern Europe | Unknown | A_1 |
| Northern Africa | Traditional cultivar/Landrace | A_1 |
| Southern Europe | Traditional cultivar/Landrace | A_3 |
| Southern Europe | Traditional cultivar/Landrace | A_1 |
| Southern Europe | Traditional cultivar/Landrace | A_1 |
| Southern Europe | Traditional cultivar/Landrace | A_1 |
| Southern Europe | Traditional cultivar/Landrace | A_1 |
| Southern Europe | Traditional cultivar/Landrace | A_1 |
| Southern Europe | Traditional cultivar/Landrace | A_1 |
| Southern Europe | Traditional cultivar/Landrace | A_1 |
| Southern Europe | Traditional cultivar/Landrace | A_1 |
| Southern Europe | Traditional cultivar/Landrace | A_1 |
| Southern Europe | Traditional cultivar/Landrace | A_1 |
| Southern Europe | Traditional cultivar/Landrace | A_1 |
| Southern Europe | Traditional cultivar/Landrace | A_1 |
| Southern Europe | Traditional cultivar/Landrace | A_1 |
| Southern Europe | Traditional cultivar/Landrace | A_1 |
| Southern Europe | Traditional cultivar/Landrace | A_1 |
| Southern Europe | Traditional cultivar/Landrace | A_1 |
| Southern Europe | Traditional cultivar/Landrace | A_1 |
| Southern Europe | Traditional cultivar/Landrace | A_1 |
| Southern Europe | Traditional cultivar/Landrace | A_1 |
| Southern Europe | Traditional cultivar/Landrace | A_1 |
| Southern Europe | Traditional cultivar/Landrace | A_1 |
| Southern Europe | Traditional cultivar/Landrace | A_1 |
| Southern Europe | Traditional cultivar/Landrace | A_1 |
| Southern Europe | Traditional cultivar/Landrace | A_1 |
| Southern Europe | Traditional cultivar/Landrace | A_1 |
| Southern Europe | Traditional cultivar/Landrace | A_1 |
| Southern Europe | Traditional cultivar/Landrace | A_1 |
| Southern Europe | Traditional cultivar/Landrace | A_1 |
| Southern Europe | Traditional cultivar/Landrace | A_1 |
| Southern Europe | Traditional cultivar/Landrace | A_1 |
| Southern Europe | Traditional cultivar/Landrace | A_1 |
| Southern Europe | Traditional cultivar/Landrace | A_1 |
| Southern Europe | Traditional cultivar/Landrace | A_1 |
| Southern Europe | Traditional cultivar/Landrace | A_1 |

| | | |
|---|---|---|
| Southern Europe | Traditional cultivar/Landrace | A_1 |
| Southern Europe | Traditional cultivar/Landrace | A_1 |
| Southern Europe | Traditional cultivar/Landrace | A_1 |
| Southern Europe | Traditional cultivar/Landrace | A_1 |
| Southern Europe | Traditional cultivar/Landrace | A_1 |
| Southern Europe | Unknown | A_1 |
| Southern Europe | Traditional cultivar/Landrace | B |
| Southern Europe | Traditional cultivar/Landrace | B |
| Southern Europe | Traditional cultivar/Landrace | A_1 |
| Southern Europe | Traditional cultivar/Landrace | A_1 |
| Southern Europe | Traditional cultivar/Landrace | A_1 |
| Southern Europe | Traditional cultivar/Landrace | A_1 |
| Southern Europe | Traditional cultivar/Landrace | A_1 |
| Southern Europe | Traditional cultivar/Landrace | A_1 |
| Southern Europe | Traditional cultivar/Landrace | A_1 |
| Southern Europe | Traditional cultivar/Landrace | A_1 |
| Southern Europe | Traditional cultivar/Landrace | A_1 |
| Southern Europe | Traditional cultivar/Landrace | A_1 |
| Southern Europe | Traditional cultivar/Landrace | A_2 |
| Southern Europe | Traditional cultivar/Landrace | A_1 |
| Southern Europe | Traditional cultivar/Landrace | A_1 |
| Southern Europe | Traditional cultivar/Landrace | A_1 |
| Southern Europe | Traditional cultivar/Landrace | A_1 |
| Southern Europe | Traditional cultivar/Landrace | A_1 |
| Southern Europe | Traditional cultivar/Landrace | A_1 |
| Southern Europe | Traditional cultivar/Landrace | A_1 |
| Unknown | Unknown | A_1 |
| Unknown | Advanced/improved cultivar | A_1 |
| Unknown | Unknown | A_1 |
| Unknown | Unknown | A_1 |
| Unknown | Unknown | A_1 |
| Unknown | Unknown | A_1 |
| Unknown | Unknown | A_1 |
| Unknown | Unknown | A_1 |
| Southern Europe | Unknown | A_3 |
| Unknown | Unknown | A_1 |
| Northern Africa | Unknown | A_1 |
| Southern Europe | Traditional cultivar/Landrace | A_1 |
| Southern Europe | Traditional cultivar/Landrace | A_1 |
| Southern Europe | Unknown | A_1 |
| Southern Europe | Unknown | A_1 |
| Southern Europe | Unknown | A_1 |
| Southern Europe | Unknown | A_1 |
| Southern Europe | Unknown | A_1 |
| Southern Europe | Unknown | A_1 |
| Southern Europe | Unknown | A_1 |
| Southern Europe | Unknown | A_1 |
| Unknown | Breeding/Research Material | A_1 |
| Unknown | Advanced/improved cultivar | A_3 |
| Northern Africa | Unknown | A_1 |
| Unknown | Advanced/improved cultivar | A_1 |
| Unknown | Unknown | A_1 |

| | | |
|---|---|---|
| Southern Europe | Traditional cultivar/Landrace | A_1 |
| Southern Europe | Traditional cultivar/Landrace | A_1 |
| Eastern Europe | Advanced/improved cultivar | A_1 |
| Southern Europe | Traditional cultivar/Landrace | A_1 |
| Southern Europe | Traditional cultivar/Landrace | A_1 |
| Southern Europe | Traditional cultivar/Landrace | A_1 |
| Southern Europe | Traditional cultivar/Landrace | A_1 |
| Southern Europe | Traditional cultivar/Landrace | A_1 |
| Southern Europe | Traditional cultivar/Landrace | A_1 |
| Southern Europe | Traditional cultivar/Landrace | A_1 |
| Southern Europe | Traditional cultivar/Landrace | A_1 |
| Southern Europe | Traditional cultivar/Landrace | A_1 |
| Southern Europe | Traditional cultivar/Landrace | A_1 |
| Southern Europe | Traditional cultivar/Landrace | A_1 |
| Southern Europe | Traditional cultivar/Landrace | A_1 |
| Southern Europe | Traditional cultivar/Landrace | A_1 |
| Southern Europe | Traditional cultivar/Landrace | A_1 |
| Southern Europe | Traditional cultivar/Landrace | A_1 |
| Southern Europe | Traditional cultivar/Landrace | A_1 |
| Southern Europe | Traditional cultivar/Landrace | A_1 |
| Southern Europe | Unknown | A_1 |
| Unknown | Advanced/improved cultivar | A_1 |
| Southern Europe | Unknown | A_1 |
| Southern Europe | Unknown | A_1 |
| Middle East | Wild | A_1 |
| Eastern Africa | Unknown | A_1 |
| Western Europe | Traditional cultivar/Landrace | A_1 |
| Western Europe | Unknown | A_1 |
| Eastern Europe | Advanced/improved cultivar | A_1 |
| Southern Europe | Traditional cultivar/Landrace | A_1 |
| Southern Europe | Traditional cultivar/Landrace | A_1 |
| Southern Europe | Traditional cultivar/Landrace | A_1 |
| Southern Europe | Traditional cultivar/Landrace | A_1 |
| Southern Europe | Traditional cultivar/Landrace | A_1 |
| Eastern Europe | Advanced/improved cultivar | A_1 |
| Unknown | Advanced/improved cultivar | A_1 |
| Unknown | Advanced/improved cultivar | A_1 |
| Southern Europe | Traditional cultivar/Landrace | A_1 |
| Western Europe | Breeding/Research Material | A_1 |
| Southern Europe | Unknown | A_1 |
| Southern Europe | Traditional cultivar/Landrace | A_1 |
| Southern Europe | Traditional cultivar/Landrace | A_1 |
| Southern Europe | Traditional cultivar/Landrace | A_3 |
| Southern Europe | Traditional cultivar/Landrace | A_1 |
| Southern Europe | Traditional cultivar/Landrace | A_1 |
| Southern Europe | Traditional cultivar/Landrace | A_1 |
| Southern Europe | Traditional cultivar/Landrace | A_1 |
| Unknown | Breeding/Research Material | A_1 |
| Unknown | Breeding/Research Material | A_1 |
| Unknown | Breeding/Research Material | A_1 |
| Unknown | Breeding/Research Material | A_1 |
| Unknown | Breeding/Research Material | A_1 |

| | | |
|---|---|---|
| Unknown | Breeding/Research Material | A_1 |
| Eastern Europe | Breeding/Research Material | A_1 |
| Unknown | Breeding/Research Material | A_1 |
| Western Europe | Breeding/Research Material | A_1 |
| Unknown | Breeding/Research Material | A_1 |
| Western Europe | Breeding/Research Material | A_1 |
| Western Europe | Advanced/improved cultivar | A_1 |
| Western Europe | Breeding/Research Material | A_1 |
| Western Europe | Breeding/Research Material | A_1 |
| Western Europe | Advanced/improved cultivar | A_1 |
| Western Europe | Breeding/Research Material | A_1 |
| Southern Europe | Breeding/Research Material | A_1 |
| Western Europe | Breeding/Research Material | A_1 |
| Western Europe | Breeding/Research Material | A_1 |
| Eastern Europe | Advanced/improved cultivar | A_1 |
| Western Europe | Breeding/Research Material | A_1 |
| Eastern Europe | Breeding/Research Material | A_1 |
| Western Europe | Breeding/Research Material | A_1 |
| Western Europe | Breeding/Research Material | A_1 |
| Western Europe | Breeding/Research Material | A_1 |
| Eastern Europe | Advanced/improved cultivar | A_1 |
| Western Europe | Breeding/Research Material | A_1 |
| Unknown | Breeding/Research Material | A_1 |
| Western Europe | Advanced/improved cultivar | A_1 |
| Unknown | Advanced/improved cultivar | A_1 |
| Southern Europe | Breeding/Research Material | A_1 |
| Western Europe | Breeding/Research Material | A_1 |
| Unknown | Advanced/improved cultivar | A_1 |
| Unknown | Breeding/Research Material | A_1 |
| Unknown | Breeding/Research Material | A_1 |
| Unknown | Breeding/Research Material | A_1 |
| Unknown | Breeding/Research Material | A_1 |
| Southern Europe | Advanced/improved cultivar | A_1 |
| Eastern Europe | Advanced/improved cultivar | A_1 |
| Southern Europe | Breeding/Research Material | A_1 |
| Southern Europe | Breeding/Research Material | A_1 |
| Southern Europe | Breeding/Research Material | A_1 |
| Northern Africa | Breeding/Research Material | A_1 |
| Southern Europe | Breeding/Research Material | A_1 |
| Western Europe | Breeding/Research Material | A_1 |
| Unknown | Advanced/improved cultivar | A_1 |
| Southern Europe | Breeding/Research Material | A_1 |
| Western Europe | Advanced/improved cultivar | A_1 |
| Southern Europe | Breeding/Research Material | A_1 |
| Western Europe | Advanced/improved cultivar | A_1 |
| Southern Europe | Unknown | A_1 |
| Southern Europe | Breeding/Research Material | A_1 |
| Unknown | Breeding/Research Material | A_1 |
| Unknown | Breeding/Research Material | A_1 |
| Unknown | Breeding/Research Material | A_1 |
| Unknown | Unknown | A_1 |
| Unknown | Breeding/Research Material | A_1 |

| | | |
|---|---|---|
| Unknown | Advanced/improved cultivar | A_1 |
| Southern Europe | Unknown | A_1 |
| Unknown | Advanced/improved cultivar | A_1 |
| Northern Africa | Wild | A_1 |
| Southern Europe | Breeding/Research Material | A_1 |
| Southern Europe | Breeding/Research Material | A_1 |
| Southern Europe | Breeding/Research Material | A_1 |
| Southern Europe | Breeding/Research Material | A_1 |
| Southern Europe | Breeding/Research Material | A_1 |
| Western Europe | Advanced/improved cultivar | A_1 |
| Northern Africa | Cultivar | A_1 |
| Southern Europe | Breeding/Research Material | A_1 |
| Southern Europe | Breeding/Research Material | A_1 |
| Western Europe | Advanced/improved cultivar | A_1 |
| Southern Europe | Breeding/Research Material | A_1 |
| Unknown | Breeding/Research Material | A_1 |
| Western Europe | Advanced/improved cultivar | A_1 |
| Northern Africa | Breeding/Research Material | A_1 |
| Southern Europe | Wild | A_3 |
| Southern Europe | Advanced/improved cultivar | A_1 |
| Northern Africa | Breeding/Research Material | A_1 |
| Northern Africa | Breeding/Research Material | A_1 |
| Northern Africa | Breeding/Research Material | A_1 |
| Southern Europe | Landrace | A_1 |
| Western Europe | Breeding/Research Material | A_1 |
| Eastern Europe | Breeding/Research Material | A_1 |
| Northern Africa | Unknown | A_1 |
| Southern Europe | Breeding/Research Material | A_1 |
| Western Europe | Advanced/improved cultivar | A_1 |
| Northern Africa | Breeding/Research Material | A_1 |
| Northern Africa | Breeding/Research Material | A_1 |
| Southern Europe | Breeding/Research Material | A_1 |
| Southern Europe | Breeding/Research Material | A_1 |
| Western Europe | Advanced/improved cultivar | A_1 |
| Northern Africa | Breeding/Research Material | A_1 |
| America | Traditional cultivar/Landrace | A_1 |
| Western Europe | Breeding/Research Material | A_1 |
| Northern Africa | Breeding/Research Material | A_1 |
| Northern Africa | Breeding/Research Material | A_1 |
| Western Europe | Advanced/improved cultivar | A_1 |
| Western Europe | Advanced/improved cultivar | A_1 |
| Eastern Europe | Unknown | A_1 |
| Southern Europe | Traditional cultivar/Landrace | A_1 |
| Western Europe | Advanced/improved cultivar | A_1 |
| Southern Europe | Traditional cultivar/Landrace | A_1 |
| Eastern Europe | Advanced/improved cultivar | A_1 |
| Western Europe | Advanced/improved cultivar | A_1 |
| Southern Europe | Traditional cultivar/Landrace | A_1 |
| Southern Europe | Breeding/Research Material | A_1 |
| Southern Europe | Breeding/Research Material | A_1 |
| Southern Europe | Breeding/Research Material | A_1 |
| Southern Europe | Breeding/Research Material | A_1 |

| | | |
|---|---|---|
| Southern Europe | Breeding/Research Material | A_1 |
| Southern Europe | Breeding/Research Material | A_1 |
| Southern Europe | Breeding/Research Material | A_1 |
| Southern Europe | Breeding/Research Material | A_1 |
| Southern Europe | Breeding/Research Material | A_1 |
| Western Europe | Breeding/Research Material | A_1 |
| Western Europe | Advanced/improved cultivar | A_1 |
| Eastern Europe | Advanced/improved cultivar | A_1 |
| Eastern Europe | Traditional cultivar/Landrace | A_1 |
| Eastern Europe | Advanced/improved cultivar | A_1 |
| Eastern Europe | Advanced/improved cultivar | A_1 |
| Eastern Europe | Advanced/improved cultivar | A_1 |
| Eastern Europe | Advanced/improved cultivar | A_1 |
| Eastern Europe | Breeding/Research Material | A_1 |
| Eastern Europe | Advanced/improved cultivar | A_1 |
| Eastern Europe | Breeding/Research Material | A_1 |
| Eastern Europe | Advanced/improved cultivar | A_1 |
| Eastern Europe | Advanced/improved cultivar | A_1 |
| Eastern Europe | Advanced/improved cultivar | A_1 |
| Eastern Europe | Advanced/improved cultivar | A_1 |
| Eastern Europe | Breeding/Research Material | A_1 |
| Eastern Europe | Breeding/Research Material | A_1 |
| Eastern Europe | Advanced/improved cultivar | A_1 |
| Eastern Europe | Advanced/improved cultivar | A_1 |
| Eastern Europe | Advanced/improved cultivar | A_1 |
| Southern Europe | Traditional cultivar/Landrace | A_1 |
| Southern Europe | Traditional cultivar/Landrace | A_1 |
| Southern Europe | Traditional cultivar/Landrace | A_3 |
| Southern Europe | Traditional cultivar/Landrace | A_1 |
| Southern Europe | Traditional cultivar/Landrace | A_1 |
| Southern Europe | Traditional cultivar/Landrace | A_1 |
| Southern Europe | Traditional cultivar/Landrace | A_1 |
| Eastern Europe | Advanced/improved cultivar | A_1 |
| Australia and New Zealand | Advanced/improved cultivar | A_1 |
| Eastern Europe | Advanced/improved cultivar | A_1 |
| Eastern Europe | Advanced/improved cultivar | A_1 |
| Eastern Europe | Advanced/improved cultivar | A_1 |
| Eastern Europe | Advanced/improved cultivar | A_1 |
| Western Europe | Advanced/improved cultivar | A_1 |
| Western Europe | Advanced/improved cultivar | A_1 |
| Western Europe | Advanced/improved cultivar | A_1 |
| Western Europe | Advanced/improved cultivar | A_1 |
| Western Europe | Advanced/improved cultivar | A_1 |
| Eastern Europe | Advanced/improved cultivar | A_1 |
| Eastern Europe | Advanced/improved cultivar | A_1 |
| Eastern Europe | Advanced/improved cultivar | A_1 |
| Western Europe | Advanced/improved cultivar | A_1 |
| Northern Africa | Cultivar | A_1 |
| Southern Europe | Wild | A_1 |
| Unknown | Unknown | A_1 |
| Unknown | Unknown | A_1 |
| Australia and New Zealand | Landrace | A_1 |

| | | |
|---|---|---|
| Southern Europe | Unknown | B |
| Southern Europe | Wild | B |
| Northern Africa | Breeding/Research Material | A_1 |
| Northern Africa | Wild | A_1 |
| Northern Africa | Breeding/Research Material | A_1 |
| Eastern Africa | Wild | A_1 |
| Southern Europe | Wild | A_1 |
| Southern Europe | Wild | A_1 |
| Southern Europe | Wild | A_1 |
| Southern Europe | Wild | A_1 |
| Southern Europe | Wild | A_1 |
| Southern Europe | Wild | A_1 |
| Northern Africa | Traditional cultivar/Landrace | A_1 |
| Eastern Africa | Unknown | A_1 |
| Western Europe | Advanced/improved cultivar | A_1 |
| Southern Europe | Unknown | A_1 |
| Southern Europe | Unknown | A_1 |
| Southern Europe | Unknown | A_1 |
| Eastern Europe | Unknown | A_1 |
| Western Europe | Unknown | A_1 |
| Eastern Europe | Unknown | A_1 |
| Western Europe | Unknown | A_1 |
| Western Europe | Unknown | A_1 |
| Southern Europe | Unknown | A_1 |
| Eastern Europe | Unknown | A_1 |
| Eastern Europe | Unknown | A_1 |
| Eastern Europe | Unknown | A_1 |
| Western Europe | Unknown | A_1 |
| Northern Africa | Unknown | A_1 |
| Eastern Europe | Unknown | A_1 |
| Eastern Africa | Unknown | A_1 |
| Western Europe | Advanced/improved cultivar | A_1 |
| Western Europe | Unknown | A_1 |
| America | Unknown | A_1 |
| Northern Africa | Unknown | A_1 |
| Northern Africa | Unknown | A_1 |
| Southern Europe | Unknown | A_1 |
| Southern Europe | Unknown | A_1 |
| Southern Europe | Unknown | A_1 |
| Southern Europe | Unknown | A_1 |
| Southern Europe | Unknown | A_1 |
| Southern Europe | Unknown | A_1 |
| Southern Europe | Unknown | A_1 |
| Southern Europe | Unknown | A_1 |
| Southern Europe | Unknown | A_1 |
| Southern Europe | Unknown | A_1 |
| Southern Europe | Unknown | A_3 |
| Southern Europe | Unknown | A_1 |
| Southern Europe | Unknown | A_1 |
| Southern Europe | Unknown | A_1 |
| Southern Europe | Unknown | A_1 |
| Southern Europe | Unknown | A_1 |

| | | |
|---|---|---|
| Southern Europe | Unknown | A_1 |
| Southern Europe | Unknown | A_1 |
| Southern Europe | Unknown | A_1 |
| Southern Europe | Unknown | A_1 |
| Southern Europe | Unknown | A_1 |
| Southern Europe | Unknown | A_1 |
| Southern Europe | Unknown | A_1 |
| Southern Europe | Unknown | A_1 |
| Southern Europe | Unknown | A_1 |
| Southern Europe | Unknown | A_1 |
| Southern Europe | Unknown | A_1 |
| Southern Europe | Unknown | A_1 |
| Southern Europe | Unknown | A_1 |
| Southern Europe | Unknown | A_1 |
| Southern Europe | Unknown | A_1 |
| Southern Europe | Unknown | A_1 |
| Southern Europe | Unknown | A_1 |
| Southern Europe | Unknown | A_1 |
| Southern Europe | Unknown | A_1 |
| Southern Europe | Unknown | A_1 |
| Southern Europe | Unknown | A_1 |
| Southern Europe | Unknown | A_1 |
| Southern Europe | Unknown | A_1 |
| Southern Europe | Unknown | A_1 |
| Southern Europe | Unknown | A_1 |
| Southern Europe | Unknown | A_1 |
| Southern Europe | Unknown | A_1 |
| Southern Europe | Unknown | A_1 |
| Southern Europe | Unknown | A_1 |
| Southern Europe | Unknown | A_1 |
| Southern Europe | Unknown | A_1 |
| Southern Europe | Unknown | A_1 |
| Southern Europe | Unknown | A_1 |
| Southern Europe | Unknown | A_1 |
| Southern Europe | Unknown | A_1 |
| Southern Europe | Unknown | A_1 |
| Southern Europe | Unknown | A_1 |
| Southern Europe | Unknown | A_1 |
| Southern Europe | Unknown | A_1 |
| Southern Europe | Unknown | A_1 |
| Southern Europe | Unknown | A_1 |
| Southern Europe | Unknown | A_1 |
| Southern Europe | Unknown | B |
| Southern Europe | Unknown | A_1 |
| Southern Europe | Unknown | B |
| Southern Europe | Unknown | B |
| Southern Europe | Unknown | B |
| Southern Europe | Unknown | A_1 |
| Southern Europe | Unknown | A_1 |
| Southern Europe | Unknown | A_1 |
| Southern Europe | Unknown | A_1 |
| Southern Europe | Unknown | A_1 |

| | | |
|---|---|---|
| Southern Europe | Unknown | A_1 |
| Southern Europe | Unknown | B |
| Southern Europe | Unknown | A_1 |
| Southern Europe | Unknown | A_1 |
| Southern Europe | Unknown | A_1 |
| Southern Europe | Unknown | A_1 |
| Southern Europe | Unknown | A_1 |
| Southern Europe | Unknown | A_1 |
| Southern Europe | Unknown | A_1 |
| Southern Europe | Unknown | A_1 |
| Southern Europe | Unknown | A_1 |
| Southern Europe | Unknown | A_3 |
| Southern Europe | Unknown | A_1 |
| Southern Europe | Unknown | A_1 |
| Southern Europe | Unknown | A_1 |
| Southern Europe | Unknown | A_1 |
| Southern Europe | Unknown | A_1 |
| Eastern Africa | Unknown | A_1 |
| Eastern Africa | Unknown | A_1 |
| Eastern Africa | Unknown | A_1 |
| Eastern Africa | Unknown | A_1 |
| Eastern Africa | Unknown | A_1 |
| Eastern Africa | Unknown | A_1 |
| Southern Europe | Unknown | A_1 |
| Southern Europe | Unknown | A_1 |
| Southern Europe | Unknown | A_1 |
| Eastern Europe | Unknown | A_1 |
| Southern Europe | Unknown | A_1 |
| Middle East | Unknown | A_3 |
| Northern Africa | Unknown | A_1 |
| Southern Europe | Unknown | A_1 |
| Southern Europe | Unknown | A_1 |
| Southern Europe | Unknown | B |
| Middle East | Unknown | A_1 |
| Eastern Africa | Unknown | A_1 |
| Northern Africa | Unknown | A_1 |
| Eastern Africa | Unknown | A_1 |
| Eastern Africa | Unknown | A_1 |
| Northern Africa | Unknown | A_1 |
| Middle East | Unknown | A_1 |
| Northern Africa | Unknown | A_1 |
| Southern Europe | Unknown | B |
| Northern Africa | Unknown | A_1 |
| Northern Africa | Unknown | A_1 |
| Western Europe | Unknown | A_1 |
| Australia and New Zealand | Unknown | A_1 |
| Southern Europe | Unknown | B |
| America | Unknown | A_1 |
| Northern Africa | Unknown | A_1 |
| Middle East | Unknown | A_1 |
| Middle East | Unknown | A_1 |
| Northern Africa | Unknown | A_1 |

| | | |
|---|---|---|
| Western Europe | Unknown | A_1 |
| America | Unknown | A_1 |
| Eastern Europe | Unknown | B |
| Unknown | Unknown | A_1 |
| Southern Europe | Unknown | A_1 |
| Southern Europe | Unknown | A_1 |
| Southern Europe | Unknown | A_1 |
| Western Europe | Landrace | A_1 |
| Western Europe | Breeding/Research Material | A_1 |
| Southern Europe | Unknown | A_1 |
| Southern Europe | Landrace | A_1 |
| Southern Europe | Landrace | A_1 |
| Southern Europe | Landrace | A_1 |
| Southern Europe | Landrace | A_2 |
| Southern Europe | Landrace | A_1 |
| Southern Europe | Landrace | A_1 |
| Southern Europe | Landrace | A_1 |
| Southern Europe | Landrace | A_1 |
| Southern Europe | Landrace | A_1 |
| Southern Europe | Landrace | A_2 |
| Southern Europe | Landrace | A_2 |
| Southern Europe | Landrace | A_1 |
| Southern Europe | Landrace | A_1 |
| Southern Europe | Landrace | A_1 |
| Southern Europe | Landrace | A_1 |
| Southern Europe | Landrace | A_2 |
| Southern Europe | Landrace | A_1 |
| Southern Europe | Wild | B |
| Southern Europe | Wild | B |
| Southern Europe | Wild | A_1 |
| Southern Europe | Landrace | A_1 |
| Southern Europe | Wild | A_1 |
| Southern Europe | Wild | A_1 |
| Southern Europe | Landrace | A_1 |
| Southern Europe | Landrace | A_1 |
| Southern Europe | Landrace | A_2 |
| Southern Europe | Landrace | A_2 |
| Southern Europe | Landrace | A_1 |
| Southern Europe | Landrace | A_1 |
| Southern Europe | Landrace | A_1 |
| Southern Europe | Landrace | A_1 |
| Southern Europe | Landrace | A_1 |
| Southern Europe | Landrace | A_1 |
| Southern Europe | Landrace | A_1 |
| Southern Europe | Landrace | A_1 |
| Southern Europe | Wild | B |
| Southern Europe | Wild | A_2 |
| Southern Europe | Wild | A_2 |
| Southern Europe | Landrace | A_2 |
| Southern Europe | Landrace | A_1 |
| Southern Europe | Wild | A_1 |
| Southern Europe | Wild | A_1 |

| | | |
|---|---|---|
| Southern Europe | Wild | B |
| Southern Europe | Wild | A_1 |
| Southern Europe | Wild | B |
| Southern Europe | Wild | A_1 |
| Western Europe | Unknown | B |
| Western Europe | Unknown | A_1 |
| Western Europe | Unknown | A_1 |
| Eastern Europe | Unknown | A_1 |
| Eastern Europe | Unknown | A_1 |
| Eastern Europe | Unknown | A_1 |
| Unknown | Unknown | A_1 |
| Northern Africa | Unknown | A_2 |
| Northern Africa | Unknown | A_1 |
| Northern Africa | Unknown | A_1 |
| America | Unknown | A_1 |
| America | Unknown | A_1 |
| America | Unknown | A_1 |
| Eastern Europe | Unknown | A_2 |
| Southern Europe | Unknown | A_1 |
| Southern Europe | Unknown | A_1 |
| Eastern Europe | Unknown | A_1 |
| Eastern Europe | Unknown | A_1 |
| Southern Europe | Unknown | A_1 |
| Eastern Europe | Unknown | A_1 |
| Western Europe | Unknown | A_1 |
| Southern Europe | Wild | A_1 |
| Southern Europe | Wild | A_1 |
| Southern Europe | Landrace | A_1 |
| Southern Europe | Landrace | A_1 |
| Southern Europe | Landrace | A_1 |
| Southern Europe | Landrace | A_1 |
| Southern Europe | Landrace | A_1 |
| Southern Europe | Landrace | A_1 |
| Southern Europe | Landrace | A_1 |
| Southern Europe | Wild | A_1 |
| Southern Europe | Landrace | A_1 |
| Southern Europe | Landrace | A_1 |
| Southern Europe | Landrace | A_1 |
| Southern Europe | Landrace | A_1 |
| Southern Europe | Landrace | A_1 |
| Southern Europe | Landrace | A_1 |
| Southern Europe | Landrace | A_1 |
| Southern Europe | Wild | A_1 |
| Southern Europe | Wild | A_1 |
| Southern Europe | Wild | A_1 |
| Southern Europe | Wild | A_1 |
| Southern Europe | Wild | A_1 |
| Southern Europe | Wild | A_1 |
| Southern Europe | Wild | A_1 |
| Southern Europe | Wild | A_1 |
| Southern Europe | Wild | A_1 |
| Southern Europe | Wild | A_1 |

| | | |
|---|---|---|
| Southern Europe | Wild | B |
| Southern Europe | Wild | B |
| Southern Europe | Wild | B |
| Southern Europe | Wild | A_1 |
| Southern Europe | Wild | B |
| Southern Europe | Wild | B |
| Southern Europe | Wild | B |
| Southern Europe | Wild | A_1 |
| Southern Europe | Landrace | A_1 |
| Southern Europe | Wild | A_1 |
| Southern Europe | Wild | A_1 |
| Southern Europe | Wild | B |
| Southern Europe | Wild | B |
| Southern Europe | Wild | A_1 |
| Southern Europe | Wild | A_1 |
| Southern Europe | Wild | A_1 |
| Southern Europe | Wild | B |
| Northern Africa | Landrace | A_1 |
| Northern Africa | Landrace | A_1 |
| Middle East | Landrace | A_2 |
| Southern Europe | Wild | B |
| Eastern Africa | Landrace | A_1 |
| Eastern Africa | Landrace | A_1 |
| Southern Europe | Landrace | A_1 |
| Northern Africa | Landrace | A_1 |
| Western Europe | Cultivar | A_1 |
| Eastern Europe | Cultivar | A_1 |
| Southern Europe | Landrace | A_1 |
| Southern Europe | Landrace | A_1 |
| Southern Europe | Landrace | A_1 |
| Southern Europe | Landrace | A_1 |
| Southern Europe | Landrace | A_1 |
| America | Breeding/Research Material | A_1 |
| Eastern Europe | Cultivar | A_1 |
| Western Europe | Breeding/Research Material | A_1 |
| Western Europe | Breeding/Research Material | A_1 |
| Western Europe | Breeding/Research Material | A_1 |
| Eastern Europe | Cultivar | A_1 |
| Eastern Europe | Cultivar | A_2 |
| Eastern Europe | Unknown | A_1 |
| Eastern Europe | Unknown | A_2 |
| Eastern Europe | Unknown | A_1 |
| America | Breeding/Research Material | A_1 |
| America | Breeding/Research Material | A_1 |
| America | Breeding/Research Material | A_1 |
| America | Breeding/Research Material | A_1 |
| America | Breeding/Research Material | A_1 |
| Southern Europe | Landrace | A_1 |
| Southern Europe | Landrace | A_1 |
| Northern Africa | Landrace | A_1 |
| Northern Africa | Landrace | A_1 |
| Northern Africa | Landrace | A_1 |

| | | |
|---|---|---|
| Northern Africa | Landrace | A_1 |
| Northern Africa | Landrace | A_2 |
| Northern Africa | Landrace | A_1 |
| Northern Africa | Landrace | A_1 |
| Northern Africa | Landrace | A_1 |
| Northern Africa | Landrace | A_1 |
| Northern Africa | Landrace | A_1 |
| Northern Africa | Landrace | A_1 |
| Northern Africa | Landrace | A_1 |
| Northern Africa | Landrace | A_1 |
| Northern Africa | Landrace | A_1 |
| Northern Africa | Landrace | A_1 |
| Northern Africa | Landrace | A_1 |
| Northern Africa | Landrace | A_1 |
| Northern Africa | Landrace | A_1 |
| Northern Africa | Landrace | A_1 |
| Northern Africa | Landrace | A_2 |
| Northern Africa | Landrace | A_1 |
| Northern Africa | Landrace | A_2 |
| Northern Africa | Landrace | A_1 |
| Northern Africa | Landrace | A_1 |
| Northern Africa | Landrace | A_1 |
| Northern Africa | Landrace | A_1 |
| Northern Africa | Landrace | A_1 |
| Northern Africa | Landrace | A_1 |
| Northern Africa | Landrace | A_1 |
| Northern Africa | Landrace | A_1 |
| Northern Africa | Landrace | A_1 |
| Northern Africa | Landrace | A_1 |
| Northern Africa | Landrace | A_1 |
| Northern Africa | Landrace | A_1 |
| Northern Africa | Landrace | A_1 |
| Northern Africa | Landrace | A_2 |
| Northern Africa | Landrace | A_1 |
| Northern Africa | Landrace | A_1 |
| Northern Africa | Landrace | A_1 |
| Northern Africa | Landrace | A_1 |
| Northern Africa | Landrace | A_1 |
| Northern Africa | Landrace | A_1 |
| Northern Africa | Landrace | A_1 |
| Northern Africa | Landrace | A_1 |
| Northern Africa | Landrace | A_1 |
| Northern Africa | Landrace | A_1 |
| Northern Africa | Landrace | A_1 |
| Northern Africa | Landrace | A_1 |
| Northern Africa | Landrace | A_1 |
| Northern Africa | Landrace | A_1 |
| Northern Africa | Landrace | A_1 |
| Northern Africa | Landrace | A_1 |
| Northern Africa | Landrace | A_1 |
| Western Europe | Unknown | A_1 |
| Western Europe | Unknown | A_2 |

| | | |
|---|---|---|
| Western Europe | Breeding/Research Material | A_1 |
| Northern Africa | Landrace | A_1 |
| Northern Africa | Landrace | A_1 |
| Northern Africa | Landrace | A_1 |
| Northern Africa | Landrace | A_1 |
| Northern Africa | Landrace | A_1 |
| Northern Africa | Landrace | A_1 |
| Southern Europe | Unknown | A_1 |
| Eastern Africa | Unknown | A_1 |
| Northern Africa | Landrace | A_1 |
| Unknown | Breeding/Research Material | A_1 |
| Western Europe | Landrace | A_2 |
| Western Europe | Landrace | A_1 |
| Western Europe | Landrace | A_1 |
| Western Europe | Landrace | A_1 |
| Western Europe | Landrace | A_1 |
| Western Europe | Landrace | A_1 |
| Western Europe | Landrace | A_1 |
| Western Europe | Landrace | A_1 |
| Western Europe | Landrace | A_1 |
| Western Europe | Landrace | A_2 |
| Western Europe | Landrace | A_1 |
| Western Europe | Landrace | A_1 |
| Western Europe | Landrace | A_1 |
| Western Europe | Landrace | A_1 |
| Western Europe | Breeding/Research Material | A_1 |
| Western Europe | Breeding/Research Material | A_2 |
| Western Europe | Breeding/Research Material | A_2 |
| Western Europe | Breeding/Research Material | A_1 |
| Northern Africa | Landrace | A_1 |
| Northern Africa | Landrace | A_1 |
| Northern Africa | Landrace | A_1 |
| Northern Africa | Landrace | A_1 |
| Northern Africa | Landrace | A_1 |
| Northern Africa | Landrace | A_1 |
| Northern Africa | Landrace | A_1 |
| Northern Africa | Landrace | A_1 |
| Northern Africa | Landrace | A_1 |
| Northern Africa | Landrace | A_1 |
| Northern Africa | Landrace | A_1 |
| Northern Africa | Landrace | A_1 |
| Northern Africa | Landrace | A_1 |
| Northern Africa | Landrace | A_1 |
| Northern Africa | Landrace | A_1 |
| Northern Africa | Landrace | A_1 |
| Northern Africa | Landrace | A_1 |
| Eastern Europe | Unknown | B |
| Middle East | Wild | A_1 |
| Western Europe | Breeding/Research Material | A_1 |
| Western Europe | Breeding/Research Material | A_1 |
| Western Europe | Breeding/Research Material | A_1 |
| Western Europe | Breeding/Research Material | A_1 |

| | | |
|---|---|---|
| Unknown | Cultivar | A_1 |
| Unknown | Unknown | A_2 |
| Unknown | Unknown | B |
| Unknown | Unknown | B |
| Unknown | Unknown | A_1 |
| Unknown | Unknown | B |
| Western Europe | Unknown | A_1 |
| Western Europe | Unknown | A_1 |
| Western Europe | Unknown | A_1 |
| Western Europe | Unknown | A_1 |
| Western Europe | Unknown | A_1 |
| Southern Europe | Wild | B |
| Southern Europe | Landrace | A_1 |
| Southern Europe | Landrace | A_1 |
| Southern Europe | Landrace | A_1 |
| Southern Europe | Landrace | A_1 |
| Southern Europe | Landrace | A_1 |
| Western Europe | Breeding/Research Material | A_1 |
| Northern Africa | Landrace | A_1 |
| Northern Africa | Landrace | A_1 |
| Northern Africa | Landrace | A_1 |
| Southern Europe | Wild | A_1 |
| Western Europe | Breeding/Research Material | A_1 |
| Western Europe | Breeding/Research Material | A_1 |
| Northern Africa | Unknown | A_1 |
| Southern Europe | Unknown | A_1 |
| Southern Europe | Unknown | A_1 |
| Western Europe | Unknown | A_1 |
| Western Europe | Unknown | A_1 |
| Unknown | Cultivar | A_1 |
| Unknown | Landrace | A_1 |
| Australia and New Zealand | Breeding/Research Material | A_1 |
| Southern Europe | Landrace | A_2 |
| Southern Europe | Landrace | A_1 |
| Southern Europe | Landrace | A_1 |
| Southern Europe | Landrace | A_1 |
| Southern Europe | Landrace | A_1 |
| Southern Europe | Landrace | A_1 |
| Southern Europe | Landrace | A_1 |
| Southern Europe | Landrace | A_1 |
| Southern Europe | Landrace | A_2 |
| Southern Europe | Landrace | A_1 |
| Southern Europe | Landrace | A_1 |
| Southern Europe | Landrace | A_1 |
| Southern Europe | Landrace | A_1 |
| Southern Europe | Landrace | A_1 |
| Southern Europe | Landrace | A_1 |
| Southern Europe | Landrace | A_1 |
| Southern Europe | Landrace | A_1 |
| Southern Europe | Landrace | A_2 |
| Southern Europe | Landrace | A_2 |
| Southern Europe | Landrace | A_2 |

| | | |
|---|---|---|
| Southern Europe | Landrace | A_1 |
| Southern Europe | Landrace | A_1 |
| Southern Europe | Landrace | A_1 |
| Southern Europe | Landrace | A_1 |
| Southern Europe | Landrace | A_1 |
| Southern Europe | Landrace | A_1 |
| Southern Europe | Landrace | A_1 |
| Southern Europe | Landrace | A_1 |
| Southern Europe | Landrace | A_1 |
| Southern Europe | Landrace | A_2 |
| Southern Europe | Landrace | A_1 |
| Southern Europe | Landrace | A_1 |
| Southern Europe | Landrace | A_1 |
| Southern Europe | Landrace | A_1 |
| Southern Europe | Landrace | A_1 |
| Southern Europe | Landrace | A_1 |
| Southern Europe | Landrace | A_1 |
| Southern Europe | Landrace | A_1 |
| Southern Europe | Landrace | A_1 |
| Southern Europe | Landrace | A_1 |
| Southern Europe | Landrace | A_1 |
| Southern Europe | Landrace | A_1 |
| Southern Europe | Landrace | A_1 |
| Southern Europe | Landrace | A_1 |
| Southern Europe | Landrace | A_1 |
| Southern Europe | Landrace | A_1 |
| Southern Europe | Landrace | A_1 |
| Southern Europe | Landrace | A_1 |
| Southern Europe | Landrace | A_1 |
| Southern Europe | Landrace | A_1 |
| Southern Europe | Landrace | A_1 |
| Southern Europe | Landrace | A_1 |
| Southern Europe | Landrace | A_1 |
| Southern Europe | Landrace | A_1 |
| Southern Europe | Landrace | A_1 |
| Southern Europe | Landrace | A_1 |
| Southern Europe | Landrace | A_1 |
| Southern Europe | Breeding/Research Material | A_2 |
| Southern Europe | Landrace | A_1 |
| Southern Europe | Landrace | A_2 |
| Eastern Africa | Unknown | A_2 |
| Northern Africa | Landrace | A_1 |
| Northern Africa | Landrace | A_1 |
| Northern Africa | Landrace | A_1 |
| Northern Africa | Landrace | A_1 |
| Northern Africa | Landrace | A_1 |
| Northern Africa | Landrace | A_1 |
| Northern Africa | Landrace | A_1 |
| Northern Africa | Landrace | A_1 |
| Northern Africa | Landrace | A_1 |
| Northern Africa | Landrace | A_1 |
| Northern Africa | Landrace | A_1 |

| | | |
|---|---|---|
| Northern Africa | Landrace | A_1 |
| Northern Africa | Landrace | A_1 |
| Northern Africa | Landrace | A_1 |
| Northern Africa | Landrace | A_1 |
| Northern Africa | Landrace | A_1 |
| Northern Africa | Landrace | A_1 |
| Western Europe | Landrace | A_1 |
| Western Europe | Landrace | A_1 |
| Western Europe | Landrace | A_1 |
| Western Europe | Landrace | A_1 |
| Western Europe | Landrace | A_1 |
| Western Europe | Landrace | A_2 |
| Northern Africa | Landrace | A_1 |
| Northern Africa | Landrace | A_1 |
| Northern Africa | Landrace | A_1 |
| Northern Africa | Landrace | A_1 |
| Northern Africa | Landrace | A_3 |
| Northern Africa | Landrace | A_1 |
| Northern Africa | Landrace | A_2 |
| Northern Africa | Landrace | A_1 |
| Northern Africa | Landrace | A_1 |
| Northern Africa | Landrace | A_1 |
| Northern Africa | Landrace | A_1 |
| Northern Africa | Landrace | A_1 |
| Northern Africa | Landrace | A_1 |
| Northern Africa | Landrace | A_1 |
| Northern Africa | Landrace | A_1 |
| Northern Africa | Landrace | A_1 |
| Northern Africa | Landrace | A_1 |
| Northern Africa | Landrace | A_1 |
| Northern Africa | Landrace | A_1 |
| Northern Africa | Landrace | A_1 |
| Northern Africa | Landrace | A_1 |
| Eastern Africa | Landrace | A_1 |
| Western Europe | Breeding/Research Material | A_1 |
| Northern Africa | Landrace | A_1 |
| Northern Africa | Landrace | A_1 |
| Northern Africa | Landrace | A_1 |
| Northern Africa | Landrace | A_1 |
| Northern Africa | Landrace | A_1 |
| Northern Africa | Landrace | A_1 |
| Northern Africa | Landrace | A_1 |
| Northern Africa | Landrace | A_1 |
| Northern Africa | Landrace | A_1 |
| Eastern Africa | Landrace | A_1 |
| Western Europe | Landrace | A_1 |
| Southern Europe | Landrace | A_1 |
| Southern Europe | Landrace | A_1 |
| Southern Europe | Landrace | A_1 |
| Southern Europe | Landrace | A_1 |
| Southern Europe | Landrace | A_1 |
| Eastern Africa | Landrace | A_1 |

| | | |
|---|---|---|
| Eastern Africa | Landrace | A_1 |
| Eastern Africa | Landrace | A_1 |
| Eastern Africa | Landrace | A_1 |
| Eastern Africa | Landrace | A_1 |
| Eastern Europe | Unknown | A_1 |
| Unknown | Unknown | A_1 |
| Unknown | Unknown | A_1 |
| Unknown | Unknown | A_1 |
| Unknown | Unknown | A_1 |
| Unknown | Unknown | A_1 |
| Unknown | Unknown | A_1 |
| Unknown | Unknown | A_1 |
| Unknown | Unknown | A_2 |
| Unknown | Unknown | A_1 |
| Unknown | Unknown | A_1 |
| Unknown | Unknown | A_1 |
| Unknown | Unknown | A_1 |
| Unknown | Unknown | A_2 |
| Unknown | Unknown | A_1 |
| Unknown | Unknown | A_1 |
| Unknown | Unknown | A_1 |
| Unknown | Unknown | A_1 |
| Unknown | Unknown | A_1 |
| Unknown | Unknown | A_1 |
| Unknown | Unknown | A_1 |
| Unknown | Unknown | A_1 |
| Unknown | Unknown | A_1 |
| Unknown | Unknown | A_1 |
| Unknown | Unknown | A_1 |
| Unknown | Unknown | A_1 |
| Unknown | Unknown | A_2 |
| Unknown | Unknown | A_1 |
| Unknown | Unknown | A_1 |
| Unknown | Unknown | A_1 |
| Unknown | Unknown | A_2 |
| Unknown | Unknown | A_1 |
| Unknown | Unknown | A_1 |
| Unknown | Unknown | A_1 |
| Unknown | Unknown | A_1 |
| Unknown | Unknown | A_1 |
| Unknown | Unknown | A_1 |
| Unknown | Unknown | A_1 |
| Unknown | Unknown | A_1 |
| Australia and New Zealand | Unknown | A_2 |
| Eastern Europe | Traditional cultivar/Landrace | A_1 |
| Eastern Europe | Traditional cultivar/Landrace | A_2 |
| Western Europe | Advanced/improved cultivar | A_1 |
| Western Europe | Advanced/improved cultivar | A_1 |
| Eastern Europe | Breeding/Research Material | A_1 |
| Eastern Europe | Advanced/improved cultivar | A_1 |
| Western Europe | Advanced/improved cultivar | A_1 |
| Western Europe | Advanced/improved cultivar | A_1 |

| | | |
|---|---|---|
| Western Europe | Advanced/improved cultivar | A_1 |
| Eastern Europe | Advanced/improved cultivar | A_1 |
| Eastern Europe | Advanced/improved cultivar | A_1 |
| Eastern Europe | Breeding/Research Material | A_1 |
| Eastern Europe | Advanced/improved cultivar | A_1 |
| Western Europe | Advanced/improved cultivar | A_1 |
| Eastern Europe | Advanced/improved cultivar | A_1 |
| Western Europe | Advanced/improved cultivar | A_1 |
| Western Europe | Advanced/improved cultivar | A_1 |
| Western Europe | Advanced/improved cultivar | A_1 |
| Western Europe | Advanced/improved cultivar | A_1 |
| Eastern Europe | Breeding/Research Material | A_1 |
| Eastern Europe | Advanced/improved cultivar | A_1 |
| Western Europe | Advanced/improved cultivar | A_1 |
| Eastern Europe | Advanced/improved cultivar | A_1 |
| Unknown | Unknown | A_1 |
| Southern Europe | Wild | A_1 |
| Unknown | Unknown | A_1 |
| Eastern Europe | Traditional cultivar/Landrace | A_1 |
| Eastern Europe | Traditional cultivar/Landrace | A_1 |
| Eastern Europe | Traditional cultivar/Landrace | A_1 |
| Eastern Europe | Traditional cultivar/Landrace | A_1 |
| Eastern Europe | Traditional cultivar/Landrace | A_1 |
| Eastern Europe | Traditional cultivar/Landrace | A_1 |
| Eastern Europe | Breeding/Research Material | A_1 |
| Eastern Europe | Breeding/Research Material | A_1 |
| Southern Europe | Traditional cultivar/Landrace | A_1 |
| Southern Europe | Traditional cultivar/Landrace | A_1 |
| Southern Europe | Traditional cultivar/Landrace | A_1 |
| Southern Europe | Traditional cultivar/Landrace | A_1 |
| Southern Europe | Traditional cultivar/Landrace | A_1 |
| Southern Europe | Traditional cultivar/Landrace | A_1 |
| Southern Europe | Traditional cultivar/Landrace | A_1 |
| Southern Europe | Traditional cultivar/Landrace | A_1 |
| Southern Europe | Traditional cultivar/Landrace | A_1 |
| Southern Europe | Traditional cultivar/Landrace | A_1 |
| Southern Europe | Traditional cultivar/Landrace | A_1 |
| Southern Europe | Traditional cultivar/Landrace | A_1 |
| Southern Europe | Traditional cultivar/Landrace | A_2 |
| Southern Europe | Traditional cultivar/Landrace | A_1 |
| Southern Europe | Traditional cultivar/Landrace | A_1 |
| Southern Europe | Traditional cultivar/Landrace | A_1 |
| Southern Europe | Traditional cultivar/Landrace | A_1 |
| Eastern Europe | Advanced/improved cultivar | A_1 |
| Eastern Europe | Advanced/improved cultivar | A_1 |
| Eastern Europe | Advanced/improved cultivar | A_1 |
| Western Europe | Advanced/improved cultivar | A_1 |
| Western Europe | Advanced/improved cultivar | A_1 |
| Western Europe | Advanced/improved cultivar | A_1 |
| Western Europe | Advanced/improved cultivar | A_1 |
| Eastern Europe | Advanced/improved cultivar | A_1 |
| Eastern Europe | Advanced/improved cultivar | A_1 |

| | | |
|---|---|---|
| Eastern Europe | Advanced/improved cultivar | A_1 |
| Eastern Europe | Advanced/improved cultivar | A_1 |
| Western Europe | Advanced/improved cultivar | A_1 |
| Southern Europe | Advanced/improved cultivar | A_1 |
| Southern Europe | Advanced/improved cultivar | A_1 |
| Northern Africa | Advanced/improved cultivar | A_1 |
| Western Europe | Advanced/improved cultivar | A_1 |
| Eastern Europe | Advanced/improved cultivar | A_2 |
| Western Europe | Advanced/improved cultivar | A_1 |
| Western Europe | Advanced/improved cultivar | A_1 |
| Eastern Europe | Advanced/improved cultivar | A_1 |
| Northern Africa | Traditional cultivar/Landrace | A_1 |
| Southern Europe | Traditional cultivar/Landrace | A_1 |
| America | Advanced/improved cultivar | A_1 |
| America | Advanced/improved cultivar | A_1 |
| America | Breeding/Research Material | A_1 |
| Eastern Europe | Advanced/improved cultivar | A_1 |
| Western Europe | Advanced/improved cultivar | A_1 |
| Unknown | Advanced/improved cultivar | A_1 |
| America | Advanced/improved cultivar | A_1 |
| America | Breeding/Research Material | A_1 |
| Southern Europe | Traditional cultivar/Landrace | A_1 |
| Southern Europe | Traditional cultivar/Landrace | A_1 |
| Unknown | Traditional cultivar/Landrace | A_1 |
| Southern Europe | Traditional cultivar/Landrace | A_1 |
| Southern Europe | Traditional cultivar/Landrace | A_1 |
| America | Advanced/improved cultivar | A_1 |
| America | Advanced/improved cultivar | A_1 |
| America | Advanced/improved cultivar | A_1 |
| Eastern Europe | Advanced/improved cultivar | A_1 |
| Eastern Europe | Breeding/Research Material | A_2 |
| Eastern Europe | Breeding/Research Material | A_1 |
| Eastern Europe | Breeding/Research Material | A_1 |
| Western Europe | Advanced/improved cultivar | A_1 |
| Eastern Europe | Advanced/improved cultivar | A_1 |
| Eastern Europe | Advanced/improved cultivar | A_2 |
| Southern Europe | Traditional cultivar/Landrace | A_1 |
| Southern Europe | Traditional cultivar/Landrace | A_1 |
| Southern Europe | Traditional cultivar/Landrace | A_1 |
| Southern Europe | Traditional cultivar/Landrace | A_1 |
| Southern Europe | Traditional cultivar/Landrace | A_1 |
| Southern Europe | Traditional cultivar/Landrace | A_1 |
| Southern Europe | Traditional cultivar/Landrace | A_2 |
| Southern Europe | Traditional cultivar/Landrace | A_1 |
| Unknown | Traditional cultivar/Landrace | A_1 |
| Southern Europe | Traditional cultivar/Landrace | A_1 |
| Southern Europe | Traditional cultivar/Landrace | A_1 |
| Southern Europe | Traditional cultivar/Landrace | A_1 |
| Southern Europe | Traditional cultivar/Landrace | A_1 |
| Southern Europe | Traditional cultivar/Landrace | A_1 |
| Southern Europe | Traditional cultivar/Landrace | A_1 |
| Southern Europe | Traditional cultivar/Landrace | A_1 |

| | | |
|---|---|---|
| Southern Europe | Traditional cultivar/Landrace | A_2 |
| Southern Europe | Traditional cultivar/Landrace | A_1 |
| Southern Europe | Traditional cultivar/Landrace | A_1 |
| Southern Europe | Traditional cultivar/Landrace | A_1 |
| Southern Europe | Traditional cultivar/Landrace | A_1 |
| Southern Europe | Traditional cultivar/Landrace | A_1 |
| Southern Europe | Traditional cultivar/Landrace | A_1 |
| Southern Europe | Traditional cultivar/Landrace | A_1 |
| Southern Europe | Traditional cultivar/Landrace | A_1 |
| Southern Europe | Traditional cultivar/Landrace | A_1 |
| Southern Europe | Traditional cultivar/Landrace | A_1 |
| Southern Europe | Traditional cultivar/Landrace | A_2 |
| Southern Europe | Traditional cultivar/Landrace | A_1 |
| Southern Europe | Traditional cultivar/Landrace | A_1 |
| Southern Europe | Traditional cultivar/Landrace | A_1 |
| Southern Europe | Traditional cultivar/Landrace | A_2 |
| Southern Europe | Traditional cultivar/Landrace | A_1 |
| Southern Europe | Traditional cultivar/Landrace | A_1 |
| Southern Europe | Traditional cultivar/Landrace | A_1 |
| America | Traditional cultivar/Landrace | A_1 |
| Northern Africa | Traditional cultivar/Landrace | A_1 |
| Eastern Europe | Traditional cultivar/Landrace | A_1 |
| Southern Asia | Traditional cultivar/Landrace | A_1 |
| Southern Europe | Traditional cultivar/Landrace | A_1 |
| Southern Europe | Traditional cultivar/Landrace | A_1 |
| Southern Europe | Traditional cultivar/Landrace | A_1 |
| Southern Europe | Traditional cultivar/Landrace | A_1 |
| Southern Europe | Traditional cultivar/Landrace | A_1 |
| Southern Europe | Traditional cultivar/Landrace | A_1 |
| America | Traditional cultivar/Landrace | A_1 |
| Southern Europe | Traditional cultivar/Landrace | A_1 |
| Southern Europe | Traditional cultivar/Landrace | A_1 |
| Southern Europe | Traditional cultivar/Landrace | A_1 |
| Southern Europe | Traditional cultivar/Landrace | A_2 |
| Southern Europe | Traditional cultivar/Landrace | A_1 |
| Southern Europe | Traditional cultivar/Landrace | A_1 |
| Southern Europe | Traditional cultivar/Landrace | A_1 |
| Southern Europe | Traditional cultivar/Landrace | A_1 |
| Southern Europe | Traditional cultivar/Landrace | A_1 |
| Southern Europe | Traditional cultivar/Landrace | A_2 |
| Southern Europe | Traditional cultivar/Landrace | A_1 |
| Southern Europe | Traditional cultivar/Landrace | A_1 |
| Southern Europe | Traditional cultivar/Landrace | A_1 |
| Southern Europe | Traditional cultivar/Landrace | A_1 |
| Southern Europe | Traditional cultivar/Landrace | A_1 |
| Southern Europe | Traditional cultivar/Landrace | A_1 |
| Southern Europe | Traditional cultivar/Landrace | A_1 |
| Southern Europe | Traditional cultivar/Landrace | A_3 |
| Southern Europe | Traditional cultivar/Landrace | A_1 |
| Southern Europe | Traditional cultivar/Landrace | A_1 |
| Southern Europe | Traditional cultivar/Landrace | A_1 |
| Southern Europe | Traditional cultivar/Landrace | A_1 |

| | | |
|---|---|---|
| Southern Europe | Traditional cultivar/Landrace | A_1 |
| Southern Europe | Traditional cultivar/Landrace | A_2 |
| Southern Europe | Traditional cultivar/Landrace | A_1 |
| Southern Europe | Traditional cultivar/Landrace | A_1 |
| Southern Europe | Traditional cultivar/Landrace | A_1 |
| Southern Europe | Traditional cultivar/Landrace | A_1 |
| Southern Europe | Traditional cultivar/Landrace | A_1 |
| Southern Europe | Traditional cultivar/Landrace | A_2 |
| Southern Europe | Traditional cultivar/Landrace | A_1 |
| Southern Europe | Traditional cultivar/Landrace | A_2 |
| Southern Europe | Traditional cultivar/Landrace | A_1 |
| Southern Europe | Traditional cultivar/Landrace | A_1 |
| Southern Europe | Traditional cultivar/Landrace | A_1 |
| Southern Europe | Traditional cultivar/Landrace | A_1 |
| Southern Europe | Traditional cultivar/Landrace | A_1 |
| Southern Europe | Traditional cultivar/Landrace | A_1 |
| Southern Europe | Traditional cultivar/Landrace | A_2 |
| Southern Europe | Traditional cultivar/Landrace | A_1 |
| Southern Europe | Traditional cultivar/Landrace | A_1 |
| Southern Europe | Traditional cultivar/Landrace | A_1 |
| Southern Europe | Traditional cultivar/Landrace | A_2 |
| Southern Europe | Traditional cultivar/Landrace | A_2 |
| Southern Europe | Traditional cultivar/Landrace | A_1 |
| Southern Europe | Traditional cultivar/Landrace | A_1 |
| Southern Europe | Traditional cultivar/Landrace | A_1 |
| Southern Europe | Traditional cultivar/Landrace | A_1 |
| Southern Europe | Traditional cultivar/Landrace | A_1 |
| Southern Europe | Traditional cultivar/Landrace | A_1 |
| Southern Europe | Traditional cultivar/Landrace | A_2 |
| Southern Europe | Traditional cultivar/Landrace | A_2 |
| Southern Europe | Wild | A_2 |
| Southern Europe | Traditional cultivar/Landrace | A_1 |
| Southern Europe | Traditional cultivar/Landrace | A_2 |
| Southern Europe | Traditional cultivar/Landrace | A_1 |
| Southern Europe | Traditional cultivar/Landrace | A_1 |
| Southern Europe | Traditional cultivar/Landrace | A_1 |
| Southern Europe | Traditional cultivar/Landrace | A_1 |
| Southern Europe | Traditional cultivar/Landrace | A_1 |
| Southern Europe | Weedy | A_1 |
| Southern Europe | Traditional cultivar/Landrace | A_1 |
| Southern Europe | Traditional cultivar/Landrace | A_1 |
| Southern Europe | Traditional cultivar/Landrace | A_1 |
| Southern Europe | Traditional cultivar/Landrace | A_1 |
| Southern Europe | Traditional cultivar/Landrace | A_2 |
| Eastern Europe | Breeding/Research Material | A_1 |
| Southern Europe | Traditional cultivar/Landrace | A_1 |
| Southern Europe | Traditional cultivar/Landrace | A_1 |
| Southern Europe | Traditional cultivar/Landrace | A_2 |
| Southern Europe | Traditional cultivar/Landrace | A_1 |
| Southern Europe | Traditional cultivar/Landrace | A_1 |
| Southern Europe | Wild | A_1 |
| Southern Europe | Traditional cultivar/Landrace | A_1 |

| | | |
|---|---|---|
| Southern Europe | Traditional cultivar/Landrace | A_1 |
| Southern Europe | Traditional cultivar/Landrace | A_2 |
| Southern Europe | Traditional cultivar/Landrace | A_1 |
| Southern Europe | Traditional cultivar/Landrace | A_1 |
| Southern Europe | Traditional cultivar/Landrace | A_1 |
| Southern Europe | Traditional cultivar/Landrace | A_2 |
| Southern Europe | Traditional cultivar/Landrace | A_1 |
| Southern Europe | Traditional cultivar/Landrace | A_1 |
| Southern Europe | Traditional cultivar/Landrace | A_1 |
| Southern Europe | Traditional cultivar/Landrace | A_1 |
| Southern Europe | Traditional cultivar/Landrace | A_1 |
| Southern Europe | Traditional cultivar/Landrace | A_1 |
| Southern Europe | Traditional cultivar/Landrace | A_1 |
| Southern Europe | Traditional cultivar/Landrace | A_1 |
| Southern Europe | Traditional cultivar/Landrace | A_1 |
| Southern Europe | Traditional cultivar/Landrace | A_1 |
| Southern Europe | Traditional cultivar/Landrace | A_1 |
| Southern Europe | Traditional cultivar/Landrace | A_1 |
| Southern Europe | Traditional cultivar/Landrace | A_2 |
| Southern Europe | Traditional cultivar/Landrace | A_1 |
| Southern Europe | Traditional cultivar/Landrace | A_1 |
| Southern Europe | Traditional cultivar/Landrace | A_2 |
| Southern Europe | Traditional cultivar/Landrace | A_1 |
| Southern Europe | Traditional cultivar/Landrace | A_1 |
| Southern Europe | Traditional cultivar/Landrace | A_1 |
| Southern Europe | Traditional cultivar/Landrace | A_1 |
| Southern Europe | Traditional cultivar/Landrace | A_1 |
| Southern Europe | Traditional cultivar/Landrace | A_1 |
| Southern Europe | Traditional cultivar/Landrace | A_1 |
| Southern Europe | Traditional cultivar/Landrace | A_1 |
| Southern Europe | Traditional cultivar/Landrace | A_1 |
| Southern Europe | Traditional cultivar/Landrace | A_1 |
| Southern Europe | Traditional cultivar/Landrace | A_1 |
| Southern Europe | Traditional cultivar/Landrace | A_1 |
| Southern Europe | Traditional cultivar/Landrace | A_1 |
| Southern Europe | Traditional cultivar/Landrace | A_1 |
| Southern Europe | Traditional cultivar/Landrace | A_1 |
| Southern Europe | Traditional cultivar/Landrace | A_1 |
| Southern Europe | Traditional cultivar/Landrace | A_1 |
| Southern Europe | Traditional cultivar/Landrace | A_1 |
| Southern Europe | Traditional cultivar/Landrace | A_1 |
| Southern Europe | Traditional cultivar/Landrace | A_1 |
| Southern Europe | Traditional cultivar/Landrace | A_1 |
| Southern Europe | Traditional cultivar/Landrace | A_1 |
| Southern Europe | Traditional cultivar/Landrace | A_2 |
| Southern Europe | Traditional cultivar/Landrace | A_1 |
| Southern Europe | Traditional cultivar/Landrace | A_1 |
| Southern Europe | Traditional cultivar/Landrace | A_1 |
| Southern Europe | Traditional cultivar/Landrace | A_1 |
| Southern Europe | Traditional cultivar/Landrace | A_1 |
| Southern Europe | Traditional cultivar/Landrace | A_2 |
| Southern Europe | Traditional cultivar/Landrace | A_1 |

| | | |
|---|---|---|
| Southern Europe | Traditional cultivar/Landrace | A_2 |
| Southern Europe | Traditional cultivar/Landrace | A_1 |
| Southern Europe | Traditional cultivar/Landrace | A_1 |
| Southern Europe | Traditional cultivar/Landrace | A_1 |
| Southern Europe | Traditional cultivar/Landrace | A_1 |
| Southern Europe | Traditional cultivar/Landrace | A_2 |
| Southern Europe | Traditional cultivar/Landrace | A_1 |
| Southern Europe | Traditional cultivar/Landrace | A_1 |
| Southern Europe | Traditional cultivar/Landrace | A_2 |
| Southern Europe | Traditional cultivar/Landrace | A_1 |
| Southern Europe | Traditional cultivar/Landrace | A_2 |
| Southern Europe | Traditional cultivar/Landrace | A_1 |
| Southern Europe | Traditional cultivar/Landrace | A_1 |
| Southern Europe | Traditional cultivar/Landrace | A_2 |
| Southern Europe | Traditional cultivar/Landrace | A_1 |
| Southern Europe | Traditional cultivar/Landrace | A_1 |
| Southern Europe | Traditional cultivar/Landrace | A_2 |
| Southern Europe | Traditional cultivar/Landrace | A_1 |
| Southern Europe | Traditional cultivar/Landrace | A_1 |
| Southern Europe | Traditional cultivar/Landrace | A_2 |
| Southern Europe | Traditional cultivar/Landrace | A_1 |
| Southern Europe | Traditional cultivar/Landrace | A_2 |
| Southern Europe | Traditional cultivar/Landrace | A_1 |
| Southern Europe | Traditional cultivar/Landrace | A_1 |
| Southern Europe | Traditional cultivar/Landrace | A_1 |
| Southern Europe | Traditional cultivar/Landrace | A_1 |
| Southern Europe | Traditional cultivar/Landrace | A_1 |
| Southern Europe | Traditional cultivar/Landrace | A_1 |
| Southern Europe | Traditional cultivar/Landrace | A_1 |
| Southern Europe | Traditional cultivar/Landrace | A_2 |
| Southern Europe | Traditional cultivar/Landrace | A_1 |
| Southern Europe | Traditional cultivar/Landrace | A_1 |
| Southern Europe | Traditional cultivar/Landrace | A_1 |
| Southern Europe | Traditional cultivar/Landrace | A_1 |
| Southern Europe | Traditional cultivar/Landrace | A_1 |
| Southern Europe | Traditional cultivar/Landrace | A_1 |
| Southern Europe | Traditional cultivar/Landrace | A_1 |
| Southern Europe | Traditional cultivar/Landrace | A_1 |
| Southern Europe | Traditional cultivar/Landrace | A_1 |
| Southern Europe | Traditional cultivar/Landrace | A_2 |
| Southern Europe | Traditional cultivar/Landrace | A_1 |
| Southern Europe | Traditional cultivar/Landrace | A_1 |
| Southern Europe | Traditional cultivar/Landrace | A_1 |
| Southern Europe | Traditional cultivar/Landrace | A_1 |
| Southern Europe | Traditional cultivar/Landrace | A_1 |
| Southern Europe | Traditional cultivar/Landrace | A_2 |
| Southern Europe | Traditional cultivar/Landrace | A_1 |
| Southern Europe | Traditional cultivar/Landrace | A_1 |
| Southern Europe | Traditional cultivar/Landrace | A_1 |
| Southern Europe | Traditional cultivar/Landrace | A_1 |
| Southern Europe | Traditional cultivar/Landrace | A_1 |
| Southern Europe | Traditional cultivar/Landrace | A_1 |

| | | |
|---|---|---|
| Southern Europe | Traditional cultivar/Landrace | A_1 |
| Southern Europe | Traditional cultivar/Landrace | A_1 |
| Southern Europe | Traditional cultivar/Landrace | A_1 |
| Southern Europe | Traditional cultivar/Landrace | A_2 |
| Southern Europe | Traditional cultivar/Landrace | A_1 |
| Southern Europe | Traditional cultivar/Landrace | A_2 |
| Southern Europe | Traditional cultivar/Landrace | A_1 |
| Southern Europe | Traditional cultivar/Landrace | A_1 |
| Southern Europe | Traditional cultivar/Landrace | A_2 |
| Southern Europe | Traditional cultivar/Landrace | A_1 |
| Southern Europe | Traditional cultivar/Landrace | A_1 |
| Southern Europe | Traditional cultivar/Landrace | A_1 |
| Southern Europe | Traditional cultivar/Landrace | A_1 |
| Southern Europe | Traditional cultivar/Landrace | A_1 |
| Southern Europe | Traditional cultivar/Landrace | A_1 |
| Southern Europe | Traditional cultivar/Landrace | A_1 |
| Southern Europe | Traditional cultivar/Landrace | A_1 |
| Southern Europe | Traditional cultivar/Landrace | A_1 |
| Southern Europe | Traditional cultivar/Landrace | A_2 |
| Southern Europe | Traditional cultivar/Landrace | A_1 |
| Southern Europe | Traditional cultivar/Landrace | A_2 |
| Southern Europe | Traditional cultivar/Landrace | A_1 |
| Southern Europe | Traditional cultivar/Landrace | A_1 |
| Southern Europe | Traditional cultivar/Landrace | A_1 |
| Southern Europe | Traditional cultivar/Landrace | A_1 |
| Southern Europe | Traditional cultivar/Landrace | A_2 |
| Southern Europe | Traditional cultivar/Landrace | A_1 |
| Southern Europe | Traditional cultivar/Landrace | A_2 |
| Southern Europe | Traditional cultivar/Landrace | A_1 |
| Southern Europe | Traditional cultivar/Landrace | A_1 |
| Southern Europe | Traditional cultivar/Landrace | A_1 |
| Southern Europe | Traditional cultivar/Landrace | A_1 |
| Southern Europe | Traditional cultivar/Landrace | A_1 |
| Southern Europe | Traditional cultivar/Landrace | A_2 |
| Southern Europe | Traditional cultivar/Landrace | A_2 |
| Southern Europe | Traditional cultivar/Landrace | A_2 |
| Southern Europe | Traditional cultivar/Landrace | A_2 |
| Southern Europe | Traditional cultivar/Landrace | A_2 |
| Southern Europe | Traditional cultivar/Landrace | A_1 |
| Southern Europe | Traditional cultivar/Landrace | A_2 |
| Southern Europe | Traditional cultivar/Landrace | A_1 |
| Southern Europe | Traditional cultivar/Landrace | A_1 |
| Southern Europe | Traditional cultivar/Landrace | A_1 |
| Southern Europe | Traditional cultivar/Landrace | A_1 |
| Southern Europe | Traditional cultivar/Landrace | A_1 |
| Southern Europe | Traditional cultivar/Landrace | A_1 |
| Southern Europe | Traditional cultivar/Landrace | A_1 |
| Southern Europe | Traditional cultivar/Landrace | A_1 |
| Southern Europe | Traditional cultivar/Landrace | A_1 |
| Southern Europe | Traditional cultivar/Landrace | A_1 |
| Southern Europe | Traditional cultivar/Landrace | A_1 |
| Southern Europe | Traditional cultivar/Landrace | A_1 |

| | | |
|---|---|---|
| Southern Europe | Traditional cultivar/Landrace | A_1 |
| Southern Europe | Traditional cultivar/Landrace | A_1 |
| Southern Europe | Traditional cultivar/Landrace | A_1 |
| Southern Europe | Traditional cultivar/Landrace | A_1 |
| Southern Europe | Traditional cultivar/Landrace | A_2 |
| Southern Europe | Traditional cultivar/Landrace | A_1 |
| Southern Europe | Traditional cultivar/Landrace | A_2 |
| Southern Europe | Traditional cultivar/Landrace | A_1 |
| Southern Europe | Traditional cultivar/Landrace | A_2 |
| Western Europe | Advanced/improved cultivar | A_1 |
| Southern Europe | Traditional cultivar/Landrace | A_1 |
| Western Europe | Advanced/improved cultivar | A_1 |
| Southern Europe | Traditional cultivar/Landrace | A_2 |
| Western Europe | Breeding/Research Material | A_1 |
| Western Europe | Breeding/Research Material | A_1 |
| Western Europe | Breeding/Research Material | A_1 |
| Southern Europe | Traditional cultivar/Landrace | A_2 |
| Southern Europe | Traditional cultivar/Landrace | A_1 |
| Southern Europe | Traditional cultivar/Landrace | A_1 |
| Southern Europe | Traditional cultivar/Landrace | A_1 |
| Southern Europe | Traditional cultivar/Landrace | A_1 |
| Southern Europe | Traditional cultivar/Landrace | A_1 |
| Southern Europe | Traditional cultivar/Landrace | A_1 |
| Southern Europe | Traditional cultivar/Landrace | A_1 |
| Southern Europe | Traditional cultivar/Landrace | A_1 |
| Southern Europe | Traditional cultivar/Landrace | A_1 |
| Southern Europe | Traditional cultivar/Landrace | A_1 |
| Southern Europe | Traditional cultivar/Landrace | A_1 |
| Southern Europe | Traditional cultivar/Landrace | A_1 |
| Southern Europe | Traditional cultivar/Landrace | A_1 |
| Southern Europe | Traditional cultivar/Landrace | A_1 |
| Southern Europe | Traditional cultivar/Landrace | A_1 |
| Southern Europe | Traditional cultivar/Landrace | A_1 |
| Southern Europe | Traditional cultivar/Landrace | A_1 |
| Southern Europe | Traditional cultivar/Landrace | A_1 |
| Southern Europe | Traditional cultivar/Landrace | A_2 |
| Southern Europe | Traditional cultivar/Landrace | A_1 |
| Southern Europe | Traditional cultivar/Landrace | A_1 |
| Southern Europe | Traditional cultivar/Landrace | A_1 |
| Southern Europe | Traditional cultivar/Landrace | A_1 |
| Southern Europe | Traditional cultivar/Landrace | A_1 |
| Southern Europe | Traditional cultivar/Landrace | A_1 |
| Southern Europe | Traditional cultivar/Landrace | A_1 |
| Southern Europe | Traditional cultivar/Landrace | A_1 |
| Southern Europe | Traditional cultivar/Landrace | A_1 |
| Southern Europe | Traditional cultivar/Landrace | A_1 |
| Southern Europe | Traditional cultivar/Landrace | A_1 |
| Southern Europe | Traditional cultivar/Landrace | A_1 |
| Southern Europe | Traditional cultivar/Landrace | A_1 |
| Southern Europe | Traditional cultivar/Landrace | A_1 |
| Southern Europe | Traditional cultivar/Landrace | A_1 |
| Southern Europe | Traditional cultivar/Landrace | A_1 |

| | | |
|---|---|---|
| Southern Europe | Traditional cultivar/Landrace | A_1 |
| Southern Europe | Traditional cultivar/Landrace | A_1 |
| Southern Europe | Traditional cultivar/Landrace | A_1 |
| Southern Europe | Traditional cultivar/Landrace | A_1 |
| Southern Europe | Traditional cultivar/Landrace | A_1 |
| Southern Europe | Traditional cultivar/Landrace | A_1 |
| Southern Europe | Traditional cultivar/Landrace | A_1 |
| Southern Europe | Traditional cultivar/Landrace | A_1 |
| Southern Europe | Traditional cultivar/Landrace | A_1 |
| Southern Europe | Traditional cultivar/Landrace | A_1 |
| Southern Europe | Traditional cultivar/Landrace | A_1 |
| Southern Europe | Traditional cultivar/Landrace | A_1 |
| Southern Europe | Traditional cultivar/Landrace | A_1 |
| Southern Europe | Traditional cultivar/Landrace | A_1 |
| Southern Europe | Traditional cultivar/Landrace | A_1 |
| Southern Europe | Traditional cultivar/Landrace | A_1 |
| Southern Europe | Traditional cultivar/Landrace | A_1 |
| Southern Europe | Traditional cultivar/Landrace | A_1 |
| Southern Europe | Traditional cultivar/Landrace | A_2 |
| Southern Europe | Traditional cultivar/Landrace | A_1 |
| Southern Europe | Traditional cultivar/Landrace | A_1 |
| Southern Europe | Traditional cultivar/Landrace | A_1 |
| Southern Europe | Traditional cultivar/Landrace | A_1 |
| Unknown | Traditional cultivar/Landrace | A_1 |
| Unknown | Traditional cultivar/Landrace | A_1 |
| Unknown | Traditional cultivar/Landrace | A_1 |
| Unknown | Traditional cultivar/Landrace | A_1 |
| Unknown | Traditional cultivar/Landrace | A_1 |
| Unknown | Traditional cultivar/Landrace | A_1 |
| Unknown | Traditional cultivar/Landrace | A_1 |
| Unknown | Traditional cultivar/Landrace | A_2 |
| Unknown | Traditional cultivar/Landrace | A_1 |
| Unknown | Traditional cultivar/Landrace | A_1 |
| Unknown | Traditional cultivar/Landrace | A_1 |
| Unknown | Traditional cultivar/Landrace | A_1 |
| Unknown | Traditional cultivar/Landrace | A_1 |
| Unknown | Traditional cultivar/Landrace | A_1 |
| Unknown | Traditional cultivar/Landrace | A_1 |
| Unknown | Traditional cultivar/Landrace | A_1 |
| Unknown | Traditional cultivar/Landrace | A_1 |
| Unknown | Traditional cultivar/Landrace | A_1 |
| Unknown | Traditional cultivar/Landrace | A_1 |
| Unknown | Traditional cultivar/Landrace | A_1 |
| Unknown | Traditional cultivar/Landrace | A_1 |
| Unknown | Traditional cultivar/Landrace | A_1 |
| Unknown | Traditional cultivar/Landrace | A_1 |
| Unknown | Traditional cultivar/Landrace | A_1 |
| Unknown | Traditional cultivar/Landrace | A_1 |
| Unknown | Traditional cultivar/Landrace | A_1 |
| Unknown | Traditional cultivar/Landrace | A_1 |
| Unknown | Traditional cultivar/Landrace | A_1 |
| Unknown | Traditional cultivar/Landrace | A_1 |

| | | |
|---|---|---|
| Unknown | Traditional cultivar/Landrace | A_1 |
| Unknown | Traditional cultivar/Landrace | A_1 |
| Unknown | Traditional cultivar/Landrace | A_1 |
| Unknown | Traditional cultivar/Landrace | A_1 |
| Unknown | Traditional cultivar/Landrace | A_1 |
| Unknown | Traditional cultivar/Landrace | A_1 |
| Unknown | Traditional cultivar/Landrace | A_1 |
| Unknown | Traditional cultivar/Landrace | A_1 |
| Unknown | Traditional cultivar/Landrace | A_1 |
| Unknown | Traditional cultivar/Landrace | A_1 |
| Unknown | Traditional cultivar/Landrace | A_1 |
| Unknown | Traditional cultivar/Landrace | A_1 |
| Unknown | Traditional cultivar/Landrace | A_1 |
| Unknown | Traditional cultivar/Landrace | A_1 |
| Unknown | Traditional cultivar/Landrace | A_1 |
| Unknown | Traditional cultivar/Landrace | A_1 |
| Unknown | Traditional cultivar/Landrace | A_1 |
| Unknown | Traditional cultivar/Landrace | A_1 |
| Southern Europe | Traditional cultivar/Landrace | A_1 |
| Southern Europe | Traditional cultivar/Landrace | A_1 |
| Southern Europe | Traditional cultivar/Landrace | A_1 |
| Southern Europe | Traditional cultivar/Landrace | A_1 |
| Southern Europe | Traditional cultivar/Landrace | A_1 |
| Southern Europe | Traditional cultivar/Landrace | A_1 |
| Southern Europe | Traditional cultivar/Landrace | A_2 |
| Southern Europe | Traditional cultivar/Landrace | A_1 |
| Southern Europe | Traditional cultivar/Landrace | A_2 |
| Southern Europe | Traditional cultivar/Landrace | A_1 |
| Southern Europe | Traditional cultivar/Landrace | A_1 |
| Southern Europe | Traditional cultivar/Landrace | A_1 |
| Southern Europe | Traditional cultivar/Landrace | A_1 |
| Southern Europe | Traditional cultivar/Landrace | A_1 |
| Southern Europe | Traditional cultivar/Landrace | A_1 |
| Southern Europe | Traditional cultivar/Landrace | A_1 |
| Southern Europe | Traditional cultivar/Landrace | A_1 |
| Southern Europe | Traditional cultivar/Landrace | A_1 |
| Southern Europe | Traditional cultivar/Landrace | A_1 |
| Southern Europe | Traditional cultivar/Landrace | A_2 |
| Southern Europe | Traditional cultivar/Landrace | A_1 |
| Southern Europe | Traditional cultivar/Landrace | A_1 |
| Southern Europe | Traditional cultivar/Landrace | A_1 |
| Southern Europe | Traditional cultivar/Landrace | A_1 |
| Southern Europe | Traditional cultivar/Landrace | A_1 |
| Southern Europe | Traditional cultivar/Landrace | A_1 |
| Southern Europe | Traditional cultivar/Landrace | A_1 |
| Southern Europe | Traditional cultivar/Landrace | A_1 |
| Southern Europe | Traditional cultivar/Landrace | A_1 |
| Southern Europe | Traditional cultivar/Landrace | A_1 |
| Southern Europe | Traditional cultivar/Landrace | A_1 |
| Southern Europe | Traditional cultivar/Landrace | A_1 |
| Southern Europe | Traditional cultivar/Landrace | A_1 |
| Southern Europe | Traditional cultivar/Landrace | A_1 |

| | | |
|---|---|---|
| Southern Europe | Traditional cultivar/Landrace | A_1 |
| Southern Europe | Traditional cultivar/Landrace | A_1 |
| Southern Europe | Traditional cultivar/Landrace | A_1 |
| Southern Europe | Traditional cultivar/Landrace | A_2 |
| Unknown | Traditional cultivar/Landrace | A_1 |
| Unknown | Traditional cultivar/Landrace | A_1 |
| Western Europe | Advanced/improved cultivar | A_1 |
| Western Europe | Advanced/improved cultivar | A_1 |
| Southern Europe | Traditional cultivar/Landrace | A_1 |
| Southern Europe | Traditional cultivar/Landrace | A_1 |
| Southern Europe | Traditional cultivar/Landrace | A_1 |
| Southern Europe | Traditional cultivar/Landrace | A_1 |
| Southern Europe | Advanced/improved cultivar | A_1 |
| Southern Europe | Advanced/improved cultivar | A_1 |
| Southern Europe | Traditional cultivar/Landrace | A_1 |
| Southern Europe | Traditional cultivar/Landrace | A_1 |
| Southern Europe | Traditional cultivar/Landrace | A_1 |
| Southern Europe | Traditional cultivar/Landrace | A_1 |
| Southern Europe | Traditional cultivar/Landrace | A_1 |
| Southern Europe | Traditional cultivar/Landrace | A_1 |
| Southern Europe | Traditional cultivar/Landrace | A_1 |
| Southern Europe | Traditional cultivar/Landrace | A_1 |
| Southern Europe | Traditional cultivar/Landrace | A_1 |
| Southern Europe | Traditional cultivar/Landrace | A_1 |
| Southern Europe | Traditional cultivar/Landrace | A_1 |
| Southern Europe | Traditional cultivar/Landrace | A_1 |
| Southern Europe | Traditional cultivar/Landrace | A_1 |
| Southern Europe | Traditional cultivar/Landrace | A_1 |
| Southern Europe | Traditional cultivar/Landrace | A_1 |
| Southern Europe | Traditional cultivar/Landrace | A_1 |
| Southern Europe | Traditional cultivar/Landrace | A_1 |
| Southern Europe | Traditional cultivar/Landrace | A_1 |
| Southern Europe | Traditional cultivar/Landrace | A_2 |
| Southern Europe | Traditional cultivar/Landrace | A_1 |
| Southern Europe | Traditional cultivar/Landrace | A_1 |
| Southern Europe | Traditional cultivar/Landrace | A_1 |
| Southern Europe | Traditional cultivar/Landrace | A_1 |
| Southern Europe | Traditional cultivar/Landrace | A_1 |
| Southern Europe | Traditional cultivar/Landrace | A_1 |
| Southern Europe | Traditional cultivar/Landrace | A_1 |
| Southern Europe | Traditional cultivar/Landrace | A_1 |
| Southern Europe | Traditional cultivar/Landrace | A_1 |
| Southern Europe | Traditional cultivar/Landrace | A_1 |
| Southern Europe | Traditional cultivar/Landrace | A_1 |
| Southern Europe | Traditional cultivar/Landrace | A_1 |
| Southern Europe | Traditional cultivar/Landrace | A_1 |
| Southern Europe | Traditional cultivar/Landrace | A_1 |
| Southern Europe | Traditional cultivar/Landrace | A_1 |
| Southern Europe | Traditional cultivar/Landrace | A_2 |
| Southern Europe | Traditional cultivar/Landrace | A_1 |
| Southern Europe | Traditional cultivar/Landrace | A_1 |
| Southern Europe | Traditional cultivar/Landrace | A_1 |

| | | |
|---|---|---|
| Southern Europe | Traditional cultivar/Landrace | A_1 |
| Southern Europe | Traditional cultivar/Landrace | A_1 |
| Southern Europe | Traditional cultivar/Landrace | A_1 |
| Southern Europe | Traditional cultivar/Landrace | A_2 |
| Southern Europe | Traditional cultivar/Landrace | A_2 |
| Southern Europe | Traditional cultivar/Landrace | A_1 |
| Southern Europe | Traditional cultivar/Landrace | A_1 |
| Southern Europe | Traditional cultivar/Landrace | A_1 |
| Southern Europe | Traditional cultivar/Landrace | A_1 |
| Southern Europe | Traditional cultivar/Landrace | A_1 |
| Southern Europe | Traditional cultivar/Landrace | A_1 |
| Southern Europe | Traditional cultivar/Landrace | A_1 |
| Southern Europe | Traditional cultivar/Landrace | A_1 |
| Southern Europe | Traditional cultivar/Landrace | A_1 |
| Southern Europe | Traditional cultivar/Landrace | A_1 |
| Southern Europe | Traditional cultivar/Landrace | A_1 |
| Southern Europe | Traditional cultivar/Landrace | A_1 |
| Southern Europe | Traditional cultivar/Landrace | A_1 |
| Southern Europe | Traditional cultivar/Landrace | A_1 |
| Southern Europe | Traditional cultivar/Landrace | A_1 |
| Southern Europe | Traditional cultivar/Landrace | A_1 |
| Southern Europe | Traditional cultivar/Landrace | A_1 |
| Southern Europe | Traditional cultivar/Landrace | A_1 |
| Southern Europe | Traditional cultivar/Landrace | A_1 |
| Southern Europe | Traditional cultivar/Landrace | A_2 |
| Southern Europe | Traditional cultivar/Landrace | A_1 |
| Southern Europe | Traditional cultivar/Landrace | A_1 |
| Southern Europe | Traditional cultivar/Landrace | A_1 |
| Southern Europe | Traditional cultivar/Landrace | A_1 |
| Southern Europe | Traditional cultivar/Landrace | A_2 |
| Southern Europe | Traditional cultivar/Landrace | A_1 |
| Southern Europe | Traditional cultivar/Landrace | A_2 |
| Southern Europe | Traditional cultivar/Landrace | A_1 |
| Southern Europe | Traditional cultivar/Landrace | A_1 |
| Southern Europe | Traditional cultivar/Landrace | A_1 |
| Southern Europe | Traditional cultivar/Landrace | A_1 |
| Southern Europe | Traditional cultivar/Landrace | A_1 |
| Southern Europe | Traditional cultivar/Landrace | A_1 |
| Southern Europe | Traditional cultivar/Landrace | A_1 |
| Southern Europe | Traditional cultivar/Landrace | A_1 |
| Southern Europe | Traditional cultivar/Landrace | A_1 |
| Southern Europe | Traditional cultivar/Landrace | A_1 |
| Southern Europe | Traditional cultivar/Landrace | A_1 |
| Southern Europe | Traditional cultivar/Landrace | A_1 |
| Southern Europe | Traditional cultivar/Landrace | A_1 |
| Southern Europe | Traditional cultivar/Landrace | A_1 |
| Southern Europe | Traditional cultivar/Landrace | A_1 |
| Southern Europe | Traditional cultivar/Landrace | A_1 |
| Southern Europe | Traditional cultivar/Landrace | A_1 |
| Southern Europe | Traditional cultivar/Landrace | A_1 |
| Southern Europe | Traditional cultivar/Landrace | A_2 |
| Southern Europe | Traditional cultivar/Landrace | A_1 |
| Southern Europe | Traditional cultivar/Landrace | A_2 |

| | | |
|---|---|---|
| Southern Europe | Traditional cultivar/Landrace | A_1 |
| Southern Europe | Traditional cultivar/Landrace | A_1 |
| Southern Europe | Traditional cultivar/Landrace | A_1 |
| Southern Europe | Traditional cultivar/Landrace | A_1 |
| Southern Europe | Traditional cultivar/Landrace | A_1 |
| Southern Europe | Traditional cultivar/Landrace | A_2 |
| Southern Europe | Traditional cultivar/Landrace | A_1 |
| Southern Europe | Traditional cultivar/Landrace | A_1 |
| Southern Europe | Traditional cultivar/Landrace | A_2 |
| Southern Europe | Traditional cultivar/Landrace | A_1 |
| Southern Europe | Traditional cultivar/Landrace | A_1 |
| Southern Europe | Traditional cultivar/Landrace | A_1 |
| Southern Europe | Traditional cultivar/Landrace | A_1 |
| Southern Europe | Traditional cultivar/Landrace | A_2 |
| Southern Europe | Traditional cultivar/Landrace | A_1 |
| Southern Europe | Traditional cultivar/Landrace | A_1 |
| Southern Europe | Traditional cultivar/Landrace | A_1 |
| Southern Europe | Traditional cultivar/Landrace | A_1 |
| Southern Europe | Traditional cultivar/Landrace | A_1 |
| Southern Europe | Traditional cultivar/Landrace | A_1 |
| Southern Europe | Traditional cultivar/Landrace | A_1 |
| Southern Europe | Traditional cultivar/Landrace | A_1 |
| Southern Europe | Wild | A_1 |
| Southern Europe | Traditional cultivar/Landrace | A_1 |
| Southern Europe | Traditional cultivar/Landrace | A_1 |
| Southern Europe | Traditional cultivar/Landrace | A_1 |
| Southern Europe | Traditional cultivar/Landrace | A_1 |
| Unknown | Traditional cultivar/Landrace | A_2 |
| Unknown | Traditional cultivar/Landrace | A_1 |
| Unknown | Traditional cultivar/Landrace | A_1 |
| Unknown | Traditional cultivar/Landrace | A_1 |
| Unknown | Traditional cultivar/Landrace | A_1 |
| Unknown | Traditional cultivar/Landrace | A_1 |
| Southern Europe | Traditional cultivar/Landrace | A_2 |
| Southern Europe | Traditional cultivar/Landrace | A_1 |
| Southern Europe | Traditional cultivar/Landrace | A_1 |
| Southern Europe | Traditional cultivar/Landrace | A_1 |
| Southern Europe | Traditional cultivar/Landrace | A_1 |
| Southern Europe | Traditional cultivar/Landrace | A_2 |
| Southern Europe | Traditional cultivar/Landrace | A_1 |
| Southern Europe | Traditional cultivar/Landrace | A_1 |
| Southern Europe | Traditional cultivar/Landrace | A_1 |
| Southern Europe | Traditional cultivar/Landrace | A_1 |
| Southern Europe | Traditional cultivar/Landrace | A_1 |
| Southern Europe | Traditional cultivar/Landrace | A_1 |
| Southern Europe | Traditional cultivar/Landrace | A_1 |
| Southern Europe | Traditional cultivar/Landrace | A_1 |
| Southern Europe | Traditional cultivar/Landrace | A_2 |
| Southern Europe | Traditional cultivar/Landrace | A_1 |
| Southern Europe | Traditional cultivar/Landrace | A_1 |
| Southern Europe | Traditional cultivar/Landrace | A_1 |
| Southern Europe | Traditional cultivar/Landrace | A_1 |

| | | |
|---|---|---|
| Southern Europe | Traditional cultivar/Landrace | A_1 |
| Eastern Europe | Breeding/Research Material | A_1 |
| Southern Europe | Traditional cultivar/Landrace | A_1 |
| Southern Europe | Traditional cultivar/Landrace | A_1 |
| Southern Europe | Traditional cultivar/Landrace | A_1 |
| Southern Europe | Traditional cultivar/Landrace | A_1 |
| Southern Europe | Traditional cultivar/Landrace | A_1 |
| Southern Europe | Traditional cultivar/Landrace | A_1 |
| Southern Europe | Traditional cultivar/Landrace | A_1 |
| Southern Europe | Traditional cultivar/Landrace | A_1 |
| Southern Europe | Traditional cultivar/Landrace | A_1 |
| Southern Europe | Traditional cultivar/Landrace | A_1 |
| Southern Europe | Traditional cultivar/Landrace | A_1 |
| Southern Europe | Traditional cultivar/Landrace | A_1 |
| Southern Europe | Traditional cultivar/Landrace | A_1 |
| Southern Europe | Traditional cultivar/Landrace | A_1 |
| Southern Europe | Traditional cultivar/Landrace | A_1 |
| Southern Europe | Traditional cultivar/Landrace | A_2 |
| Eastern Europe | Breeding/Research Material | A_2 |
| Southern Europe | Traditional cultivar/Landrace | A_1 |
| Western Europe | Advanced/improved cultivar | A_1 |
| Eastern Europe | Breeding/Research Material | B |
| Southern Europe | Traditional cultivar/Landrace | A_1 |
| Southern Europe | Traditional cultivar/Landrace | A_2 |
| Southern Europe | Traditional cultivar/Landrace | A_1 |
| Southern Europe | Traditional cultivar/Landrace | A_1 |
| Unknown | Traditional cultivar/Landrace | A_1 |
| Unknown | Traditional cultivar/Landrace | A_1 |
| Unknown | Traditional cultivar/Landrace | A_1 |
| Southern Europe | Traditional cultivar/Landrace | A_1 |
| Southern Europe | Traditional cultivar/Landrace | A_1 |
| Southern Europe | Traditional cultivar/Landrace | A_1 |
| Southern Europe | Traditional cultivar/Landrace | A_2 |
| Southern Europe | Traditional cultivar/Landrace | A_1 |
| Southern Europe | Traditional cultivar/Landrace | A_1 |
| Southern Europe | Traditional cultivar/Landrace | A_1 |
| Southern Europe | Traditional cultivar/Landrace | A_1 |
| Southern Europe | Traditional cultivar/Landrace | A_1 |
| Southern Europe | Traditional cultivar/Landrace | A_1 |
| Southern Europe | Traditional cultivar/Landrace | A_1 |
| Southern Europe | Traditional cultivar/Landrace | A_1 |
| Southern Europe | Traditional cultivar/Landrace | A_1 |
| Southern Europe | Traditional cultivar/Landrace | A_1 |
| Southern Europe | Traditional cultivar/Landrace | A_2 |
| Southern Europe | Traditional cultivar/Landrace | A_1 |
| Southern Europe | Traditional cultivar/Landrace | A_1 |
| Southern Europe | Traditional cultivar/Landrace | A_1 |
| Southern Europe | Traditional cultivar/Landrace | A_1 |
| Southern Europe | Traditional cultivar/Landrace | A_1 |
| Southern Europe | Traditional cultivar/Landrace | A_1 |
| Southern Europe | Traditional cultivar/Landrace | A_1 |
| Southern Europe | Traditional cultivar/Landrace | A_1 |

| | | |
|---|---|---|
| Southern Europe | Traditional cultivar/Landrace | A_1 |
| Southern Europe | Traditional cultivar/Landrace | A_1 |
| Southern Europe | Traditional cultivar/Landrace | A_1 |
| Southern Europe | Traditional cultivar/Landrace | A_1 |
| Southern Europe | Traditional cultivar/Landrace | A_1 |
| Southern Europe | Traditional cultivar/Landrace | A_1 |
| Southern Europe | Wild | A_1 |
| Southern Europe | Traditional cultivar/Landrace | A_1 |
| Southern Europe | Traditional cultivar/Landrace | A_1 |
| Southern Europe | Traditional cultivar/Landrace | A_1 |
| Southern Europe | Traditional cultivar/Landrace | A_1 |
| Southern Europe | Traditional cultivar/Landrace | A_2 |
| Southern Europe | Traditional cultivar/Landrace | A_1 |
| Southern Europe | Traditional cultivar/Landrace | A_1 |
| Unknown | Traditional cultivar/Landrace | A_1 |
| Unknown | Traditional cultivar/Landrace | A_1 |
| Southern Europe | Traditional cultivar/Landrace | A_1 |
| Unknown | Traditional cultivar/Landrace | A_1 |
| Southern Europe | Traditional cultivar/Landrace | A_1 |
| Southern Europe | Traditional cultivar/Landrace | A_1 |
| Southern Europe | Traditional cultivar/Landrace | A_1 |
| Southern Europe | Traditional cultivar/Landrace | A_1 |
| Southern Europe | Wild | A_1 |
| Southern Europe | Traditional cultivar/Landrace | A_1 |
| Southern Europe | Traditional cultivar/Landrace | A_1 |
| Southern Europe | Traditional cultivar/Landrace | A_1 |
| Southern Europe | Traditional cultivar/Landrace | A_1 |
| Eastern Europe | Traditional cultivar/Landrace | A_1 |
| Southern Europe | Traditional cultivar/Landrace | A_1 |
| Southern Europe | Traditional cultivar/Landrace | A_2 |
| Southern Europe | Traditional cultivar/Landrace | A_1 |
| Southern Europe | Traditional cultivar/Landrace | A_1 |
| Southern Europe | Traditional cultivar/Landrace | A_1 |
| Southern Europe | Traditional cultivar/Landrace | A_1 |
| Southern Europe | Traditional cultivar/Landrace | A_1 |
| Southern Europe | Traditional cultivar/Landrace | A_1 |
| Southern Europe | Wild | A_1 |
| Southern Europe | Unknown | A_1 |
| Southern Europe | Traditional cultivar/Landrace | A_1 |
| Unknown | Advanced/improved cultivar | A_1 |
| Western Europe | Advanced/improved cultivar | A_1 |
| America | Cultivar | A_1 |
| Eastern Europe | Advanced/improved cultivar | A_1 |
| Western Europe | Cultivar | A_1 |
| Western Europe | Cultivar | A_1 |
| Western Europe | Cultivar | A_1 |
| Western Europe | Cultivar | A_1 |
| Eastern Europe | Cultivar | A_1 |
| Southern Europe | Cultivar | A_1 |
| Western Europe | Cultivar | A_1 |
| Western Europe | Cultivar | A_1 |
| Middle East | Cultivar | A_1 |

| | | |
|---|---|---|
| Western Europe | Cultivar | A_1 |
| Eastern Europe | Cultivar | A_2 |
| Western Europe | Cultivar | A_1 |
| Western Europe | Cultivar | A_1 |
| Middle East | Unknown | A_1 |
| Middle East | Cultivar | A_1 |
| Eastern Europe | Cultivar | A_1 |
| Western Europe | Cultivar | A_1 |
| Eastern Africa | Cultivar | A_1 |
| Unknown | Unknown | A_1 |
| Northern Africa | Cultivar | A_1 |
| Northern Africa | Cultivar | A_1 |
| Eastern Europe | Unknown | A_1 |
| Eastern Europe | Unknown | A_1 |
| Eastern Europe | Unknown | A_1 |
| Western Europe | Advanced/improved cultivar | A_1 |
| Eastern Europe | Advanced/improved cultivar | A_1 |
| Western Europe | Advanced/improved cultivar | A_1 |
| America | Advanced/improved cultivar | A_1 |
| Eastern Europe | Unknown | A_2 |
| Eastern Europe | Unknown | A_1 |
| Eastern Europe | Unknown | A_1 |
| Eastern Europe | Cultivar | A_1 |
| Eastern Europe | Advanced/improved cultivar | A_1 |
| Australia and New Zealand | Advanced/improved cultivar | A_1 |
| Australia and New Zealand | Advanced/improved cultivar | A_2 |
| Southern Europe | Cultivar | A_1 |
| Southern Europe | Cultivar | A_1 |
| Southern Europe | Cultivar | A_1 |
| Southern Europe | Cultivar | A_1 |
| Southern Europe | Cultivar | A_1 |
| Southern Europe | Cultivar | A_1 |
| Southern Europe | Cultivar | A_1 |
| Southern Europe | Cultivar | A_2 |
| Southern Europe | Cultivar | A_1 |
| Southern Europe | Cultivar | A_1 |
| Western Europe | Cultivar | A_1 |
| Western Europe | Cultivar | A_1 |
| Western Europe | Cultivar | A_1 |
| Southern Europe | Cultivar | A_1 |
| Southern Europe | Cultivar | A_1 |
| Northern Africa | Cultivar | A_1 |
| Southern Europe | Cultivar | A_1 |
| Northern Africa | Cultivar | A_1 |
| Southern Europe | Breeding/Research Material | A_1 |
| Southern Europe | Breeding/Research Material | A_1 |
| Southern Europe | Breeding/Research Material | A_1 |
| Southern Europe | Breeding/Research Material | A_1 |
| Southern Europe | Breeding/Research Material | A_1 |
| Southern Europe | Breeding/Research Material | A_1 |
| Southern Europe | Breeding/Research Material | A_1 |
| Southern Europe | Breeding/Research Material | A_1 |

| | | |
|---|---|---|
| Southern Europe | Breeding/Research Material | A_1 |
| Southern Europe | Breeding/Research Material | A_1 |
| Southern Europe | Breeding/Research Material | A_1 |
| Southern Europe | Breeding/Research Material | A_1 |
| Southern Europe | Breeding/Research Material | A_1 |
| Southern Europe | Breeding/Research Material | A_1 |
| Southern Europe | Breeding/Research Material | A_1 |
| Eastern Europe | Cultivar | A_1 |
| Southern Europe | Cultivar | A_1 |
| Southern Europe | Cultivar | A_1 |
| Western Europe | Advanced/improved cultivar | A_1 |
| Western Europe | Advanced/improved cultivar | A_1 |
| Western Europe | Advanced/improved cultivar | A_1 |
| Eastern Europe | Advanced/improved cultivar | A_2 |
| Eastern Europe | Advanced/improved cultivar | A_1 |
| Eastern Europe | Breeding/Research Material | A_1 |
| Australia and New Zealand | Advanced/improved cultivar | A_1 |
| Australia and New Zealand | Advanced/improved cultivar | A_1 |
| Northern Africa | Advanced/improved cultivar | A_1 |
| Northern Africa | Traditional cultivar/Landrace | A_1 |
| Eastern Europe | Advanced/improved cultivar | A_2 |
| Eastern Europe | Advanced/improved cultivar | A_1 |
| Eastern Europe | Advanced/improved cultivar | A_1 |
| Eastern Europe | Advanced/improved cultivar | A_1 |
| Eastern Europe | Advanced/improved cultivar | A_1 |
| Middle East | Advanced/improved cultivar | A_1 |
| Eastern Africa | Cultivar | A_1 |
| Southern Europe | Cultivar | A_1 |
| Eastern Europe | Cultivar | A_1 |
| Eastern Europe | Cultivar | A_1 |
| Eastern Europe | Cultivar | A_1 |
| Southern Europe | Cultivar | A_1 |
| Northern Africa | Cultivar | A_1 |
| Northern Africa | Cultivar | A_1 |
| Western Europe | Cultivar | A_1 |
| Western Europe | Cultivar | A_1 |
| Western Europe | Cultivar | A_1 |
| Northern Africa | Cultivar | A_1 |
| Eastern Europe | Cultivar | A_1 |
| Middle East | Cultivar | A_1 |
| Southern Europe | Cultivar | A_1 |
| Eastern Europe | Cultivar | A_1 |
| Middle East | Cultivar | A_2 |
| Northern Africa | Cultivar | A_1 |
| Middle East | Cultivar | A_1 |
| Middle East | Cultivar | A_1 |
| Middle East | Cultivar | A_1 |
| Western Europe | Advanced/improved cultivar | A_1 |
| Eastern Europe | Advanced/improved cultivar | A_1 |
| Eastern Europe | Advanced/improved cultivar | A_1 |
| Eastern Europe | Cultivar | A_1 |
| Western Europe | Cultivar | A_1 |

| | | |
|---|---|---|
| America | Advanced/improved cultivar | A_1 |
| Eastern Europe | Advanced/improved cultivar | A_1 |
| Eastern Europe | Advanced/improved cultivar | A_1 |
| Eastern Europe | Cultivar | A_1 |
| Eastern Europe | Cultivar | A_1 |
| Eastern Europe | Cultivar | A_1 |
| Eastern Europe | Cultivar | A_1 |
| Eastern Europe | Cultivar | A_1 |
| Eastern Europe | Cultivar | A_1 |
| Eastern Europe | Advanced/improved cultivar | A_1 |
| Northern Africa | Cultivar | A_1 |
| Northern Africa | Cultivar | A_1 |
| Southern Europe | Cultivar | A_1 |
| Western Europe | Cultivar | A_1 |
| Western Europe | Cultivar | A_1 |
| Southern Europe | Cultivar | A_1 |
| Northern Africa | Cultivar | A_1 |
| Western Europe | Cultivar | A_1 |
| Northern Africa | Cultivar | A_1 |
| Southern Europe | Cultivar | A_1 |
| Southern Europe | Cultivar | A_1 |
| Eastern Europe | Cultivar | A_1 |
| Southern Europe | Cultivar | A_1 |
| Southern Europe | Cultivar | A_1 |
| Southern Europe | Cultivar | A_1 |
| Southern Europe | Cultivar | A_1 |
| Southern Europe | Cultivar | A_1 |
| Southern Europe | Cultivar | A_1 |
| Southern Europe | Cultivar | A_1 |
| Southern Europe | Cultivar | B |
| Southern Europe | Cultivar | B |
| Southern Europe | Cultivar | A_1 |
| Southern Europe | Cultivar | A_1 |
| Southern Europe | Cultivar | A_1 |
| Southern Europe | Cultivar | A_1 |
| Southern Europe | Cultivar | A_1 |
| Southern Europe | Cultivar | A_1 |
| Southern Europe | Cultivar | A_1 |
| Southern Europe | Cultivar | A_2 |
| Southern Europe | Cultivar | A_1 |
| Southern Europe | Cultivar | A_1 |
| Southern Europe | Cultivar | A_1 |
| Southern Europe | Cultivar | A_1 |
| Southern Europe | Cultivar | A_2 |
| Southern Europe | Cultivar | A_1 |
| Southern Europe | Cultivar | A_1 |
| Southern Europe | Cultivar | A_1 |
| America | Advanced/improved cultivar | A_1 |
| America | Advanced/improved cultivar | A_1 |
| America | Unknown | A_1 |
| Southern Europe | Cultivar | A_1 |
| Southern Europe | Cultivar | A_1 |

| | | |
|---|---|---|
| Northern Africa | Cultivar | A_1 |
| Northern Africa | Cultivar | A_1 |
| America | Cultivar | A_1 |
| America | Cultivar | A_1 |
| America | Cultivar | A_1 |
| America | Cultivar | A_1 |
| America | Cultivar | A_1 |
| America | Cultivar | A_1 |
| America | Advanced/improved cultivar | A_1 |
| Southern Europe | Wild | A_2 |
| Southern Europe | Advanced/improved cultivar | A_1 |
| Eastern Africa | Advanced/improved cultivar | A_1 |
| Middle East | Advanced/improved cultivar | A_1 |
| Northern Africa | Advanced/improved cultivar | A_1 |
| Western Europe | Advanced/improved cultivar | A_1 |
| Western Europe | Advanced/improved cultivar | A_1 |
| America | Advanced/improved cultivar | A_1 |
| Middle East | Cultivar | A_2 |
| Australia and New Zealand | Cultivar | A_1 |
| Southern Europe | Wild | A_2 |
| Southern Europe | Wild | A_1 |
| Southern Europe | Wild | A_1 |
| Australia and New Zealand | Cultivar | A_1 |
| Northern Africa | Advanced/improved cultivar | A_1 |
| Western Europe | Cultivar | A_1 |
| Western Europe | Cultivar | A_1 |
| Western Europe | Cultivar | A_1 |
| Western Europe | Cultivar | A_1 |
| Southern Europe | Cultivar | A_1 |
| Middle East | Cultivar | A_1 |
| Western Europe | Cultivar | A_1 |
| Western Europe | Cultivar | A_2 |
| Western Europe | Cultivar | A_1 |
| Western Europe | Cultivar | A_1 |
| Western Europe | Cultivar | A_1 |
| Western Europe | Cultivar | A_1 |
| Eastern Europe | Cultivar | A_1 |
| Western Europe | Advanced/improved cultivar | B |
| Unknown | Unknown | A_1 |
| Unknown | Unknown | A_1 |
| Unknown | Unknown | A_1 |
| Unknown | Unknown | B |
| Unknown | Unknown | A_1 |
| Unknown | Unknown | B |
| Unknown | Advanced/improved cultivar | A_1 |
| Unknown | Unknown | A_2 |
| Unknown | Unknown | A_1 |
| Unknown | Unknown | A_1 |
| Unknown | Unknown | A_1 |
| Unknown | Unknown | A_1 |
| Northern Africa | Landrace | A_1 |
| Unknown | Cultivar | A_1 |

| | | |
|---|---|---|
| Unknown | Landrace | A_1 |
| Southern Europe | Landrace | A_1 |

**GENEBANK**
P13-IHAR-PIB
P13-IHAR-PIB
P13-IHAR-PIB
P13-IHAR-PIB
P13-IHAR-PIB
P13-IHAR-PIB
P13-IHAR-PIB
P13-IHAR-PIB
P13-IHAR-PIB
P13-IHAR-PIB
P13-IHAR-PIB
P13-IHAR-PIB
P13-IHAR-PIB
P13-IHAR-PIB
P13-IHAR-PIB
P13-IHAR-PIB
P13-IHAR-PIB
P13-IHAR-PIB
P13-IHAR-PIB
P13-IHAR-PIB
P13-IHAR-PIB
P13-IHAR-PIB
P13-IHAR-PIB
P13-IHAR-PIB
P13-IHAR-PIB
P13-IHAR-PIB
P13-IHAR-PIB
P13-IHAR-PIB
P13-IHAR-PIB
P13-IHAR-PIB
P13-IHAR-PIB
P13-IHAR-PIB
P13-IHAR-PIB
P13-IHAR-PIB
P13-IHAR-PIB
P13-IHAR-PIB
P13-IHAR-PIB
P13-IHAR-PIB
P13-IHAR-PIB
P13-IHAR-PIB
P13-IHAR-PIB
P13-IHAR-PIB
P13-IHAR-PIB
P13-IHAR-PIB
P13-IHAR-PIB
P13-IHAR-PIB
P13-IHAR-PIB
P13-IHAR-PIB
P13-IHAR-PIB
P13-IHAR-PIB

P13-IHAR-PIB
P13-IHAR-PIB
P13-IHAR-PIB
P13-IHAR-PIB
P13-IHAR-PIB
P13-IHAR-PIB
P13-IHAR-PIB
P13-IHAR-PIB
P13-IHAR-PIB
P13-IHAR-PIB
P13-IHAR-PIB
P13-IHAR-PIB
P13-IHAR-PIB
P13-IHAR-PIB
P13-IHAR-PIB
P13-IHAR-PIB
P13-IHAR-PIB
P13-IHAR-PIB
P13-IHAR-PIB
P13-IHAR-PIB
P13-IHAR-PIB
P13-IHAR-PIB
P13-IHAR-PIB
P13-IHAR-PIB
P13-IHAR-PIB
P13-IHAR-PIB
P13-IHAR-PIB
P13-IHAR-PIB
P13-IHAR-PIB
P13-IHAR-PIB
P13-IHAR-PIB
P13-IHAR-PIB
P13-IHAR-PIB
P13-IHAR-PIB
P13-IHAR-PIB
P13-IHAR-PIB
P13-IHAR-PIB
P13-IHAR-PIB
P13-IHAR-PIB
P13-IHAR-PIB
P13-IHAR-PIB
P13-IHAR-PIB
P13-IHAR-PIB
P13-IHAR-PIB
P13-IHAR-PIB
P13-IHAR-PIB
P13-IHAR-PIB
P13-IHAR-PIB
P13-IHAR-PIB
P13-IHAR-PIB
P13-IHAR-PIB
P13-IHAR-PIB

P13-IHAR-PIB
P13-IHAR-PIB
P13-IHAR-PIB
P13-IHAR-PIB
P13-IHAR-PIB
P13-IHAR-PIB
P13-IHAR-PIB
P13-IHAR-PIB
P13-IHAR-PIB
P13-IHAR-PIB
P13-IHAR-PIB
P13-IHAR-PIB
P13-IHAR-PIB
P13-IHAR-PIB
P13-IHAR-PIB
P13-IHAR-PIB
P13-IHAR-PIB
P13-IHAR-PIB
P13-IHAR-PIB
P13-IHAR-PIB
P13-IHAR-PIB
P13-IHAR-PIB
P13-IHAR-PIB
P13-IHAR-PIB
P13-IHAR-PIB
P13-IHAR-PIB
P13-IHAR-PIB
P13-IHAR-PIB
P13-IHAR-PIB
P13-IHAR-PIB
P13-IHAR-PIB
P13-IHAR-PIB
P13-IHAR-PIB
P13-IHAR-PIB
P13-IHAR-PIB
P13-IHAR-PIB
P13-IHAR-PIB
P13-IHAR-PIB
P13-IHAR-PIB
P13-IHAR-PIB
P13-IHAR-PIB
P13-IHAR-PIB
P13-IHAR-PIB
P13-IHAR-PIB
P13-IHAR-PIB
P13-IHAR-PIB
P13-IHAR-PIB
P13-IHAR-PIB
P13-IHAR-PIB
P13-IHAR-PIB
P13-IHAR-PIB
P13-IHAR-PIB

P13-IHAR-PIB
P13-IHAR-PIB
P13-IHAR-PIB
P13-IHAR-PIB
P13-IHAR-PIB
P13-IHAR-PIB
P13-IHAR-PIB
P13-IHAR-PIB
P13-IHAR-PIB
P13-IHAR-PIB
P13-IHAR-PIB
P13-IHAR-PIB
P13-IHAR-PIB
P13-IHAR-PIB
P13-IHAR-PIB
P13-IHAR-PIB
P13-IHAR-PIB
P13-IHAR-PIB
P13-IHAR-PIB
P13-IHAR-PIB
P13-IHAR-PIB
P13-IHAR-PIB
P13-IHAR-PIB
P13-IHAR-PIB
P13-IHAR-PIB
P13-IHAR-PIB
P13-IHAR-PIB
P13-IHAR-PIB
P13-IHAR-PIB
P13-IHAR-PIB
P13-IHAR-PIB
P13-IHAR-PIB
P13-IHAR-PIB
P13-IHAR-PIB
P13-IHAR-PIB
P13-IHAR-PIB
P13-IHAR-PIB
P13-IHAR-PIB
P13-IHAR-PIB
P13-IHAR-PIB
P13-IHAR-PIB
P13-IHAR-PIB
P13-IHAR-PIB
P13-IHAR-PIB
P13-IHAR-PIB
P13-IHAR-PIB
P13-IHAR-PIB
P13-IHAR-PIB
P13-IHAR-PIB
P13-IHAR-PIB
P13-IHAR-PIB
P13-IHAR-PIB

P13-IHAR-PIB
P13-IHAR-PIB
P13-IHAR-PIB
P13-IHAR-PIB
P13-IHAR-PIB
P13-IHAR-PIB
P13-IHAR-PIB
P13-IHAR-PIB
P13-IHAR-PIB
P13-IHAR-PIB
P13-IHAR-PIB
P13-IHAR-PIB
P13-IHAR-PIB
P13-IHAR-PIB
P13-IHAR-PIB
P13-IHAR-PIB
P13-IHAR-PIB
P13-IHAR-PIB
P13-IHAR-PIB
P13-IHAR-PIB
P13-IHAR-PIB
P13-IHAR-PIB
P13-IHAR-PIB
P13-IHAR-PIB
P13-IHAR-PIB
P13-IHAR-PIB
P13-IHAR-PIB
P13-IHAR-PIB
P13-IHAR-PIB
P13-IHAR-PIB
P13-IHAR-PIB
P13-IHAR-PIB
P13-IHAR-PIB
P13-IHAR-PIB
P13-IHAR-PIB
P13-IHAR-PIB
P13-IHAR-PIB
P13-IHAR-PIB
P13-IHAR-PIB
P13-IHAR-PIB
P13-IHAR-PIB
P13-IHAR-PIB
P13-IHAR-PIB
P13-IHAR-PIB
P13-IHAR-PIB
P13-IHAR-PIB
P13-IHAR-PIB
P13-IHAR-PIB
P13-IHAR-PIB
P13-IHAR-PIB
P13-IHAR-PIB
P13-IHAR-PIB

P13-IHAR-PIB
P13-IHAR-PIB
P13-IHAR-PIB
P13-IHAR-PIB
P13-IHAR-PIB
P13-IHAR-PIB
P13-IHAR-PIB
P13-IHAR-PIB
P13-IHAR-PIB
P13-IHAR-PIB
P13-IHAR-PIB
P13-IHAR-PIB
P13-IHAR-PIB
P13-IHAR-PIB
P13-IHAR-PIB
P13-IHAR-PIB
P13-IHAR-PIB
P13-IHAR-PIB
P13-IHAR-PIB
P13-IHAR-PIB
P13-IHAR-PIB
P13-IHAR-PIB
P13-IHAR-PIB
P13-IHAR-PIB
P13-IHAR-PIB
P13-IHAR-PIB
P13-IHAR-PIB
P13-IHAR-PIB
P13-IHAR-PIB
P13-IHAR-PIB
P13-IHAR-PIB
P13-IHAR-PIB
P13-IHAR-PIB
P13-IHAR-PIB
P13-IHAR-PIB
P13-IHAR-PIB
P13-IHAR-PIB
P13-IHAR-PIB
P13-IHAR-PIB
P13-IHAR-PIB
P13-IHAR-PIB
P13-IHAR-PIB
P13-IHAR-PIB
P13-IHAR-PIB
P13-IHAR-PIB
P13-IHAR-PIB
P13-IHAR-PIB
P13-IHAR-PIB
P13-IHAR-PIB
P13-IHAR-PIB
P13-IHAR-PIB
P13-IHAR-PIB

P13-IHAR-PIB
P13-IHAR-PIB
P13-IHAR-PIB
P13-IHAR-PIB
P13-IHAR-PIB
P13-IHAR-PIB
P13-IHAR-PIB
P13-IHAR-PIB
P13-IHAR-PIB
P13-IHAR-PIB
P13-IHAR-PIB
P13-IHAR-PIB
P13-IHAR-PIB
P13-IHAR-PIB
P13-IHAR-PIB
P13-IHAR-PIB
P13-IHAR-PIB
P13-IHAR-PIB
P13-IHAR-PIB
P13-IHAR-PIB
P13-IHAR-PIB
P2-IPK
P2-IPK
P2-IPK
P2-IPK
P2-IPK
P2-IPK
P2-IPK
P2-IPK
P2-IPK
P2-IPK
P2-IPK
P2-IPK
P2-IPK
P2-IPK
P2-IPK
P2-IPK
P2-IPK
P2-IPK
P2-IPK
P2-IPK
P2-IPK
P2-IPK
P2-IPK
P2-IPK
P2-IPK
P2-IPK
P2-IPK
P2-IPK
P2-IPK
P2-IPK
P2-IPK

P2-IPK
P2-IPK
P2-IPK
P2-IPK
P2-IPK
P2-IPK
P2-IPK
P2-IPK
P2-IPK
P2-IPK
P2-IPK
P2-IPK
P2-IPK
P2-IPK
P2-IPK
P2-IPK
P2-IPK
P2-IPK
P2-IPK
P2-IPK
P2-IPK
P2-IPK
P2-IPK
P2-IPK
P2-IPK
P2-IPK
P2-IPK
P2-IPK
P2-IPK
P2-IPK
P2-IPK
P2-IPK
P2-IPK
P2-IPK
P2-IPK
P2-IPK
P2-IPK
P2-IPK
P2-IPK
P2-IPK
P2-IPK
P2-IPK
P2-IPK
P2-IPK
P2-IPK
P2-IPK
P2-IPK
P2-IPK
P2-IPK
P2-IPK
P2-IPK
P2-IPK

P2-IPK
P2-IPK
P2-IPK
P2-IPK
P2-IPK
P2-IPK
P2-IPK
P2-IPK
P2-IPK
P2-IPK
P2-IPK
P2-IPK
P2-IPK
P2-IPK
P2-IPK
P2-IPK
P2-IPK
P2-IPK
P2-IPK
P2-IPK
P2-IPK
P2-IPK
P2-IPK
P2-IPK
P2-IPK
P2-IPK
P2-IPK
P2-IPK
P2-IPK
P2-IPK
P2-IPK
P2-IPK
P2-IPK
P2-IPK
P2-IPK
P2-IPK
P2-IPK
P2-IPK
P2-IPK
P2-IPK
P2-IPK
P2-IPK
P2-IPK
P2-IPK
P2-IPK
P2-IPK
P2-IPK
P2-IPK
P2-IPK
P2-IPK
P2-IPK
P2-IPK

P2-IPK
P2-IPK
P2-IPK
P2-IPK
P2-IPK
P2-IPK
P2-IPK
P2-IPK
P2-IPK
P2-IPK
P2-IPK
P2-IPK
P2-IPK
P2-IPK
P2-IPK
P2-IPK
P2-IPK
P2-IPK
P2-IPK
P2-IPK
P2-IPK
P2-IPK
P2-IPK
P2-IPK
P2-IPK
P2-IPK
P2-IPK
P2-IPK
P2-IPK
P2-IPK
P2-IPK
P2-IPK
P2-IPK
P2-IPK
P2-IPK
P2-IPK
P2-IPK
P2-IPK
P2-IPK
P2-IPK
P2-IPK
P2-IPK
P2-IPK
P2-IPK
P2-IPK
P2-IPK
P2-IPK
P2-IPK
P2-IPK
P2-IPK
P2-IPK
P2-IPK

P2-IPK
P2-IPK
P2-IPK
P2-IPK
P2-IPK
P2-IPK
P2-IPK
P2-IPK
P2-IPK
P2-IPK
P2-IPK
P2-IPK
P2-IPK
P2-IPK
P2-IPK
P2-IPK
P2-IPK
P2-IPK
P2-IPK
P2-IPK
P2-IPK
P2-IPK
P2-IPK
P2-IPK
P2-IPK
P2-IPK
P2-IPK
P2-IPK
P2-IPK
P2-IPK
P2-IPK
P2-IPK
P2-IPK
P2-IPK
P2-IPK
P2-IPK
P2-IPK
P2-IPK
P2-IPK
P2-IPK
P2-IPK
P2-IPK
P2-IPK
P2-IPK
P2-IPK
P2-IPK
P2-IPK
P2-IPK
P2-IPK
P2-IPK
P2-IPK
P2-IPK

P2-IPK
P2-IPK
P2-IPK
P2-IPK
P2-IPK
P2-IPK
P2-IPK
P2-IPK
P2-IPK
P2-IPK
P2-IPK
P2-IPK
P2-IPK
P2-IPK
P2-IPK
P2-IPK
P2-IPK
P2-IPK
P2-IPK
P2-IPK
P2-IPK
P2-IPK
P2-IPK
P2-IPK
P2-IPK
P2-IPK
P2-IPK
P2-IPK
P2-IPK
P2-IPK
P2-IPK
P2-IPK
P2-IPK
P2-IPK
P2-IPK
P2-IPK
P2-IPK
P2-IPK
P2-IPK
P2-IPK
P2-IPK
P2-IPK
P2-IPK
P2-IPK
P2-IPK
P2-IPK
P2-IPK
P2-IPK
Wageningen Uni
Wageningen Uni
Wageningen Uni
Wageningen Uni

Wageningen Uni
Wageningen Uni
Wageningen Uni
Wageningen Uni
Wageningen Uni
Wageningen Uni
Wageningen Uni
Wageningen Uni
Wageningen Uni
GRIN CZEH
GRIN CZEH
GRIN CZEH
GRIN CZEH
GRIN CZEH
GRIN CZEH
GRIN CZEH
GRIN CZEH
GRIN CZEH
GRIN CZEH
GRIN CZEH
GRIN CZEH
GRIN CZEH
GRIN CZEH
GRIN CZEH
GRIN CZEH
GRIN CZEH
GRIN CZEH
GRIN CZEH
GRIN CZEH
GRIN CZEH
GRIN CZEH
GRIN CZEH
GRIN CZEH
GRIN CZEH
GRIN CZEH
GRIN CZEH
GRIN CZEH
GRIN CZEH
GRIN CZEH
GRIN CZEH
GRIN CZEH
GRIN CZEH
GRIN CZEH
GRIN CZEH
GRIN CZEH
GRIN CZEH
GRIN CZEH
USDA
P13-IHAR-PIB
Private donor
Private donor
P13-IHAR-PIB

P13-IHAR-PIB
P13-IHAR-PIB
P13-IHAR-PIB
P13-IHAR-PIB
P13-IHAR-PIB
P13-IHAR-PIB
P13-IHAR-PIB
P13-IHAR-PIB
P13-IHAR-PIB
P13-IHAR-PIB
P13-IHAR-PIB
P13-IHAR-PIB
MSB Kew Garden
MSB Kew Garden
MSB Kew Garden
MSB Kew Garden
MSB Kew Garden
MSB Kew Garden
MSB Kew Garden
MSB Kew Garden
MSB Kew Garden
MSB Kew Garden
MSB Kew Garden
MSB Kew Garden
MSB Kew Garden
MSB Kew Garden
MSB Kew Garden
MSB Kew Garden
MSB Kew Garden
MSB Kew Garden
MSB Kew Garden
MSB Kew Garden
MSB Kew Garden
MSB Kew Garden
MSB Kew Garden
MSB Kew Garden
MSB Kew Garden
MSB Kew Garden
MSB Kew Garden
MSB Kew Garden
MSB Kew Garden
MSB Kew Garden
MSB Kew Garden
MSB Kew Garden
MSB Kew Garden
MSB Kew Garden
MSB Kew Garden
MSB Kew Garden
MSB Kew Garden
MSB Kew Garden
MSB Kew Garden
MSB Kew Garden

MSB Kew Garden
MSB Kew Garden
MSB Kew Garden
MSB Kew Garden
MSB Kew Garden
MSB Kew Garden
MSB Kew Garden
MSB Kew Garden
MSB Kew Garden
MSB Kew Garden
MSB Kew Garden
MSB Kew Garden
MSB Kew Garden
MSB Kew Garden
MSB Kew Garden
MSB Kew Garden
MSB Kew Garden
MSB Kew Garden
MSB Kew Garden
MSB Kew Garden
MSB Kew Garden
MSB Kew Garden
MSB Kew Garden
MSB Kew Garden
MSB Kew Garden
MSB Kew Garden
MSB Kew Garden
MSB Kew Garden
MSB Kew Garden
MSB Kew Garden
MSB Kew Garden
MSB Kew Garden
MSB Kew Garden
MSB Kew Garden
MSB Kew Garden
MSB Kew Garden
MSB Kew Garden
MSB Kew Garden
MSB Kew Garden
MSB Kew Garden
MSB Kew Garden
MSB Kew Garden
MSB Kew Garden
MSB Kew Garden
MSB Kew Garden
MSB Kew Garden
MSB Kew Garden
MSB Kew Garden
MSB Kew Garden
MSB Kew Garden
MSB Kew Garden
MSB Kew Garden

MSB Kew Garden
MSB Kew Garden
MSB Kew Garden
MSB Kew Garden
MSB Kew Garden
MSB Kew Garden
MSB Kew Garden
MSB Kew Garden
MSB Kew Garden
MSB Kew Garden
MSB Kew Garden
MSB Kew Garden
MSB Kew Garden
MSB Kew Garden
MSB Kew Garden
MSB Kew Garden
MSB Kew Garden
MSB Kew Garden
MSB Kew Garden
MSB Kew Garden
MSB Kew Garden
MSB Kew Garden
MSB Kew Garden
MSB Kew Garden
MSB Kew Garden
MSB Kew Garden
MSB Kew Garden
MSB Kew Garden
MSB Kew Garden
MSB Kew Garden
MSB Kew Garden
MSB Kew Garden
MSB Kew Garden
MSB Kew Garden
MSB Kew Garden
MSB Kew Garden
MSB Kew Garden
MSB Kew Garden
MSB Kew Garden
MSB Kew Garden
MSB Kew Garden
MSB Kew Garden
MSB Kew Garden
MSB Kew Garden
MSB Kew Garden
MSB Kew Garden
MSB Kew Garden
MSB Kew Garden
MSB Kew Garden
MSB Kew Garden
MSB Kew Garden
MSB Kew Garden

MSB Kew Garden
MSB Kew Garden
MSB Kew Garden
AUSTRALIAN GENBANK
AUSTRALIAN GENBANK
AUSTRALIAN GENBANK
AUSTRALIAN GENBANK
AUSTRALIAN GENBANK
AUSTRALIAN GENBANK
AUSTRALIAN GENBANK
AUSTRALIAN GENBANK
AUSTRALIAN GENBANK
AUSTRALIAN GENBANK
AUSTRALIAN GENBANK
AUSTRALIAN GENBANK
AUSTRALIAN GENBANK
AUSTRALIAN GENBANK
AUSTRALIAN GENBANK
AUSTRALIAN GENBANK
AUSTRALIAN GENBANK
AUSTRALIAN GENBANK
AUSTRALIAN GENBANK
AUSTRALIAN GENBANK
AUSTRALIAN GENBANK
AUSTRALIAN GENBANK
AUSTRALIAN GENBANK
AUSTRALIAN GENBANK
AUSTRALIAN GENBANK
AUSTRALIAN GENBANK
AUSTRALIAN GENBANK
AUSTRALIAN GENBANK
AUSTRALIAN GENBANK
AUSTRALIAN GENBANK
AUSTRALIAN GENBANK
AUSTRALIAN GENBANK
AUSTRALIAN GENBANK
AUSTRALIAN GENBANK
AUSTRALIAN GENBANK
AUSTRALIAN GENBANK
AUSTRALIAN GENBANK
AUSTRALIAN GENBANK
AUSTRALIAN GENBANK
AUSTRALIAN GENBANK
AUSTRALIAN GENBANK
AUSTRALIAN GENBANK
AUSTRALIAN GENBANK
AUSTRALIAN GENBANK
AUSTRALIAN GENBANK
AUSTRALIAN GENBANK
AUSTRALIAN GENBANK
AUSTRALIAN GENBANK
AUSTRALIAN GENBANK

AUSTRALIAN GENBANK
AUSTRALIAN GENBANK
AUSTRALIAN GENBANK
AUSTRALIAN GENBANK
AUSTRALIAN GENBANK
AUSTRALIAN GENBANK
AUSTRALIAN GENBANK
AUSTRALIAN GENBANK
AUSTRALIAN GENBANK
AUSTRALIAN GENBANK
AUSTRALIAN GENBANK
AUSTRALIAN GENBANK
AUSTRALIAN GENBANK
AUSTRALIAN GENBANK
AUSTRALIAN GENBANK
AUSTRALIAN GENBANK
AUSTRALIAN GENBANK
AUSTRALIAN GENBANK
AUSTRALIAN GENBANK
AUSTRALIAN GENBANK
AUSTRALIAN GENBANK
AUSTRALIAN GENBANK
AUSTRALIAN GENBANK
AUSTRALIAN GENBANK
AUSTRALIAN GENBANK
AUSTRALIAN GENBANK
AUSTRALIAN GENBANK
AUSTRALIAN GENBANK
AUSTRALIAN GENBANK
AUSTRALIAN GENBANK
AUSTRALIAN GENBANK
AUSTRALIAN GENBANK
AUSTRALIAN GENBANK
AUSTRALIAN GENBANK
AUSTRALIAN GENBANK
AUSTRALIAN GENBANK
AUSTRALIAN GENBANK
AUSTRALIAN GENBANK
AUSTRALIAN GENBANK
AUSTRALIAN GENBANK
AUSTRALIAN GENBANK
AUSTRALIAN GENBANK
AUSTRALIAN GENBANK
AUSTRALIAN GENBANK
AUSTRALIAN GENBANK
AUSTRALIAN GENBANK
AUSTRALIAN GENBANK
AUSTRALIAN GENBANK
AUSTRALIAN GENBANK
AUSTRALIAN GENBANK
AUSTRALIAN GENBANK
AUSTRALIAN GENBANK

AUSTRALIAN GENBANK
AUSTRALIAN GENBANK
AUSTRALIAN GENBANK
AUSTRALIAN GENBANK
AUSTRALIAN GENBANK
AUSTRALIAN GENBANK
AUSTRALIAN GENBANK
AUSTRALIAN GENBANK
AUSTRALIAN GENBANK
AUSTRALIAN GENBANK
AUSTRALIAN GENBANK
AUSTRALIAN GENBANK
AUSTRALIAN GENBANK
AUSTRALIAN GENBANK
AUSTRALIAN GENBANK
AUSTRALIAN GENBANK
AUSTRALIAN GENBANK
AUSTRALIAN GENBANK
AUSTRALIAN GENBANK
AUSTRALIAN GENBANK
AUSTRALIAN GENBANK
AUSTRALIAN GENBANK
AUSTRALIAN GENBANK
AUSTRALIAN GENBANK
AUSTRALIAN GENBANK
AUSTRALIAN GENBANK
AUSTRALIAN GENBANK
AUSTRALIAN GENBANK
AUSTRALIAN GENBANK
AUSTRALIAN GENBANK
AUSTRALIAN GENBANK
AUSTRALIAN GENBANK
AUSTRALIAN GENBANK
AUSTRALIAN GENBANK
AUSTRALIAN GENBANK
AUSTRALIAN GENBANK
AUSTRALIAN GENBANK
AUSTRALIAN GENBANK
AUSTRALIAN GENBANK
AUSTRALIAN GENBANK
AUSTRALIAN GENBANK
AUSTRALIAN GENBANK
AUSTRALIAN GENBANK
AUSTRALIAN GENBANK
AUSTRALIAN GENBANK
AUSTRALIAN GENBANK
AUSTRALIAN GENBANK
AUSTRALIAN GENBANK
AUSTRALIAN GENBANK
AUSTRALIAN GENBANK
AUSTRALIAN GENBANK
AUSTRALIAN GENBANK

AUSTRALIAN GENBANK
AUSTRALIAN GENBANK
AUSTRALIAN GENBANK
AUSTRALIAN GENBANK
AUSTRALIAN GENBANK
AUSTRALIAN GENBANK
AUSTRALIAN GENBANK
AUSTRALIAN GENBANK
AUSTRALIAN GENBANK
AUSTRALIAN GENBANK
AUSTRALIAN GENBANK
AUSTRALIAN GENBANK
AUSTRALIAN GENBANK
AUSTRALIAN GENBANK
AUSTRALIAN GENBANK
AUSTRALIAN GENBANK
AUSTRALIAN GENBANK
AUSTRALIAN GENBANK
AUSTRALIAN GENBANK
AUSTRALIAN GENBANK
AUSTRALIAN GENBANK
AUSTRALIAN GENBANK
AUSTRALIAN GENBANK
AUSTRALIAN GENBANK
AUSTRALIAN GENBANK
AUSTRALIAN GENBANK
AUSTRALIAN GENBANK
AUSTRALIAN GENBANK
AUSTRALIAN GENBANK
AUSTRALIAN GENBANK
AUSTRALIAN GENBANK
AUSTRALIAN GENBANK
AUSTRALIAN GENBANK
AUSTRALIAN GENBANK
AUSTRALIAN GENBANK
AUSTRALIAN GENBANK
AUSTRALIAN GENBANK
AUSTRALIAN GENBANK
AUSTRALIAN GENBANK
AUSTRALIAN GENBANK
AUSTRALIAN GENBANK
AUSTRALIAN GENBANK
AUSTRALIAN GENBANK
AUSTRALIAN GENBANK
AUSTRALIAN GENBANK
AUSTRALIAN GENBANK
AUSTRALIAN GENBANK
AUSTRALIAN GENBANK
AUSTRALIAN GENBANK
AUSTRALIAN GENBANK
AUSTRALIAN GENBANK
AUSTRALIAN GENBANK

AUSTRALIAN GENBANK
AUSTRALIAN GENBANK
AUSTRALIAN GENBANK
AUSTRALIAN GENBANK
AUSTRALIAN GENBANK
AUSTRALIAN GENBANK
AUSTRALIAN GENBANK
AUSTRALIAN GENBANK
AUSTRALIAN GENBANK
AUSTRALIAN GENBANK
AUSTRALIAN GENBANK
AUSTRALIAN GENBANK
AUSTRALIAN GENBANK
AUSTRALIAN GENBANK
AUSTRALIAN GENBANK
AUSTRALIAN GENBANK
AUSTRALIAN GENBANK
AUSTRALIAN GENBANK
AUSTRALIAN GENBANK
AUSTRALIAN GENBANK
AUSTRALIAN GENBANK
AUSTRALIAN GENBANK
AUSTRALIAN GENBANK
AUSTRALIAN GENBANK
AUSTRALIAN GENBANK
AUSTRALIAN GENBANK
AUSTRALIAN GENBANK
AUSTRALIAN GENBANK
AUSTRALIAN GENBANK
AUSTRALIAN GENBANK
AUSTRALIAN GENBANK
AUSTRALIAN GENBANK
AUSTRALIAN GENBANK
AUSTRALIAN GENBANK
AUSTRALIAN GENBANK
AUSTRALIAN GENBANK
AUSTRALIAN GENBANK
AUSTRALIAN GENBANK
AUSTRALIAN GENBANK
AUSTRALIAN GENBANK
AUSTRALIAN GENBANK
AUSTRALIAN GENBANK
AUSTRALIAN GENBANK
AUSTRALIAN GENBANK
AUSTRALIAN GENBANK
AUSTRALIAN GENBANK
AUSTRALIAN GENBANK
AUSTRALIAN GENBANK
AUSTRALIAN GENBANK
AUSTRALIAN GENBANK
AUSTRALIAN GENBANK
AUSTRALIAN GENBANK

AUSTRALIAN GENBANK
AUSTRALIAN GENBANK
AUSTRALIAN GENBANK
AUSTRALIAN GENBANK
AUSTRALIAN GENBANK
AUSTRALIAN GENBANK
AUSTRALIAN GENBANK
AUSTRALIAN GENBANK
AUSTRALIAN GENBANK
AUSTRALIAN GENBANK
AUSTRALIAN GENBANK
AUSTRALIAN GENBANK
AUSTRALIAN GENBANK
AUSTRALIAN GENBANK
AUSTRALIAN GENBANK
AUSTRALIAN GENBANK
AUSTRALIAN GENBANK
AUSTRALIAN GENBANK
AUSTRALIAN GENBANK
AUSTRALIAN GENBANK
AUSTRALIAN GENBANK
AUSTRALIAN GENBANK
AUSTRALIAN GENBANK
AUSTRALIAN GENBANK
AUSTRALIAN GENBANK
AUSTRALIAN GENBANK
AUSTRALIAN GENBANK
AUSTRALIAN GENBANK
AUSTRALIAN GENBANK
AUSTRALIAN GENBANK
AUSTRALIAN GENBANK
AUSTRALIAN GENBANK
AUSTRALIAN GENBANK
AUSTRALIAN GENBANK
AUSTRALIAN GENBANK
AUSTRALIAN GENBANK
AUSTRALIAN GENBANK
AUSTRALIAN GENBANK
AUSTRALIAN GENBANK
AUSTRALIAN GENBANK
AUSTRALIAN GENBANK
AUSTRALIAN GENBANK
AUSTRALIAN GENBANK
AUSTRALIAN GENBANK
AUSTRALIAN GENBANK
AUSTRALIAN GENBANK
AUSTRALIAN GENBANK
AUSTRALIAN GENBANK
AUSTRALIAN GENBANK
AUSTRALIAN GENBANK
AUSTRALIAN GENBANK
AUSTRALIAN GENBANK

AUSTRALIAN GENBANK
AUSTRALIAN GENBANK
AUSTRALIAN GENBANK
AUSTRALIAN GENBANK
AUSTRALIAN GENBANK
AUSTRALIAN GENBANK
AUSTRALIAN GENBANK
AUSTRALIAN GENBANK
AUSTRALIAN GENBANK
AUSTRALIAN GENBANK
AUSTRALIAN GENBANK
AUSTRALIAN GENBANK
AUSTRALIAN GENBANK
AUSTRALIAN GENBANK
AUSTRALIAN GENBANK
AUSTRALIAN GENBANK
AUSTRALIAN GENBANK
AUSTRALIAN GENBANK
AUSTRALIAN GENBANK
AUSTRALIAN GENBANK
AUSTRALIAN GENBANK
AUSTRALIAN GENBANK
AUSTRALIAN GENBANK
AUSTRALIAN GENBANK
AUSTRALIAN GENBANK
AUSTRALIAN GENBANK
AUSTRALIAN GENBANK
AUSTRALIAN GENBANK
AUSTRALIAN GENBANK
AUSTRALIAN GENBANK
AUSTRALIAN GENBANK
AUSTRALIAN GENBANK
AUSTRALIAN GENBANK
AUSTRALIAN GENBANK
AUSTRALIAN GENBANK
AUSTRALIAN GENBANK
AUSTRALIAN GENBANK
AUSTRALIAN GENBANK
AUSTRALIAN GENBANK
AUSTRALIAN GENBANK
AUSTRALIAN GENBANK
AUSTRALIAN GENBANK
AUSTRALIAN GENBANK
AUSTRALIAN GENBANK
AUSTRALIAN GENBANK
AUSTRALIAN GENBANK
AUSTRALIAN GENBANK
AUSTRALIAN GENBANK
AUSTRALIAN GENBANK
AUSTRALIAN GENBANK
AUSTRALIAN GENBANK
AUSTRALIAN GENBANK

AUSTRALIAN GENBANK
AUSTRALIAN GENBANK
AUSTRALIAN GENBANK
AUSTRALIAN GENBANK
AUSTRALIAN GENBANK
AUSTRALIAN GENBANK
AUSTRALIAN GENBANK
AUSTRALIAN GENBANK
AUSTRALIAN GENBANK
AUSTRALIAN GENBANK
AUSTRALIAN GENBANK
AUSTRALIAN GENBANK
AUSTRALIAN GENBANK
AUSTRALIAN GENBANK
AUSTRALIAN GENBANK
AUSTRALIAN GENBANK
AUSTRALIAN GENBANK
AUSTRALIAN GENBANK
AUSTRALIAN GENBANK
AUSTRALIAN GENBANK
AUSTRALIAN GENBANK
AUSTRALIAN GENBANK
AUSTRALIAN GENBANK
AUSTRALIAN GENBANK
AUSTRALIAN GENBANK
AUSTRALIAN GENBANK
AUSTRALIAN GENBANK
AUSTRALIAN GENBANK
AUSTRALIAN GENBANK
AUSTRALIAN GENBANK
AUSTRALIAN GENBANK
AUSTRALIAN GENBANK
AUSTRALIAN GENBANK
AUSTRALIAN GENBANK
AUSTRALIAN GENBANK
AUSTRALIAN GENBANK
AUSTRALIAN GENBANK
AUSTRALIAN GENBANK
AUSTRALIAN GENBANK
AUSTRALIAN GENBANK
AUSTRALIAN GENBANK
AUSTRALIAN GENBANK
AUSTRALIAN GENBANK
AUSTRALIAN GENBANK
AUSTRALIAN GENBANK
AUSTRALIAN GENBANK
AUSTRALIAN GENBANK
AUSTRALIAN GENBANK
AUSTRALIAN GENBANK
AUSTRALIAN GENBANK
AUSTRALIAN GENBANK
AUSTRALIAN GENBANK

AUSTRALIAN GENBANK
AUSTRALIAN GENBANK
AUSTRALIAN GENBANK
AUSTRALIAN GENBANK
AUSTRALIAN GENBANK
AUSTRALIAN GENBANK
AUSTRALIAN GENBANK
AUSTRALIAN GENBANK
AUSTRALIAN GENBANK
AUSTRALIAN GENBANK
AUSTRALIAN GENBANK
AUSTRALIAN GENBANK
AUSTRALIAN GENBANK
AUSTRALIAN GENBANK
AUSTRALIAN GENBANK
AUSTRALIAN GENBANK
AUSTRALIAN GENBANK
AUSTRALIAN GENBANK
AUSTRALIAN GENBANK
AUSTRALIAN GENBANK
AUSTRALIAN GENBANK
AUSTRALIAN GENBANK
AUSTRALIAN GENBANK
AUSTRALIAN GENBANK
AUSTRALIAN GENBANK
AUSTRALIAN GENBANK
AUSTRALIAN GENBANK
AUSTRALIAN GENBANK
AUSTRALIAN GENBANK
AUSTRALIAN GENBANK
AUSTRALIAN GENBANK
AUSTRALIAN GENBANK
AUSTRALIAN GENBANK
AUSTRALIAN GENBANK
AUSTRALIAN GENBANK
AUSTRALIAN GENBANK
AUSTRALIAN GENBANK
AUSTRALIAN GENBANK
AUSTRALIAN GENBANK
AUSTRALIAN GENBANK
AUSTRALIAN GENBANK
AUSTRALIAN GENBANK
AUSTRALIAN GENBANK
AUSTRALIAN GENBANK
P15-BRGV Suceava
P15-BRGV Suceava
P15-BRGV Suceava
P15-BRGV Suceava
P15-BRGV Suceava
P15-BRGV Suceava
P15-BRGV Suceava
P15-BRGV Suceava

| | |
|---|---|
| P15-BRGV | Suceava |
| P15-BRGV | Suceava |
| P15-BRGV | Suceava |
| P15-BRGV | Suceava |
| P15-BRGV | Suceava |
| P15-BRGV | Suceava |
| P15-BRGV | Suceava |
| P15-BRGV | Suceava |
| P15-BRGV | Suceava |
| P15-BRGV | Suceava |
| P15-BRGV | Suceava |
| P15-BRGV | Suceava |
| P15-BRGV | Suceava |
| P15-BRGV | Suceava |
| P15-BRGV | Suceava |
| P19-ISEA | |
| P19-ISEA | |
| P19-ISEA | |
| P14-SCDL-BACAU | |
| P14-SCDL-BACAU | |
| P14-SCDL-BACAU | |
| P14-SCDL-BACAU | |
| P14-SCDL-BACAU | |
| P14-SCDL-BACAU | |
| P14-SCDL-BACAU | |
| P14-SCDL-BACAU | |
| P16-INIA | |
| P16-INIA | |
| P16-INIA | |
| P16-INIA | |
| P16-INIA | |
| P16-INIA | |
| P16-INIA | |
| P16-INIA | |
| P16-INIA | |
| P16-INIA | |
| P16-INIA | |
| P16-INIA | |
| P16-INIA | |
| P16-INIA | |
| P16-INIA | |
| P16-INIA | |
| P16-INIA | |
| P16-INIA | |
| P16-INIA | |
| P16-INIA | |
| P16-INIA | |
| P16-INIA | |
| P16-INIA | |
| P16-INIA | |
| P16-INIA | |
| P16-INIA | |

P16-INIA
P16-INIA
P16-INIA
P16-INIA
P16-INIA
P16-INIA
P16-INIA
P16-INIA
P16-INIA
P16-INIA
P16-INIA
P16-INIA
P16-INIA
P16-INIA
P16-INIA
P16-INIA
P16-INIA
P16-INIA
P16-INIA
P16-INIA
P16-INIA
P16-INIA
P16-INIA
P16-INIA
P16-INIA
P16-INIA
P16-INIA
P16-INIA
P16-INIA
P16-INIA
P16-INIA
P16-INIA
P16-INIA
P16-INIA
P16-INIA
P16-INIA
P16-INIA
P16-INIA
P16-INIA
P16-INIA
P16-INIA
P16-INIA
P16-INIA
P16-INIA
P16-INIA
P16-INIA
P16-INIA
P16-INIA
P16-INIA
P16-INIA
P16-INIA
P16-INIA

P16-INIA
P16-INIA
P16-INIA
P16-INIA
P16-INIA
P16-INIA
P16-INIA
P16-INIA
P16-INIA
P16-INIA
P16-INIA
P16-INIA
P16-INIA
P16-INIA
P16-INIA
P16-INIA
P16-INIA
P16-INIA
P16-INIA
P16-INIA
P16-INIA
P16-INIA
P16-INIA
P16-INIA
P16-INIA
P16-INIA
P16-INIA
P16-INIA
P16-INIA
P16-INIA
P16-INIA
P16-INIA
P16-INIA
P16-INIA
P16-INIA
P16-INIA
P16-INIA
P16-INIA
P16-INIA
P16-INIA
P16-INIA
P16-INIA
P16-INIA
P16-INIA
P16-INIA
P16-INIA
P16-INIA
P16-INIA
P16-INIA
P16-INIA
P16-INIA
P16-INIA

P16-INIA
P16-INIA
P16-INIA
P16-INIA
P16-INIA
P16-INIA
P16-INIA
P16-INIA
P16-INIA
P16-INIA
P16-INIA
P16-INIA
P16-INIA
P16-INIA
P16-INIA
P16-INIA
P16-INIA
P16-INIA
P16-INIA
P16-INIA
P16-INIA
P16-INIA
P16-INIA
P16-INIA
P16-INIA
P16-INIA
P16-INIA
P16-INIA
P16-INIA
P16-INIA
P16-INIA
P16-INIA
P16-INIA
P16-INIA
P16-INIA
P16-INIA
P16-INIA
P16-INIA
P16-INIA
P16-INIA
P16-INIA
P16-INIA
P16-INIA
P16-INIA
P16-INIA
P16-INIA
P16-INIA
P16-INIA
P16-INIA
P16-INIA
P16-INIA
P16-INIA

P16-INIA
P16-INIA
P16-INIA
P16-INIA
P16-INIA
P16-INIA
P16-INIA
P16-INIA
P16-INIA
P16-INIA
P16-INIA
P16-INIA
P16-INIA
P16-INIA
P16-INIA
P16-INIA
P16-INIA
P16-INIA
P16-INIA
P16-INIA
P16-INIA
P16-INIA
P16-INIA
P16-INIA
P16-INIA
P16-INIA
P16-INIA
P16-INIA
P16-INIA
P16-INIA
P16-INIA
P16-INIA
P16-INIA
P16-INIA
P16-INIA
P16-INIA
P16-INIA
P16-INIA
P16-INIA
P16-INIA
P16-INIA
P16-INIA
P16-INIA
P16-INIA
P16-INIA
P16-INIA
P16-INIA
P16-INIA
P16-INIA
P16-INIA
P16-INIA
P16-INIA

P16-INIA
P16-INIA
P16-INIA
P16-INIA
P16-INIA
P16-INIA
P16-INIA
P16-INIA
P16-INIA
P16-INIA
P16-INIA
P16-INIA
P16-INIA
P16-INIA
P16-INIA
P16-INIA
P16-INIA
P16-INIA
P16-INIA
P16-INIA
P16-INIA
P16-INIA
P16-INIA
P16-INIA
P16-INIA
P16-INIA
P16-INIA
P16-INIA
P16-INIA
P16-INIA
P16-INIA
P16-INIA
P16-INIA
P16-INIA
P16-INIA
P16-INIA
P16-INIA
P16-INIA
P16-INIA
P16-INIA
P16-INIA
P16-INIA
P16-INIA
P16-INIA
P16-INIA
P16-INIA
P16-INIA
P16-INIA
P16-INIA
P16-INIA
P16-INIA
P16-INIA

P16-INIA
P16-INIA
P16-INIA
P16-INIA
P16-INIA
P16-INIA
P16-INIA
P16-INIA
P16-INIA
P16-INIA
P16-INIA
P16-INIA
P16-INIA
P16-INIA
P16-INIA
P16-INIA
P16-INIA
P16-INIA
P16-INIA
P16-INIA
P16-INIA
P16-INIA
P16-INIA
P16-INIA
P16-INIA
P16-INIA
P16-INIA
P16-INIA
P16-INIA
P16-INIA
P16-INIA
P16-INIA
P16-INIA
P16-INIA
P16-INIA
P16-INIA
P16-INIA
P16-INIA
P16-INIA
P16-INIA
P16-INIA
P16-INIA
P16-INIA
P16-INIA
P16-INIA
P16-INIA
P16-INIA
P16-INIA
P16-INIA
P16-INIA
P16-INIA
P16-INIA

P16-INIA
P16-INIA
P16-INIA
P16-INIA
P16-INIA
P16-INIA
P16-INIA
P16-INIA
P16-INIA
P16-INIA
P16-INIA
P16-INIA
P16-INIA
P16-INIA
P16-INIA
P16-INIA
P16-INIA
P16-INIA
P16-INIA
P16-INIA
P16-INIA
P16-INIA
P16-INIA
P16-INIA
P16-INIA
P16-INIA
P16-INIA
P16-INIA
P16-INIA
P16-INIA
P16-INIA
P16-INIA
P16-INIA
P16-INIA
P16-INIA
P16-INIA
P16-INIA
P16-INIA
P16-INIA
P16-INIA
P16-INIA
P16-INIA
P16-INIA
P16-INIA
P16-INIA
P16-INIA
P16-INIA
P16-INIA
P16-INIA
P16-INIA
P16-INIA
P16-INIA

P16-INIA
P16-INIA
P16-INIA
P16-INIA
P16-INIA
P16-INIA
P16-INIA
P16-INIA
P16-INIA
P16-INIA
P16-INIA
P16-INIA
P16-INIA
P16-INIA
P16-INIA
P16-INIA
P16-INIA
P16-INIA
P16-INIA
P16-INIA
P16-INIA
P16-INIA
P16-INIA
P16-INIA
P16-INIA
P16-INIA
P16-INIA
P16-INIA
P16-INIA
P16-INIA
P16-INIA
P16-INIA
P16-INIA
P16-INIA
P16-INIA
P16-INIA
P16-INIA
P16-INIA
P16-INIA
P16-INIA
P16-INIA
P16-INIA
P16-INIA
P16-INIA
P16-INIA
P16-INIA
P16-INIA
P16-INIA
P16-INIA
P16-INIA
P16-INIA
P16-INIA

P16-INIA
P16-INIA
P16-INIA
P16-INIA
P16-INIA
P16-INIA
P16-INIA
P16-INIA
P16-INIA
P16-INIA
P16-INIA
P16-INIA
P16-INIA
P16-INIA
P16-INIA
P16-INIA
P16-INIA
P16-INIA
P16-INIA
P16-INIA
P16-INIA
P16-INIA
P16-INIA
P16-INIA
P16-INIA
P16-INIA
P16-INIA
P16-INIA
P16-INIA
P16-INIA
P16-INIA
P16-INIA
P16-INIA
P16-INIA
P16-INIA
P16-INIA
P16-INIA
P16-INIA
P16-INIA
P16-INIA
P16-INIA
P16-INIA
P16-INIA
P16-INIA
P16-INIA
P16-INIA
P16-INIA
P16-INIA
P16-INIA
P16-INIA
P16-INIA
P16-INIA

P16-INIA
P16-INIA
P16-INIA
P16-INIA
P16-INIA
P16-INIA
P16-INIA
P16-INIA
P16-INIA
P16-INIA
P16-INIA
P16-INIA
P16-INIA
P16-INIA
P16-INIA
P16-INIA
P16-INIA
P16-INIA
P16-INIA
P16-INIA
P16-INIA
P16-INIA
P16-INIA
P16-INIA
P16-INIA
P16-INIA
P16-INIA
P16-INIA
P16-INIA
P16-INIA
P16-INIA
P16-INIA
P16-INIA
P16-INIA
P16-INIA
P16-INIA
P16-INIA
P16-INIA
P16-INIA
P16-INIA
P16-INIA
P16-INIA
P16-INIA
P16-INIA
P16-INIA
P16-INIA
P16-INIA
P16-INIA
P16-INIA
P16-INIA
P16-INIA
P16-INIA

P16-INIA
P16-INIA
P16-INIA
P16-INIA
P16-INIA
P16-INIA
P16-INIA
P16-INIA
P16-INIA
P16-INIA
P16-INIA
P16-INIA
P16-INIA
P16-INIA
P16-INIA
P16-INIA
P16-INIA
P16-INIA
P16-INIA
P16-INIA
P16-INIA
P16-INIA
P16-INIA
P16-INIA
P16-INIA
P16-INIA
P16-INIA
P16-INIA
P16-INIA
P16-INIA
P16-INIA
P16-INIA
P16-INIA
P16-INIA
P16-INIA
P16-INIA
P16-INIA
P16-INIA
P16-INIA
P16-INIA
P16-INIA
P16-INIA
P16-INIA
P16-INIA
P16-INIA
P16-INIA
P16-INIA
P16-INIA
P16-INIA
P16-INIA
P16-INIA
P16-INIA

P16-INIA
P16-INIA
P16-INIA
P16-INIA
P16-INIA
P16-INIA
P16-INIA
P16-INIA
P16-INIA
P16-INIA
P16-INIA
P16-INIA
P16-INIA
P16-INIA
P16-INIA
P16-INIA
P16-INIA
P16-INIA
P16-INIA
P16-INIA
P16-INIA
P16-INIA
P16-INIA
P16-INIA
P16-INIA
P16-INIA
P16-INIA
P16-INIA
P16-INIA
P16-INIA
P16-INIA
P16-INIA
P16-INIA
P16-INIA
P16-INIA
P16-INIA
P16-INIA
P16-INIA
P16-INIA
P16-INIA
P16-INIA
P16-INIA
P16-INIA
P16-INIA
P16-INIA
P16-INIA
P16-INIA
P16-INIA
P16-INIA
P16-INIA
P16-INIA
P16-INIA

P16-INIA
P16-INIA
P16-INIA
P16-INIA
P16-INIA
P16-INIA
P16-INIA
P16-INIA
P16-INIA
P16-INIA
P16-INIA
P16-INIA
P16-INIA
P16-INIA
P16-INIA
P16-INIA
P16-INIA
P16-INIA
P16-INIA
P16-INIA
P16-INIA
P16-INIA
P16-INIA
P16-INIA
P16-INIA
P16-INIA
P16-INIA
P16-INIA
P16-INIA
P16-INIA
P16-INIA
P16-INIA
P16-INIA
P16-INIA
P16-INIA
P16-INIA
P16-INIA
P16-INIA
P16-INIA
P16-INIA
P16-INIA
P16-INIA
P16-INIA
P16-INIA
P16-INIA
P16-INIA
P16-INIA
P16-INIA
P16-INIA
P16-INIA
P16-INIA
P16-INIA

P16-INIA
P16-INIA
P16-INIA
P16-INIA
P16-INIA
P16-INIA
P16-INIA
P16-INIA
P16-INIA
P16-INIA
P16-INIA
P16-INIA
P16-INIA
P16-INIA
P16-INIA
P16-INIA
P16-INIA
P16-INIA
P16-INIA
P16-INIA
P16-INIA
P16-INIA
P16-INIA
P16-INIA
P16-INIA
P16-INIA
P16-INIA
P16-INIA
P16-INIA
P16-INIA
P16-INIA
P16-INIA
P16-INIA
P16-INIA
P16-INIA
P16-INIA
P16-INIA
P16-INIA
P16-INIA
P16-INIA
USDA
USDA
USDA
USDA
USDA
USDA
USDA
USDA
USDA
USDA
USDA
USDA

USDA
USDA
USDA
USDA
USDA
USDA
USDA
USDA
USDA
USDA
USDA
USDA
USDA
USDA
USDA
USDA
USDA
USDA
USDA
USDA
USDA
USDA
USDA
USDA
USDA
USDA
USDA
USDA
USDA
USDA
USDA
USDA
USDA
USDA
USDA
USDA
USDA
USDA
USDA
USDA
USDA
USDA
USDA
USDA
USDA
USDA
USDA
USDA
USDA
USDA
USDA
USDA

USDA
USDA
USDA
USDA
USDA
USDA
USDA
USDA
USDA
USDA
USDA
USDA
USDA
USDA
USDA
USDA
USDA
USDA
USDA
USDA
USDA
USDA
USDA
USDA
USDA
USDA
USDA
USDA
USDA
USDA
USDA
USDA
USDA
USDA
USDA
USDA
USDA
USDA
USDA
USDA
USDA
USDA
USDA
USDA
USDA
USDA
USDA
USDA
USDA
USDA
USDA
USDA

USDA
USDA
USDA
USDA
USDA
USDA
USDA
USDA
USDA
USDA
USDA
USDA
USDA
USDA
USDA
USDA
USDA
USDA
USDA
USDA
USDA
USDA
USDA
USDA
USDA
USDA
USDA
USDA
USDA
USDA
USDA
USDA
USDA
USDA
USDA
USDA
USDA
USDA
USDA
USDA
USDA
USDA
USDA
USDA
USDA
USDA
USDA
USDA
USDA
USDA
USDA
USDA

USDA
USDA
USDA
USDA
USDA
USDA
USDA
USDA
USDA
USDA
USDA
USDA
USDA
USDA
USDA
USDA
USDA
USDA
USDA
USDA
USDA
USDA
USDA
USDA
USDA
USDA
USDA
USDA
USDA
USDA
USDA
USDA
USDA
USDA
USDA
USDA
USDA
Jouffray-Drillaud
Jouffray-Drillaud
Jouffray-Drillaud
Jouffray-Drillaud
Jouffray-Drillaud
Jouffray-Drillaud
Jouffray-Drillaud
Jouffray-Drillaud
Jouffray-Drillaud
Jouffray-Drillaud
Jouffray-Drillaud
Jouffray-Drillaud
Jouffray-Drillaud
Jouffray-Drillaud
P8-CREA

P8-CREA
Private donor

| Source |
| --- |
| https://www.ecpgr.cgiar.org/resources/germplasm-databases/list-of-germplasm-databases/crop-databases/crop-database-windows/lupin |
| https://www.ecpgr.cgiar.org/resources/germplasm-databases/list-of-germplasm-databases/crop-databases/crop-database-windows/lupin |
| https://www.ecpgr.cgiar.org/resources/germplasm-databases/list-of-germplasm-databases/crop-databases/crop-database-windows/lupin |
| https://www.ecpgr.cgiar.org/resources/germplasm-databases/list-of-germplasm-databases/crop-databases/crop-database-windows/lupin |
| https://www.ecpgr.cgiar.org/resources/germplasm-databases/list-of-germplasm-databases/crop-databases/crop-database-windows/lupin |
| https://www.ecpgr.cgiar.org/resources/germplasm-databases/list-of-germplasm-databases/crop-databases/crop-database-windows/lupin |
| https://www.ecpgr.cgiar.org/resources/germplasm-databases/list-of-germplasm-databases/crop-databases/crop-database-windows/lupin |
| https://www.ecpgr.cgiar.org/resources/germplasm-databases/list-of-germplasm-databases/crop-databases/crop-database-windows/lupin |
| https://www.ecpgr.cgiar.org/resources/germplasm-databases/list-of-germplasm-databases/crop-databases/crop-database-windows/lupin |
| https://www.ecpgr.cgiar.org/resources/germplasm-databases/list-of-germplasm-databases/crop-databases/crop-database-windows/lupin |
| https://www.ecpgr.cgiar.org/resources/germplasm-databases/list-of-germplasm-databases/crop-databases/crop-database-windows/lupin |
| https://www.ecpgr.cgiar.org/resources/germplasm-databases/list-of-germplasm-databases/crop-databases/crop-database-windows/lupin |
| https://www.ecpgr.cgiar.org/resources/germplasm-databases/list-of-germplasm-databases/crop-databases/crop-database-windows/lupin |
| https://www.ecpgr.cgiar.org/resources/germplasm-databases/list-of-germplasm-databases/crop-databases/crop-database-windows/lupin |
| https://www.ecpgr.cgiar.org/resources/germplasm-databases/list-of-germplasm-databases/crop-databases/crop-database-windows/lupin |
| https://www.ecpgr.cgiar.org/resources/germplasm-databases/list-of-germplasm-databases/crop-databases/crop-database-windows/lupin |
| https://www.ecpgr.cgiar.org/resources/germplasm-databases/list-of-germplasm-databases/crop-databases/crop-database-windows/lupin |
| https://www.ecpgr.cgiar.org/resources/germplasm-databases/list-of-germplasm-databases/crop-databases/crop-database-windows/lupin |
| https://www.ecpgr.cgiar.org/resources/germplasm-databases/list-of-germplasm-databases/crop-databases/crop-database-windows/lupin |
| https://www.ecpgr.cgiar.org/resources/germplasm-databases/list-of-germplasm-databases/crop-databases/crop-database-windows/lupin |
| https://www.ecpgr.cgiar.org/resources/germplasm-databases/list-of-germplasm-databases/crop-databases/crop-database-windows/lupin |
| https://www.ecpgr.cgiar.org/resources/germplasm-databases/list-of-germplasm-databases/crop-databases/crop-database-windows/lupin |
| https://www.ecpgr.cgiar.org/resources/germplasm-databases/list-of-germplasm-databases/crop-databases/crop-database-windows/lupin |
| https://www.ecpgr.cgiar.org/resources/germplasm-databases/list-of-germplasm-databases/crop-databases/crop-database-windows/lupin |
| https://www.ecpgr.cgiar.org/resources/germplasm-databases/list-of-germplasm-databases/crop-databases/crop-database-windows/lupin |
| https://www.ecpgr.cgiar.org/resources/germplasm-databases/list-of-germplasm-databases/crop-databases/crop-database-windows/lupin |
| https://www.ecpgr.cgiar.org/resources/germplasm-databases/list-of-germplasm-databases/crop-databases/crop-database-windows/lupin |
| https://www.ecpgr.cgiar.org/resources/germplasm-databases/list-of-germplasm-databases/crop-databases/crop-database-windows/lupin |
| https://www.ecpgr.cgiar.org/resources/germplasm-databases/list-of-germplasm-databases/crop-databases/crop-database-windows/lupin |
| https://www.ecpgr.cgiar.org/resources/germplasm-databases/list-of-germplasm-databases/crop-databases/crop-database-windows/lupin |
| https://www.ecpgr.cgiar.org/resources/germplasm-databases/list-of-germplasm-databases/crop-databases/crop-database-windows/lupin |
| https://www.ecpgr.cgiar.org/resources/germplasm-databases/list-of-germplasm-databases/crop-databases/crop-database-windows/lupin |
| https://www.ecpgr.cgiar.org/resources/germplasm-databases/list-of-germplasm-databases/crop-databases/crop-database-windows/lupin |
| https://www.ecpgr.cgiar.org/resources/germplasm-databases/list-of-germplasm-databases/crop-databases/crop-database-windows/lupin |
| https://www.ecpgr.cgiar.org/resources/germplasm-databases/list-of-germplasm-databases/crop-databases/crop-database-windows/lupin |
| https://www.ecpgr.cgiar.org/resources/germplasm-databases/list-of-germplasm-databases/crop-databases/crop-database-windows/lupin |
| https://www.ecpgr.cgiar.org/resources/germplasm-databases/list-of-germplasm-databases/crop-databases/crop-database-windows/lupin |
| https://www.ecpgr.cgiar.org/resources/germplasm-databases/list-of-germplasm-databases/crop-databases/crop-database-windows/lupin |
| https://www.ecpgr.cgiar.org/resources/germplasm-databases/list-of-germplasm-databases/crop-databases/crop-database-windows/lupin |
| https://www.ecpgr.cgiar.org/resources/germplasm-databases/list-of-germplasm-databases/crop-databases/crop-database-windows/lupin |
| https://www.ecpgr.cgiar.org/resources/germplasm-databases/list-of-germplasm-databases/crop-databases/crop-database-windows/lupin |
| https://www.ecpgr.cgiar.org/resources/germplasm-databases/list-of-germplasm-databases/crop-databases/crop-database-windows/lupin |
| https://www.ecpgr.cgiar.org/resources/germplasm-databases/list-of-germplasm-databases/crop-databases/crop-database-windows/lupin |
| https://www.ecpgr.cgiar.org/resources/germplasm-databases/list-of-germplasm-databases/crop-databases/crop-database-windows/lupin |
| https://www.ecpgr.cgiar.org/resources/germplasm-databases/list-of-germplasm-databases/crop-databases/crop-database-windows/lupin |
| https://www.ecpgr.cgiar.org/resources/germplasm-databases/list-of-germplasm-databases/crop-databases/crop-database-windows/lupin |
| https://www.ecpgr.cgiar.org/resources/germplasm-databases/list-of-germplasm-databases/crop-databases/crop-database-windows/lupin |
| https://www.ecpgr.cgiar.org/resources/germplasm-databases/list-of-germplasm-databases/crop-databases/crop-database-windows/lupin |
| https://www.ecpgr.cgiar.org/resources/germplasm-databases/list-of-germplasm-databases/crop-databases/crop-database-windows/lupin |
| https://www.ecpgr.cgiar.org/resources/germplasm-databases/list-of-germplasm-databases/crop-databases/crop-database-windows/lupin |

https://www.ecpgr.cgiar.org/resources/germplasm-databases/list-of-germplasm-databases/crop-databases/crop-database-windows/lupin
https://www.ecpgr.cgiar.org/resources/germplasm-databases/list-of-germplasm-databases/crop-databases/crop-database-windows/lupin
https://www.ecpgr.cgiar.org/resources/germplasm-databases/list-of-germplasm-databases/crop-databases/crop-database-windows/lupin
https://www.ecpgr.cgiar.org/resources/germplasm-databases/list-of-germplasm-databases/crop-databases/crop-database-windows/lupin
https://www.ecpgr.cgiar.org/resources/germplasm-databases/list-of-germplasm-databases/crop-databases/crop-database-windows/lupin
https://www.ecpgr.cgiar.org/resources/germplasm-databases/list-of-germplasm-databases/crop-databases/crop-database-windows/lupin
https://www.ecpgr.cgiar.org/resources/germplasm-databases/list-of-germplasm-databases/crop-databases/crop-database-windows/lupin
https://www.ecpgr.cgiar.org/resources/germplasm-databases/list-of-germplasm-databases/crop-databases/crop-database-windows/lupin
https://www.ecpgr.cgiar.org/resources/germplasm-databases/list-of-germplasm-databases/crop-databases/crop-database-windows/lupin
https://www.ecpgr.cgiar.org/resources/germplasm-databases/list-of-germplasm-databases/crop-databases/crop-database-windows/lupin
https://www.ecpgr.cgiar.org/resources/germplasm-databases/list-of-germplasm-databases/crop-databases/crop-database-windows/lupin
https://www.ecpgr.cgiar.org/resources/germplasm-databases/list-of-germplasm-databases/crop-databases/crop-database-windows/lupin
https://www.ecpgr.cgiar.org/resources/germplasm-databases/list-of-germplasm-databases/crop-databases/crop-database-windows/lupin
https://www.ecpgr.cgiar.org/resources/germplasm-databases/list-of-germplasm-databases/crop-databases/crop-database-windows/lupin
https://www.ecpgr.cgiar.org/resources/germplasm-databases/list-of-germplasm-databases/crop-databases/crop-database-windows/lupin
https://www.ecpgr.cgiar.org/resources/germplasm-databases/list-of-germplasm-databases/crop-databases/crop-database-windows/lupin
https://www.ecpgr.cgiar.org/resources/germplasm-databases/list-of-germplasm-databases/crop-databases/crop-database-windows/lupin
https://www.genesys-pgr.org/a/c30728db-7632-4457-84bd-c3197be87ba4
https://www.ecpgr.cgiar.org/resources/germplasm-databases/list-of-germplasm-databases/crop-databases/crop-database-windows/lupin
https://www.ecpgr.cgiar.org/resources/germplasm-databases/list-of-germplasm-databases/crop-databases/crop-database-windows/lupin
https://www.ecpgr.cgiar.org/resources/germplasm-databases/list-of-germplasm-databases/crop-databases/crop-database-windows/lupin
https://www.ecpgr.cgiar.org/resources/germplasm-databases/list-of-germplasm-databases/crop-databases/crop-database-windows/lupin
https://www.ecpgr.cgiar.org/resources/germplasm-databases/list-of-germplasm-databases/crop-databases/crop-database-windows/lupin
https://www.ecpgr.cgiar.org/resources/germplasm-databases/list-of-germplasm-databases/crop-databases/crop-database-windows/lupin
https://www.ecpgr.cgiar.org/resources/germplasm-databases/list-of-germplasm-databases/crop-databases/crop-database-windows/lupin
https://www.ecpgr.cgiar.org/resources/germplasm-databases/list-of-germplasm-databases/crop-databases/crop-database-windows/lupin
https://www.ecpgr.cgiar.org/resources/germplasm-databases/list-of-germplasm-databases/crop-databases/crop-database-windows/lupin
https://www.genesys-pgr.org/a/d11766ef-4b1c-4887-ad9e-3d404aad11b1
https://www.ecpgr.cgiar.org/resources/germplasm-databases/list-of-germplasm-databases/crop-databases/crop-database-windows/lupin
https://www.ecpgr.cgiar.org/resources/germplasm-databases/list-of-germplasm-databases/crop-databases/crop-database-windows/lupin
https://www.ecpgr.cgiar.org/resources/germplasm-databases/list-of-germplasm-databases/crop-databases/crop-database-windows/lupin
https://www.ecpgr.cgiar.org/resources/germplasm-databases/list-of-germplasm-databases/crop-databases/crop-database-windows/lupin
https://www.ecpgr.cgiar.org/resources/germplasm-databases/list-of-germplasm-databases/crop-databases/crop-database-windows/lupin
https://www.ecpgr.cgiar.org/resources/germplasm-databases/list-of-germplasm-databases/crop-databases/crop-database-windows/lupin
https://www.ecpgr.cgiar.org/resources/germplasm-databases/list-of-germplasm-databases/crop-databases/crop-database-windows/lupin
https://www.ecpgr.cgiar.org/resources/germplasm-databases/list-of-germplasm-databases/crop-databases/crop-database-windows/lupin
https://www.ecpgr.cgiar.org/resources/germplasm-databases/list-of-germplasm-databases/crop-databases/crop-database-windows/lupin
https://www.ecpgr.cgiar.org/resources/germplasm-databases/list-of-germplasm-databases/crop-databases/crop-database-windows/lupin
https://www.ecpgr.cgiar.org/resources/germplasm-databases/list-of-germplasm-databases/crop-databases/crop-database-windows/lupin
https://www.ecpgr.cgiar.org/resources/germplasm-databases/list-of-germplasm-databases/crop-databases/crop-database-windows/lupin
https://www.ecpgr.cgiar.org/resources/germplasm-databases/list-of-germplasm-databases/crop-databases/crop-database-windows/lupin
https://www.ecpgr.cgiar.org/resources/germplasm-databases/list-of-germplasm-databases/crop-databases/crop-database-windows/lupin
https://www.ecpgr.cgiar.org/resources/germplasm-databases/list-of-germplasm-databases/crop-databases/crop-database-windows/lupin
https://www.ecpgr.cgiar.org/resources/germplasm-databases/list-of-germplasm-databases/crop-databases/crop-database-windows/lupin
https://www.ecpgr.cgiar.org/resources/germplasm-databases/list-of-germplasm-databases/crop-databases/crop-database-windows/lupin
https://www.ecpgr.cgiar.org/resources/germplasm-databases/list-of-germplasm-databases/crop-databases/crop-database-windows/lupin
https://www.ecpgr.cgiar.org/resources/germplasm-databases/list-of-germplasm-databases/crop-databases/crop-database-windows/lupin
https://www.genesys-pgr.org/a/bf13afa6-a0bc-4d42-8a30-d64329cd3658
https://www.ecpgr.cgiar.org/resources/germplasm-databases/list-of-germplasm-databases/crop-databases/crop-database-windows/lupin
https://www.genesys-pgr.org/a/999ed2c0-de33-40b2-99f3-baab3776a6c0
https://www.ecpgr.cgiar.org/resources/germplasm-databases/list-of-germplasm-databases/crop-databases/crop-database-windows/lupin
https://www.ecpgr.cgiar.org/resources/germplasm-databases/list-of-germplasm-databases/crop-databases/crop-database-windows/lupin

https://www.ecpgr.cgiar.org/resources/germplasm-databases/list-of-germplasm-databases/crop-databases/crop-database-windows/lupin
https://www.genesys-pgr.org/a/47276023-511c-4d8d-8378-76ee3cd64f3b
https://www.ecpgr.cgiar.org/resources/germplasm-databases/list-of-germplasm-databases/crop-databases/crop-database-windows/lupin
https://www.ecpgr.cgiar.org/resources/germplasm-databases/list-of-germplasm-databases/crop-databases/crop-database-windows/lupin
https://www.ecpgr.cgiar.org/resources/germplasm-databases/list-of-germplasm-databases/crop-databases/crop-database-windows/lupin
https://www.ecpgr.cgiar.org/resources/germplasm-databases/list-of-germplasm-databases/crop-databases/crop-database-windows/lupin
https://www.ecpgr.cgiar.org/resources/germplasm-databases/list-of-germplasm-databases/crop-databases/crop-database-windows/lupin
https://www.ecpgr.cgiar.org/resources/germplasm-databases/list-of-germplasm-databases/crop-databases/crop-database-windows/lupin
https://www.ecpgr.cgiar.org/resources/germplasm-databases/list-of-germplasm-databases/crop-databases/crop-database-windows/lupin
https://www.ecpgr.cgiar.org/resources/germplasm-databases/list-of-germplasm-databases/crop-databases/crop-database-windows/lupin
https://www.ecpgr.cgiar.org/resources/germplasm-databases/list-of-germplasm-databases/crop-databases/crop-database-windows/lupin
https://www.ecpgr.cgiar.org/resources/germplasm-databases/list-of-germplasm-databases/crop-databases/crop-database-windows/lupin
https://www.ecpgr.cgiar.org/resources/germplasm-databases/list-of-germplasm-databases/crop-databases/crop-database-windows/lupin
https://www.ecpgr.cgiar.org/resources/germplasm-databases/list-of-germplasm-databases/crop-databases/crop-database-windows/lupin
https://www.ecpgr.cgiar.org/resources/germplasm-databases/list-of-germplasm-databases/crop-databases/crop-database-windows/lupin
https://www.ecpgr.cgiar.org/resources/germplasm-databases/list-of-germplasm-databases/crop-databases/crop-database-windows/lupin
https://www.ecpgr.cgiar.org/resources/germplasm-databases/list-of-germplasm-databases/crop-databases/crop-database-windows/lupin
https://www.ecpgr.cgiar.org/resources/germplasm-databases/list-of-germplasm-databases/crop-databases/crop-database-windows/lupin
https://www.ecpgr.cgiar.org/resources/germplasm-databases/list-of-germplasm-databases/crop-databases/crop-database-windows/lupin
https://www.ecpgr.cgiar.org/resources/germplasm-databases/list-of-germplasm-databases/crop-databases/crop-database-windows/lupin
https://www.genesys-pgr.org/a/376c4110-f28e-4644-b2a9-fe9af9842759
https://www.ecpgr.cgiar.org/resources/germplasm-databases/list-of-germplasm-databases/crop-databases/crop-database-windows/lupin
https://www.ecpgr.cgiar.org/resources/germplasm-databases/list-of-germplasm-databases/crop-databases/crop-database-windows/lupin
https://eurisco.ipk-gatersleben.de/apex/f?p=103:16:::::P16_EURISCO_ACC_ID:565043
https://www.ecpgr.cgiar.org/resources/germplasm-databases/list-of-germplasm-databases/crop-databases/crop-database-windows/lupin
https://www.ecpgr.cgiar.org/resources/germplasm-databases/list-of-germplasm-databases/crop-databases/crop-database-windows/lupin
https://www.ecpgr.cgiar.org/resources/germplasm-databases/list-of-germplasm-databases/crop-databases/crop-database-windows/lupin
https://www.ecpgr.cgiar.org/resources/germplasm-databases/list-of-germplasm-databases/crop-databases/crop-database-windows/lupin
https://www.ecpgr.cgiar.org/resources/germplasm-databases/list-of-germplasm-databases/crop-databases/crop-database-windows/lupin
https://www.ecpgr.cgiar.org/resources/germplasm-databases/list-of-germplasm-databases/crop-databases/crop-database-windows/lupin
https://www.ecpgr.cgiar.org/resources/germplasm-databases/list-of-germplasm-databases/crop-databases/crop-database-windows/lupin
https://www.ecpgr.cgiar.org/resources/germplasm-databases/list-of-germplasm-databases/crop-databases/crop-database-windows/lupin
https://www.ecpgr.cgiar.org/resources/germplasm-databases/list-of-germplasm-databases/crop-databases/crop-database-windows/lupin
https://www.ecpgr.cgiar.org/resources/germplasm-databases/list-of-germplasm-databases/crop-databases/crop-database-windows/lupin
https://www.ecpgr.cgiar.org/resources/germplasm-databases/list-of-germplasm-databases/crop-databases/crop-database-windows/lupin
https://www.ecpgr.cgiar.org/resources/germplasm-databases/list-of-germplasm-databases/crop-databases/crop-database-windows/lupin
https://www.ecpgr.cgiar.org/resources/germplasm-databases/list-of-germplasm-databases/crop-databases/crop-database-windows/lupin
https://www.ecpgr.cgiar.org/resources/germplasm-databases/list-of-germplasm-databases/crop-databases/crop-database-windows/lupin
https://www.ecpgr.cgiar.org/resources/germplasm-databases/list-of-germplasm-databases/crop-databases/crop-database-windows/lupin
https://www.ecpgr.cgiar.org/resources/germplasm-databases/list-of-germplasm-databases/crop-databases/crop-database-windows/lupin
https://www.ecpgr.cgiar.org/resources/germplasm-databases/list-of-germplasm-databases/crop-databases/crop-database-windows/lupin
https://www.ecpgr.cgiar.org/resources/germplasm-databases/list-of-germplasm-databases/crop-databases/crop-database-windows/lupin
https://www.ecpgr.cgiar.org/resources/germplasm-databases/list-of-germplasm-databases/crop-databases/crop-database-windows/lupin
https://www.ecpgr.cgiar.org/resources/germplasm-databases/list-of-germplasm-databases/crop-databases/crop-database-windows/lupin
https://www.ecpgr.cgiar.org/resources/germplasm-databases/list-of-germplasm-databases/crop-databases/crop-database-windows/lupin
https://www.ecpgr.cgiar.org/resources/germplasm-databases/list-of-germplasm-databases/crop-databases/crop-database-windows/lupin
https://www.ecpgr.cgiar.org/resources/germplasm-databases/list-of-germplasm-databases/crop-databases/crop-database-windows/lupin
https://www.ecpgr.cgiar.org/resources/germplasm-databases/list-of-germplasm-databases/crop-databases/crop-database-windows/lupin
https://www.ecpgr.cgiar.org/resources/germplasm-databases/list-of-germplasm-databases/crop-databases/crop-database-windows/lupin
https://www.ecpgr.cgiar.org/resources/germplasm-databases/list-of-germplasm-databases/crop-databases/crop-database-windows/lupin
https://www.ecpgr.cgiar.org/resources/germplasm-databases/list-of-germplasm-databases/crop-databases/crop-database-windows/lupin
https://www.ecpgr.cgiar.org/resources/germplasm-databases/list-of-germplasm-databases/crop-databases/crop-database-windows/lupin
https://www.ecpgr.cgiar.org/resources/germplasm-databases/list-of-germplasm-databases/crop-databases/crop-database-windows/lupin
https://www.ecpgr.cgiar.org/resources/germplasm-databases/list-of-germplasm-databases/crop-databases/crop-database-windows/lupin

https://www.ecpgr.cgiar.org/resources/germplasm-databases/list-of-germplasm-databases/crop-databases/crop-database-windows/lupin
https://www.ecpgr.cgiar.org/resources/germplasm-databases/list-of-germplasm-databases/crop-databases/crop-database-windows/lupin
https://www.ecpgr.cgiar.org/resources/germplasm-databases/list-of-germplasm-databases/crop-databases/crop-database-windows/lupin
https://www.ecpgr.cgiar.org/resources/germplasm-databases/list-of-germplasm-databases/crop-databases/crop-database-windows/lupin
https://www.ecpgr.cgiar.org/resources/germplasm-databases/list-of-germplasm-databases/crop-databases/crop-database-windows/lupin
https://www.ecpgr.cgiar.org/resources/germplasm-databases/list-of-germplasm-databases/crop-databases/crop-database-windows/lupin
https://www.ecpgr.cgiar.org/resources/germplasm-databases/list-of-germplasm-databases/crop-databases/crop-database-windows/lupin
https://www.ecpgr.cgiar.org/resources/germplasm-databases/list-of-germplasm-databases/crop-databases/crop-database-windows/lupin
https://www.ecpgr.cgiar.org/resources/germplasm-databases/list-of-germplasm-databases/crop-databases/crop-database-windows/lupin
https://www.ecpgr.cgiar.org/resources/germplasm-databases/list-of-germplasm-databases/crop-databases/crop-database-windows/lupin
https://www.ecpgr.cgiar.org/resources/germplasm-databases/list-of-germplasm-databases/crop-databases/crop-database-windows/lupin
https://www.ecpgr.cgiar.org/resources/germplasm-databases/list-of-germplasm-databases/crop-databases/crop-database-windows/lupin
https://www.ecpgr.cgiar.org/resources/germplasm-databases/list-of-germplasm-databases/crop-databases/crop-database-windows/lupin
https://www.ecpgr.cgiar.org/resources/germplasm-databases/list-of-germplasm-databases/crop-databases/crop-database-windows/lupin
https://www.ecpgr.cgiar.org/resources/germplasm-databases/list-of-germplasm-databases/crop-databases/crop-database-windows/lupin
https://www.ecpgr.cgiar.org/resources/germplasm-databases/list-of-germplasm-databases/crop-databases/crop-database-windows/lupin
https://www.ecpgr.cgiar.org/resources/germplasm-databases/list-of-germplasm-databases/crop-databases/crop-database-windows/lupin
https://www.ecpgr.cgiar.org/resources/germplasm-databases/list-of-germplasm-databases/crop-databases/crop-database-windows/lupin
https://www.ecpgr.cgiar.org/resources/germplasm-databases/list-of-germplasm-databases/crop-databases/crop-database-windows/lupin
https://www.ecpgr.cgiar.org/resources/germplasm-databases/list-of-germplasm-databases/crop-databases/crop-database-windows/lupin
https://www.ecpgr.cgiar.org/resources/germplasm-databases/list-of-germplasm-databases/crop-databases/crop-database-windows/lupin
https://www.ecpgr.cgiar.org/resources/germplasm-databases/list-of-germplasm-databases/crop-databases/crop-database-windows/lupin
https://www.ecpgr.cgiar.org/resources/germplasm-databases/list-of-germplasm-databases/crop-databases/crop-database-windows/lupin
https://www.ecpgr.cgiar.org/resources/germplasm-databases/list-of-germplasm-databases/crop-databases/crop-database-windows/lupin
https://www.ecpgr.cgiar.org/resources/germplasm-databases/list-of-germplasm-databases/crop-databases/crop-database-windows/lupin
https://www.ecpgr.cgiar.org/resources/germplasm-databases/list-of-germplasm-databases/crop-databases/crop-database-windows/lupin
https://www.ecpgr.cgiar.org/resources/germplasm-databases/list-of-germplasm-databases/crop-databases/crop-database-windows/lupin
https://www.ecpgr.cgiar.org/resources/germplasm-databases/list-of-germplasm-databases/crop-databases/crop-database-windows/lupin
https://www.ecpgr.cgiar.org/resources/germplasm-databases/list-of-germplasm-databases/crop-databases/crop-database-windows/lupin
https://www.ecpgr.cgiar.org/resources/germplasm-databases/list-of-germplasm-databases/crop-databases/crop-database-windows/lupin
https://www.ecpgr.cgiar.org/resources/germplasm-databases/list-of-germplasm-databases/crop-databases/crop-database-windows/lupin
https://www.ecpgr.cgiar.org/resources/germplasm-databases/list-of-germplasm-databases/crop-databases/crop-database-windows/lupin
https://www.ecpgr.cgiar.org/resources/germplasm-databases/list-of-germplasm-databases/crop-databases/crop-database-windows/lupin
https://www.ecpgr.cgiar.org/resources/germplasm-databases/list-of-germplasm-databases/crop-databases/crop-database-windows/lupin
https://www.ecpgr.cgiar.org/resources/germplasm-databases/list-of-germplasm-databases/crop-databases/crop-database-windows/lupin
https://www.ecpgr.cgiar.org/resources/germplasm-databases/list-of-germplasm-databases/crop-databases/crop-database-windows/lupin
https://www.ecpgr.cgiar.org/resources/germplasm-databases/list-of-germplasm-databases/crop-databases/crop-database-windows/lupin
https://www.ecpgr.cgiar.org/resources/germplasm-databases/list-of-germplasm-databases/crop-databases/crop-database-windows/lupin
https://www.ecpgr.cgiar.org/resources/germplasm-databases/list-of-germplasm-databases/crop-databases/crop-database-windows/lupin
https://www.ecpgr.cgiar.org/resources/germplasm-databases/list-of-germplasm-databases/crop-databases/crop-database-windows/lupin
https://www.ecpgr.cgiar.org/resources/germplasm-databases/list-of-germplasm-databases/crop-databases/crop-database-windows/lupin
https://www.ecpgr.cgiar.org/resources/germplasm-databases/list-of-germplasm-databases/crop-databases/crop-database-windows/lupin
https://www.ecpgr.cgiar.org/resources/germplasm-databases/list-of-germplasm-databases/crop-databases/crop-database-windows/lupin
https://www.ecpgr.cgiar.org/resources/germplasm-databases/list-of-germplasm-databases/crop-databases/crop-database-windows/lupin
https://www.ecpgr.cgiar.org/resources/germplasm-databases/list-of-germplasm-databases/crop-databases/crop-database-windows/lupin
https://www.ecpgr.cgiar.org/resources/germplasm-databases/list-of-germplasm-databases/crop-databases/crop-database-windows/lupin
https://www.ecpgr.cgiar.org/resources/germplasm-databases/list-of-germplasm-databases/crop-databases/crop-database-windows/lupin
https://www.ecpgr.cgiar.org/resources/germplasm-databases/list-of-germplasm-databases/crop-databases/crop-database-windows/lupin
https://www.genesys-pgr.org/a/38fa7c3d-42b0-4589-a418-ad7b74348cb7
https://www.ecpgr.cgiar.org/resources/germplasm-databases/list-of-germplasm-databases/crop-databases/crop-database-windows/lupin
https://www.ecpgr.cgiar.org/resources/germplasm-databases/list-of-germplasm-databases/crop-databases/crop-database-windows/lupin
https://www.ecpgr.cgiar.org/resources/germplasm-databases/list-of-germplasm-databases/crop-databases/crop-database-windows/lupin

https://www.ecpgr.cgiar.org/resources/germplasm-databases/list-of-germplasm-databases/crop-databases/crop-database-windows/lupin
https://www.ecpgr.cgiar.org/resources/germplasm-databases/list-of-germplasm-databases/crop-databases/crop-database-windows/lupin
https://www.ecpgr.cgiar.org/resources/germplasm-databases/list-of-germplasm-databases/crop-databases/crop-database-windows/lupin
https://www.ecpgr.cgiar.org/resources/germplasm-databases/list-of-germplasm-databases/crop-databases/crop-database-windows/lupin
https://www.ecpgr.cgiar.org/resources/germplasm-databases/list-of-germplasm-databases/crop-databases/crop-database-windows/lupin
https://www.ecpgr.cgiar.org/resources/germplasm-databases/list-of-germplasm-databases/crop-databases/crop-database-windows/lupin
https://www.ecpgr.cgiar.org/resources/germplasm-databases/list-of-germplasm-databases/crop-databases/crop-database-windows/lupin
https://www.ecpgr.cgiar.org/resources/germplasm-databases/list-of-germplasm-databases/crop-databases/crop-database-windows/lupin
https://www.ecpgr.cgiar.org/resources/germplasm-databases/list-of-germplasm-databases/crop-databases/crop-database-windows/lupin
https://www.ecpgr.cgiar.org/resources/germplasm-databases/list-of-germplasm-databases/crop-databases/crop-database-windows/lupin
https://www.ecpgr.cgiar.org/resources/germplasm-databases/list-of-germplasm-databases/crop-databases/crop-database-windows/lupin
https://www.ecpgr.cgiar.org/resources/germplasm-databases/list-of-germplasm-databases/crop-databases/crop-database-windows/lupin
https://www.ecpgr.cgiar.org/resources/germplasm-databases/list-of-germplasm-databases/crop-databases/crop-database-windows/lupin
https://www.ecpgr.cgiar.org/resources/germplasm-databases/list-of-germplasm-databases/crop-databases/crop-database-windows/lupin
https://www.ecpgr.cgiar.org/resources/germplasm-databases/list-of-germplasm-databases/crop-databases/crop-database-windows/lupin
https://www.ecpgr.cgiar.org/resources/germplasm-databases/list-of-germplasm-databases/crop-databases/crop-database-windows/lupin
https://www.ecpgr.cgiar.org/resources/germplasm-databases/list-of-germplasm-databases/crop-databases/crop-database-windows/lupin
https://www.ecpgr.cgiar.org/resources/germplasm-databases/list-of-germplasm-databases/crop-databases/crop-database-windows/lupin
https://www.ecpgr.cgiar.org/resources/germplasm-databases/list-of-germplasm-databases/crop-databases/crop-database-windows/lupin
https://www.ecpgr.cgiar.org/resources/germplasm-databases/list-of-germplasm-databases/crop-databases/crop-database-windows/lupin
https://www.ecpgr.cgiar.org/resources/germplasm-databases/list-of-germplasm-databases/crop-databases/crop-database-windows/lupin
https://www.ecpgr.cgiar.org/resources/germplasm-databases/list-of-germplasm-databases/crop-databases/crop-database-windows/lupin
https://www.ecpgr.cgiar.org/resources/germplasm-databases/list-of-germplasm-databases/crop-databases/crop-database-windows/lupin
https://www.ecpgr.cgiar.org/resources/germplasm-databases/list-of-germplasm-databases/crop-databases/crop-database-windows/lupin
https://www.ecpgr.cgiar.org/resources/germplasm-databases/list-of-germplasm-databases/crop-databases/crop-database-windows/lupin
https://www.ecpgr.cgiar.org/resources/germplasm-databases/list-of-germplasm-databases/crop-databases/crop-database-windows/lupin
https://www.ecpgr.cgiar.org/resources/germplasm-databases/list-of-germplasm-databases/crop-databases/crop-database-windows/lupin
https://www.ecpgr.cgiar.org/resources/germplasm-databases/list-of-germplasm-databases/crop-databases/crop-database-windows/lupin
https://www.ecpgr.cgiar.org/resources/germplasm-databases/list-of-germplasm-databases/crop-databases/crop-database-windows/lupin
https://www.ecpgr.cgiar.org/resources/germplasm-databases/list-of-germplasm-databases/crop-databases/crop-database-windows/lupin
https://www.ecpgr.cgiar.org/resources/germplasm-databases/list-of-germplasm-databases/crop-databases/crop-database-windows/lupin
https://www.ecpgr.cgiar.org/resources/germplasm-databases/list-of-germplasm-databases/crop-databases/crop-database-windows/lupin
https://www.ecpgr.cgiar.org/resources/germplasm-databases/list-of-germplasm-databases/crop-databases/crop-database-windows/lupin
https://www.ecpgr.cgiar.org/resources/germplasm-databases/list-of-germplasm-databases/crop-databases/crop-database-windows/lupin
https://www.ecpgr.cgiar.org/resources/germplasm-databases/list-of-germplasm-databases/crop-databases/crop-database-windows/lupin
https://www.genesys-pgr.org/a/92024e7a-5d2f-44c5-a6b8-1bd4917ab9f4
https://www.ecpgr.cgiar.org/resources/germplasm-databases/list-of-germplasm-databases/crop-databases/crop-database-windows/lupin
https://www.ecpgr.cgiar.org/resources/germplasm-databases/list-of-germplasm-databases/crop-databases/crop-database-windows/lupin
https://www.ecpgr.cgiar.org/resources/germplasm-databases/list-of-germplasm-databases/crop-databases/crop-database-windows/lupin
https://www.ecpgr.cgiar.org/resources/germplasm-databases/list-of-germplasm-databases/crop-databases/crop-database-windows/lupin
https://www.ecpgr.cgiar.org/resources/germplasm-databases/list-of-germplasm-databases/crop-databases/crop-database-windows/lupin
https://www.ecpgr.cgiar.org/resources/germplasm-databases/list-of-germplasm-databases/crop-databases/crop-database-windows/lupin
https://www.ecpgr.cgiar.org/resources/germplasm-databases/list-of-germplasm-databases/crop-databases/crop-database-windows/lupin
https://www.ecpgr.cgiar.org/resources/germplasm-databases/list-of-germplasm-databases/crop-databases/crop-database-windows/lupin
https://www.ecpgr.cgiar.org/resources/germplasm-databases/list-of-germplasm-databases/crop-databases/crop-database-windows/lupin
https://www.ecpgr.cgiar.org/resources/germplasm-databases/list-of-germplasm-databases/crop-databases/crop-database-windows/lupin
https://www.ecpgr.cgiar.org/resources/germplasm-databases/list-of-germplasm-databases/crop-databases/crop-database-windows/lupin
https://www.ecpgr.cgiar.org/resources/germplasm-databases/list-of-germplasm-databases/crop-databases/crop-database-windows/lupin
https://www.ecpgr.cgiar.org/resources/germplasm-databases/list-of-germplasm-databases/crop-databases/crop-database-windows/lupin
https://www.ecpgr.cgiar.org/resources/germplasm-databases/list-of-germplasm-databases/crop-databases/crop-database-windows/lupin
https://www.ecpgr.cgiar.org/resources/germplasm-databases/list-of-germplasm-databases/crop-databases/crop-database-windows/lupin
https://www.ecpgr.cgiar.org/resources/germplasm-databases/list-of-germplasm-databases/crop-databases/crop-database-windows/lupin

https://www.ecpgr.cgiar.org/resources/germplasm-databases/list-of-germplasm-databases/crop-databases/crop-database-windows/lupin
https://www.ecpgr.cgiar.org/resources/germplasm-databases/list-of-germplasm-databases/crop-databases/crop-database-windows/lupin
https://www.genesys-pgr.org/a/577787d5-3683-4d97-9c3a-df43af5975a4
https://www.ecpgr.cgiar.org/resources/germplasm-databases/list-of-germplasm-databases/crop-databases/crop-database-windows/lupin
https://www.genesys-pgr.org/a/c20ba857-7d8c-4c8a-96f9-32e3abe8950a
https://www.ecpgr.cgiar.org/resources/germplasm-databases/list-of-germplasm-databases/crop-databases/crop-database-windows/lupin
https://www.ecpgr.cgiar.org/resources/germplasm-databases/list-of-germplasm-databases/crop-databases/crop-database-windows/lupin
https://www.ecpgr.cgiar.org/resources/germplasm-databases/list-of-germplasm-databases/crop-databases/crop-database-windows/lupin
https://www.genesys-pgr.org/a/ce59c5e4-94c7-46db-8b93-e3ef1cc4cc23
https://www.ecpgr.cgiar.org/resources/germplasm-databases/list-of-germplasm-databases/crop-databases/crop-database-windows/lupin
https://www.ecpgr.cgiar.org/resources/germplasm-databases/list-of-germplasm-databases/crop-databases/crop-database-windows/lupin
https://www.ecpgr.cgiar.org/resources/germplasm-databases/list-of-germplasm-databases/crop-databases/crop-database-windows/lupin
https://www.ecpgr.cgiar.org/resources/germplasm-databases/list-of-germplasm-databases/crop-databases/crop-database-windows/lupin
https://www.ecpgr.cgiar.org/resources/germplasm-databases/list-of-germplasm-databases/crop-databases/crop-database-windows/lupin
https://www.ecpgr.cgiar.org/resources/germplasm-databases/list-of-germplasm-databases/crop-databases/crop-database-windows/lupin
https://www.ecpgr.cgiar.org/resources/germplasm-databases/list-of-germplasm-databases/crop-databases/crop-database-windows/lupin
https://www.genesys-pgr.org/a/4f80170a-470c-4ec0-9979-8884b64dab67
https://www.ecpgr.cgiar.org/resources/germplasm-databases/list-of-germplasm-databases/crop-databases/crop-database-windows/lupin
https://www.ecpgr.cgiar.org/resources/germplasm-databases/list-of-germplasm-databases/crop-databases/crop-database-windows/lupin
https://www.ecpgr.cgiar.org/resources/germplasm-databases/list-of-germplasm-databases/crop-databases/crop-database-windows/lupin
https://www.ecpgr.cgiar.org/resources/germplasm-databases/list-of-germplasm-databases/crop-databases/crop-database-windows/lupin
https://www.ecpgr.cgiar.org/resources/germplasm-databases/list-of-germplasm-databases/crop-databases/crop-database-windows/lupin
https://www.ecpgr.cgiar.org/resources/germplasm-databases/list-of-germplasm-databases/crop-databases/crop-database-windows/lupin
https://www.ecpgr.cgiar.org/resources/germplasm-databases/list-of-germplasm-databases/crop-databases/crop-database-windows/lupin
https://www.ecpgr.cgiar.org/resources/germplasm-databases/list-of-germplasm-databases/crop-databases/crop-database-windows/lupin
https://www.ecpgr.cgiar.org/resources/germplasm-databases/list-of-germplasm-databases/crop-databases/crop-database-windows/lupin
https://www.ecpgr.cgiar.org/resources/germplasm-databases/list-of-germplasm-databases/crop-databases/crop-database-windows/lupin
https://www.ecpgr.cgiar.org/resources/germplasm-databases/list-of-germplasm-databases/crop-databases/crop-database-windows/lupin
https://www.ecpgr.cgiar.org/resources/germplasm-databases/list-of-germplasm-databases/crop-databases/crop-database-windows/lupin
https://www.ecpgr.cgiar.org/resources/germplasm-databases/list-of-germplasm-databases/crop-databases/crop-database-windows/lupin
https://www.ecpgr.cgiar.org/resources/germplasm-databases/list-of-germplasm-databases/crop-databases/crop-database-windows/lupin
https://www.ecpgr.cgiar.org/resources/germplasm-databases/list-of-germplasm-databases/crop-databases/crop-database-windows/lupin
https://www.ecpgr.cgiar.org/resources/germplasm-databases/list-of-germplasm-databases/crop-databases/crop-database-windows/lupin
https://www.ecpgr.cgiar.org/resources/germplasm-databases/list-of-germplasm-databases/crop-databases/crop-database-windows/lupin
https://www.ecpgr.cgiar.org/resources/germplasm-databases/list-of-germplasm-databases/crop-databases/crop-database-windows/lupin
https://www.ecpgr.cgiar.org/resources/germplasm-databases/list-of-germplasm-databases/crop-databases/crop-database-windows/lupin
https://www.ecpgr.cgiar.org/resources/germplasm-databases/list-of-germplasm-databases/crop-databases/crop-database-windows/lupin
https://www.ecpgr.cgiar.org/resources/germplasm-databases/list-of-germplasm-databases/crop-databases/crop-database-windows/lupin
https://www.ecpgr.cgiar.org/resources/germplasm-databases/list-of-germplasm-databases/crop-databases/crop-database-windows/lupin
https://www.ecpgr.cgiar.org/resources/germplasm-databases/list-of-germplasm-databases/crop-databases/crop-database-windows/lupin
https://www.ecpgr.cgiar.org/resources/germplasm-databases/list-of-germplasm-databases/crop-databases/crop-database-windows/lupin
https://www.ecpgr.cgiar.org/resources/germplasm-databases/list-of-germplasm-databases/crop-databases/crop-database-windows/lupin
https://www.ecpgr.cgiar.org/resources/germplasm-databases/list-of-germplasm-databases/crop-databases/crop-database-windows/lupin
https://www.ecpgr.cgiar.org/resources/germplasm-databases/list-of-germplasm-databases/crop-databases/crop-database-windows/lupin
https://www.ecpgr.cgiar.org/resources/germplasm-databases/list-of-germplasm-databases/crop-databases/crop-database-windows/lupin
https://www.ecpgr.cgiar.org/resources/germplasm-databases/list-of-germplasm-databases/crop-databases/crop-database-windows/lupin
https://www.ecpgr.cgiar.org/resources/germplasm-databases/list-of-germplasm-databases/crop-databases/crop-database-windows/lupin
https://www.genesys-pgr.org/a/0d8c5d8f-1206-4dac-9051-5737e6996c22
https://www.genesys-pgr.org/a/4af569a5-0b1d-4f04-a0b5-cbbeffe7162a
https://www.ecpgr.cgiar.org/resources/germplasm-databases/list-of-germplasm-databases/crop-databases/crop-database-windows/lupin
https://www.ecpgr.cgiar.org/resources/germplasm-databases/list-of-germplasm-databases/crop-databases/crop-database-windows/lupin
https://www.ecpgr.cgiar.org/resources/germplasm-databases/list-of-germplasm-databases/crop-databases/crop-database-windows/lupin

https://www.ecpgr.cgiar.org/resources/germplasm-databases/list-of-germplasm-databases/crop-databases/crop-database-windows/lupin
https://www.ecpgr.cgiar.org/resources/germplasm-databases/list-of-germplasm-databases/crop-databases/crop-database-windows/lupin
https://www.ecpgr.cgiar.org/resources/germplasm-databases/list-of-germplasm-databases/crop-databases/crop-database-windows/lupin
https://www.ecpgr.cgiar.org/resources/germplasm-databases/list-of-germplasm-databases/crop-databases/crop-database-windows/lupin
https://www.ecpgr.cgiar.org/resources/germplasm-databases/list-of-germplasm-databases/crop-databases/crop-database-windows/lupin
https://www.ecpgr.cgiar.org/resources/germplasm-databases/list-of-germplasm-databases/crop-databases/crop-database-windows/lupin
https://www.ecpgr.cgiar.org/resources/germplasm-databases/list-of-germplasm-databases/crop-databases/crop-database-windows/lupin
https://www.genesys-pgr.org/a/f66fe44d-392e-4267-8c7d-93a2aaf15d1d
https://www.genesys-pgr.org/a/8e920f26-adee-45b5-ad83-f21d5a88a8d0
https://www.ecpgr.cgiar.org/resources/germplasm-databases/list-of-germplasm-databases/crop-databases/crop-database-windows/lupin
https://www.ecpgr.cgiar.org/resources/germplasm-databases/list-of-germplasm-databases/crop-databases/crop-database-windows/lupin
https://www.ecpgr.cgiar.org/resources/germplasm-databases/list-of-germplasm-databases/crop-databases/crop-database-windows/lupin
https://www.ecpgr.cgiar.org/resources/germplasm-databases/list-of-germplasm-databases/crop-databases/crop-database-windows/lupin
https://www.ecpgr.cgiar.org/resources/germplasm-databases/list-of-germplasm-databases/crop-databases/crop-database-windows/lupin
https://www.ecpgr.cgiar.org/resources/germplasm-databases/list-of-germplasm-databases/crop-databases/crop-database-windows/lupin
https://www.ecpgr.cgiar.org/resources/germplasm-databases/list-of-germplasm-databases/crop-databases/crop-database-windows/lupin
https://www.ecpgr.cgiar.org/resources/germplasm-databases/list-of-germplasm-databases/crop-databases/crop-database-windows/lupin
https://www.ecpgr.cgiar.org/resources/germplasm-databases/list-of-germplasm-databases/crop-databases/crop-database-windows/lupin
https://www.ecpgr.cgiar.org/resources/germplasm-databases/list-of-germplasm-databases/crop-databases/crop-database-windows/lupin
https://www.ecpgr.cgiar.org/resources/germplasm-databases/list-of-germplasm-databases/crop-databases/crop-database-windows/lupin
https://www.ecpgr.cgiar.org/resources/germplasm-databases/list-of-germplasm-databases/crop-databases/crop-database-windows/lupin
https://www.genesys-pgr.org/10.25642/IPK/GBIS/45613
https://www.genesys-pgr.org/a/4c61f9a1-4413-4811-a403-40830a75f623
https://www.genesys-pgr.org/10.25642/IPK/GBIS/45615
https://www.genesys-pgr.org/a/8567e57a-beb2-48b9-8661-09f889703b61
https://www.genesys-pgr.org/a/e14052a2-141d-4b42-b180-6438af3d5617
https://www.genesys-pgr.org/10.25642/IPK/GBIS/45618
https://www.genesys-pgr.org/10.25642/IPK/GBIS/45619
https://www.genesys-pgr.org/a/8f9137d6-6d94-4e95-b928-96595b8fbec9
https://www.ecpgr.cgiar.org/resources/germplasm-databases/list-of-germplasm-databases/crop-databases/crop-database-windows/lupin
https://www.genesys-pgr.org/10.25642/IPK/GBIS/45622
https://www.genesys-pgr.org/10.25642/IPK/GBIS/45623
https://www.genesys-pgr.org/10.25642/IPK/GBIS/45624
https://www.genesys-pgr.org/10.25642/IPK/GBIS/77732
https://www.genesys-pgr.org/10.25642/IPK/GBIS/45626
https://www.genesys-pgr.org/10.25642/IPK/GBIS/45627
https://www.genesys-pgr.org/10.25642/IPK/GBIS/78125
https://www.genesys-pgr.org/10.25642/IPK/GBIS/78133
https://www.genesys-pgr.org/10.25642/IPK/GBIS/78150
https://www.genesys-pgr.org/10.25642/IPK/GBIS/78209
https://gbis.ipk-gatersleben.de/gbis2i/faces/pages/detail.jsf?akzessionId=45632
https://gbis.ipk-gatersleben.de/gbis2i/faces/pages/detail.jsf?akzessionId=45633
https://gbis.ipk-gatersleben.de/gbis2i/faces/pages/detail.jsf?akzessionId=45635
https://www.genesys-pgr.org/10.25642/IPK/GBIS/45639
https://www.genesys-pgr.org/10.25642/IPK/GBIS/45640
https://www.genesys-pgr.org/10.25642/IPK/GBIS/45641
https://www.genesys-pgr.org/10.25642/IPK/GBIS/45642
https://eurisco.ipk-gatersleben.de/apex/f?p=103:16:::::P16_EURISCO_ACC_ID:138205
https://www.genesys-pgr.org/10.25642/IPK/GBIS/45645
https://www.genesys-pgr.org/10.25642/IPK/GBIS/45646
https://gbis.ipk-gatersleben.de/gbis2i/faces/pages/detail.jsf?akzessionId=45647
https://www.genesys-pgr.org/10.25642/IPK/GBIS/45648

https://www.genesys-pgr.org/10.25642/IPK/GBIS/45649
https://www.genesys-pgr.org/10.25642/IPK/GBIS/45650
https://www.genesys-pgr.org/10.25642/IPK/GBIS/45651
https://www.genesys-pgr.org/10.25642/IPK/GBIS/45652
https://www.genesys-pgr.org/10.25642/IPK/GBIS/45653
https://www.genesys-pgr.org/10.25642/IPK/GBIS/45654
https://eurisco.ipk-gatersleben.de/apex/f?p=103:16:::::P16_EURISCO_ACC_ID:138139
https://gbis.ipk-gatersleben.de/gbis2i/faces/pages/detail.jsf?akzessionId=45656
https://www.genesys-pgr.org/10.25642/IPK/GBIS/45657
https://www.genesys-pgr.org/10.25642/IPK/GBIS/45658
https://www.genesys-pgr.org/10.25642/IPK/GBIS/45659
https://www.genesys-pgr.org/10.25642/IPK/GBIS/45660
https://www.genesys-pgr.org/10.25642/IPK/GBIS/45661
https://www.genesys-pgr.org/10.25642/IPK/GBIS/45662
https://gbis.ipk-gatersleben.de/gbis2i/faces/pages/detail.jsf;jsessionid=joa3dzwt6hqjcYYlbOphiEpGPfAuj1FVFYJ6dqsZ0S2-6sAEo
https://gbis.ipk-gatersleben.de/gbis2i/faces/pages/detail.jsf?akzessionId=45664
https://www.genesys-pgr.org/10.25642/IPK/GBIS/45665
https://www.genesys-pgr.org/10.25642/IPK/GBIS/77408
https://www.genesys-pgr.org/10.25642/IPK/GBIS/77409
https://www.genesys-pgr.org/10.25642/IPK/GBIS/77426
https://www.genesys-pgr.org/10.25642/IPK/GBIS/77430
https://www.genesys-pgr.org/10.25642/IPK/GBIS/77447
https://www.genesys-pgr.org/10.25642/IPK/GBIS/77450
https://www.genesys-pgr.org/10.25642/IPK/GBIS/77452
https://www.genesys-pgr.org/10.25642/IPK/GBIS/77453
https://www.genesys-pgr.org/10.25642/IPK/GBIS/77440
https://www.genesys-pgr.org/10.25642/IPK/GBIS/77454
https://www.genesys-pgr.org/10.25642/IPK/GBIS/77473
https://www.genesys-pgr.org/10.25642/IPK/GBIS/77484
https://www.genesys-pgr.org/10.25642/IPK/GBIS/77486
https://www.genesys-pgr.org/10.25642/IPK/GBIS/77488
https://www.genesys-pgr.org/10.25642/IPK/GBIS/77517
https://www.genesys-pgr.org/10.25642/IPK/GBIS/77519
https://www.genesys-pgr.org/10.25642/IPK/GBIS/77524
https://www.genesys-pgr.org/10.25642/IPK/GBIS/77526
https://www.genesys-pgr.org/10.25642/IPK/GBIS/77527
https://www.genesys-pgr.org/10.25642/IPK/GBIS/77530
https://www.genesys-pgr.org/10.25642/IPK/GBIS/77546
https://www.genesys-pgr.org/10.25642/IPK/GBIS/77547
https://www.genesys-pgr.org/10.25642/IPK/GBIS/77555
https://www.genesys-pgr.org/10.25642/IPK/GBIS/77560
https://www.genesys-pgr.org/10.25642/IPK/GBIS/77562
https://www.genesys-pgr.org/10.25642/IPK/GBIS/77567
https://www.genesys-pgr.org/10.25642/IPK/GBIS/77571
https://www.genesys-pgr.org/10.25642/IPK/GBIS/77572
https://www.genesys-pgr.org/10.25642/IPK/GBIS/77574
https://www.genesys-pgr.org/10.25642/IPK/GBIS/77576
https://www.genesys-pgr.org/10.25642/IPK/GBIS/77577
https://www.genesys-pgr.org/10.25642/IPK/GBIS/77592
https://www.genesys-pgr.org/10.25642/IPK/GBIS/77607
https://www.genesys-pgr.org/10.25642/IPK/GBIS/77613
https://www.genesys-pgr.org/10.25642/IPK/GBIS/77620

https://www.genesys-pgr.org/10.25642/IPK/GBIS/77621
https://www.genesys-pgr.org/10.25642/IPK/GBIS/77622
https://www.genesys-pgr.org/10.25642/IPK/GBIS/77626
https://www.genesys-pgr.org/10.25642/IPK/GBIS/77634
https://www.genesys-pgr.org/10.25642/IPK/GBIS/77638
https://gbis.ipk-gatersleben.de/gbis2i/faces/pages/detail.jsf?akzessionId=73618
https://www.genesys-pgr.org/10.25642/IPK/GBIS/79636
https://www.genesys-pgr.org/10.25642/IPK/GBIS/79654
https://www.genesys-pgr.org/10.25642/IPK/GBIS/77649
https://www.genesys-pgr.org/10.25642/IPK/GBIS/77650
https://www.genesys-pgr.org/10.25642/IPK/GBIS/77655
https://www.genesys-pgr.org/10.25642/IPK/GBIS/77665
https://www.genesys-pgr.org/10.25642/IPK/GBIS/77694
https://www.genesys-pgr.org/10.25642/IPK/GBIS/77699
https://www.genesys-pgr.org/10.25642/IPK/GBIS/77714
https://www.genesys-pgr.org/10.25642/IPK/GBIS/77715
https://www.genesys-pgr.org/10.25642/IPK/GBIS/77718
https://www.genesys-pgr.org/10.25642/IPK/GBIS/77722
https://www.genesys-pgr.org/10.25642/IPK/GBIS/77723
https://www.genesys-pgr.org/10.25642/IPK/GBIS/77724
https://www.genesys-pgr.org/10.25642/IPK/GBIS/77732
https://www.genesys-pgr.org/10.25642/IPK/GBIS/77743
https://www.genesys-pgr.org/10.25642/IPK/GBIS/77765
https://www.genesys-pgr.org/10.25642/IPK/GBIS/77776
https://www.genesys-pgr.org/10.25642/IPK/GBIS/77786
https://www.genesys-pgr.org/10.25642/IPK/GBIS/77787
https://gbis.ipk-gatersleben.de/gbis2i/faces/pages/detail.jsf?akzessionId=46048
https://www.genesys-pgr.org/10.25642/IPK/GBIS/46049
https://apex.ipk-gatersleben.de/apex/f?p=DOI:RESOLVE:::NO:RP:DOI:10.25642/IPK/GBIS/46051
https://apex.ipk-gatersleben.de/apex/f?p=DOI:RESOLVE:::NO:RP:DOI:10.25642/IPK/GBIS/46052
https://apex.ipk-gatersleben.de/apex/f?p=DOI:RESOLVE:::NO:RP:DOI:10.25642/IPK/GBIS/46053
https://apex.ipk-gatersleben.de/apex/f?p=DOI:RESOLVE:::NO:RP:DOI:10.25642/IPK/GBIS/46054
https://apex.ipk-gatersleben.de/apex/f?p=DOI:RESOLVE:::NO:RP:DOI:10.25642/IPK/GBIS/46055
https://apex.ipk-gatersleben.de/apex/f?p=DOI:RESOLVE:::NO:RP:DOI:10.25642/IPK/GBIS/46056
https://www.genesys-pgr.org/10.25642/IPK/GBIS/46057
https://apex.ipk-gatersleben.de/apex/f?p=DOI:RESOLVE:::NO:RP:DOI:10.25642/IPK/GBIS/46058
https://gbis.ipk-gatersleben.de/gbis2i/faces/pages/detail.jsf?akzessionId=46059
https://www.genesys-pgr.org/10.25642/IPK/GBIS/78715
https://www.genesys-pgr.org/10.25642/IPK/GBIS/46063
https://gbis.ipk-gatersleben.de/gbis2i/faces/pages/detail.jsf?akzessionId=46064
https://www.genesys-pgr.org/10.25642/IPK/GBIS/46065
https://www.genesys-pgr.org/10.25642/IPK/GBIS/46066
https://www.genesys-pgr.org/10.25642/IPK/GBIS/46067
https://www.genesys-pgr.org/10.25642/IPK/GBIS/46068
https://www.genesys-pgr.org/10.25642/IPK/GBIS/46069
https://www.genesys-pgr.org/10.25642/IPK/GBIS/46070
https://www.genesys-pgr.org/10.25642/IPK/GBIS/46071
https://www.genesys-pgr.org/10.25642/IPK/GBIS/46072
https://www.genesys-pgr.org/10.25642/IPK/GBIS/46073
https://www.genesys-pgr.org/10.25642/IPK/GBIS/46074
https://gbis.ipk-gatersleben.de/gbis2i/faces/pages/detail.jsf?akzessionId=46075
https://apex.ipk-gatersleben.de/apex/f?p=DOI:RESOLVE:::NO:RP:DOI:10.25642/IPK/GBIS/46076

https://www.genesys-pgr.org/10.25642/IPK/GBIS/78784
https://www.genesys-pgr.org/10.25642/IPK/GBIS/78882
https://www.genesys-pgr.org/10.25642/IPK/GBIS/46080
https://www.genesys-pgr.org/10.25642/IPK/GBIS/79359
https://www.genesys-pgr.org/10.25642/IPK/GBIS/79392
https://www.genesys-pgr.org/10.25642/IPK/GBIS/79400
https://www.genesys-pgr.org/10.25642/IPK/GBIS/79128
https://www.genesys-pgr.org/10.25642/IPK/GBIS/46086
https://www.genesys-pgr.org/10.25642/IPK/GBIS/79530
https://www.genesys-pgr.org/10.25642/IPK/GBIS/79627
https://www.genesys-pgr.org/10.25642/IPK/GBIS/79661
https://www.genesys-pgr.org/10.25642/IPK/GBIS/79701
https://www.genesys-pgr.org/10.25642/IPK/GBIS/79721
https://www.genesys-pgr.org/10.25642/IPK/GBIS/79748
https://www.genesys-pgr.org/10.25642/IPK/GBIS/78700
https://www.genesys-pgr.org/10.25642/IPK/GBIS/78849
https://www.genesys-pgr.org/10.25642/IPK/GBIS/78654
https://www.genesys-pgr.org/10.25642/IPK/GBIS/45436
https://www.genesys-pgr.org/10.25642/IPK/GBIS/45437
https://www.genesys-pgr.org/10.25642/IPK/GBIS/45438
https://gbis.ipk-gatersleben.de/gbis2i/faces/pages/detail.jsf?akzessionId=45439
https://gbis.ipk-gatersleben.de/gbis2i/faces/pages/detail.jsf?akzessionId=45371
https://gbis.ipk-gatersleben.de/gbis2i/faces/pages/detail.jsf?akzessionId=72929
https://gbis.ipk-gatersleben.de/gbis2i/faces/pages/detail.jsf?akzessionId=72930
https://apex.ipk-gatersleben.de/apex/f?p=DOI:RESOLVE:::NO:RP:DOI:10.25642/IPK/GBIS/72932
https://gbis.ipk-gatersleben.de/gbis2i/faces/pages/detail.jsf?akzessionId=72933
https://gbis.ipk-gatersleben.de/gbis2i/faces/pages/detail.jsf?akzessionId=72937
https://gbis.ipk-gatersleben.de/gbis2i/faces/pages/detail.jsf?akzessionId=72936
https://www.genesys-pgr.org/10.25642/IPK/GBIS/72940
https://www.genesys-pgr.org/10.25642/IPK/GBIS/79774
https://www.genesys-pgr.org/10.25642/IPK/GBIS/79850
https://www.genesys-pgr.org/10.25642/IPK/GBIS/79862
https://www.genesys-pgr.org/10.25642/IPK/GBIS/80256
https://www.genesys-pgr.org/10.25642/IPK/GBIS/80848
https://www.genesys-pgr.org/10.25642/IPK/GBIS/45366
https://www.genesys-pgr.org/10.25642/IPK/GBIS/45373
https://www.genesys-pgr.org/10.25642/IPK/GBIS/45372
https://www.genesys-pgr.org/10.25642/IPK/GBIS/23168
https://gbis.ipk-gatersleben.de/gbis2i/faces/pages/detail.jsf?akzessionId=97798
https://gbis.ipk-gatersleben.de/gbis2i/faces/pages/detail.jsf?akzessionId=45365
https://www.genesys-pgr.org/10.25642/IPK/GBIS/59323
https://www.genesys-pgr.org/10.25642/IPK/GBIS/68462
https://www.genesys-pgr.org/10.25642/IPK/GBIS/45346
https://www.genesys-pgr.org/10.25642/IPK/GBIS/45347
https://www.genesys-pgr.org/10.25642/IPK/GBIS/45348
https://www.genesys-pgr.org/10.25642/IPK/GBIS/45349
https://www.genesys-pgr.org/10.25642/IPK/GBIS/45350
https://www.genesys-pgr.org/10.25642/IPK/GBIS/262896
https://www.genesys-pgr.org/10.25642/IPK/GBIS/262897
https://www.genesys-pgr.org/10.25642/IPK/GBIS/262898
https://www.genesys-pgr.org/10.25642/IPK/GBIS/262899
https://www.genesys-pgr.org/10.25642/IPK/GBIS/262900

https://www.genesys-pgr.org/10.25642/IPK/GBIS/262905
https://www.genesys-pgr.org/10.25642/IPK/GBIS/228963
https://www.genesys-pgr.org/10.25642/IPK/GBIS/259603
https://eurisco.ipk-gatersleben.de/apex/f?p=103:16:::::P16_EURISCO_ACC_ID:134917
https://www.genesys-pgr.org/10.25642/IPK/GBIS/223389
https://eurisco.ipk-gatersleben.de/apex/f?p=103:16:::::P16_EURISCO_ACC_ID:134916
https://gbis.ipk-gatersleben.de/gbis2i/faces/pages/detail.jsf?akzessionId=246227
https://eurisco.ipk-gatersleben.de/apex/f?p=103:16:::::P16_EURISCO_ACC_ID:134915
https://eurisco.ipk-gatersleben.de/apex/f?p=103:16:::::P16_EURISCO_ACC_ID:134167
https://gbis.ipk-gatersleben.de/gbis2i/faces/pages/detail.jsf?akzessionId=249317
https://eurisco.ipk-gatersleben.de/apex/f?p=103:16:::::P16_EURISCO_ACC_ID:134165
https://www.genesys-pgr.org/10.25642/IPK/GBIS/259894
https://eurisco.ipk-gatersleben.de/apex/f?p=103:16:::::P16_EURISCO_ACC_ID:134163
https://eurisco.ipk-gatersleben.de/apex/f?p=103:16:::::P16_EURISCO_ACC_ID:134161
https://www.genesys-pgr.org/10.25642/IPK/GBIS/233812
https://eurisco.ipk-gatersleben.de/apex/f?p=103:16:::::P16_EURISCO_ACC_ID:134157
https://www.genesys-pgr.org/10.25642/IPK/GBIS/228956
https://eurisco.ipk-gatersleben.de/apex/f?p=103:16:::::P16_EURISCO_ACC_ID:134156
https://eurisco.ipk-gatersleben.de/apex/f?p=103:16:::::P16_EURISCO_ACC_ID:134154
https://eurisco.ipk-gatersleben.de/apex/f?p=103:16:::::P16_EURISCO_ACC_ID:134153
https://www.genesys-pgr.org/10.25642/IPK/GBIS/248181
https://eurisco.ipk-gatersleben.de/apex/f?p=103:16:::::P16_EURISCO_ACC_ID:134151
https://www.genesys-pgr.org/10.25642/IPK/GBIS/242528
https://gbis.ipk-gatersleben.de/gbis2i/faces/pages/detail.jsf?akzessionId=238804
https://www.genesys-pgr.org/10.25642/IPK/GBIS/228204
https://www.genesys-pgr.org/10.25642/IPK/GBIS/224576
https://eurisco.ipk-gatersleben.de/apex/f?p=103:16:::::P16_EURISCO_ACC_ID:136098
https://www.genesys-pgr.org/10.25642/IPK/GBIS/247212
https://www.genesys-pgr.org/10.25642/IPK/GBIS/230438
https://www.genesys-pgr.org/10.25642/IPK/GBIS/228175
https://www.genesys-pgr.org/10.25642/IPK/GBIS/239066
https://www.genesys-pgr.org/10.25642/IPK/GBIS/247926
https://www.genesys-pgr.org/10.25642/IPK/GBIS/254306
https://www.genesys-pgr.org/10.25642/IPK/GBIS/228197
https://www.genesys-pgr.org/10.25642/IPK/GBIS/247924
https://www.genesys-pgr.org/10.25642/IPK/GBIS/228148
https://www.genesys-pgr.org/10.25642/IPK/GBIS/246449
https://www.genesys-pgr.org/10.25642/IPK/GBIS/233441
https://www.genesys-pgr.org/10.25642/IPK/GBIS/242530
https://eurisco.ipk-gatersleben.de/apex/f?p=103:16:::::P16_EURISCO_ACC_ID:135877
https://www.genesys-pgr.org/10.25642/IPK/GBIS/247196
https://www.genesys-pgr.org/10.25642/IPK/GBIS/259585
https://www.genesys-pgr.org/10.25642/IPK/GBIS/239071
https://www.genesys-pgr.org/10.25642/IPK/GBIS/224174
https://www.genesys-pgr.org/10.25642/IPK/GBIS/239070
https://apex.ipk-gatersleben.de/apex/f?p=DOI:RESOLVE:::NO:RP:DOI:10.25642/IPK/GBIS/224587
https://www.genesys-pgr.org/10.25642/IPK/GBIS/223829
https://www.genesys-pgr.org/10.25642/IPK/GBIS/247888
https://www.genesys-pgr.org/10.25642/IPK/GBIS/230441
https://www.genesys-pgr.org/10.25642/IPK/GBIS/239052
https://apex.ipk-gatersleben.de/apex/f?p=DOI:RESOLVE:::NO:RP:DOI:10.25642/IPK/GBIS/259849
https://www.genesys-pgr.org/10.25642/IPK/GBIS/247202

https://www.genesys-pgr.org/10.25642/IPK/GBIS/228205
https://apex.ipk-gatersleben.de/apex/f?p=DOI:RESOLVE:::NO:RP:DOI:10.25642/IPK/GBIS/234245
https://www.genesys-pgr.org/10.25642/IPK/GBIS/242545
https://www.genesys-pgr.org/10.25642/IPK/GBIS/234274
https://www.genesys-pgr.org/10.25642/IPK/GBIS/230439
https://www.genesys-pgr.org/10.25642/IPK/GBIS/242527
https://www.genesys-pgr.org/10.25642/IPK/GBIS/235263
https://www.genesys-pgr.org/10.25642/IPK/GBIS/254000
https://www.genesys-pgr.org/10.25642/IPK/GBIS/228207
https://gbis.ipk-gatersleben.de/gbis2i/faces/pages/detail.jsf?akzessionId=223836
https://npgsweb.ars-grin.gov/gringlobal/accessiondetail?id=1854300
https://www.genesys-pgr.org/10.25642/IPK/GBIS/234250
https://www.genesys-pgr.org/10.25642/IPK/GBIS/224590
https://gbis.ipk-gatersleben.de/gbis2i/faces/pages/detail.jsf?akzessionId=239083
https://www.genesys-pgr.org/10.25642/IPK/GBIS/229777
https://www.genesys-pgr.org/10.25642/IPK/GBIS/242512
https://gbis.ipk-gatersleben.de/gbis2i/faces/pages/detail.jsf?akzessionId=248162
https://www.genesys-pgr.org/10.25642/IPK/GBIS/259875
https://www.genesys-pgr.org/10.25642/IPK/GBIS/259869
https://www.genesys-pgr.org/10.25642/IPK/GBIS/247199
https://www.genesys-pgr.org/10.25642/IPK/GBIS/254028
https://www.genesys-pgr.org/10.25642/IPK/GBIS/246460
https://www.genesys-pgr.org/10.25642/IPK/GBIS/254003
https://apex.ipk-gatersleben.de/apex/f?p=DOI:RESOLVE:::NO:RP:DOI:10.25642/IPK/GBIS/242539
https://eurisco.ipk-gatersleben.de/apex/f?p=103:16:::::P16_EURISCO_ACC_ID:136028
https://www.genesys-pgr.org/10.25642/IPK/GBIS/238820
https://apex.ipk-gatersleben.de/apex/f?p=DOI:RESOLVE:::NO:RP:DOI:10.25642/IPK/GBIS/239454
https://www.genesys-pgr.org/10.25642/IPK/GBIS/239453
https://gbis.ipk-gatersleben.de/gbis2i/faces/pages/detail.jsf?akzessionId=234304
https://www.genesys-pgr.org/10.25642/IPK/GBIS/228150
https://www.genesys-pgr.org/10.25642/IPK/GBIS/239464
https://www.genesys-pgr.org/10.25642/IPK/GBIS/253992
https://www.genesys-pgr.org/10.25642/IPK/GBIS/233436
https://www.genesys-pgr.org/10.25642/IPK/GBIS/254298
https://www.genesys-pgr.org/10.25642/IPK/GBIS/238859
https://www.genesys-pgr.org/10.25642/IPK/GBIS/259882
https://eurisco.ipk-gatersleben.de/apex/f?p=103:16:::::P16_EURISCO_ACC_ID:136013
https://www.genesys-pgr.org/10.25642/IPK/GBIS/260753
https://www.genesys-pgr.org/10.25642/IPK/GBIS/260725
https://www.genesys-pgr.org/10.25642/IPK/GBIS/228154
https://gbis.ipk-gatersleben.de/gbis2i/faces/pages/detail.jsf?akzessionId=1490669
https://gbis.ipk-gatersleben.de/gbis2i/faces/pages/detail.jsf?akzessionId=94639
https://www.genesys-pgr.org/10.25642/IPK/GBIS/94951
https://www.genesys-pgr.org/10.25642/IPK/GBIS/1805739
https://www.genesys-pgr.org/10.25642/IPK/GBIS/99326
https://www.genesys-pgr.org/10.25642/IPK/GBIS/2245402
https://www.genesys-pgr.org/10.25642/IPK/GBIS/2447670
https://www.genesys-pgr.org/10.25642/IPK/GBIS/264000
https://www.genesys-pgr.org/10.18730/1D2JQ
https://www.genesys-pgr.org/10.18730/1D2KR
https://www.genesys-pgr.org/10.18730/1D2MS
https://www.genesys-pgr.org/10.18730/1D2NT

https://www.genesys-pgr.org/10.18730/1D2PV
https://www.genesys-pgr.org/10.18730/1D2QW
https://www.genesys-pgr.org/10.18730/1D2RX
https://www.genesys-pgr.org/10.18730/1D2SY
https://www.genesys-pgr.org/10.18730/1D2TZ
https://www.genesys-pgr.org/10.18730/1D2V*
https://www.genesys-pgr.org/10.18730/1D2W~
https://www.genesys-pgr.org/10.18730/1D2X$
https://www.genesys-pgr.org/10.18730/1D2Y=
https://grinczech.vurv.cz/gringlobal/accessiondetail.aspx?id=28711
https://grinczech.vurv.cz/gringlobal/accessiondetail.aspx?id=28712
https://grinczech.vurv.cz/gringlobal/accessiondetail.aspx?id=28713
https://www.genesys-pgr.org/a/31abbee4-7026-4b50-bb14-7a43695c386c
https://grinczech.vurv.cz/gringlobal/accessiondetail.aspx?id=28723
https://www.genesys-pgr.org/a/a40b153c-4310-4ad8-b3c7-e31f08f30489
https://www.genesys-pgr.org/a/100f7d08-c825-4a78-8d2f-6059320a376c
https://www.genesys-pgr.org/a/0f9240c5-b31d-4c87-b45d-5513952e8505
https://www.genesys-pgr.org/a/b66cb378-72ed-411f-89e8-c6d53066d477
https://grinczech.vurv.cz/gringlobal/accessiondetail.aspx?id=28737
https://www.genesys-pgr.org/a/960c5682-9eb1-47fd-8fea-b28ea1053830
https://www.genesys-pgr.org/a/8eb6a73c-861e-4dbd-a6f7-928b78cb00ec
https://www.genesys-pgr.org/a/852703e0-8e10-457f-a9cd-f2b628465963
https://www.genesys-pgr.org/a/3e517732-14d7-4ec5-a511-c54ececaa175
https://www.genesys-pgr.org/a/75e0f266-5fea-4afe-9059-433d503ffb9b
https://www.genesys-pgr.org/a/5e08e4ad-0b10-4585-bec9-bfc28c4217cb
https://www.genesys-pgr.org/a/ed743901-c7f0-4644-8dfa-416be463da9b
https://www.genesys-pgr.org/a/cf8df296-14d0-447f-985b-c05a6ed635a1
https://www.genesys-pgr.org/a/eb95e43f-cda3-4e6d-a204-dfb745ef76c2
https://www.genesys-pgr.org/a/9e81d5d6-4677-44f7-ba1e-ba32b8e691ad
https://www.genesys-pgr.org/a/706e86a5-c8cc-4e94-986a-909a84fb5021
https://www.genesys-pgr.org/a/fbc15b96-4df8-4fed-95c3-ae8a8d52e45d
https://www.genesys-pgr.org/a/35eff782-df2f-469b-ac4a-62fe5daddf99
https://www.genesys-pgr.org/a/809106de-2533-4d91-856b-a942f1ada1f2
https://www.genesys-pgr.org/a/078537de-d9be-44fc-94ba-ec466f1bce53
https://www.genesys-pgr.org/a/a1f85428-53f2-4d4b-9250-0e982998d7c9
https://www.genesys-pgr.org/a/8f76ea9c-4e2c-4587-ad55-72be0dd9cdd1
https://www.genesys-pgr.org/a/4ade3b6f-b1c1-472e-a4ae-c3691193d86c
https://www.genesys-pgr.org/a/1405852c-0d37-4000-a74e-59462c4b7a34
https://www.genesys-pgr.org/a/61f4e070-fb41-4c43-8414-4523c3070e8b
https://www.genesys-pgr.org/a/923cc81d-0d20-49db-b9d1-53c03afc8aac
https://www.genesys-pgr.org/a/ec89a16b-c264-429e-84c7-1a34d50f76c2
https://www.genesys-pgr.org/a/b8690a9f-8811-4a94-ab91-c71dcd3f697d
https://www.genesys-pgr.org/a/f64836c8-a181-4b92-a2dc-e130f61b7374
https://www.genesys-pgr.org/a/193f08ea-e0b7-439c-88d4-8c78be4120f8
https://www.genesys-pgr.org/a/c3e912a5-0bd8-4961-8074-a3a77511fa9b
https://www.genesys-pgr.org/a/f2cc09ce-de08-4007-96ef-7f7ecd4ab3aa
https://www.genesys-pgr.org/a/a4cae986-4484-488a-8668-a83e55d97961
https://npgsweb.ars-grin.gov/gringlobal/accessiondetail?id=1193152
https://www.ecpgr.cgiar.org/resources/germplasm-databases/list-of-germplasm-databases/crop-databases/crop-database-windows/lupin

https://www.ecpgr.cgiar.org/resources/germplasm-databases/list-of-germplasm-databases/crop-databases/crop-database-windows/lupin

https://wyszukiwarka.ihar.edu.pl/pl/details/644316
https://www.ecpgr.cgiar.org/resources/germplasm-databases/list-of-germplasm-databases/crop-databases/crop-database-windows/lupin
https://www.genesys-pgr.org/a/5d97699e-ea40-45ab-b761-d04d8f5c4962
https://www.ecpgr.cgiar.org/resources/germplasm-databases/list-of-germplasm-databases/crop-databases/crop-database-windows/lupin
https://www.genesys-pgr.org/a/c558f87a-1fb3-4240-a8b6-7e0746aa28ab
https://www.ecpgr.cgiar.org/resources/germplasm-databases/list-of-germplasm-databases/crop-databases/crop-database-windows/lupin
https://www.ecpgr.cgiar.org/resources/germplasm-databases/list-of-germplasm-databases/crop-databases/crop-database-windows/lupin
https://www.genesys-pgr.org/a/cfa0137e-db30-4599-b22a-e032678894be
https://www.genesys-pgr.org/a/23a8e58e-268e-4cae-b1d1-6b5e4d35f3aa
https://www.genesys-pgr.org/a/f78ae048-10ca-4d4c-999e-0fd5684eef42
https://www.genesys-pgr.org/a/272d8102-a154-4cd9-9c56-9500c74aa2bf
https://www.genesys-pgr.org/a/edaa4cf4-73f2-4fca-b2ac-1def9561f119
https://www.genesys-pgr.org/a/19bc61e2-4811-4a9e-9837-3e4369433307
https://www.ecpgr.cgiar.org/resources/germplasm-databases/list-of-germplasm-databases/crop-databases/crop-database-windows/lupin

https://www.genesys-pgr.org/10.18730/VPR4K

https://www.genesys-pgr.org/a/e6ba9821-e563-4d46-80d4-c3c8a6820d33

https://www.genesys-pgr.org/a/ead4b6c5-a318-49f0-95c2-0466740b738d

https://www.genesys-pgr.org/a/cf4e7d2d-fa90-41db-94cc-4ef93b22e564
https://www.genesys-pgr.org/a/8e5e4589-f78b-463b-95ce-79f4107c4687
https://www.genesys-pgr.org/a/44950684-f16a-4e1c-bf0f-75cd997eeaa7
https://www.genesys-pgr.org/a/d45725ef-8bff-4489-aea0-5981b34c7ec6
https://www.ecpgr.cgiar.org/resources/germplasm-databases/list-of-germplasm-databases/crop-databases/crop-database-windows/lupin
https://www.ecpgr.cgiar.org/resources/germplasm-databases/list-of-germplasm-databases/crop-databases/crop-database-windows/lupin
https://www.genesys-pgr.org/a/f635e9c4-190e-4b83-a8f8-8e599d171a9e
https://www.ecpgr.cgiar.org/resources/germplasm-databases/list-of-germplasm-databases/crop-databases/crop-database-windows/lupin
https://www.ecpgr.cgiar.org/resources/germplasm-databases/list-of-germplasm-databases/crop-databases/crop-database-windows/lupin
https://www.ecpgr.cgiar.org/resources/germplasm-databases/list-of-germplasm-databases/crop-databases/crop-database-windows/lupin
https://www.ecpgr.cgiar.org/resources/germplasm-databases/list-of-germplasm-databases/crop-databases/crop-database-windows/lupin
https://www.ecpgr.cgiar.org/resources/germplasm-databases/list-of-germplasm-databases/crop-databases/crop-database-windows/lupin
https://www.ecpgr.cgiar.org/resources/germplasm-databases/list-of-germplasm-databases/crop-databases/crop-database-windows/lupin
https://www.ecpgr.cgiar.org/resources/germplasm-databases/list-of-germplasm-databases/crop-databases/crop-database-windows/lupin
https://www.ecpgr.cgiar.org/resources/germplasm-databases/list-of-germplasm-databases/crop-databases/crop-database-windows/lupin
https://www.ecpgr.cgiar.org/resources/germplasm-databases/list-of-germplasm-databases/crop-databases/crop-database-windows/lupin
https://www.ecpgr.cgiar.org/resources/germplasm-databases/list-of-germplasm-databases/crop-databases/crop-database-windows/lupin
https://www.ecpgr.cgiar.org/resources/germplasm-databases/list-of-germplasm-databases/crop-databases/crop-database-windows/lupin
https://www.ecpgr.cgiar.org/resources/germplasm-databases/list-of-germplasm-databases/crop-databases/crop-database-windows/lupin
https://www.ecpgr.cgiar.org/resources/germplasm-databases/list-of-germplasm-databases/crop-databases/crop-database-windows/lupin
https://www.ecpgr.cgiar.org/resources/germplasm-databases/list-of-germplasm-databases/crop-databases/crop-database-windows/lupin
https://www.ecpgr.cgiar.org/resources/germplasm-databases/list-of-germplasm-databases/crop-databases/crop-database-windows/lupin
https://www.ecpgr.cgiar.org/resources/germplasm-databases/list-of-germplasm-databases/crop-databases/crop-database-windows/lupin
https://www.ecpgr.cgiar.org/resources/germplasm-databases/list-of-germplasm-databases/crop-databases/crop-database-windows/lupin
https://www.ecpgr.cgiar.org/resources/germplasm-databases/list-of-germplasm-databases/crop-databases/crop-database-windows/lupin
https://www.ecpgr.cgiar.org/resources/germplasm-databases/list-of-germplasm-databases/crop-databases/crop-database-windows/lupin
https://www.ecpgr.cgiar.org/resources/germplasm-databases/list-of-germplasm-databases/crop-databases/crop-database-windows/lupin
https://www.ecpgr.cgiar.org/resources/germplasm-databases/list-of-germplasm-databases/crop-databases/crop-database-windows/lupin
https://www.ecpgr.cgiar.org/resources/germplasm-databases/list-of-germplasm-databases/crop-databases/crop-database-windows/lupin
https://www.ecpgr.cgiar.org/resources/germplasm-databases/list-of-germplasm-databases/crop-databases/crop-database-windows/lupin
https://www.ecpgr.cgiar.org/resources/germplasm-databases/list-of-germplasm-databases/crop-databases/crop-database-windows/lupin
https://www.ecpgr.cgiar.org/resources/germplasm-databases/list-of-germplasm-databases/crop-databases/crop-database-windows/lupin
https://www.ecpgr.cgiar.org/resources/germplasm-databases/list-of-germplasm-databases/crop-databases/crop-database-windows/lupin
https://www.ecpgr.cgiar.org/resources/germplasm-databases/list-of-germplasm-databases/crop-databases/crop-database-windows/lupin
https://www.ecpgr.cgiar.org/resources/germplasm-databases/list-of-germplasm-databases/crop-databases/crop-database-windows/lupin
https://www.ecpgr.cgiar.org/resources/germplasm-databases/list-of-germplasm-databases/crop-databases/crop-database-windows/lupin
https://www.ecpgr.cgiar.org/resources/germplasm-databases/list-of-germplasm-databases/crop-databases/crop-database-windows/lupin
https://www.ecpgr.cgiar.org/resources/germplasm-databases/list-of-germplasm-databases/crop-databases/crop-database-windows/lupin
https://www.ecpgr.cgiar.org/resources/germplasm-databases/list-of-germplasm-databases/crop-databases/crop-database-windows/lupin
https://www.ecpgr.cgiar.org/resources/germplasm-databases/list-of-germplasm-databases/crop-databases/crop-database-windows/lupin
https://www.ecpgr.cgiar.org/resources/germplasm-databases/list-of-germplasm-databases/crop-databases/crop-database-windows/lupin
https://www.ecpgr.cgiar.org/resources/germplasm-databases/list-of-germplasm-databases/crop-databases/crop-database-windows/lupin
https://www.ecpgr.cgiar.org/resources/germplasm-databases/list-of-germplasm-databases/crop-databases/crop-database-windows/lupin
https://www.ecpgr.cgiar.org/resources/germplasm-databases/list-of-germplasm-databases/crop-databases/crop-database-windows/lupin
https://www.ecpgr.cgiar.org/resources/germplasm-databases/list-of-germplasm-databases/crop-databases/crop-database-windows/lupin
https://www.ecpgr.cgiar.org/resources/germplasm-databases/list-of-germplasm-databases/crop-databases/crop-database-windows/lupin
https://www.ecpgr.cgiar.org/resources/germplasm-databases/list-of-germplasm-databases/crop-databases/crop-database-windows/lupin
https://www.ecpgr.cgiar.org/resources/germplasm-databases/list-of-germplasm-databases/crop-databases/crop-database-windows/lupin
https://www.ecpgr.cgiar.org/resources/germplasm-databases/list-of-germplasm-databases/crop-databases/crop-database-windows/lupin

https://www.ecpgr.cgiar.org/resources/germplasm-databases/list-of-germplasm-databases/crop-databases/crop-database-windows/lupin
https://www.ecpgr.cgiar.org/resources/germplasm-databases/list-of-germplasm-databases/crop-databases/crop-database-windows/lupin
https://www.ecpgr.cgiar.org/resources/germplasm-databases/list-of-germplasm-databases/crop-databases/crop-database-windows/lupin
https://www.ecpgr.cgiar.org/resources/germplasm-databases/list-of-germplasm-databases/crop-databases/crop-database-windows/lupin
https://www.genesys-pgr.org/a/bd3c4a4b-ffb1-4677-b1d4-1e5eb126d8de
https://www.genesys-pgr.org/a/caf1f0af-ab97-418b-b674-6892114b868f
https://www.genesys-pgr.org/a/e3716f79-06ff-4421-816e-286b01c91e2e
https://www.genesys-pgr.org/a/07e7d83b-d40f-4613-be20-9e17c0ede189
https://www.genesys-pgr.org/a/17a84c9f-3176-4df3-8fa0-0c9e3d75ca43
https://www.genesys-pgr.org/a/846df4ca-cf96-4680-b119-3412956f06b1
https://www.genesys-pgr.org/a/9c6a8124-b796-47da-9c0f-ceb164784e45
https://www.genesys-pgr.org/a/bb57b4eb-657b-4efa-95ce-b29f3490db3d
https://www.genesys-pgr.org/a/a248b105-295d-4ad0-990d-e9edea510f7f
https://www.genesys-pgr.org/a/10b91a70-7eb8-4fe9-b7e2-121beadb9c94
https://www.genesys-pgr.org/a/9d54fe05-2fce-4737-86dc-fa0b54dfd643
https://www.genesys-pgr.org/a/c65365e6-bc30-4dbc-accd-95aa1c4fe42a
https://www.genesys-pgr.org/a/baa57cfc-875b-4418-9214-a0d13ddae832
https://www.genesys-pgr.org/a/b4c86909-fb82-497c-9963-aa970b2d2133
https://www.ecpgr.cgiar.org/resources/germplasm-databases/list-of-germplasm-databases/crop-databases/crop-database-windows/lupin
https://www.ecpgr.cgiar.org/resources/germplasm-databases/list-of-germplasm-databases/crop-databases/crop-database-windows/lupin
https://www.genesys-pgr.org/a/71f8a976-0ca4-4dbf-ba97-c7f96a589d0b
https://www.genesys-pgr.org/a/2fb09cb4-2f6a-44e6-9d1f-2a1c7025cc5c
https://www.genesys-pgr.org/a/7ac07a5a-3a28-40b0-b1eb-675703b4a142
https://www.genesys-pgr.org/a/93b8112b-883a-481e-bcd9-5783f6406747
https://www.genesys-pgr.org/a/3ed9f6ef-b451-46bc-aba6-657d6308aaab
https://www.ecpgr.cgiar.org/resources/germplasm-databases/list-of-germplasm-databases/crop-databases/crop-database-windows/lupin
https://www.ecpgr.cgiar.org/resources/germplasm-databases/list-of-germplasm-databases/crop-databases/crop-database-windows/lupin
https://www.ecpgr.cgiar.org/resources/germplasm-databases/list-of-germplasm-databases/crop-databases/crop-database-windows/lupin
https://www.ecpgr.cgiar.org/resources/germplasm-databases/list-of-germplasm-databases/crop-databases/crop-database-windows/lupin
https://www.ecpgr.cgiar.org/resources/germplasm-databases/list-of-germplasm-databases/crop-databases/crop-database-windows/lupin
https://www.ecpgr.cgiar.org/resources/germplasm-databases/list-of-germplasm-databases/crop-databases/crop-database-windows/lupin
https://www.ecpgr.cgiar.org/resources/germplasm-databases/list-of-germplasm-databases/crop-databases/crop-database-windows/lupin
https://www.ecpgr.cgiar.org/resources/germplasm-databases/list-of-germplasm-databases/crop-databases/crop-database-windows/lupin
https://www.ecpgr.cgiar.org/resources/germplasm-databases/list-of-germplasm-databases/crop-databases/crop-database-windows/lupin
https://www.ecpgr.cgiar.org/resources/germplasm-databases/list-of-germplasm-databases/crop-databases/crop-database-windows/lupin
https://www.ecpgr.cgiar.org/resources/germplasm-databases/list-of-germplasm-databases/crop-databases/crop-database-windows/lupin
https://www.ecpgr.cgiar.org/resources/germplasm-databases/list-of-germplasm-databases/crop-databases/crop-database-windows/lupin
https://www.ecpgr.cgiar.org/resources/germplasm-databases/list-of-germplasm-databases/crop-databases/crop-database-windows/lupin
https://www.ecpgr.cgiar.org/resources/germplasm-databases/list-of-germplasm-databases/crop-databases/crop-database-windows/lupin
https://www.ecpgr.cgiar.org/resources/germplasm-databases/list-of-germplasm-databases/crop-databases/crop-database-windows/lupin
https://www.ecpgr.cgiar.org/resources/germplasm-databases/list-of-germplasm-databases/crop-databases/crop-database-windows/lupin
https://www.ecpgr.cgiar.org/resources/germplasm-databases/list-of-germplasm-databases/crop-databases/crop-database-windows/lupin
https://www.ecpgr.cgiar.org/resources/germplasm-databases/list-of-germplasm-databases/crop-databases/crop-database-windows/lupin
https://www.ecpgr.cgiar.org/resources/germplasm-databases/list-of-germplasm-databases/crop-databases/crop-database-windows/lupin
https://www.ecpgr.cgiar.org/resources/germplasm-databases/list-of-germplasm-databases/crop-databases/crop-database-windows/lupin
https://www.ecpgr.cgiar.org/resources/germplasm-databases/list-of-germplasm-databases/crop-databases/crop-database-windows/lupin
https://www.ecpgr.cgiar.org/resources/germplasm-databases/list-of-germplasm-databases/crop-databases/crop-database-windows/lupin
https://www.ecpgr.cgiar.org/resources/germplasm-databases/list-of-germplasm-databases/crop-databases/crop-database-windows/lupin
https://www.ecpgr.cgiar.org/resources/germplasm-databases/list-of-germplasm-databases/crop-databases/crop-database-windows/lupin
https://www.ecpgr.cgiar.org/resources/germplasm-databases/list-of-germplasm-databases/crop-databases/crop-database-windows/lupin
https://www.ecpgr.cgiar.org/resources/germplasm-databases/list-of-germplasm-databases/crop-databases/crop-database-windows/lupin

https://www.ecpgr.cgiar.org/resources/germplasm-databases/list-of-germplasm-databases/crop-databases/crop-database-windows/lupin
https://www.ecpgr.cgiar.org/resources/germplasm-databases/list-of-germplasm-databases/crop-databases/crop-database-windows/lupin
https://www.ecpgr.cgiar.org/resources/germplasm-databases/list-of-germplasm-databases/crop-databases/crop-database-windows/lupin
https://www.ecpgr.cgiar.org/resources/germplasm-databases/list-of-germplasm-databases/crop-databases/crop-database-windows/lupin
https://www.ecpgr.cgiar.org/resources/germplasm-databases/list-of-germplasm-databases/crop-databases/crop-database-windows/lupin
https://www.ecpgr.cgiar.org/resources/germplasm-databases/list-of-germplasm-databases/crop-databases/crop-database-windows/lupin
https://www.ecpgr.cgiar.org/resources/germplasm-databases/list-of-germplasm-databases/crop-databases/crop-database-windows/lupin
https://www.ecpgr.cgiar.org/resources/germplasm-databases/list-of-germplasm-databases/crop-databases/crop-database-windows/lupin
https://www.ecpgr.cgiar.org/resources/germplasm-databases/list-of-germplasm-databases/crop-databases/crop-database-windows/lupin
https://www.ecpgr.cgiar.org/resources/germplasm-databases/list-of-germplasm-databases/crop-databases/crop-database-windows/lupin
https://www.ecpgr.cgiar.org/resources/germplasm-databases/list-of-germplasm-databases/crop-databases/crop-database-windows/lupin
https://www.ecpgr.cgiar.org/resources/germplasm-databases/list-of-germplasm-databases/crop-databases/crop-database-windows/lupin
https://www.ecpgr.cgiar.org/resources/germplasm-databases/list-of-germplasm-databases/crop-databases/crop-database-windows/lupin
https://www.ecpgr.cgiar.org/resources/germplasm-databases/list-of-germplasm-databases/crop-databases/crop-database-windows/lupin
https://www.ecpgr.cgiar.org/resources/germplasm-databases/list-of-germplasm-databases/crop-databases/crop-database-windows/lupin
https://www.ecpgr.cgiar.org/resources/germplasm-databases/list-of-germplasm-databases/crop-databases/crop-database-windows/lupin
https://www.ecpgr.cgiar.org/resources/germplasm-databases/list-of-germplasm-databases/crop-databases/crop-database-windows/lupin
https://www.ecpgr.cgiar.org/resources/germplasm-databases/list-of-germplasm-databases/crop-databases/crop-database-windows/lupin
https://www.ecpgr.cgiar.org/resources/germplasm-databases/list-of-germplasm-databases/crop-databases/crop-database-windows/lupin
https://www.ecpgr.cgiar.org/resources/germplasm-databases/list-of-germplasm-databases/crop-databases/crop-database-windows/lupin
https://www.ecpgr.cgiar.org/resources/germplasm-databases/list-of-germplasm-databases/crop-databases/crop-database-windows/lupin
https://www.ecpgr.cgiar.org/resources/germplasm-databases/list-of-germplasm-databases/crop-databases/crop-database-windows/lupin
https://www.ecpgr.cgiar.org/resources/germplasm-databases/list-of-germplasm-databases/crop-databases/crop-database-windows/lupin
https://www.ecpgr.cgiar.org/resources/germplasm-databases/list-of-germplasm-databases/crop-databases/crop-database-windows/lupin
https://www.ecpgr.cgiar.org/resources/germplasm-databases/list-of-germplasm-databases/crop-databases/crop-database-windows/lupin
https://www.ecpgr.cgiar.org/resources/germplasm-databases/list-of-germplasm-databases/crop-databases/crop-database-windows/lupin
https://www.ecpgr.cgiar.org/resources/germplasm-databases/list-of-germplasm-databases/crop-databases/crop-database-windows/lupin
https://www.ecpgr.cgiar.org/resources/germplasm-databases/list-of-germplasm-databases/crop-databases/crop-database-windows/lupin
https://www.ecpgr.cgiar.org/resources/germplasm-databases/list-of-germplasm-databases/crop-databases/crop-database-windows/lupin
https://www.ecpgr.cgiar.org/resources/germplasm-databases/list-of-germplasm-databases/crop-databases/crop-database-windows/lupin
https://www.ecpgr.cgiar.org/resources/germplasm-databases/list-of-germplasm-databases/crop-databases/crop-database-windows/lupin
https://www.ecpgr.cgiar.org/resources/germplasm-databases/list-of-germplasm-databases/crop-databases/crop-database-windows/lupin
https://www.ecpgr.cgiar.org/resources/germplasm-databases/list-of-germplasm-databases/crop-databases/crop-database-windows/lupin
https://www.ecpgr.cgiar.org/resources/germplasm-databases/list-of-germplasm-databases/crop-databases/crop-database-windows/lupin
https://www.ecpgr.cgiar.org/resources/germplasm-databases/list-of-germplasm-databases/crop-databases/crop-database-windows/lupin
https://www.ecpgr.cgiar.org/resources/germplasm-databases/list-of-germplasm-databases/crop-databases/crop-database-windows/lupin
https://www.ecpgr.cgiar.org/resources/germplasm-databases/list-of-germplasm-databases/crop-databases/crop-database-windows/lupin
https://www.ecpgr.cgiar.org/resources/germplasm-databases/list-of-germplasm-databases/crop-databases/crop-database-windows/lupin
https://www.ecpgr.cgiar.org/resources/germplasm-databases/list-of-germplasm-databases/crop-databases/crop-database-windows/lupin
https://www.genesys-pgr.org/a/298a98da-8bf1-45f1-b2d9-ec77d733235e
https://www.genesys-pgr.org/a/2f0acc9d-382c-4d6a-b687-ee7c0213fb3a
https://www.genesys-pgr.org/a/906e62e0-ef8a-4ac2-b701-eeb74f865c7b
https://www.ecpgr.cgiar.org/resources/germplasm-databases/list-of-germplasm-databases/crop-databases/crop-database-windows/lupin
https://www.ecpgr.cgiar.org/resources/germplasm-databases/list-of-germplasm-databases/crop-databases/crop-database-windows/lupin
https://www.ecpgr.cgiar.org/resources/germplasm-databases/list-of-germplasm-databases/crop-databases/crop-database-windows/lupin
https://www.ecpgr.cgiar.org/resources/germplasm-databases/list-of-germplasm-databases/crop-databases/crop-database-windows/lupin
https://www.ecpgr.cgiar.org/resources/germplasm-databases/list-of-germplasm-databases/crop-databases/crop-database-windows/lupin
https://www.ecpgr.cgiar.org/resources/germplasm-databases/list-of-germplasm-databases/crop-databases/crop-database-windows/lupin
https://www.ecpgr.cgiar.org/resources/germplasm-databases/list-of-germplasm-databases/crop-databases/crop-database-windows/lupin
https://www.ecpgr.cgiar.org/resources/germplasm-databases/list-of-germplasm-databases/crop-databases/crop-database-windows/lupin
https://www.ecpgr.cgiar.org/resources/germplasm-databases/list-of-germplasm-databases/crop-databases/crop-database-windows/lupin
https://www.ecpgr.cgiar.org/resources/germplasm-databases/list-of-germplasm-databases/crop-databases/crop-database-windows/lupin

https://www.ecpgr.cgiar.org/resources/germplasm-databases/list-of-germplasm-databases/crop-databases/crop-database-windows/lupin
https://www.ecpgr.cgiar.org/resources/germplasm-databases/list-of-germplasm-databases/crop-databases/crop-database-windows/lupin
https://www.ecpgr.cgiar.org/resources/germplasm-databases/list-of-germplasm-databases/crop-databases/crop-database-windows/lupin
https://www.ecpgr.cgiar.org/resources/germplasm-databases/list-of-germplasm-databases/crop-databases/crop-database-windows/lupin
https://www.ecpgr.cgiar.org/resources/germplasm-databases/list-of-germplasm-databases/crop-databases/crop-database-windows/lupin
https://www.ecpgr.cgiar.org/resources/germplasm-databases/list-of-germplasm-databases/crop-databases/crop-database-windows/lupin
https://www.ecpgr.cgiar.org/resources/germplasm-databases/list-of-germplasm-databases/crop-databases/crop-database-windows/lupin
https://www.ecpgr.cgiar.org/resources/germplasm-databases/list-of-germplasm-databases/crop-databases/crop-database-windows/lupin
https://www.ecpgr.cgiar.org/resources/germplasm-databases/list-of-germplasm-databases/crop-databases/crop-database-windows/lupin
https://www.ecpgr.cgiar.org/resources/germplasm-databases/list-of-germplasm-databases/crop-databases/crop-database-windows/lupin
https://www.ecpgr.cgiar.org/resources/germplasm-databases/list-of-germplasm-databases/crop-databases/crop-database-windows/lupin
https://www.ecpgr.cgiar.org/resources/germplasm-databases/list-of-germplasm-databases/crop-databases/crop-database-windows/lupin
https://www.ecpgr.cgiar.org/resources/germplasm-databases/list-of-germplasm-databases/crop-databases/crop-database-windows/lupin
https://www.ecpgr.cgiar.org/resources/germplasm-databases/list-of-germplasm-databases/crop-databases/crop-database-windows/lupin
https://www.ecpgr.cgiar.org/resources/germplasm-databases/list-of-germplasm-databases/crop-databases/crop-database-windows/lupin
https://www.ecpgr.cgiar.org/resources/germplasm-databases/list-of-germplasm-databases/crop-databases/crop-database-windows/lupin
https://www.ecpgr.cgiar.org/resources/germplasm-databases/list-of-germplasm-databases/crop-databases/crop-database-windows/lupin
https://www.ecpgr.cgiar.org/resources/germplasm-databases/list-of-germplasm-databases/crop-databases/crop-database-windows/lupin
https://www.ecpgr.cgiar.org/resources/germplasm-databases/list-of-germplasm-databases/crop-databases/crop-database-windows/lupin
https://www.ecpgr.cgiar.org/resources/germplasm-databases/list-of-germplasm-databases/crop-databases/crop-database-windows/lupin
https://www.ecpgr.cgiar.org/resources/germplasm-databases/list-of-germplasm-databases/crop-databases/crop-database-windows/lupin
https://www.ecpgr.cgiar.org/resources/germplasm-databases/list-of-germplasm-databases/crop-databases/crop-database-windows/lupin
https://www.ecpgr.cgiar.org/resources/germplasm-databases/list-of-germplasm-databases/crop-databases/crop-database-windows/lupin
https://www.ecpgr.cgiar.org/resources/germplasm-databases/list-of-germplasm-databases/crop-databases/crop-database-windows/lupin
https://www.ecpgr.cgiar.org/resources/germplasm-databases/list-of-germplasm-databases/crop-databases/crop-database-windows/lupin
https://www.ecpgr.cgiar.org/resources/germplasm-databases/list-of-germplasm-databases/crop-databases/crop-database-windows/lupin
https://www.ecpgr.cgiar.org/resources/germplasm-databases/list-of-germplasm-databases/crop-databases/crop-database-windows/lupin
https://www.ecpgr.cgiar.org/resources/germplasm-databases/list-of-germplasm-databases/crop-databases/crop-database-windows/lupin
https://www.ecpgr.cgiar.org/resources/germplasm-databases/list-of-germplasm-databases/crop-databases/crop-database-windows/lupin
https://www.ecpgr.cgiar.org/resources/germplasm-databases/list-of-germplasm-databases/crop-databases/crop-database-windows/lupin
https://www.ecpgr.cgiar.org/resources/germplasm-databases/list-of-germplasm-databases/crop-databases/crop-database-windows/lupin
https://www.ecpgr.cgiar.org/resources/germplasm-databases/list-of-germplasm-databases/crop-databases/crop-database-windows/lupin
https://www.ecpgr.cgiar.org/resources/germplasm-databases/list-of-germplasm-databases/crop-databases/crop-database-windows/lupin
https://www.ecpgr.cgiar.org/resources/germplasm-databases/list-of-germplasm-databases/crop-databases/crop-database-windows/lupin
https://www.ecpgr.cgiar.org/resources/germplasm-databases/list-of-germplasm-databases/crop-databases/crop-database-windows/lupin
https://www.ecpgr.cgiar.org/resources/germplasm-databases/list-of-germplasm-databases/crop-databases/crop-database-windows/lupin
https://www.ecpgr.cgiar.org/resources/germplasm-databases/list-of-germplasm-databases/crop-databases/crop-database-windows/lupin
https://www.ecpgr.cgiar.org/resources/germplasm-databases/list-of-germplasm-databases/crop-databases/crop-database-windows/lupin
https://www.ecpgr.cgiar.org/resources/germplasm-databases/list-of-germplasm-databases/crop-databases/crop-database-windows/lupin
https://www.ecpgr.cgiar.org/resources/germplasm-databases/list-of-germplasm-databases/crop-databases/crop-database-windows/lupin
https://www.ecpgr.cgiar.org/resources/germplasm-databases/list-of-germplasm-databases/crop-databases/crop-database-windows/lupin
https://www.ecpgr.cgiar.org/resources/germplasm-databases/list-of-germplasm-databases/crop-databases/crop-database-windows/lupin
https://www.ecpgr.cgiar.org/resources/germplasm-databases/list-of-germplasm-databases/crop-databases/crop-database-windows/lupin
https://www.ecpgr.cgiar.org/resources/germplasm-databases/list-of-germplasm-databases/crop-databases/crop-database-windows/lupin
https://www.ecpgr.cgiar.org/resources/germplasm-databases/list-of-germplasm-databases/crop-databases/crop-database-windows/lupin
https://www.ecpgr.cgiar.org/resources/germplasm-databases/list-of-germplasm-databases/crop-databases/crop-database-windows/lupin
https://www.ecpgr.cgiar.org/resources/germplasm-databases/list-of-germplasm-databases/crop-databases/crop-database-windows/lupin
https://www.ecpgr.cgiar.org/resources/germplasm-databases/list-of-germplasm-databases/crop-databases/crop-database-windows/lupin
https://www.ecpgr.cgiar.org/resources/germplasm-databases/list-of-germplasm-databases/crop-databases/crop-database-windows/lupin
https://www.ecpgr.cgiar.org/resources/germplasm-databases/list-of-germplasm-databases/crop-databases/crop-database-windows/lupin
https://www.genesys-pgr.org/a/3d51b65a-0b0c-4b85-a113-304b9ffed147
https://www.genesys-pgr.org/a/50ba925c-722f-4ef7-ade5-81379d47c104

https://www.ecpgr.cgiar.org/resources/germplasm-databases/list-of-germplasm-databases/crop-databases/crop-database-windows/lupin
https://www.ecpgr.cgiar.org/resources/germplasm-databases/list-of-germplasm-databases/crop-databases/crop-database-windows/lupin
https://www.ecpgr.cgiar.org/resources/germplasm-databases/list-of-germplasm-databases/crop-databases/crop-database-windows/lupin
https://www.ecpgr.cgiar.org/resources/germplasm-databases/list-of-germplasm-databases/crop-databases/crop-database-windows/lupin
https://www.ecpgr.cgiar.org/resources/germplasm-databases/list-of-germplasm-databases/crop-databases/crop-database-windows/lupin
https://www.ecpgr.cgiar.org/resources/germplasm-databases/list-of-germplasm-databases/crop-databases/crop-database-windows/lupin
https://www.ecpgr.cgiar.org/resources/germplasm-databases/list-of-germplasm-databases/crop-databases/crop-database-windows/lupin
https://www.genesys-pgr.org/a/bd9e5f94-35b8-4845-8e83-ec46641c1457
https://www.genesys-pgr.org/a/ad266428-a9b7-461c-81c1-7dba7d70aaed
https://www.ecpgr.cgiar.org/resources/germplasm-databases/list-of-germplasm-databases/crop-databases/crop-database-windows/lupin
https://www.ecpgr.cgiar.org/resources/germplasm-databases/list-of-germplasm-databases/crop-databases/crop-database-windows/lupin
https://www.ecpgr.cgiar.org/resources/germplasm-databases/list-of-germplasm-databases/crop-databases/crop-database-windows/lupin
https://www.ecpgr.cgiar.org/resources/germplasm-databases/list-of-germplasm-databases/crop-databases/crop-database-windows/lupin
https://www.ecpgr.cgiar.org/resources/germplasm-databases/list-of-germplasm-databases/crop-databases/crop-database-windows/lupin
https://www.ecpgr.cgiar.org/resources/germplasm-databases/list-of-germplasm-databases/crop-databases/crop-database-windows/lupin
https://www.ecpgr.cgiar.org/resources/germplasm-databases/list-of-germplasm-databases/crop-databases/crop-database-windows/lupin
https://www.ecpgr.cgiar.org/resources/germplasm-databases/list-of-germplasm-databases/crop-databases/crop-database-windows/lupin
https://www.ecpgr.cgiar.org/resources/germplasm-databases/list-of-germplasm-databases/crop-databases/crop-database-windows/lupin
https://www.ecpgr.cgiar.org/resources/germplasm-databases/list-of-germplasm-databases/crop-databases/crop-database-windows/lupin
https://www.ecpgr.cgiar.org/resources/germplasm-databases/list-of-germplasm-databases/crop-databases/crop-database-windows/lupin
https://www.ecpgr.cgiar.org/resources/germplasm-databases/list-of-germplasm-databases/crop-databases/crop-database-windows/lupin
https://www.ecpgr.cgiar.org/resources/germplasm-databases/list-of-germplasm-databases/crop-databases/crop-database-windows/lupin
https://www.ecpgr.cgiar.org/resources/germplasm-databases/list-of-germplasm-databases/crop-databases/crop-database-windows/lupin
https://www.ecpgr.cgiar.org/resources/germplasm-databases/list-of-germplasm-databases/crop-databases/crop-database-windows/lupin
https://www.ecpgr.cgiar.org/resources/germplasm-databases/list-of-germplasm-databases/crop-databases/crop-database-windows/lupin
https://www.ecpgr.cgiar.org/resources/germplasm-databases/list-of-germplasm-databases/crop-databases/crop-database-windows/lupin
https://www.ecpgr.cgiar.org/resources/germplasm-databases/list-of-germplasm-databases/crop-databases/crop-database-windows/lupin
https://www.ecpgr.cgiar.org/resources/germplasm-databases/list-of-germplasm-databases/crop-databases/crop-database-windows/lupin
https://www.ecpgr.cgiar.org/resources/germplasm-databases/list-of-germplasm-databases/crop-databases/crop-database-windows/lupin
https://www.ecpgr.cgiar.org/resources/germplasm-databases/list-of-germplasm-databases/crop-databases/crop-database-windows/lupin
https://www.ecpgr.cgiar.org/resources/germplasm-databases/list-of-germplasm-databases/crop-databases/crop-database-windows/lupin
https://www.ecpgr.cgiar.org/resources/germplasm-databases/list-of-germplasm-databases/crop-databases/crop-database-windows/lupin
https://www.ecpgr.cgiar.org/resources/germplasm-databases/list-of-germplasm-databases/crop-databases/crop-database-windows/lupin
https://www.ecpgr.cgiar.org/resources/germplasm-databases/list-of-germplasm-databases/crop-databases/crop-database-windows/lupin
https://www.ecpgr.cgiar.org/resources/germplasm-databases/list-of-germplasm-databases/crop-databases/crop-database-windows/lupin
https://www.ecpgr.cgiar.org/resources/germplasm-databases/list-of-germplasm-databases/crop-databases/crop-database-windows/lupin
https://www.ecpgr.cgiar.org/resources/germplasm-databases/list-of-germplasm-databases/crop-databases/crop-database-windows/lupin
https://www.ecpgr.cgiar.org/resources/germplasm-databases/list-of-germplasm-databases/crop-databases/crop-database-windows/lupin
https://www.ecpgr.cgiar.org/resources/germplasm-databases/list-of-germplasm-databases/crop-databases/crop-database-windows/lupin
https://www.ecpgr.cgiar.org/resources/germplasm-databases/list-of-germplasm-databases/crop-databases/crop-database-windows/lupin
https://www.ecpgr.cgiar.org/resources/germplasm-databases/list-of-germplasm-databases/crop-databases/crop-database-windows/lupin
https://www.ecpgr.cgiar.org/resources/germplasm-databases/list-of-germplasm-databases/crop-databases/crop-database-windows/lupin
https://www.ecpgr.cgiar.org/resources/germplasm-databases/list-of-germplasm-databases/crop-databases/crop-database-windows/lupin
https://www.ecpgr.cgiar.org/resources/germplasm-databases/list-of-germplasm-databases/crop-databases/crop-database-windows/lupin
https://www.ecpgr.cgiar.org/resources/germplasm-databases/list-of-germplasm-databases/crop-databases/crop-database-windows/lupin
https://www.ecpgr.cgiar.org/resources/germplasm-databases/list-of-germplasm-databases/crop-databases/crop-database-windows/lupin
https://www.genesys-pgr.org/a/3ee571c2-2373-4703-8121-92803d08e88e
https://www.ecpgr.cgiar.org/resources/germplasm-databases/list-of-germplasm-databases/crop-databases/crop-database-windows/lupin
https://www.ecpgr.cgiar.org/resources/germplasm-databases/list-of-germplasm-databases/crop-databases/crop-database-windows/lupin
https://www.ecpgr.cgiar.org/resources/germplasm-databases/list-of-germplasm-databases/crop-databases/crop-database-windows/lupin
https://www.ecpgr.cgiar.org/resources/germplasm-databases/list-of-germplasm-databases/crop-databases/crop-database-windows/lupin
https://www.ecpgr.cgiar.org/resources/germplasm-databases/list-of-germplasm-databases/crop-databases/crop-database-windows/lupin

https://www.ecpgr.cgiar.org/resources/germplasm-databases/list-of-germplasm-databases/crop-databases/crop-database-windows/lupin

https://www.genesys-pgr.org/a/dacf3db2-ae4b-4ebb-87b6-6f67f2d6a8c6

https://www.genesys-pgr.org/a/200e0460-0cca-4e7d-9e9a-f2d636b8f89b

https://www.genesys-pgr.org/a/03da2ffe-b8eb-43a4-b56a-fef5be3bac6a

https://www.genesys-pgr.org/a/d27af507-1dc5-43fe-a908-9208f8cda094

https://www.genesys-pgr.org/a/d351eebc-229c-4d04-b3d6-75951a599f25

https://www.genesys-pgr.org/a/486bb5e3-cbc8-44d8-add1-fa2a0f4284ba

https://www.genesys-pgr.org/a/97f574fc-78e9-413b-b9d7-f3aea77da3bb

https://www.genesys-pgr.org/a/bed9e710-0075-452d-afbc-ae27bed35652

https://www.genesys-pgr.org/a/da7f5a86-e7f0-4901-959a-e7da459ac343

https://www.genesys-pgr.org/a/b23edd02-9df2-4b29-9bb8-84f3cf49b5ae

https://www.ecpgr.cgiar.org/resources/germplasm-databases/list-of-germplasm-databases/crop-databases/crop-database-windows/lupin

https://www.ecpgr.cgiar.org/resources/germplasm-databases/list-of-germplasm-databases/crop-databases/crop-database-windows/lupin

https://www.ecpgr.cgiar.org/resources/germplasm-databases/list-of-germplasm-databases/crop-databases/crop-database-windows/lupin

https://www.ecpgr.cgiar.org/resources/germplasm-databases/list-of-germplasm-databases/crop-databases/crop-database-windows/lupin

https://www.ecpgr.cgiar.org/resources/germplasm-databases/list-of-germplasm-databases/crop-databases/crop-database-windows/lupin

https://www.ecpgr.cgiar.org/resources/germplasm-databases/list-of-germplasm-databases/crop-databases/crop-database-windows/lupin

https://www.ecpgr.cgiar.org/resources/germplasm-databases/list-of-germplasm-databases/crop-databases/crop-database-windows/lupin

https://www.ecpgr.cgiar.org/resources/germplasm-databases/list-of-germplasm-databases/crop-databases/crop-database-windows/lupin

https://www.ecpgr.cgiar.org/resources/germplasm-databases/list-of-germplasm-databases/crop-databases/crop-database-windows/lupin

https://www.ecpgr.cgiar.org/resources/germplasm-databases/list-of-germplasm-databases/crop-databases/crop-database-windows/lupin

https://www.ecpgr.cgiar.org/resources/germplasm-databases/list-of-germplasm-databases/crop-databases/crop-database-windows/lupin

https://www.ecpgr.cgiar.org/resources/germplasm-databases/list-of-germplasm-databases/crop-databases/crop-database-windows/lupin

https://www.ecpgr.cgiar.org/resources/germplasm-databases/list-of-germplasm-databases/crop-databases/crop-database-windows/lupin

https://www.genesys-pgr.org/a/cba55f73-e52d-4059-86cd-6eb9c7c1f000

https://www.genesys-pgr.org/a/44cdae40-95c9-4c99-91dc-97167fef91be

https://www.genesys-pgr.org/a/8a5a987e-a1c8-4011-8115-ddeb4d333097

https://www.genesys-pgr.org/a/11aee2d4-7987-40e9-878c-479207d3aba3

https://www.genesys-pgr.org/a/550fea10-e198-478e-aa9a-e1851daffc83

https://www.ecpgr.cgiar.org/resources/germplasm-databases/list-of-germplasm-databases/crop-databases/crop-database-windows/lupin

https://www.ecpgr.cgiar.org/resources/germplasm-databases/list-of-germplasm-databases/crop-databases/crop-database-windows/lupin

https://www.ecpgr.cgiar.org/resources/germplasm-databases/list-of-germplasm-databases/crop-databases/crop-database-windows/lupin

https://www.ecpgr.cgiar.org/resources/germplasm-databases/list-of-germplasm-databases/crop-databases/crop-database-windows/lupin

https://www.ecpgr.cgiar.org/resources/germplasm-databases/list-of-germplasm-databases/crop-databases/crop-database-windows/lupin

https://www.ecpgr.cgiar.org/resources/germplasm-databases/list-of-germplasm-databases/crop-databases/crop-database-windows/lupin

https://www.ecpgr.cgiar.org/resources/germplasm-databases/list-of-germplasm-databases/crop-databases/crop-database-windows/lupin

https://www.ecpgr.cgiar.org/resources/germplasm-databases/list-of-germplasm-databases/crop-databases/crop-database-windows/lupin

https://www.ecpgr.cgiar.org/resources/germplasm-databases/list-of-germplasm-databases/crop-databases/crop-database-windows/lupin

https://www.ecpgr.cgiar.org/resources/germplasm-databases/list-of-germplasm-databases/crop-databases/crop-database-windows/lupin

https://www.ecpgr.cgiar.org/resources/germplasm-databases/list-of-germplasm-databases/crop-databases/crop-database-windows/lupin

https://www.ecpgr.cgiar.org/resources/germplasm-databases/list-of-germplasm-databases/crop-databases/crop-database-windows/lupin

https://www.ecpgr.cgiar.org/resources/germplasm-databases/list-of-germplasm-databases/crop-databases/crop-database-windows/lupin

https://www.ecpgr.cgiar.org/resources/germplasm-databases/list-of-germplasm-databases/crop-databases/crop-database-windows/lupin

https://www.ecpgr.cgiar.org/resources/germplasm-databases/list-of-germplasm-databases/crop-databases/crop-database-windows/lupin

https://www.ecpgr.cgiar.org/resources/germplasm-databases/list-of-germplasm-databases/crop-databases/crop-database-windows/lupin

https://www.ecpgr.cgiar.org/resources/germplasm-databases/list-of-germplasm-databases/crop-databases/crop-database-windows/lupin

https://www.ecpgr.cgiar.org/resources/germplasm-databases/list-of-germplasm-databases/crop-databases/crop-database-windows/lupin

https://www.ecpgr.cgiar.org/resources/germplasm-databases/list-of-germplasm-databases/crop-databases/crop-database-windows/lupin

https://www.ecpgr.cgiar.org/resources/germplasm-databases/list-of-germplasm-databases/crop-databases/crop-database-windows/lupin

https://www.ecpgr.cgiar.org/resources/germplasm-databases/list-of-germplasm-databases/crop-databases/crop-database-windows/lupin

https://www.ecpgr.cgiar.org/resources/germplasm-databases/list-of-germplasm-databases/crop-databases/crop-database-windows/lupin

https://www.ecpgr.cgiar.org/resources/germplasm-databases/list-of-germplasm-databases/crop-databases/crop-database-windows/lupin

https://www.ecpgr.cgiar.org/resources/germplasm-databases/list-of-germplasm-databases/crop-databases/crop-database-windows/lupin
https://www.ecpgr.cgiar.org/resources/germplasm-databases/list-of-germplasm-databases/crop-databases/crop-database-windows/lupin
https://www.ecpgr.cgiar.org/resources/germplasm-databases/list-of-germplasm-databases/crop-databases/crop-database-windows/lupin
https://www.ecpgr.cgiar.org/resources/germplasm-databases/list-of-germplasm-databases/crop-databases/crop-database-windows/lupin
https://www.ecpgr.cgiar.org/resources/germplasm-databases/list-of-germplasm-databases/crop-databases/crop-database-windows/lupin
https://www.ecpgr.cgiar.org/resources/germplasm-databases/list-of-germplasm-databases/crop-databases/crop-database-windows/lupin
https://www.ecpgr.cgiar.org/resources/germplasm-databases/list-of-germplasm-databases/crop-databases/crop-database-windows/lupin
https://www.ecpgr.cgiar.org/resources/germplasm-databases/list-of-germplasm-databases/crop-databases/crop-database-windows/lupin
https://www.ecpgr.cgiar.org/resources/germplasm-databases/list-of-germplasm-databases/crop-databases/crop-database-windows/lupin
https://www.ecpgr.cgiar.org/resources/germplasm-databases/list-of-germplasm-databases/crop-databases/crop-database-windows/lupin
https://www.ecpgr.cgiar.org/resources/germplasm-databases/list-of-germplasm-databases/crop-databases/crop-database-windows/lupin
https://www.ecpgr.cgiar.org/resources/germplasm-databases/list-of-germplasm-databases/crop-databases/crop-database-windows/lupin
https://www.ecpgr.cgiar.org/resources/germplasm-databases/list-of-germplasm-databases/crop-databases/crop-database-windows/lupin
https://www.ecpgr.cgiar.org/resources/germplasm-databases/list-of-germplasm-databases/crop-databases/crop-database-windows/lupin
https://www.ecpgr.cgiar.org/resources/germplasm-databases/list-of-germplasm-databases/crop-databases/crop-database-windows/lupin
https://www.ecpgr.cgiar.org/resources/germplasm-databases/list-of-germplasm-databases/crop-databases/crop-database-windows/lupin
https://www.ecpgr.cgiar.org/resources/germplasm-databases/list-of-germplasm-databases/crop-databases/crop-database-windows/lupin
https://www.ecpgr.cgiar.org/resources/germplasm-databases/list-of-germplasm-databases/crop-databases/crop-database-windows/lupin
https://www.ecpgr.cgiar.org/resources/germplasm-databases/list-of-germplasm-databases/crop-databases/crop-database-windows/lupin
https://www.ecpgr.cgiar.org/resources/germplasm-databases/list-of-germplasm-databases/crop-databases/crop-database-windows/lupin
https://www.ecpgr.cgiar.org/resources/germplasm-databases/list-of-germplasm-databases/crop-databases/crop-database-windows/lupin
https://www.ecpgr.cgiar.org/resources/germplasm-databases/list-of-germplasm-databases/crop-databases/crop-database-windows/lupin
https://www.ecpgr.cgiar.org/resources/germplasm-databases/list-of-germplasm-databases/crop-databases/crop-database-windows/lupin
https://www.ecpgr.cgiar.org/resources/germplasm-databases/list-of-germplasm-databases/crop-databases/crop-database-windows/lupin
https://www.ecpgr.cgiar.org/resources/germplasm-databases/list-of-germplasm-databases/crop-databases/crop-database-windows/lupin
https://www.ecpgr.cgiar.org/resources/germplasm-databases/list-of-germplasm-databases/crop-databases/crop-database-windows/lupin
https://www.ecpgr.cgiar.org/resources/germplasm-databases/list-of-germplasm-databases/crop-databases/crop-database-windows/lupin
https://www.ecpgr.cgiar.org/resources/germplasm-databases/list-of-germplasm-databases/crop-databases/crop-database-windows/lupin
https://www.ecpgr.cgiar.org/resources/germplasm-databases/list-of-germplasm-databases/crop-databases/crop-database-windows/lupin
https://www.ecpgr.cgiar.org/resources/germplasm-databases/list-of-germplasm-databases/crop-databases/crop-database-windows/lupin
https://www.ecpgr.cgiar.org/resources/germplasm-databases/list-of-germplasm-databases/crop-databases/crop-database-windows/lupin
https://www.ecpgr.cgiar.org/resources/germplasm-databases/list-of-germplasm-databases/crop-databases/crop-database-windows/lupin
https://www.ecpgr.cgiar.org/resources/germplasm-databases/list-of-germplasm-databases/crop-databases/crop-database-windows/lupin
https://www.ecpgr.cgiar.org/resources/germplasm-databases/list-of-germplasm-databases/crop-databases/crop-database-windows/lupin
https://www.ecpgr.cgiar.org/resources/germplasm-databases/list-of-germplasm-databases/crop-databases/crop-database-windows/lupin
https://www.ecpgr.cgiar.org/resources/germplasm-databases/list-of-germplasm-databases/crop-databases/crop-database-windows/lupin
https://www.ecpgr.cgiar.org/resources/germplasm-databases/list-of-germplasm-databases/crop-databases/crop-database-windows/lupin
https://www.ecpgr.cgiar.org/resources/germplasm-databases/list-of-germplasm-databases/crop-databases/crop-database-windows/lupin
https://www.ecpgr.cgiar.org/resources/germplasm-databases/list-of-germplasm-databases/crop-databases/crop-database-windows/lupin
https://www.ecpgr.cgiar.org/resources/germplasm-databases/list-of-germplasm-databases/crop-databases/crop-database-windows/lupin
https://www.genesys-pgr.org/a/13f17d1e-1434-44b8-a166-575c6daae94b
https://www.ecpgr.cgiar.org/resources/germplasm-databases/list-of-germplasm-databases/crop-databases/crop-database-windows/lupin
https://www.ecpgr.cgiar.org/resources/germplasm-databases/list-of-germplasm-databases/crop-databases/crop-database-windows/lupin
https://www.ecpgr.cgiar.org/resources/germplasm-databases/list-of-germplasm-databases/crop-databases/crop-database-windows/lupin
https://www.ecpgr.cgiar.org/resources/germplasm-databases/list-of-germplasm-databases/crop-databases/crop-database-windows/lupin
https://www.ecpgr.cgiar.org/resources/germplasm-databases/list-of-germplasm-databases/crop-databases/crop-database-windows/lupin
https://www.ecpgr.cgiar.org/resources/germplasm-databases/list-of-germplasm-databases/crop-databases/crop-database-windows/lupin
https://www.ecpgr.cgiar.org/resources/germplasm-databases/list-of-germplasm-databases/crop-databases/crop-database-windows/lupin
https://www.ecpgr.cgiar.org/resources/germplasm-databases/list-of-germplasm-databases/crop-databases/crop-database-windows/lupin
https://www.ecpgr.cgiar.org/resources/germplasm-databases/list-of-germplasm-databases/crop-databases/crop-database-windows/lupin
https://www.ecpgr.cgiar.org/resources/germplasm-databases/list-of-germplasm-databases/crop-databases/crop-database-windows/lupin
https://www.ecpgr.cgiar.org/resources/germplasm-databases/list-of-germplasm-databases/crop-databases/crop-database-windows/lupin

https://www.ecpgr.cgiar.org/resources/germplasm-databases/list-of-germplasm-databases/crop-databases/crop-database-windows/lupin
https://www.ecpgr.cgiar.org/resources/germplasm-databases/list-of-germplasm-databases/crop-databases/crop-database-windows/lupin
https://www.ecpgr.cgiar.org/resources/germplasm-databases/list-of-germplasm-databases/crop-databases/crop-database-windows/lupin
https://www.ecpgr.cgiar.org/resources/germplasm-databases/list-of-germplasm-databases/crop-databases/crop-database-windows/lupin
https://www.ecpgr.cgiar.org/resources/germplasm-databases/list-of-germplasm-databases/crop-databases/crop-database-windows/lupin
https://www.ecpgr.cgiar.org/resources/germplasm-databases/list-of-germplasm-databases/crop-databases/crop-database-windows/lupin
https://www.ecpgr.cgiar.org/resources/germplasm-databases/list-of-germplasm-databases/crop-databases/crop-database-windows/lupin
https://www.ecpgr.cgiar.org/resources/germplasm-databases/list-of-germplasm-databases/crop-databases/crop-database-windows/lupin
https://www.ecpgr.cgiar.org/resources/germplasm-databases/list-of-germplasm-databases/crop-databases/crop-database-windows/lupin
https://www.ecpgr.cgiar.org/resources/germplasm-databases/list-of-germplasm-databases/crop-databases/crop-database-windows/lupin
https://www.ecpgr.cgiar.org/resources/germplasm-databases/list-of-germplasm-databases/crop-databases/crop-database-windows/lupin
https://www.ecpgr.cgiar.org/resources/germplasm-databases/list-of-germplasm-databases/crop-databases/crop-database-windows/lupin
https://www.ecpgr.cgiar.org/resources/germplasm-databases/list-of-germplasm-databases/crop-databases/crop-database-windows/lupin
https://www.ecpgr.cgiar.org/resources/germplasm-databases/list-of-germplasm-databases/crop-databases/crop-database-windows/lupin
https://www.ecpgr.cgiar.org/resources/germplasm-databases/list-of-germplasm-databases/crop-databases/crop-database-windows/lupin
https://www.ecpgr.cgiar.org/resources/germplasm-databases/list-of-germplasm-databases/crop-databases/crop-database-windows/lupin
https://www.ecpgr.cgiar.org/resources/germplasm-databases/list-of-germplasm-databases/crop-databases/crop-database-windows/lupin
https://www.ecpgr.cgiar.org/resources/germplasm-databases/list-of-germplasm-databases/crop-databases/crop-database-windows/lupin
https://www.ecpgr.cgiar.org/resources/germplasm-databases/list-of-germplasm-databases/crop-databases/crop-database-windows/lupin
https://www.ecpgr.cgiar.org/resources/germplasm-databases/list-of-germplasm-databases/crop-databases/crop-database-windows/lupin
https://www.ecpgr.cgiar.org/resources/germplasm-databases/list-of-germplasm-databases/crop-databases/crop-database-windows/lupin
https://www.ecpgr.cgiar.org/resources/germplasm-databases/list-of-germplasm-databases/crop-databases/crop-database-windows/lupin
https://www.ecpgr.cgiar.org/resources/germplasm-databases/list-of-germplasm-databases/crop-databases/crop-database-windows/lupin
https://www.ecpgr.cgiar.org/resources/germplasm-databases/list-of-germplasm-databases/crop-databases/crop-database-windows/lupin
https://www.ecpgr.cgiar.org/resources/germplasm-databases/list-of-germplasm-databases/crop-databases/crop-database-windows/lupin
https://www.ecpgr.cgiar.org/resources/germplasm-databases/list-of-germplasm-databases/crop-databases/crop-database-windows/lupin
https://www.ecpgr.cgiar.org/resources/germplasm-databases/list-of-germplasm-databases/crop-databases/crop-database-windows/lupin
https://www.ecpgr.cgiar.org/resources/germplasm-databases/list-of-germplasm-databases/crop-databases/crop-database-windows/lupin
https://www.ecpgr.cgiar.org/resources/germplasm-databases/list-of-germplasm-databases/crop-databases/crop-database-windows/lupin
https://www.ecpgr.cgiar.org/resources/germplasm-databases/list-of-germplasm-databases/crop-databases/crop-database-windows/lupin
https://www.ecpgr.cgiar.org/resources/germplasm-databases/list-of-germplasm-databases/crop-databases/crop-database-windows/lupin
https://www.ecpgr.cgiar.org/resources/germplasm-databases/list-of-germplasm-databases/crop-databases/crop-database-windows/lupin
https://www.ecpgr.cgiar.org/resources/germplasm-databases/list-of-germplasm-databases/crop-databases/crop-database-windows/lupin
https://www.ecpgr.cgiar.org/resources/germplasm-databases/list-of-germplasm-databases/crop-databases/crop-database-windows/lupin
https://www.ecpgr.cgiar.org/resources/germplasm-databases/list-of-germplasm-databases/crop-databases/crop-database-windows/lupin
https://www.ecpgr.cgiar.org/resources/germplasm-databases/list-of-germplasm-databases/crop-databases/crop-database-windows/lupin
https://www.ecpgr.cgiar.org/resources/germplasm-databases/list-of-germplasm-databases/crop-databases/crop-database-windows/lupin
https://www.ecpgr.cgiar.org/resources/germplasm-databases/list-of-germplasm-databases/crop-databases/crop-database-windows/lupin
https://www.ecpgr.cgiar.org/resources/germplasm-databases/list-of-germplasm-databases/crop-databases/crop-database-windows/lupin
https://www.ecpgr.cgiar.org/resources/germplasm-databases/list-of-germplasm-databases/crop-databases/crop-database-windows/lupin
https://www.ecpgr.cgiar.org/resources/germplasm-databases/list-of-germplasm-databases/crop-databases/crop-database-windows/lupin
https://www.ecpgr.cgiar.org/resources/germplasm-databases/list-of-germplasm-databases/crop-databases/crop-database-windows/lupin
https://www.ecpgr.cgiar.org/resources/germplasm-databases/list-of-germplasm-databases/crop-databases/crop-database-windows/lupin
https://www.ecpgr.cgiar.org/resources/germplasm-databases/list-of-germplasm-databases/crop-databases/crop-database-windows/lupin
https://www.ecpgr.cgiar.org/resources/germplasm-databases/list-of-germplasm-databases/crop-databases/crop-database-windows/lupin
https://www.ecpgr.cgiar.org/resources/germplasm-databases/list-of-germplasm-databases/crop-databases/crop-database-windows/lupin
https://www.ecpgr.cgiar.org/resources/germplasm-databases/list-of-germplasm-databases/crop-databases/crop-database-windows/lupin
https://www.ecpgr.cgiar.org/resources/germplasm-databases/list-of-germplasm-databases/crop-databases/crop-database-windows/lupin
https://www.ecpgr.cgiar.org/resources/germplasm-databases/list-of-germplasm-databases/crop-databases/crop-database-windows/lupin
https://www.ecpgr.cgiar.org/resources/germplasm-databases/list-of-germplasm-databases/crop-databases/crop-database-windows/lupin
https://www.ecpgr.cgiar.org/resources/germplasm-databases/list-of-germplasm-databases/crop-databases/crop-database-windows/lupin
https://www.ecpgr.cgiar.org/resources/germplasm-databases/list-of-germplasm-databases/crop-databases/crop-database-windows/lupin

https://www.ecpgr.cgiar.org/resources/germplasm-databases/list-of-germplasm-databases/crop-databases/crop-database-windows/lupin
https://www.ecpgr.cgiar.org/resources/germplasm-databases/list-of-germplasm-databases/crop-databases/crop-database-windows/lupin
https://www.ecpgr.cgiar.org/resources/germplasm-databases/list-of-germplasm-databases/crop-databases/crop-database-windows/lupin
https://www.ecpgr.cgiar.org/resources/germplasm-databases/list-of-germplasm-databases/crop-databases/crop-database-windows/lupin
https://www.genesys-pgr.org/a/151186c1-13f9-49cd-bc62-4120f624aafc
https://www.genesys-pgr.org/a/09faea17-3a5f-4a96-acb3-7136b5d9b5d4
https://www.genesys-pgr.org/a/8cb69691-1c57-4cd4-88bd-98f960be0f7b
https://www.genesys-pgr.org/a/cdaf0580-8e6b-4b45-a24a-ae36669b707a
https://www.genesys-pgr.org/a/db99d6d2-dc77-4ec9-958d-2248a09c46ff
https://www.genesys-pgr.org/a/db99d6d2-dc77-4ec9-958d-2248a09c46ff
https://www.genesys-pgr.org/a/586af7aa-cf6e-4ea1-ba0c-53fb5a7fa6c6
https://www.genesys-pgr.org/a/475af59e-43f6-47ab-9e88-d4a01de91b73
https://www.genesys-pgr.org/a/475af59e-43f6-47ab-9e88-d4a01de91b73
https://www.genesys-pgr.org/a/3d63fd4f-f5c0-4d33-9104-f85326a7d53d
https://www.genesys-pgr.org/a/259b1dc3-7388-4e0d-baca-cf40445613bb
https://www.genesys-pgr.org/a/3a719bca-9c20-4858-b800-3c4b7736a115
https://www.genesys-pgr.org/a/cfdd2483-4331-4e0b-9eb5-d87f3ce9c3a3
https://www.genesys-pgr.org/a/5ca98dcb-820d-4de8-8c1b-a2ae68939d1f
https://www.genesys-pgr.org/a/255d3008-d952-4911-b4f5-a788d47aa05d
https://www.genesys-pgr.org/a/d499fff4-1e6f-4853-8cf5-f927bce1d1dd
https://www.genesys-pgr.org/a/ba9c8643-3696-46ed-b138-88f0c62e3a74
https://www.genesys-pgr.org/a/e1e19390-40eb-4b0c-88ef-3629a6520460
https://www.genesys-pgr.org/a/488234b9-5000-4cbb-8680-5149fb0cf095
https://www.genesys-pgr.org/a/5928c3b7-8a73-4296-9db5-ffa5bd8ac369
https://www.genesys-pgr.org/a/bfcb09ab-bbd5-4470-b1ad-faddb8ad37ad
https://www.genesys-pgr.org/a/dbc145fe-e861-48e0-8c25-260d506f5e4f
https://www.genesys-pgr.org/a/d39d75f1-9e81-4e4b-8195-caa48da21b77
https://www.genesys-pgr.org/a/f1e4cc29-d643-490e-9939-f1459a569da5
https://www.genesys-pgr.org/a/27d17e62-9894-402b-ba7b-2bfbcddb883a
https://www.genesys-pgr.org/a/997f45e3-b547-4e01-a31d-24fda258b5a3
https://www.genesys-pgr.org/a/76880aea-4b95-4326-bacc-3997687ce59b
https://www.genesys-pgr.org/a/fd40a12c-b113-4439-b9b7-c929839afbf9
https://www.genesys-pgr.org/a/7a269568-3438-4561-907c-d5938a6fb326
https://www.genesys-pgr.org/a/f58ee57a-4740-4f5f-9534-e996e11e9d61
https://www.genesys-pgr.org/a/1e84a6ea-3592-4f70-9abb-21f261f4d5a5
https://www.genesys-pgr.org/a/91ac5910-eaff-432f-8b2c-4cd493cb7f41
https://www.genesys-pgr.org/a/54dadf82-3fe4-4125-ad26-44b4dd88b00f
https://www.genesys-pgr.org/a/2bcd5950-68ab-46a9-9c52-961577bb3cdd
https://www.genesys-pgr.org/a/b5a3c420-758b-4e90-8704-9a70e1bdf065
https://www.genesys-pgr.org/a/596f3981-d4cc-496a-9afa-316473f704ed
https://www.genesys-pgr.org/a/78afb009-5d18-4d03-bb2d-2bb314822fcd
https://www.genesys-pgr.org/a/47824be0-787e-4346-993e-fea72b58ac00
https://www.genesys-pgr.org/a/fc39fe42-cdae-4616-a7ea-68cb7d738f39

https://www.genesys-pgr.org/10.18730/VPGJU
https://www.genesys-pgr.org/10.18730/VZ6SP
https://www.genesys-pgr.org/10.18730/VPR8Q
https://www.genesys-pgr.org/10.18730/VPR4K
https://www.genesys-pgr.org/10.18730/VPPPA
https://www.genesys-pgr.org/10.18730/VP73X
https://www.genesys-pgr.org/10.18730/VPR2H
https://www.genesys-pgr.org/10.18730/VPPM8

https://www.genesys-pgr.org/10.18730/VP72W
https://www.genesys-pgr.org/10.18730/VPGK0
https://www.genesys-pgr.org/10.18730/VPGH=
https://www.genesys-pgr.org/10.18730/VPPQB
https://www.genesys-pgr.org/10.18730/VPGF~
https://www.genesys-pgr.org/10.18730/VPR5M
https://www.genesys-pgr.org/10.18730/VPR7P
https://www.genesys-pgr.org/10.18730/VP72W
https://www.genesys-pgr.org/10.18730/VZ6TQ
https://www.genesys-pgr.org/10.18730/VPRAS
https://www.genesys-pgr.org/10.18730/VP5NM
https://www.genesys-pgr.org/10.18730/VPPN9
https://www.genesys-pgr.org/10.18730/VPR6N
https://www.genesys-pgr.org/10.18730/YBKN3
https://www.genesys-pgr.org/10.18730/YBKP4

https://www.ecpgr.cgiar.org/resources/germplasm-databases/list-of-germplasm-databases/crop-databases/crop-database-windows/lupin

https://www.genesys-pgr.org/a/dfa3eabc-484a-40ec-a43e-f051afa5cb18
https://www.genesys-pgr.org/a/8c08bd9e-76a2-47b4-9bbd-ad400157fbdc
https://www.genesys-pgr.org/a/411b8ff4-19d0-4982-a9e6-395c136f0adb
https://www.genesys-pgr.org/a/125965b6-2e3e-43d6-92fd-42638f4bfc5c
https://www.genesys-pgr.org/a/c0ff1817-115a-4ba2-b91e-6e05c44bb9b4
https://www.genesys-pgr.org/a/ab1e8dd2-86fc-4650-93e4-7b2d02c16162
https://www.genesys-pgr.org/a/b05e664b-4d82-4147-b701-497be661e53c
https://www.genesys-pgr.org/a/639bd8a4-4a58-4a44-a040-b42f1d03e491
https://www.genesys-pgr.org/a/e473a086-8324-4e46-93c3-5dbc70177dca
https://www.genesys-pgr.org/a/aab991e9-9f34-4615-9bd8-5931d9e7e3db
https://www.genesys-pgr.org/a/8a6d49d1-2258-48de-8c69-5c9856691195
https://www.genesys-pgr.org/a/593e1451-bf3a-4960-acea-e3475fe3dc9a
https://www.genesys-pgr.org/a/51f209f1-d422-4d24-b62b-3864bf96620e
https://www.genesys-pgr.org/a/1737c39b-eae8-4652-8a2e-330950d1d0cc
https://www.genesys-pgr.org/a/da05f5ec-e6c8-48e7-884b-198a05ee60bb
https://www.genesys-pgr.org/a/d8b4e910-0f8f-4627-892e-92602eb057a1
https://www.genesys-pgr.org/a/558d2393-fa40-4d33-ab14-872e80985ea2
https://www.genesys-pgr.org/a/48bdfc93-90f9-4055-9112-1e2b8334aee6
https://www.genesys-pgr.org/a/405cedeb-86b7-435a-9397-e2c98f1f7ac9
https://www.genesys-pgr.org/a/c08c00cf-45aa-47e4-89ca-bed337c88a8b
https://www.genesys-pgr.org/a/dc723786-ab38-467e-9d61-0b2bf40c836c
https://www.genesys-pgr.org/a/84b1b186-5d47-41e8-b94c-bf84c667f9e0
https://www.genesys-pgr.org/a/54e689ea-83e2-403d-962e-c7ca1beda2d2
https://www.genesys-pgr.org/a/87fdfc88-3905-457d-9631-a52b552597ea
https://www.genesys-pgr.org/a/d1c6258b-11e1-4078-9b0f-f43ca7e83c37
https://www.genesys-pgr.org/a/ae0c063f-0bc4-43fc-b7a3-adb30b5bae53
https://www.genesys-pgr.org/a/bbefcd39-e728-4251-9d81-621697d8f175
https://www.genesys-pgr.org/a/7a344a0e-a480-4f93-bc92-8ca5591bf2d5
https://www.genesys-pgr.org/a/841dae69-5d76-4e8c-ad99-56c97ce2c8f2
https://www.genesys-pgr.org/a/d0a8776a-9a34-46ec-9131-eff9265e3248
https://www.genesys-pgr.org/a/20642fc2-4646-40a9-bb53-63af2acdf06a
https://www.genesys-pgr.org/a/2da8a0b5-baf5-4efe-98e6-a41fddf040ec
https://www.genesys-pgr.org/a/6f0193b2-c8f7-4c95-afb3-b220a3d7b943
https://www.genesys-pgr.org/a/92d170f2-a185-4bc7-a5a1-42cb7429703d

https://www.genesys-pgr.org/a/60ee56eb-1628-442f-af2f-302ac916420b
https://www.genesys-pgr.org/a/974969ab-9225-4e1a-ba4d-6feb95d134bd
https://www.genesys-pgr.org/a/6ca744c3-b5da-4921-9423-a47602f5ae3a
https://www.genesys-pgr.org/a/b5b39351-2b39-421b-a999-5bb7e6f96578
https://www.genesys-pgr.org/a/21ed808d-6c94-4bff-b813-70e3324743e7
https://www.genesys-pgr.org/a/1bab6e33-efe7-45ee-8634-646889573606
https://www.genesys-pgr.org/a/b40c70e9-be26-4b16-9908-2f5ba7860e06
https://www.genesys-pgr.org/a/bb412a0b-4d5f-4aac-b129-9ae3275e743f
https://www.genesys-pgr.org/a/bd259af8-2a09-47be-8fbd-620a67f7e7dc
https://www.genesys-pgr.org/a/5aaa3aea-3c29-4d51-8ffd-12adbfbf620a
https://www.genesys-pgr.org/a/4d9e6840-2253-4166-b221-6d46e06f710f
https://www.genesys-pgr.org/a/7bb3646b-118d-4ed0-a107-a2586fdb0efa
https://www.genesys-pgr.org/a/1dac95b2-77d1-4340-90a2-8f291abec4f1
https://www.genesys-pgr.org/a/26797a04-85bc-404c-bd2e-25a77cfa4ece
https://www.genesys-pgr.org/a/74e2efa2-7bfa-4a94-abc0-d257654214e5
https://www.genesys-pgr.org/a/b74d5342-9819-40f7-a91a-c882b4c8f6a8
https://www.genesys-pgr.org/a/2d236f35-5c50-4b58-9682-1e61d0e85f4f
https://www.genesys-pgr.org/a/4ffb52eb-87ed-4e27-b1fe-30a872ca1a3c
https://www.genesys-pgr.org/a/53d4fb9c-25fb-4365-8591-81f3896cbaf6
https://www.genesys-pgr.org/a/1b19d079-b199-46a8-8d67-5a9f215d7de8
https://www.genesys-pgr.org/a/1b510d31-50f1-44e4-977b-31aacec5dcf1
https://www.ecpgr.cgiar.org/resources/germplasm-databases/list-of-germplasm-databases/crop-databases/crop-database-windows/lupin
https://www.ecpgr.cgiar.org/resources/germplasm-databases/list-of-germplasm-databases/crop-databases/crop-database-windows/lupin
https://www.genesys-pgr.org/a/v2GrrEYOpaV
https://www.genesys-pgr.org/a/87002d0b-d807-4603-8acd-13004f20af2b
https://www.genesys-pgr.org/a/636cfd35-bc98-4f7f-b4bd-a99a45d4c9e9
https://www.genesys-pgr.org/a/71d7bc72-cd1b-4fb6-9d92-235f44da4ced
https://www.genesys-pgr.org/a/07360fc3-c361-4aa7-bc66-f63b07087110
https://www.genesys-pgr.org/a/85210cf4-f119-4af5-9e9c-b9f923a004f2
https://www.genesys-pgr.org/a/c57362ad-e06c-442e-a8c4-ad97bf1e9bc9
https://www.genesys-pgr.org/a/e45d0dd0-cf23-4248-bcea-6af358403508
https://www.genesys-pgr.org/a/24253aa6-6a9d-43d7-bae7-5e63263ed7ea
https://www.genesys-pgr.org/a/8e3a94f5-ebba-4cdd-80d3-af144fadd13a
https://www.genesys-pgr.org/a/9290f678-0d58-4599-9fa6-a47c2b376a0d
https://www.genesys-pgr.org/a/da85b539-f7be-4cac-96d1-7ee8ed2c6694
https://www.genesys-pgr.org/a/718118fd-b726-4559-bda3-c24dab6427b5
https://www.genesys-pgr.org/a/fe7c257e-51d5-4341-b684-01123a39ec3d
https://www.genesys-pgr.org/a/07cc7c46-8e85-4e48-8fbe-26b8d9c362be
https://www.genesys-pgr.org/a/ebcad35a-4f76-4a1d-8165-8b8bef2f405b
https://www.genesys-pgr.org/a/dfc706d6-8225-4b03-9e9a-cac15443c2c8
https://www.genesys-pgr.org/a/41f3c744-ed84-4d3b-9b54-8c3b2664801c
https://www.genesys-pgr.org/a/b1e135b8-df6b-4aa2-ac56-f4cd620b0ea4
https://www.genesys-pgr.org/a/738f458d-f5c7-4c5f-9576-51f7847638cd
https://www.genesys-pgr.org/a/662661f9-1f15-4078-82ee-5d0e861490cc
https://www.genesys-pgr.org/a/v2jLL8rBWJ4
https://www.genesys-pgr.org/a/68e82a9e-8b3f-4fab-bf51-1858e21188a5
https://www.genesys-pgr.org/a/f50ba217-7abf-472c-b710-c2559f9fbaaa
https://www.genesys-pgr.org/a/a0bc9841-8225-490f-941c-58a567aecde7
https://www.genesys-pgr.org/a/e5a69b5b-4993-4753-96d6-5c5f233ba580
https://www.genesys-pgr.org/a/4e9f81c0-c4de-496e-86f3-35e3c05f2249
https://www.genesys-pgr.org/a/948448bf-73c8-483f-9195-43aa4732d1aa
https://www.genesys-pgr.org/a/c4bc8a8f-a16b-44d9-914d-1eeea8bde04c

https://www.genesys-pgr.org/a/2f85683c-c003-495c-bd4c-6899f37e430b
https://www.genesys-pgr.org/a/cbf7e780-6847-40a8-b6c9-7a37e978a995
https://www.genesys-pgr.org/a/a73a7ab1-49b7-4654-b7be-136311336b95
https://www.genesys-pgr.org/a/e2d53283-3375-4891-a1bd-1a7e0a4189ac
https://www.genesys-pgr.org/a/c05b50a4-d5d9-44a2-b218-a9070490bff0
https://www.genesys-pgr.org/a/e83cc590-a4e7-475d-984b-d5ac6839dea5
https://www.genesys-pgr.org/a/6ef01c50-f091-4904-8cc0-6ddaa0c0bf29
https://www.genesys-pgr.org/a/1577b5d3-8777-4bfc-b3ec-0f202dfaae86
https://www.genesys-pgr.org/a/c91e2fb2-296d-4b27-a207-312d51314cc8
https://www.genesys-pgr.org/a/15860d67-deb8-465c-aecf-cc2f930cef85
https://www.genesys-pgr.org/a/41f89aca-22ac-4c13-b083-f02d52621bef
https://www.genesys-pgr.org/a/b214ad22-2a8d-4d7b-93d1-953815bcc6e4
https://www.genesys-pgr.org/a/c6209323-3348-478a-aa4c-8a2af98fc5b2
https://www.genesys-pgr.org/a/cef079e4-c818-48aa-83b9-74937724b232
https://www.genesys-pgr.org/a/610bba52-7795-4509-86ce-7496e9767ae7
https://www.genesys-pgr.org/a/43443d07-70a2-4c73-97bb-f63d63b462e0
https://www.genesys-pgr.org/a/f1957b23-7d54-4960-a115-d6b0a7bfab12
https://www.genesys-pgr.org/a/8a32f661-b526-4e89-af3f-50b96d7cfa4b
https://www.genesys-pgr.org/a/570f9671-9e1d-4dbb-a718-d38b72ad6fad
https://www.genesys-pgr.org/a/2db654fb-0474-479d-8d9c-e0c70c1740fb
https://www.genesys-pgr.org/a/f5701a2a-382f-4cdb-a44b-b5a557a8ea81
https://www.genesys-pgr.org/a/eeeed81b-11d8-4b2a-8c22-039fe0e23d81
https://www.genesys-pgr.org/a/00d1e681-a355-4bca-8443-db5e4df2fc9f
https://www.genesys-pgr.org/a/40bfd18c-76f6-49d9-944c-af9bc76de9cb
https://www.genesys-pgr.org/a/04f0588c-ca8d-4155-af5e-898bcb39b5de
https://www.genesys-pgr.org/a/d05950f7-5eec-4bf1-89ee-b5074490dfc8
https://www.genesys-pgr.org/a/a6109cb4-2b6e-45bd-a328-a9f25e1b693c
https://www.genesys-pgr.org/a/7f6cc662-2c79-47e2-a6eb-7a1cbe06780b
https://www.genesys-pgr.org/a/b4e0e9f8-40fc-4eaa-9bba-bb8a67f40b93
https://www.genesys-pgr.org/a/8ddd5eb0-c4d2-4edc-8eef-24334125ec83
https://www.genesys-pgr.org/a/c5d56803-785d-483d-83f0-8e22d259b0d3
https://www.genesys-pgr.org/a/2be50c31-9d5f-4839-ad45-eba87f446d5e
https://www.genesys-pgr.org/a/c1a1543e-4125-4cb4-bb5e-5d07e107569a
https://www.genesys-pgr.org/a/4c19afa6-dc20-4b0d-be00-b6b123e189b2
https://www.genesys-pgr.org/a/d15cbf6d-2e2b-4d1a-b9b4-e14432be802f
https://www.genesys-pgr.org/a/f7656b66-f175-4ee1-8368-8d2b630b6ec7
https://www.genesys-pgr.org/a/6c4dd9a5-9664-419b-bf13-1fee0b9a753c
https://www.genesys-pgr.org/a/e2978589-66b6-41e1-a998-539d5b4aa818
https://www.genesys-pgr.org/a/37cc0d61-6abc-49e5-a1bb-7baf53792b5f
https://www.genesys-pgr.org/a/3874ea3c-5129-4777-8999-77ac25141694
https://www.genesys-pgr.org/a/7b9ef6b0-a0a6-485e-8cc6-4ed3c3d4a34e
https://www.genesys-pgr.org/a/22593771-52ab-456a-a1ec-be3a2e7a5fe5
https://www.genesys-pgr.org/a/afefa376-8a59-45e6-ba07-66b524b4ad17
https://www.genesys-pgr.org/a/0d02a342-8408-4443-9661-6f26ff4438d3
https://www.genesys-pgr.org/a/a5178ac6-35ef-47e7-9c45-3e3fc7fcac8e
https://www.genesys-pgr.org/a/5a6c24f2-aae6-4fc8-a221-9830dbe97470
https://www.genesys-pgr.org/a/10d304b4-214d-463f-9b4f-3873f748e7d0
https://www.genesys-pgr.org/a/7cc05167-6c6c-4499-adb3-0d8bfcdc6adc
https://www.genesys-pgr.org/a/78b61e6e-f475-463b-bb60-0843e4c55fdf
https://www.genesys-pgr.org/a/36d90a96-8ea0-4b1a-ad0d-e2f4d296f37a
https://www.genesys-pgr.org/a/49a52b7d-99bc-4e88-a135-cdd42dc75f59
https://www.genesys-pgr.org/a/892872f9-d020-43f3-a4ef-d047f99d2ae5

https://www.genesys-pgr.org/a/afc50908-6342-4c90-b727-9b5601720f04
https://www.genesys-pgr.org/a/01ae6efd-ac22-4866-aa77-2b37fe751013
https://www.genesys-pgr.org/a/46ffb979-8faf-47b6-94c1-b51b7cd7117d
https://www.ecpgr.cgiar.org/resources/germplasm-databases/list-of-germplasm-databases/crop-databases/crop-database-windows/lupin
https://www.genesys-pgr.org/a/e6c6d263-6ced-47b6-afe0-bbc1cba484e2
https://www.genesys-pgr.org/a/553514ad-1423-4550-83ba-195524bbaf43
https://www.genesys-pgr.org/a/4885c8ad-e757-4994-b948-438440bdc73f
https://www.genesys-pgr.org/a/f006df93-88f6-4d49-a217-2b70824f6b20
https://www.genesys-pgr.org/a/d117c984-437b-4b2c-b6ff-6a37bb1ea754
https://www.genesys-pgr.org/a/9392e2f5-9a64-4c61-bf74-6cf86e0d5cac
https://www.genesys-pgr.org/a/0107caa5-3ac8-457b-81b3-42fb5671d888
https://www.genesys-pgr.org/a/b76160c4-be11-4e87-9390-3d513c8213c2
https://www.genesys-pgr.org/a/a0c9e1f6-5310-4e52-93f8-490950bcea13
https://www.genesys-pgr.org/a/adf77c9d-533d-4887-8858-235993c1f018
https://www.genesys-pgr.org/a/ea5aa861-207a-46c0-8eda-c8ed82cf941a
https://www.genesys-pgr.org/a/ee6133ca-b107-49b5-b072-c32c389c2caf
https://www.genesys-pgr.org/a/2083bc09-ffe3-41fb-93b5-9fb144141e9c
https://www.genesys-pgr.org/a/2a942f86-2e1f-4747-b249-642d85d41690
https://www.genesys-pgr.org/a/1953ff0d-7fd4-4b62-a6ce-6dea948e89f9
https://www.genesys-pgr.org/a/ea0dc747-0ab5-4dce-9a7c-f1a1f13e94f0
https://www.genesys-pgr.org/a/841e8960-ca7a-45b6-9575-9a75d7cf7bd4
https://www.genesys-pgr.org/a/123f85e6-78fd-41b3-9fa4-8f81ffb6b09d
https://www.genesys-pgr.org/a/566cee9d-c73b-4194-a703-8888abc5073a
https://www.genesys-pgr.org/a/2e6cdb6e-2a49-4772-a0cc-07d49e9bde8a
https://www.genesys-pgr.org/a/295afb01-325b-4cae-a534-1503d387ac3e
https://www.genesys-pgr.org/a/43006f90-d1c2-4942-a87e-5b5b317a30bd
https://www.genesys-pgr.org/a/67e330b9-47b1-4f67-8b38-8a32ede92742
https://www.genesys-pgr.org/a/69177d8c-d357-414b-a6fe-71fe93449b15
https://www.genesys-pgr.org/a/21cdcdf3-17f7-42b1-8b95-715cc78a1dbb
https://www.genesys-pgr.org/a/e9b23a85-aa53-48c5-8bfa-bd1ba0db5c31
https://www.genesys-pgr.org/a/36b71f07-42f5-4576-842d-03600fd7f919
https://www.genesys-pgr.org/a/d4b7903b-9262-4428-b9ac-060d397317ca
https://www.genesys-pgr.org/a/b8b1898e-b605-455f-91ea-ef0405e98789
https://www.genesys-pgr.org/a/82c68f1b-b2ce-4649-9ca6-8d8023d30a43
https://www.genesys-pgr.org/a/18457729-aea3-4966-805e-f49fd8280683
https://www.genesys-pgr.org/a/5ac45f7b-c5c6-42e5-a8e4-32f764722a17
https://www.genesys-pgr.org/a/4ea0a80a-f94e-4b0a-b753-0a8e6b05aae2
https://www.genesys-pgr.org/a/7035ae68-955b-4882-ac1a-1bec911ec526
https://www.genesys-pgr.org/a/2b59e26e-709f-4827-87c6-71001e30b808
https://www.genesys-pgr.org/a/71d6d5b0-b25a-4525-857b-28c3e067f2b3
https://www.genesys-pgr.org/a/9347caf4-de93-4720-bb71-efa0902675eb
https://www.genesys-pgr.org/a/3b6e1afa-b11f-4553-8423-9c4a06b86e7d
https://www.genesys-pgr.org/a/3b9b437c-8d5a-4f16-98a6-4ad6e87d23db
https://www.genesys-pgr.org/a/b305001d-f7fc-4744-acc7-bd3479ba2adf
https://www.genesys-pgr.org/a/0fcfa132-89d0-40a3-8953-81372b4ebcf2
https://www.genesys-pgr.org/a/e4f0c19d-7e3a-49fd-bdfa-d928a41c93ae
https://www.genesys-pgr.org/a/c6246512-3f2a-4fe3-80e4-c11c7a4e0508
https://www.genesys-pgr.org/a/3d1c6fe1-b08f-4d76-b509-78d23962674f
https://www.genesys-pgr.org/a/e2e86426-19fa-4a25-a4f9-b054fc06f6e5
https://www.genesys-pgr.org/a/7158a61a-108c-41d1-885e-988448f5957f
https://www.genesys-pgr.org/a/cf48b051-cf77-4c8a-9331-b98b6f2790ff
https://www.genesys-pgr.org/a/0c8e243e-32af-48f3-9d9f-393bb292ca69

https://www.genesys-pgr.org/a/85baaef9-b32c-4cd7-b169-5cbabc27f6ff
https://www.genesys-pgr.org/a/7449b870-40b4-405c-a5e5-404d51ffb4d9
https://www.genesys-pgr.org/a/f98813f4-38e0-4eab-96a5-887241a62137
https://www.genesys-pgr.org/a/48ad6ef8-4fe6-4819-af52-9c0bf24a83d5
https://www.genesys-pgr.org/a/f716e236-80ce-4341-996e-90985f83d49d
https://www.genesys-pgr.org/a/f0adb6ab-e310-4f32-9da4-03cbca5b8978
https://www.genesys-pgr.org/a/56460200-972b-49a9-882e-d1e77abf85b8
https://www.genesys-pgr.org/a/e09de388-8ea2-4462-970b-982c4eb1b558
https://www.genesys-pgr.org/a/8efdc94e-40c5-4f53-a484-57e74f08f045
https://www.genesys-pgr.org/a/34f2ce22-f3d6-483f-871d-dd490f35fef5
https://www.genesys-pgr.org/a/073548c7-21b7-4f9a-8e01-b10578f48e54
https://www.genesys-pgr.org/a/055101e9-7ad9-40e7-8372-ff4b365c0825
https://www.ecpgr.cgiar.org/resources/germplasm-databases/list-of-germplasm-databases/crop-databases/crop-database-windows/lupin
https://www.ecpgr.cgiar.org/resources/germplasm-databases/list-of-germplasm-databases/crop-databases/crop-database-windows/lupin
https://www.ecpgr.cgiar.org/resources/germplasm-databases/list-of-germplasm-databases/crop-databases/crop-database-windows/lupin
https://www.ecpgr.cgiar.org/resources/germplasm-databases/list-of-germplasm-databases/crop-databases/crop-database-windows/lupin
https://www.ecpgr.cgiar.org/resources/germplasm-databases/list-of-germplasm-databases/crop-databases/crop-database-windows/lupin
https://www.ecpgr.cgiar.org/resources/germplasm-databases/list-of-germplasm-databases/crop-databases/crop-database-windows/lupin
https://www.ecpgr.cgiar.org/resources/germplasm-databases/list-of-germplasm-databases/crop-databases/crop-database-windows/lupin
https://www.ecpgr.cgiar.org/resources/germplasm-databases/list-of-germplasm-databases/crop-databases/crop-database-windows/lupin
https://www.ecpgr.cgiar.org/resources/germplasm-databases/list-of-germplasm-databases/crop-databases/crop-database-windows/lupin
https://www.ecpgr.cgiar.org/resources/germplasm-databases/list-of-germplasm-databases/crop-databases/crop-database-windows/lupin
https://www.ecpgr.cgiar.org/resources/germplasm-databases/list-of-germplasm-databases/crop-databases/crop-database-windows/lupin
https://www.ecpgr.cgiar.org/resources/germplasm-databases/list-of-germplasm-databases/crop-databases/crop-database-windows/lupin
https://www.ecpgr.cgiar.org/resources/germplasm-databases/list-of-germplasm-databases/crop-databases/crop-database-windows/lupin
https://www.ecpgr.cgiar.org/resources/germplasm-databases/list-of-germplasm-databases/crop-databases/crop-database-windows/lupin
https://www.ecpgr.cgiar.org/resources/germplasm-databases/list-of-germplasm-databases/crop-databases/crop-database-windows/lupin
https://www.ecpgr.cgiar.org/resources/germplasm-databases/list-of-germplasm-databases/crop-databases/crop-database-windows/lupin
https://www.ecpgr.cgiar.org/resources/germplasm-databases/list-of-germplasm-databases/crop-databases/crop-database-windows/lupin
https://www.ecpgr.cgiar.org/resources/germplasm-databases/list-of-germplasm-databases/crop-databases/crop-database-windows/lupin
https://www.ecpgr.cgiar.org/resources/germplasm-databases/list-of-germplasm-databases/crop-databases/crop-database-windows/lupin
https://www.ecpgr.cgiar.org/resources/germplasm-databases/list-of-germplasm-databases/crop-databases/crop-database-windows/lupin
https://www.ecpgr.cgiar.org/resources/germplasm-databases/list-of-germplasm-databases/crop-databases/crop-database-windows/lupin
https://www.ecpgr.cgiar.org/resources/germplasm-databases/list-of-germplasm-databases/crop-databases/crop-database-windows/lupin
https://www.ecpgr.cgiar.org/resources/germplasm-databases/list-of-germplasm-databases/crop-databases/crop-database-windows/lupin
https://www.ecpgr.cgiar.org/resources/germplasm-databases/list-of-germplasm-databases/crop-databases/crop-database-windows/lupin
https://www.ecpgr.cgiar.org/resources/germplasm-databases/list-of-germplasm-databases/crop-databases/crop-database-windows/lupin
https://www.ecpgr.cgiar.org/resources/germplasm-databases/list-of-germplasm-databases/crop-databases/crop-database-windows/lupin
https://www.ecpgr.cgiar.org/resources/germplasm-databases/list-of-germplasm-databases/crop-databases/crop-database-windows/lupin
https://www.ecpgr.cgiar.org/resources/germplasm-databases/list-of-germplasm-databases/crop-databases/crop-database-windows/lupin
https://www.ecpgr.cgiar.org/resources/germplasm-databases/list-of-germplasm-databases/crop-databases/crop-database-windows/lupin
https://www.ecpgr.cgiar.org/resources/germplasm-databases/list-of-germplasm-databases/crop-databases/crop-database-windows/lupin
https://www.ecpgr.cgiar.org/resources/germplasm-databases/list-of-germplasm-databases/crop-databases/crop-database-windows/lupin
https://www.ecpgr.cgiar.org/resources/germplasm-databases/list-of-germplasm-databases/crop-databases/crop-database-windows/lupin
https://www.ecpgr.cgiar.org/resources/germplasm-databases/list-of-germplasm-databases/crop-databases/crop-database-windows/lupin
https://www.ecpgr.cgiar.org/resources/germplasm-databases/list-of-germplasm-databases/crop-databases/crop-database-windows/lupin
https://www.ecpgr.cgiar.org/resources/germplasm-databases/list-of-germplasm-databases/crop-databases/crop-database-windows/lupin
https://www.ecpgr.cgiar.org/resources/germplasm-databases/list-of-germplasm-databases/crop-databases/crop-database-windows/lupin
https://www.ecpgr.cgiar.org/resources/germplasm-databases/list-of-germplasm-databases/crop-databases/crop-database-windows/lupin
https://www.ecpgr.cgiar.org/resources/germplasm-databases/list-of-germplasm-databases/crop-databases/crop-database-windows/lupin
https://www.ecpgr.cgiar.org/resources/germplasm-databases/list-of-germplasm-databases/crop-databases/crop-database-windows/lupin
https://www.ecpgr.cgiar.org/resources/germplasm-databases/list-of-germplasm-databases/crop-databases/crop-database-windows/lupin
https://www.ecpgr.cgiar.org/resources/germplasm-databases/list-of-germplasm-databases/crop-databases/crop-database-windows/lupin
https://www.ecpgr.cgiar.org/resources/germplasm-databases/list-of-germplasm-databases/crop-databases/crop-database-windows/lupin

https://www.ecpgr.cgiar.org/resources/germplasm-databases/list-of-germplasm-databases/crop-databases/crop-database-windows/lupin
https://www.ecpgr.cgiar.org/resources/germplasm-databases/list-of-germplasm-databases/crop-databases/crop-database-windows/lupin
https://www.ecpgr.cgiar.org/resources/germplasm-databases/list-of-germplasm-databases/crop-databases/crop-database-windows/lupin
https://www.ecpgr.cgiar.org/resources/germplasm-databases/list-of-germplasm-databases/crop-databases/crop-database-windows/lupin
https://www.ecpgr.cgiar.org/resources/germplasm-databases/list-of-germplasm-databases/crop-databases/crop-database-windows/lupin
https://www.ecpgr.cgiar.org/resources/germplasm-databases/list-of-germplasm-databases/crop-databases/crop-database-windows/lupin
https://www.ecpgr.cgiar.org/resources/germplasm-databases/list-of-germplasm-databases/crop-databases/crop-database-windows/lupin
https://www.ecpgr.cgiar.org/resources/germplasm-databases/list-of-germplasm-databases/crop-databases/crop-database-windows/lupin
https://www.ecpgr.cgiar.org/resources/germplasm-databases/list-of-germplasm-databases/crop-databases/crop-database-windows/lupin
https://www.ecpgr.cgiar.org/resources/germplasm-databases/list-of-germplasm-databases/crop-databases/crop-database-windows/lupin
https://www.ecpgr.cgiar.org/resources/germplasm-databases/list-of-germplasm-databases/crop-databases/crop-database-windows/lupin
https://www.genesys-pgr.org/a/dc8ec904-8b0d-4347-981c-a625072a29e1
https://www.genesys-pgr.org/a/cdfe74d4-3d8d-4701-8b28-e81646f4dbe2
https://www.genesys-pgr.org/a/2872b1fb-79ec-4192-a4c5-8debb8a216fe
https://www.genesys-pgr.org/a/bb61b61f-1062-41fe-b7c5-ffa269e901da
https://www.genesys-pgr.org/a/74ea1d7a-cff0-44c3-a2a9-f23105e187fc
https://www.genesys-pgr.org/a/13a18524-e6ee-4f16-a941-a95b52fdb173
https://www.genesys-pgr.org/a/c4db0a6c-77a4-4360-9bbd-8a0c47b8e7d3
https://www.genesys-pgr.org/a/49d48546-7706-4717-a13d-5036bbd3fb63
https://www.genesys-pgr.org/a/9d1790d3-cee5-4aef-875a-1d1e9aa5b16a
https://www.genesys-pgr.org/a/6c81a801-a376-47ce-bb0d-b3c375e91ffd
https://www.genesys-pgr.org/a/bdbb0241-3254-471b-9ef2-d00aaec34cd8
https://www.genesys-pgr.org/a/f9b39ba7-de2a-47d3-9fb5-0a8619f39a65
https://www.genesys-pgr.org/a/b86df3eb-df21-4909-b6d8-92182c54b85f
https://www.genesys-pgr.org/a/090d522f-7276-4ddc-aff8-47bc940a037f
https://www.genesys-pgr.org/a/92d50140-b617-4ac6-af02-706572cb324a
https://www.genesys-pgr.org/a/5073c793-d8e5-46c6-9b39-1692a86be392
https://www.genesys-pgr.org/a/74d332e4-38d2-4ea0-bc92-e666300f7c72
https://www.genesys-pgr.org/a/fc970d2c-5851-4bf7-b9a7-2a07240ad9ca
https://www.genesys-pgr.org/a/3187fc55-74d0-4ce1-9923-398f9483e0aa
https://www.genesys-pgr.org/a/7579bb99-034b-456f-ba9e-e986708b97bd
https://www.genesys-pgr.org/a/011f4540-83c3-4a31-876c-e9ef44c9dba2
https://www.genesys-pgr.org/a/18118fc8-824f-42d0-87a5-5ec18156f135
https://www.genesys-pgr.org/a/a17907ba-f472-4490-b2c4-cefebc8d7276
https://www.genesys-pgr.org/a/879562bf-4188-4527-8b0e-cb5911fe64ff
https://www.genesys-pgr.org/a/c281f378-32a9-4389-8d1a-63c4839e2ba6
https://www.ecpgr.cgiar.org/resources/germplasm-databases/list-of-germplasm-databases/crop-databases/crop-database-windows/lupin
https://www.ecpgr.cgiar.org/resources/germplasm-databases/list-of-germplasm-databases/crop-databases/crop-database-windows/lupin
https://www.ecpgr.cgiar.org/resources/germplasm-databases/list-of-germplasm-databases/crop-databases/crop-database-windows/lupin
https://www.ecpgr.cgiar.org/resources/germplasm-databases/list-of-germplasm-databases/crop-databases/crop-database-windows/lupin
https://www.ecpgr.cgiar.org/resources/germplasm-databases/list-of-germplasm-databases/crop-databases/crop-database-windows/lupin
https://www.ecpgr.cgiar.org/resources/germplasm-databases/list-of-germplasm-databases/crop-databases/crop-database-windows/lupin
https://www.ecpgr.cgiar.org/resources/germplasm-databases/list-of-germplasm-databases/crop-databases/crop-database-windows/lupin
https://www.ecpgr.cgiar.org/resources/germplasm-databases/list-of-germplasm-databases/crop-databases/crop-database-windows/lupin
https://www.ecpgr.cgiar.org/resources/germplasm-databases/list-of-germplasm-databases/crop-databases/crop-database-windows/lupin
https://www.ecpgr.cgiar.org/resources/germplasm-databases/list-of-germplasm-databases/crop-databases/crop-database-windows/lupin
https://www.ecpgr.cgiar.org/resources/germplasm-databases/list-of-germplasm-databases/crop-databases/crop-database-windows/lupin
https://www.ecpgr.cgiar.org/resources/germplasm-databases/list-of-germplasm-databases/crop-databases/crop-database-windows/lupin
https://www.ecpgr.cgiar.org/resources/germplasm-databases/list-of-germplasm-databases/crop-databases/crop-database-windows/lupin
https://www.ecpgr.cgiar.org/resources/germplasm-databases/list-of-germplasm-databases/crop-databases/crop-database-windows/lupin
https://www.ecpgr.cgiar.org/resources/germplasm-databases/list-of-germplasm-databases/crop-databases/crop-database-windows/lupin
https://www.ecpgr.cgiar.org/resources/germplasm-databases/list-of-germplasm-databases/crop-databases/crop-database-windows/lupin

https://www.ecpgr.cgiar.org/resources/germplasm-databases/list-of-germplasm-databases/crop-databases/crop-database-windows/lupin
https://www.ecpgr.cgiar.org/resources/germplasm-databases/list-of-germplasm-databases/crop-databases/crop-database-windows/lupin
https://www.ecpgr.cgiar.org/resources/germplasm-databases/list-of-germplasm-databases/crop-databases/crop-database-windows/lupin
https://www.ecpgr.cgiar.org/resources/germplasm-databases/list-of-germplasm-databases/crop-databases/crop-database-windows/lupin
https://www.ecpgr.cgiar.org/resources/germplasm-databases/list-of-germplasm-databases/crop-databases/crop-database-windows/lupin
https://www.ecpgr.cgiar.org/resources/germplasm-databases/list-of-germplasm-databases/crop-databases/crop-database-windows/lupin
https://www.ecpgr.cgiar.org/resources/germplasm-databases/list-of-germplasm-databases/crop-databases/crop-database-windows/lupin
https://www.ecpgr.cgiar.org/resources/germplasm-databases/list-of-germplasm-databases/crop-databases/crop-database-windows/lupin
https://www.ecpgr.cgiar.org/resources/germplasm-databases/list-of-germplasm-databases/crop-databases/crop-database-windows/lupin
https://www.ecpgr.cgiar.org/resources/germplasm-databases/list-of-germplasm-databases/crop-databases/crop-database-windows/lupin
https://www.ecpgr.cgiar.org/resources/germplasm-databases/list-of-germplasm-databases/crop-databases/crop-database-windows/lupin
https://www.ecpgr.cgiar.org/resources/germplasm-databases/list-of-germplasm-databases/crop-databases/crop-database-windows/lupin
https://www.ecpgr.cgiar.org/resources/germplasm-databases/list-of-germplasm-databases/crop-databases/crop-database-windows/lupin
https://www.ecpgr.cgiar.org/resources/germplasm-databases/list-of-germplasm-databases/crop-databases/crop-database-windows/lupin
https://www.ecpgr.cgiar.org/resources/germplasm-databases/list-of-germplasm-databases/crop-databases/crop-database-windows/lupin
https://www.ecpgr.cgiar.org/resources/germplasm-databases/list-of-germplasm-databases/crop-databases/crop-database-windows/lupin
https://www.ecpgr.cgiar.org/resources/germplasm-databases/list-of-germplasm-databases/crop-databases/crop-database-windows/lupin
https://www.ecpgr.cgiar.org/resources/germplasm-databases/list-of-germplasm-databases/crop-databases/crop-database-windows/lupin
https://www.ecpgr.cgiar.org/resources/germplasm-databases/list-of-germplasm-databases/crop-databases/crop-database-windows/lupin
https://www.ecpgr.cgiar.org/resources/germplasm-databases/list-of-germplasm-databases/crop-databases/crop-database-windows/lupin
https://www.ecpgr.cgiar.org/resources/germplasm-databases/list-of-germplasm-databases/crop-databases/crop-database-windows/lupin
https://www.ecpgr.cgiar.org/resources/germplasm-databases/list-of-germplasm-databases/crop-databases/crop-database-windows/lupin
https://www.ecpgr.cgiar.org/resources/germplasm-databases/list-of-germplasm-databases/crop-databases/crop-database-windows/lupin
https://www.ecpgr.cgiar.org/resources/germplasm-databases/list-of-germplasm-databases/crop-databases/crop-database-windows/lupin
https://www.ecpgr.cgiar.org/resources/germplasm-databases/list-of-germplasm-databases/crop-databases/crop-database-windows/lupin
https://www.ecpgr.cgiar.org/resources/germplasm-databases/list-of-germplasm-databases/crop-databases/crop-database-windows/lupin
https://www.ecpgr.cgiar.org/resources/germplasm-databases/list-of-germplasm-databases/crop-databases/crop-database-windows/lupin
https://www.ecpgr.cgiar.org/resources/germplasm-databases/list-of-germplasm-databases/crop-databases/crop-database-windows/lupin
https://www.ecpgr.cgiar.org/resources/germplasm-databases/list-of-germplasm-databases/crop-databases/crop-database-windows/lupin
https://www.ecpgr.cgiar.org/resources/germplasm-databases/list-of-germplasm-databases/crop-databases/crop-database-windows/lupin
https://www.ecpgr.cgiar.org/resources/germplasm-databases/list-of-germplasm-databases/crop-databases/crop-database-windows/lupin
https://www.ecpgr.cgiar.org/resources/germplasm-databases/list-of-germplasm-databases/crop-databases/crop-database-windows/lupin
https://www.ecpgr.cgiar.org/resources/germplasm-databases/list-of-germplasm-databases/crop-databases/crop-database-windows/lupin
https://www.ecpgr.cgiar.org/resources/germplasm-databases/list-of-germplasm-databases/crop-databases/crop-database-windows/lupin
https://www.ecpgr.cgiar.org/resources/germplasm-databases/list-of-germplasm-databases/crop-databases/crop-database-windows/lupin
https://www.ecpgr.cgiar.org/resources/germplasm-databases/list-of-germplasm-databases/crop-databases/crop-database-windows/lupin
https://www.ecpgr.cgiar.org/resources/germplasm-databases/list-of-germplasm-databases/crop-databases/crop-database-windows/lupin
https://www.ecpgr.cgiar.org/resources/germplasm-databases/list-of-germplasm-databases/crop-databases/crop-database-windows/lupin
https://www.ecpgr.cgiar.org/resources/germplasm-databases/list-of-germplasm-databases/crop-databases/crop-database-windows/lupin
https://www.ecpgr.cgiar.org/resources/germplasm-databases/list-of-germplasm-databases/crop-databases/crop-database-windows/lupin
https://www.ecpgr.cgiar.org/resources/germplasm-databases/list-of-germplasm-databases/crop-databases/crop-database-windows/lupin
https://www.ecpgr.cgiar.org/resources/germplasm-databases/list-of-germplasm-databases/crop-databases/crop-database-windows/lupin
https://www.ecpgr.cgiar.org/resources/germplasm-databases/list-of-germplasm-databases/crop-databases/crop-database-windows/lupin
https://www.ecpgr.cgiar.org/resources/germplasm-databases/list-of-germplasm-databases/crop-databases/crop-database-windows/lupin
https://www.ecpgr.cgiar.org/resources/germplasm-databases/list-of-germplasm-databases/crop-databases/crop-database-windows/lupin
https://www.ecpgr.cgiar.org/resources/germplasm-databases/list-of-germplasm-databases/crop-databases/crop-database-windows/lupin
https://www.ecpgr.cgiar.org/resources/germplasm-databases/list-of-germplasm-databases/crop-databases/crop-database-windows/lupin
https://www.ecpgr.cgiar.org/resources/germplasm-databases/list-of-germplasm-databases/crop-databases/crop-database-windows/lupin
https://www.ecpgr.cgiar.org/resources/germplasm-databases/list-of-germplasm-databases/crop-databases/crop-database-windows/lupin
https://www.ecpgr.cgiar.org/resources/germplasm-databases/list-of-germplasm-databases/crop-databases/crop-database-windows/lupin
https://www.ecpgr.cgiar.org/resources/germplasm-databases/list-of-germplasm-databases/crop-databases/crop-database-windows/lupin
https://www.ecpgr.cgiar.org/resources/germplasm-databases/list-of-germplasm-databases/crop-databases/crop-database-windows/lupin

https://www.ecpgr.cgiar.org/resources/germplasm-databases/list-of-germplasm-databases/crop-databases/crop-database-windows/lupin
https://www.ecpgr.cgiar.org/resources/germplasm-databases/list-of-germplasm-databases/crop-databases/crop-database-windows/lupin
https://www.ecpgr.cgiar.org/resources/germplasm-databases/list-of-germplasm-databases/crop-databases/crop-database-windows/lupin
https://www.ecpgr.cgiar.org/resources/germplasm-databases/list-of-germplasm-databases/crop-databases/crop-database-windows/lupin
https://www.ecpgr.cgiar.org/resources/germplasm-databases/list-of-germplasm-databases/crop-databases/crop-database-windows/lupin
https://www.ecpgr.cgiar.org/resources/germplasm-databases/list-of-germplasm-databases/crop-databases/crop-database-windows/lupin
https://www.ecpgr.cgiar.org/resources/germplasm-databases/list-of-germplasm-databases/crop-databases/crop-database-windows/lupin
https://www.genesys-pgr.org/a/b76e525e-7883-4d26-989e-8de52f7981f3
https://www.genesys-pgr.org/a/c13a760b-f483-43a9-897a-de85af6edf8b
https://www.genesys-pgr.org/a/faa4b170-b1e8-4a14-891f-09f59a90bd49
https://www.ecpgr.cgiar.org/resources/germplasm-databases/list-of-germplasm-databases/crop-databases/crop-database-windows/lupin
https://www.genesys-pgr.org/a/49a95294-b9c3-4da9-8118-9150a741db8b
https://www.genesys-pgr.org/a/fc4a10b1-f9cb-47b2-b6ac-f672ba8c3d00
https://www.genesys-pgr.org/a/183c6b40-6ee2-475c-af1c-2a8d259098c8
https://www.genesys-pgr.org/a/6e3de5bf-77a2-427e-9506-91d1b3077168
https://www.genesys-pgr.org/a/d1889249-0c2c-4b3d-90be-b096984c0dae
https://www.genesys-pgr.org/a/c6229073-a2cd-4df8-91d7-b12f4724cd08
https://www.genesys-pgr.org/a/9aceb285-75ad-4d43-b7ba-c9f5c635d6c1
https://www.genesys-pgr.org/a/bb76379d-bc4e-40e7-93c7-a28d5c5c6dbb
https://www.genesys-pgr.org/a/63c7202d-589d-45e0-91bc-1601c95853e1
https://www.genesys-pgr.org/a/4ed71a93-6ef9-441b-bce3-ccecf524790e
https://www.genesys-pgr.org/a/36629417-4a06-4842-893c-db900d6620f9
https://www.ecpgr.cgiar.org/resources/germplasm-databases/list-of-germplasm-databases/crop-databases/crop-database-windows/lupin
https://www.ecpgr.cgiar.org/resources/germplasm-databases/list-of-germplasm-databases/crop-databases/crop-database-windows/lupin
https://www.ecpgr.cgiar.org/resources/germplasm-databases/list-of-germplasm-databases/crop-databases/crop-database-windows/lupin
https://www.ecpgr.cgiar.org/resources/germplasm-databases/list-of-germplasm-databases/crop-databases/crop-database-windows/lupin
https://www.ecpgr.cgiar.org/resources/germplasm-databases/list-of-germplasm-databases/crop-databases/crop-database-windows/lupin
https://www.ecpgr.cgiar.org/resources/germplasm-databases/list-of-germplasm-databases/crop-databases/crop-database-windows/lupin
https://www.ecpgr.cgiar.org/resources/germplasm-databases/list-of-germplasm-databases/crop-databases/crop-database-windows/lupin
https://www.ecpgr.cgiar.org/resources/germplasm-databases/list-of-germplasm-databases/crop-databases/crop-database-windows/lupin
https://www.ecpgr.cgiar.org/resources/germplasm-databases/list-of-germplasm-databases/crop-databases/crop-database-windows/lupin
https://www.ecpgr.cgiar.org/resources/germplasm-databases/list-of-germplasm-databases/crop-databases/crop-database-windows/lupin
https://www.ecpgr.cgiar.org/resources/germplasm-databases/list-of-germplasm-databases/crop-databases/crop-database-windows/lupin
https://www.ecpgr.cgiar.org/resources/germplasm-databases/list-of-germplasm-databases/crop-databases/crop-database-windows/lupin
https://www.ecpgr.cgiar.org/resources/germplasm-databases/list-of-germplasm-databases/crop-databases/crop-database-windows/lupin
https://www.ecpgr.cgiar.org/resources/germplasm-databases/list-of-germplasm-databases/crop-databases/crop-database-windows/lupin
https://www.ecpgr.cgiar.org/resources/germplasm-databases/list-of-germplasm-databases/crop-databases/crop-database-windows/lupin
https://www.ecpgr.cgiar.org/resources/germplasm-databases/list-of-germplasm-databases/crop-databases/crop-database-windows/lupin
https://www.ecpgr.cgiar.org/resources/germplasm-databases/list-of-germplasm-databases/crop-databases/crop-database-windows/lupin
https://www.ecpgr.cgiar.org/resources/germplasm-databases/list-of-germplasm-databases/crop-databases/crop-database-windows/lupin
https://www.ecpgr.cgiar.org/resources/germplasm-databases/list-of-germplasm-databases/crop-databases/crop-database-windows/lupin
https://www.ecpgr.cgiar.org/resources/germplasm-databases/list-of-germplasm-databases/crop-databases/crop-database-windows/lupin
https://www.ecpgr.cgiar.org/resources/germplasm-databases/list-of-germplasm-databases/crop-databases/crop-database-windows/lupin
https://www.ecpgr.cgiar.org/resources/germplasm-databases/list-of-germplasm-databases/crop-databases/crop-database-windows/lupin
https://www.ecpgr.cgiar.org/resources/germplasm-databases/list-of-germplasm-databases/crop-databases/crop-database-windows/lupin
https://www.ecpgr.cgiar.org/resources/germplasm-databases/list-of-germplasm-databases/crop-databases/crop-database-windows/lupin
https://www.ecpgr.cgiar.org/resources/germplasm-databases/list-of-germplasm-databases/crop-databases/crop-database-windows/lupin
https://www.ecpgr.cgiar.org/resources/germplasm-databases/list-of-germplasm-databases/crop-databases/crop-database-windows/lupin
https://www.ecpgr.cgiar.org/resources/germplasm-databases/list-of-germplasm-databases/crop-databases/crop-database-windows/lupin
https://www.ecpgr.cgiar.org/resources/germplasm-databases/list-of-germplasm-databases/crop-databases/crop-database-windows/lupin
https://www.ecpgr.cgiar.org/resources/germplasm-databases/list-of-germplasm-databases/crop-databases/crop-database-windows/lupin
https://www.ecpgr.cgiar.org/resources/germplasm-databases/list-of-germplasm-databases/crop-databases/crop-database-windows/lupin

https://www.ecpgr.cgiar.org/resources/germplasm-databases/list-of-germplasm-databases/crop-databases/crop-database-windows/lupin
https://www.ecpgr.cgiar.org/resources/germplasm-databases/list-of-germplasm-databases/crop-databases/crop-database-windows/lupin
https://www.ecpgr.cgiar.org/resources/germplasm-databases/list-of-germplasm-databases/crop-databases/crop-database-windows/lupin
https://www.ecpgr.cgiar.org/resources/germplasm-databases/list-of-germplasm-databases/crop-databases/crop-database-windows/lupin
https://www.ecpgr.cgiar.org/resources/germplasm-databases/list-of-germplasm-databases/crop-databases/crop-database-windows/lupin
https://www.ecpgr.cgiar.org/resources/germplasm-databases/list-of-germplasm-databases/crop-databases/crop-database-windows/lupin
https://www.ecpgr.cgiar.org/resources/germplasm-databases/list-of-germplasm-databases/crop-databases/crop-database-windows/lupin
https://www.ecpgr.cgiar.org/resources/germplasm-databases/list-of-germplasm-databases/crop-databases/crop-database-windows/lupin
https://www.ecpgr.cgiar.org/resources/germplasm-databases/list-of-germplasm-databases/crop-databases/crop-database-windows/lupin
https://www.ecpgr.cgiar.org/resources/germplasm-databases/list-of-germplasm-databases/crop-databases/crop-database-windows/lupin
https://www.ecpgr.cgiar.org/resources/germplasm-databases/list-of-germplasm-databases/crop-databases/crop-database-windows/lupin
https://www.ecpgr.cgiar.org/resources/germplasm-databases/list-of-germplasm-databases/crop-databases/crop-database-windows/lupin
https://www.ecpgr.cgiar.org/resources/germplasm-databases/list-of-germplasm-databases/crop-databases/crop-database-windows/lupin
https://www.ecpgr.cgiar.org/resources/germplasm-databases/list-of-germplasm-databases/crop-databases/crop-database-windows/lupin
https://www.ecpgr.cgiar.org/resources/germplasm-databases/list-of-germplasm-databases/crop-databases/crop-database-windows/lupin
https://www.ecpgr.cgiar.org/resources/germplasm-databases/list-of-germplasm-databases/crop-databases/crop-database-windows/lupin
https://www.ecpgr.cgiar.org/resources/germplasm-databases/list-of-germplasm-databases/crop-databases/crop-database-windows/lupin
https://www.ecpgr.cgiar.org/resources/germplasm-databases/list-of-germplasm-databases/crop-databases/crop-database-windows/lupin
https://www.ecpgr.cgiar.org/resources/germplasm-databases/list-of-germplasm-databases/crop-databases/crop-database-windows/lupin
https://www.ecpgr.cgiar.org/resources/germplasm-databases/list-of-germplasm-databases/crop-databases/crop-database-windows/lupin
https://www.ecpgr.cgiar.org/resources/germplasm-databases/list-of-germplasm-databases/crop-databases/crop-database-windows/lupin
https://www.ecpgr.cgiar.org/resources/germplasm-databases/list-of-germplasm-databases/crop-databases/crop-database-windows/lupin
https://www.ecpgr.cgiar.org/resources/germplasm-databases/list-of-germplasm-databases/crop-databases/crop-database-windows/lupin
https://www.genesys-pgr.org/a/v2dggZ0aQed
https://www.genesys-pgr.org/a/v2QLLE1gOkE
https://www.genesys-pgr.org/a/v2baaLW14wp
https://www.genesys-pgr.org/a/v2xKKGOp5mO
https://www.genesys-pgr.org/a/v2glla4bR4a
https://www.genesys-pgr.org/a/v2OAARBX8pb
https://www.genesys-pgr.org/a/v2QLLE1JaDY
https://www.genesys-pgr.org/a/v21DDmxX5dG
https://www.genesys-pgr.org/a/v20DDmYEpM8
https://www.genesys-pgr.org/a/v2R22pPw5Lp
https://www.genesys-pgr.org/a/v2ZddpVGwrE
https://www.genesys-pgr.org/a/v2aKKVO21ED
https://www.genesys-pgr.org/a/v21DDmxXWdb
https://www.genesys-pgr.org/a/v2OAARBQ4BX
https://www.genesys-pgr.org/a/v2V00pQ6epE
https://www.genesys-pgr.org/a/v23DDmOXEOr
https://www.genesys-pgr.org/a/v2KBBbWP4pR
https://www.genesys-pgr.org/a/v2KBBbWPwWJ
https://www.genesys-pgr.org/a/v2yzzeW7V1b
https://www.genesys-pgr.org/a/v2WOOpmaDql
https://www.genesys-pgr.org/a/v2QLLE1VXXd
https://www.genesys-pgr.org/a/v2m88GQ97JX
https://www.genesys-pgr.org/a/v2xKKGOA0aO
https://www.genesys-pgr.org/a/v2aKKVOrBYA
https://www.genesys-pgr.org/a/v2ZddpV4E8j
https://www.genesys-pgr.org/a/v2V00pQ8mOL
https://www.genesys-pgr.org/a/v28556B1QPB
https://www.genesys-pgr.org/a/v2AVVJlRYG8
https://www.genesys-pgr.org/a/v22KKmaW3D3

https://www.genesys-pgr.org/a/v2R22pPe7Ja
https://www.genesys-pgr.org/a/v23DDmOkk1L
https://www.genesys-pgr.org/a/v2baaLWj00R
https://www.genesys-pgr.org/a/v2BMMDAe3kx
https://www.genesys-pgr.org/a/v2ZddpVBA2j
https://www.genesys-pgr.org/a/v2qBBGVMa7k
https://www.genesys-pgr.org/a/v2933M4a8kx
https://www.genesys-pgr.org/a/v2aKKVOex1a
https://www.genesys-pgr.org/a/v2yzzeW2jV6
https://www.genesys-pgr.org/a/v2QLLE1d4r9
https://www.genesys-pgr.org/a/84d6e49f-1203-4175-a0da-e0ea5aaec94c
https://www.genesys-pgr.org/a/d59f5e49-5186-4787-9ab1-3a9a346015a4
https://www.genesys-pgr.org/a/a62fe5e2-e69e-42b3-ac26-de11ff14acd3
https://www.genesys-pgr.org/a/035168a9-6030-48f3-a253-7c5d70a82f08
https://www.genesys-pgr.org/a/7030a699-39d9-46ac-afa9-ed55c126d477
https://www.genesys-pgr.org/a/777633bf-4072-42cc-bd6d-762208caa008
https://www.genesys-pgr.org/a/262b21fe-3d79-4088-a2d4-39bee4096081
https://www.genesys-pgr.org/a/f7c5c242-a67f-4115-bd93-19d96154e744
https://www.genesys-pgr.org/a/d81c7733-64a6-4f13-879e-881e8fe17121
https://www.ecpgr.cgiar.org/resources/germplasm-databases/list-of-germplasm-databases/crop-databases/crop-database-windows/lupin
https://www.ecpgr.cgiar.org/resources/germplasm-databases/list-of-germplasm-databases/crop-databases/crop-database-windows/lupin
https://www.genesys-pgr.org/a/d2c16747-9a6a-4b1c-bdb6-4422656b3fd4
https://www.genesys-pgr.org/a/c484c0d4-6a03-4038-83cd-2a320c32c736
https://www.genesys-pgr.org/a/76ee0137-e180-4c7b-b346-74007443d23d
https://www.genesys-pgr.org/a/e80b4a1c-0424-401d-a5ae-fc2bb89c44e9
https://www.genesys-pgr.org/a/f13553bf-c6c6-483c-a801-31d1cebebb6e
https://www.genesys-pgr.org/a/74f5b22a-fa6d-44d3-8c8c-47594dbd3468
https://www.genesys-pgr.org/a/d7d9a737-cdda-430d-be6f-b70b50a59d1a
https://www.genesys-pgr.org/a/3d5fa866-4cfd-4208-963d-a9ffd3a53b76
https://www.genesys-pgr.org/a/0e3900ff-03e4-40eb-b0cf-4229f80ef06e
https://www.genesys-pgr.org/a/d39a3bc6-b72d-47a7-b543-9292398e4f67
https://www.genesys-pgr.org/a/6864b3e3-5887-49b9-bd99-43ef2f1bbfe9
https://www.genesys-pgr.org/a/511ff2ee-a5f9-4619-b209-0d6dfd3999c7
https://www.genesys-pgr.org/a/b91978bd-f908-47e2-b5f7-7d240f1ed0b3
https://www.genesys-pgr.org/a/9be7c083-8186-4321-82bc-504e70ba8b03
https://www.genesys-pgr.org/a/2d102bae-0267-4e95-9563-cde268880778
https://www.genesys-pgr.org/a/04a23a80-cc47-4440-bc69-150cd58c4a30
https://www.genesys-pgr.org/a/0fd6de06-15fd-41de-a747-7c57fd22205a
https://www.genesys-pgr.org/a/7afbad1e-2d94-4be0-a26e-2ab86230a2c1
https://www.genesys-pgr.org/a/0fb736b5-d87a-49eb-a4cb-314874b8fce1
https://www.genesys-pgr.org/a/1cae68a5-7ac0-4f24-bbb6-76e307ca1313
https://www.genesys-pgr.org/a/1900a8a5-a644-4849-932e-0eedbd30c85d
https://www.genesys-pgr.org/a/6585e75c-916a-4064-8436-b50f02d32e8e
https://www.genesys-pgr.org/a/5ba0d3ce-a201-49e8-84f4-591c85651dad
https://www.genesys-pgr.org/a/91e91d65-ee4e-4b9c-9d22-f3cb42ebe78b
https://www.genesys-pgr.org/a/9ae0404a-5993-41b6-be76-a37393acda1d
https://www.genesys-pgr.org/a/2ea91be9-5ce9-4577-add7-e1430e2e799f
https://www.genesys-pgr.org/a/3610e40c-8aa3-4b8f-9114-67122b024ea9
https://www.genesys-pgr.org/a/73a28dde-085f-42e3-b3f3-2469261b6d29
https://www.genesys-pgr.org/a/5e83f9cd-9c74-4474-bf6f-a6e1edf9b7c1
https://www.ecpgr.cgiar.org/resources/germplasm-databases/list-of-germplasm-databases/crop-databases/crop-database-windows/lupin
https://www.ecpgr.cgiar.org/resources/germplasm-databases/list-of-germplasm-databases/crop-databases/crop-database-windows/lupin

https://www.ecpgr.cgiar.org/resources/germplasm-databases/list-of-germplasm-databases/crop-databases/crop-database-windows/lupin
https://www.ecpgr.cgiar.org/resources/germplasm-databases/list-of-germplasm-databases/crop-databases/crop-database-windows/lupin
https://www.ecpgr.cgiar.org/resources/germplasm-databases/list-of-germplasm-databases/crop-databases/crop-database-windows/lupin
https://www.ecpgr.cgiar.org/resources/germplasm-databases/list-of-germplasm-databases/crop-databases/crop-database-windows/lupin
https://www.genesys-pgr.org/a/46b29fa6-7313-4c00-af8d-eaede1a3848b
https://www.genesys-pgr.org/a/ce4b687f-a1cc-404f-bedb-9d08a43045f3
https://www.genesys-pgr.org/a/c2ca21ba-cc12-486d-ae61-c79ad1581604
https://www.genesys-pgr.org/a/ffa5fd41-7991-413c-adc8-b6495eb7f006
https://www.genesys-pgr.org/a/6c1dd920-6fe4-434f-9bb3-c60d77591040
https://www.genesys-pgr.org/a/a23d499c-102e-4b86-9296-d66f53c24201
https://www.genesys-pgr.org/a/b5c755d7-73b4-4387-a6fa-2ceb68cb6826
https://www.ecpgr.cgiar.org/resources/germplasm-databases/list-of-germplasm-databases/crop-databases/crop-database-windows/lupin
https://www.genesys-pgr.org/a/e8d218bf-33a4-4c59-94c4-db240c00e0e8
https://www.genesys-pgr.org/a/33e0c647-71ef-4859-a6b0-49a88f01592e
https://www.ecpgr.cgiar.org/resources/germplasm-databases/list-of-germplasm-databases/crop-databases/crop-database-windows/lupin
https://www.ecpgr.cgiar.org/resources/germplasm-databases/list-of-germplasm-databases/crop-databases/crop-database-windows/lupin
https://www.ecpgr.cgiar.org/resources/germplasm-databases/list-of-germplasm-databases/crop-databases/crop-database-windows/lupin
https://www.ecpgr.cgiar.org/resources/germplasm-databases/list-of-germplasm-databases/crop-databases/crop-database-windows/lupin
https://www.ecpgr.cgiar.org/resources/germplasm-databases/list-of-germplasm-databases/crop-databases/crop-database-windows/lupin
https://www.ecpgr.cgiar.org/resources/germplasm-databases/list-of-germplasm-databases/crop-databases/crop-database-windows/lupin
https://www.ecpgr.cgiar.org/resources/germplasm-databases/list-of-germplasm-databases/crop-databases/crop-database-windows/lupin
https://www.ecpgr.cgiar.org/resources/germplasm-databases/list-of-germplasm-databases/crop-databases/crop-database-windows/lupin
https://www.ecpgr.cgiar.org/resources/germplasm-databases/list-of-germplasm-databases/crop-databases/crop-database-windows/lupin
https://www.ecpgr.cgiar.org/resources/germplasm-databases/list-of-germplasm-databases/crop-databases/crop-database-windows/lupin
https://www.ecpgr.cgiar.org/resources/germplasm-databases/list-of-germplasm-databases/crop-databases/crop-database-windows/lupin
https://www.ecpgr.cgiar.org/resources/germplasm-databases/list-of-germplasm-databases/crop-databases/crop-database-windows/lupin
https://www.ecpgr.cgiar.org/resources/germplasm-databases/list-of-germplasm-databases/crop-databases/crop-database-windows/lupin
https://www.ecpgr.cgiar.org/resources/germplasm-databases/list-of-germplasm-databases/crop-databases/crop-database-windows/lupin
https://www.ecpgr.cgiar.org/resources/germplasm-databases/list-of-germplasm-databases/crop-databases/crop-database-windows/lupin
https://www.ecpgr.cgiar.org/resources/germplasm-databases/list-of-germplasm-databases/crop-databases/crop-database-windows/lupin
https://www.ecpgr.cgiar.org/resources/germplasm-databases/list-of-germplasm-databases/crop-databases/crop-database-windows/lupin
https://www.ecpgr.cgiar.org/resources/germplasm-databases/list-of-germplasm-databases/crop-databases/crop-database-windows/lupin
https://www.ecpgr.cgiar.org/resources/germplasm-databases/list-of-germplasm-databases/crop-databases/crop-database-windows/lupin
https://www.ecpgr.cgiar.org/resources/germplasm-databases/list-of-germplasm-databases/crop-databases/crop-database-windows/lupin
https://www.ecpgr.cgiar.org/resources/germplasm-databases/list-of-germplasm-databases/crop-databases/crop-database-windows/lupin
https://www.ecpgr.cgiar.org/resources/germplasm-databases/list-of-germplasm-databases/crop-databases/crop-database-windows/lupin
https://www.ecpgr.cgiar.org/resources/germplasm-databases/list-of-germplasm-databases/crop-databases/crop-database-windows/lupin
https://www.ecpgr.cgiar.org/resources/germplasm-databases/list-of-germplasm-databases/crop-databases/crop-database-windows/lupin
https://www.ecpgr.cgiar.org/resources/germplasm-databases/list-of-germplasm-databases/crop-databases/crop-database-windows/lupin
https://www.ecpgr.cgiar.org/resources/germplasm-databases/list-of-germplasm-databases/crop-databases/crop-database-windows/lupin
https://www.ecpgr.cgiar.org/resources/germplasm-databases/list-of-germplasm-databases/crop-databases/crop-database-windows/lupin
https://www.ecpgr.cgiar.org/resources/germplasm-databases/list-of-germplasm-databases/crop-databases/crop-database-windows/lupin
https://www.ecpgr.cgiar.org/resources/germplasm-databases/list-of-germplasm-databases/crop-databases/crop-database-windows/lupin
https://www.ecpgr.cgiar.org/resources/germplasm-databases/list-of-germplasm-databases/crop-databases/crop-database-windows/lupin
https://www.ecpgr.cgiar.org/resources/germplasm-databases/list-of-germplasm-databases/crop-databases/crop-database-windows/lupin
https://www.ecpgr.cgiar.org/resources/germplasm-databases/list-of-germplasm-databases/crop-databases/crop-database-windows/lupin
https://www.ecpgr.cgiar.org/resources/germplasm-databases/list-of-germplasm-databases/crop-databases/crop-database-windows/lupin
https://www.ecpgr.cgiar.org/resources/germplasm-databases/list-of-germplasm-databases/crop-databases/crop-database-windows/lupin
https://www.ecpgr.cgiar.org/resources/germplasm-databases/list-of-germplasm-databases/crop-databases/crop-database-windows/lupin
https://www.ecpgr.cgiar.org/resources/germplasm-databases/list-of-germplasm-databases/crop-databases/crop-database-windows/lupin
https://www.ecpgr.cgiar.org/resources/germplasm-databases/list-of-germplasm-databases/crop-databases/crop-database-windows/lupin
https://www.ecpgr.cgiar.org/resources/germplasm-databases/list-of-germplasm-databases/crop-databases/crop-database-windows/lupin

https://www.ecpgr.cgiar.org/resources/germplasm-databases/list-of-germplasm-databases/crop-databases/crop-database-windows/lupin
https://www.ecpgr.cgiar.org/resources/germplasm-databases/list-of-germplasm-databases/crop-databases/crop-database-windows/lupin
https://www.ecpgr.cgiar.org/resources/germplasm-databases/list-of-germplasm-databases/crop-databases/crop-database-windows/lupin
https://www.ecpgr.cgiar.org/resources/germplasm-databases/list-of-germplasm-databases/crop-databases/crop-database-windows/lupin
https://www.ecpgr.cgiar.org/resources/germplasm-databases/list-of-germplasm-databases/crop-databases/crop-database-windows/lupin
https://www.ecpgr.cgiar.org/resources/germplasm-databases/list-of-germplasm-databases/crop-databases/crop-database-windows/lupin
https://www.ecpgr.cgiar.org/resources/germplasm-databases/list-of-germplasm-databases/crop-databases/crop-database-windows/lupin
https://www.ecpgr.cgiar.org/resources/germplasm-databases/list-of-germplasm-databases/crop-databases/crop-database-windows/lupin
https://www.ecpgr.cgiar.org/resources/germplasm-databases/list-of-germplasm-databases/crop-databases/crop-database-windows/lupin
https://www.ecpgr.cgiar.org/resources/germplasm-databases/list-of-germplasm-databases/crop-databases/crop-database-windows/lupin
https://www.ecpgr.cgiar.org/resources/germplasm-databases/list-of-germplasm-databases/crop-databases/crop-database-windows/lupin
https://www.ecpgr.cgiar.org/resources/germplasm-databases/list-of-germplasm-databases/crop-databases/crop-database-windows/lupin
https://www.ecpgr.cgiar.org/resources/germplasm-databases/list-of-germplasm-databases/crop-databases/crop-database-windows/lupin
https://www.ecpgr.cgiar.org/resources/germplasm-databases/list-of-germplasm-databases/crop-databases/crop-database-windows/lupin
https://www.ecpgr.cgiar.org/resources/germplasm-databases/list-of-germplasm-databases/crop-databases/crop-database-windows/lupin
https://www.ecpgr.cgiar.org/resources/germplasm-databases/list-of-germplasm-databases/crop-databases/crop-database-windows/lupin
https://www.ecpgr.cgiar.org/resources/germplasm-databases/list-of-germplasm-databases/crop-databases/crop-database-windows/lupin
https://www.ecpgr.cgiar.org/resources/germplasm-databases/list-of-germplasm-databases/crop-databases/crop-database-windows/lupin
https://www.ecpgr.cgiar.org/resources/germplasm-databases/list-of-germplasm-databases/crop-databases/crop-database-windows/lupin
https://www.ecpgr.cgiar.org/resources/germplasm-databases/list-of-germplasm-databases/crop-databases/crop-database-windows/lupin
https://www.ecpgr.cgiar.org/resources/germplasm-databases/list-of-germplasm-databases/crop-databases/crop-database-windows/lupin
https://www.ecpgr.cgiar.org/resources/germplasm-databases/list-of-germplasm-databases/crop-databases/crop-database-windows/lupin
https://www.ecpgr.cgiar.org/resources/germplasm-databases/list-of-germplasm-databases/crop-databases/crop-database-windows/lupin
https://www.ecpgr.cgiar.org/resources/germplasm-databases/list-of-germplasm-databases/crop-databases/crop-database-windows/lupin
https://www.ecpgr.cgiar.org/resources/germplasm-databases/list-of-germplasm-databases/crop-databases/crop-database-windows/lupin
https://www.ecpgr.cgiar.org/resources/germplasm-databases/list-of-germplasm-databases/crop-databases/crop-database-windows/lupin
https://www.ecpgr.cgiar.org/resources/germplasm-databases/list-of-germplasm-databases/crop-databases/crop-database-windows/lupin
https://www.ecpgr.cgiar.org/resources/germplasm-databases/list-of-germplasm-databases/crop-databases/crop-database-windows/lupin
https://www.genesys-pgr.org/a/59ed3a71-adeb-4e37-a0f2-01645adb238b
https://www.genesys-pgr.org/a/69701462-5629-4766-b904-7bcd32947e8f
https://www.genesys-pgr.org/a/f0c1f68a-d137-4b71-b5e2-9403a748d00b
https://www.genesys-pgr.org/a/863f7815-5178-4ee7-8092-aaa8fb9ee78c
https://www.genesys-pgr.org/a/3b7ade5a-08d1-4c19-bb03-bfb28d255bc8
https://www.genesys-pgr.org/a/d2e02cb2-8542-41d6-a558-ccd2dd370602
https://www.genesys-pgr.org/a/455cbede-9192-45f4-a7f5-3f49c44223c9
https://www.genesys-pgr.org/a/50671964-889a-486f-bcf8-37d35fdbcd2c
https://www.genesys-pgr.org/a/9a4f37d5-d56c-4981-bfe5-9295b638b653
https://www.genesys-pgr.org/a/c0a585fd-3964-42e4-9208-2e4ac088e846
https://www.genesys-pgr.org/a/520ac198-349f-4336-b7d7-9d5a7f08d19d
https://www.genesys-pgr.org/a/612e802a-2b5d-4025-830c-07a50af611b5
https://www.genesys-pgr.org/a/4f109b1e-bbe3-4584-9e57-08f47bca7df1
https://www.genesys-pgr.org/a/27e78564-53b1-4d7a-b184-d7558ef0865a
https://www.genesys-pgr.org/a/9e5b2f55-5d71-412d-b2b4-433e950bcc00
https://www.genesys-pgr.org/a/ee13d298-50f8-416b-a928-6890969cf685
https://www.genesys-pgr.org/a/d3a8d8c7-1c77-4614-b41b-8cefa362aa7f
https://www.genesys-pgr.org/a/a100aa00-bcbb-4bba-b0ad-142670abfe5f
https://www.genesys-pgr.org/a/293da733-f881-472d-9f57-d968c0dde6b4
https://www.genesys-pgr.org/a/a24fc1a2-3372-4d09-8348-310693c16f05
https://www.genesys-pgr.org/a/27763674-55ef-42e0-96d7-633b33fbd73e
https://www.genesys-pgr.org/a/b1d0b524-df12-4553-8190-900048e42734
https://www.genesys-pgr.org/a/38bd9e40-a9d4-42f7-8a61-d38505421dfe
https://www.genesys-pgr.org/a/d9bc4b83-577a-450a-9f46-7d8083327a21

https://www.genesys-pgr.org/a/06d8247c-08e9-404a-a550-a9bc061de6b1
https://www.genesys-pgr.org/a/76d53297-ce82-4127-b452-2af8d90b776e
https://www.genesys-pgr.org/a/3ba623ee-4bf7-4c65-93e0-08406577571c
https://www.genesys-pgr.org/a/cd25df23-3fd8-4c40-8cea-0c8f07d9de94
https://www.genesys-pgr.org/a/689d0127-0395-4919-b9d3-355cc30fea1b
https://www.genesys-pgr.org/a/00375fda-2cf2-4275-a4ad-e8b5f70f577e
https://www.genesys-pgr.org/a/f5683210-e7b6-4641-9dab-b4baa32bf825
https://www.genesys-pgr.org/a/d2b9a07c-9f9e-4372-88ac-30deb1dfa7a1
https://www.genesys-pgr.org/a/bd5d9749-4604-473f-b826-60600717c39d
https://www.genesys-pgr.org/a/f22d1392-a7c4-4492-9eb2-27e431c0d2ac
https://www.genesys-pgr.org/a/06d3d399-8cb8-402a-892d-46ae16114e89
https://www.genesys-pgr.org/a/77bd89f6-1b1f-434c-afa4-b4b8ebfbee86
https://www.genesys-pgr.org/a/5536441a-8354-4eea-801b-863b1b06b6ae
https://www.ecpgr.cgiar.org/resources/germplasm-databases/list-of-germplasm-databases/crop-databases/crop-database-windows/lupin
https://www.ecpgr.cgiar.org/resources/germplasm-databases/list-of-germplasm-databases/crop-databases/crop-database-windows/lupin
https://www.ecpgr.cgiar.org/resources/germplasm-databases/list-of-germplasm-databases/crop-databases/crop-database-windows/lupin
https://www.ecpgr.cgiar.org/resources/germplasm-databases/list-of-germplasm-databases/crop-databases/crop-database-windows/lupin
https://www.ecpgr.cgiar.org/resources/germplasm-databases/list-of-germplasm-databases/crop-databases/crop-database-windows/lupin
https://www.ecpgr.cgiar.org/resources/germplasm-databases/list-of-germplasm-databases/crop-databases/crop-database-windows/lupin
https://www.ecpgr.cgiar.org/resources/germplasm-databases/list-of-germplasm-databases/crop-databases/crop-database-windows/lupin
https://www.ecpgr.cgiar.org/resources/germplasm-databases/list-of-germplasm-databases/crop-databases/crop-database-windows/lupin
https://www.ecpgr.cgiar.org/resources/germplasm-databases/list-of-germplasm-databases/crop-databases/crop-database-windows/lupin
https://www.genesys-pgr.org/a/ff09782a-b64a-4fbc-8ff1-29e5b1b54f6b
https://www.genesys-pgr.org/a/9c2c6b6b-11e7-4f5b-80e7-801c99f67d80
https://www.genesys-pgr.org/a/0301f1a1-53e8-4bc6-bdfb-eaa73abff3fe
https://www.genesys-pgr.org/a/bd196d83-f48f-49b0-a049-56b066699216
https://www.genesys-pgr.org/a/f836c961-04a7-4346-84c9-b376b35a39e7
https://www.genesys-pgr.org/a/5d66bd49-fd86-4ab9-b862-aa4b33ae70a3
https://www.genesys-pgr.org/a/a4882315-4ce8-4f4c-8f63-0319b403a848
https://eurisco.ipk-gatersleben.de/apex/f?p=103:16:::::P16_EURISCO_ACC_ID:425268
https://eurisco.ipk-gatersleben.de/apex/f?p=103:16:::::P16_EURISCO_ACC_ID:425267
https://eurisco.ipk-gatersleben.de/apex/f?p=103:16:::::P16_EURISCO_ACC_ID:425261
https://eurisco.ipk-gatersleben.de/apex/f?p=103:16:::::P16_EURISCO_ACC_ID:425260
https://www.ecpgr.cgiar.org/resources/germplasm-databases/list-of-germplasm-databases/crop-databases/crop-database-windows/lupin
https://www.ecpgr.cgiar.org/resources/germplasm-databases/list-of-germplasm-databases/crop-databases/crop-database-windows/lupin
https://www.ecpgr.cgiar.org/resources/germplasm-databases/list-of-germplasm-databases/crop-databases/crop-database-windows/lupin
https://www.ecpgr.cgiar.org/resources/germplasm-databases/list-of-germplasm-databases/crop-databases/crop-database-windows/lupin
https://www.ecpgr.cgiar.org/resources/germplasm-databases/list-of-germplasm-databases/crop-databases/crop-database-windows/lupin
https://www.ecpgr.cgiar.org/resources/germplasm-databases/list-of-germplasm-databases/crop-databases/crop-database-windows/lupin
https://www.ecpgr.cgiar.org/resources/germplasm-databases/list-of-germplasm-databases/crop-databases/crop-database-windows/lupin
https://www.ecpgr.cgiar.org/resources/germplasm-databases/list-of-germplasm-databases/crop-databases/crop-database-windows/lupin
https://www.ecpgr.cgiar.org/resources/germplasm-databases/list-of-germplasm-databases/crop-databases/crop-database-windows/lupin
https://www.ecpgr.cgiar.org/resources/germplasm-databases/list-of-germplasm-databases/crop-databases/crop-database-windows/lupin
https://www.ecpgr.cgiar.org/resources/germplasm-databases/list-of-germplasm-databases/crop-databases/crop-database-windows/lupin
https://www.ecpgr.cgiar.org/resources/germplasm-databases/list-of-germplasm-databases/crop-databases/crop-database-windows/lupin
https://www.ecpgr.cgiar.org/resources/germplasm-databases/list-of-germplasm-databases/crop-databases/crop-database-windows/lupin
https://www.ecpgr.cgiar.org/resources/germplasm-databases/list-of-germplasm-databases/crop-databases/crop-database-windows/lupin
https://www.genesys-pgr.org/a/077ddfae-14b5-4012-9d58-9fa6a464a111
https://www.genesys-pgr.org/a/e6334d47-1c2b-488f-9ceb-8d8a63c3d6f5
https://www.genesys-pgr.org/a/6375f128-9a45-4412-8650-a230f0ba9b57
https://www.genesys-pgr.org/a/47dd1464-7b40-4bc4-8a1f-e79ef6a961d2
https://www.ecpgr.cgiar.org/resources/germplasm-databases/list-of-germplasm-databases/crop-databases/crop-database-windows/lupin

https://www.ecpgr.cgiar.org/resources/germplasm-databases/list-of-germplasm-databases/crop-databases/crop-database-windows/lupin
https://www.genesys-pgr.org/a/9da8fb46-fde1-4560-a447-7a235fa7f8e9
https://www.genesys-pgr.org/a/2e02e6a6-dcd4-40aa-9504-ce4d16567335
https://www.genesys-pgr.org/a/cef68d94-841c-42a9-8c4f-a00ed9c43c00
https://www.genesys-pgr.org/a/0295e498-cbf9-4ea8-b35a-8317dd99bb25
https://www.genesys-pgr.org/a/6e85bbb8-1c17-46e9-a8bb-de5b9a48493b
https://www.genesys-pgr.org/a/0d121092-41ca-461c-9d2a-d6dc3720e706
https://www.genesys-pgr.org/a/d680aa96-c250-4a91-bda4-dfeef3972b37
https://www.genesys-pgr.org/a/77d185ea-a71e-44a7-a355-c58839b7d228
https://www.genesys-pgr.org/a/408932b2-49d4-4862-af32-52942ea1b643
https://www.genesys-pgr.org/a/0363f7f3-7ff8-4519-8e8a-f858a2ca1a92
https://www.genesys-pgr.org/a/620ac92f-91d7-4c98-8b3d-c2f90052a91d
https://www.genesys-pgr.org/a/a29a1671-0e6a-4a63-9560-f9376ac609a9
https://www.genesys-pgr.org/a/6e6b06ef-dcc3-4ed1-953c-d883a40fc1b2
https://www.genesys-pgr.org/a/61cefefe-f07b-46e3-bda3-f8d0acb69e16
https://www.genesys-pgr.org/a/bed758f9-8125-4e3a-bd47-d5b7ee2f0394
https://www.genesys-pgr.org/a/8890989c-a1ce-4b34-97d3-baf82b56eae1
https://www.genesys-pgr.org/a/17956c7e-5448-49bc-bd49-e47573adb7a8
https://www.genesys-pgr.org/a/0854091d-7af0-4c8a-910d-e4b8858edf53
https://www.genesys-pgr.org/a/5a0c2a91-c20a-4484-bb2a-fcf088b266c6
https://www.genesys-pgr.org/a/353ccd09-b1c1-429e-8847-233fb1b8d7e8
https://www.genesys-pgr.org/a/2bbc1c20-be36-4152-b7ec-15dd3e73013e
https://www.genesys-pgr.org/a/bd4d1564-3a86-4e64-8b1e-3c3a424db9a9
https://www.genesys-pgr.org/a/b875715d-7a94-4014-84f6-91605db96893
https://www.genesys-pgr.org/a/fcb47764-dce5-464c-bbe3-53d0640ca1e6
https://www.genesys-pgr.org/a/0d884fbd-7afb-4bbd-aa27-ee9a37cf5730
https://eurisco.ipk-gatersleben.de/apex/f?p=103:16:::::P16_EURISCO_ACC_ID:425262
https://eurisco.ipk-gatersleben.de/apex/f?p=103:16:::::P16_EURISCO_ACC_ID:425048
https://eurisco.ipk-gatersleben.de/apex/f?p=103:16:::::P16_EURISCO_ACC_ID:425045
https://www.genesys-pgr.org/a/93703495-0e56-4cff-99af-8f5bf8dc9e70
https://www.genesys-pgr.org/a/cd395058-c6fd-41c4-8469-1d2547422335
https://www.genesys-pgr.org/a/f0485c34-7837-4f70-8d98-ff583b5d5cfc
https://www.genesys-pgr.org/a/02240984-1d1b-4329-8969-46e7025a49d0
https://www.genesys-pgr.org/a/1616907a-1b22-4339-8dd3-1fe78444794f
https://www.genesys-pgr.org/a/94b17616-cacf-46b4-9f13-c87c22b4c50e
https://www.genesys-pgr.org/a/94b17616-cacf-46b4-9f13-c87c22b4c50e
https://www.genesys-pgr.org/a/12f6201f-0880-4996-a84e-096eb0c0beb7
https://www.genesys-pgr.org/a/dfc1e289-1c6e-4f7f-b8d5-7d6aacedcdcd
https://www.genesys-pgr.org/a/2bc49a64-fba5-4b9f-92cc-60f31140a3b5
https://www.genesys-pgr.org/a/62e96402-2e56-4a59-9f82-246b549a71b8
https://www.genesys-pgr.org/a/a8fc72d4-2172-4e00-bdfa-a0da81ad4844
https://www.genesys-pgr.org/a/34922b97-263e-4d14-8c14-61bf09e93f5b
https://www.genesys-pgr.org/a/4de575a3-050c-4ed8-a0e0-a52975b66e12
https://www.genesys-pgr.org/a/32e4895d-ffcc-40a4-bd56-d9eced69e78e
https://www.genesys-pgr.org/a/a9643b68-248c-4b9b-b1dc-98343f05478d
https://www.genesys-pgr.org/a/cc5dfc53-978e-437c-ab7e-89597ab77a32
https://www.genesys-pgr.org/a/a29dc5b2-51a9-42de-bee5-308b2e3f45f9
https://www.genesys-pgr.org/a/02b1272f-ef31-4901-a734-5d3e833c87e4
https://www.genesys-pgr.org/a/495b340b-b98d-4a57-beef-81311eba08f2
https://www.genesys-pgr.org/a/c51896d4-355a-42bd-aa65-e4ad7191585b
https://www.genesys-pgr.org/a/986f3086-b86f-4a77-9cfe-173af9d3c428
https://www.genesys-pgr.org/a/v2OAAy9D58X

https://www.genesys-pgr.org/a/d3d461c1-d447-4521-bfe9-06315b5fde96
https://www.genesys-pgr.org/a/ad098a2f-ef20-4ae1-a0d3-ac216589e11f
https://www.genesys-pgr.org/a/8fde6e72-1c6b-4571-b7ea-e952681d1ff9
https://www.genesys-pgr.org/a/2ee3fdc3-9853-46a7-ba46-b8a4678880c1
https://www.genesys-pgr.org/a/5f944497-358d-47df-9d69-99815593448d
https://www.genesys-pgr.org/a/423b4418-a27a-488f-a84a-abc4088fa046
https://www.genesys-pgr.org/a/21f5a51a-4c26-45ba-8111-49b0ac551ac0
https://www.genesys-pgr.org/a/852e490a-0318-41d7-abd6-6160c033a1a6
https://www.genesys-pgr.org/a/12fdfd29-029d-4d2d-9835-a8f404df4055
https://www.genesys-pgr.org/a/22312008-c65d-4704-836d-7da212b8510c
https://www.genesys-pgr.org/a/61c7d9ac-832a-4d0e-8b0b-f93885731119
https://www.genesys-pgr.org/a/eaf5e9a4-ab6e-42d1-b6a0-216be076acb6
https://www.genesys-pgr.org/a/ed73ed64-07be-44d6-8e68-178436bb478b
https://www.genesys-pgr.org/a/f03f36ee-acdf-40fe-80b2-20854e2dfe14
https://eurisco.ipk-gatersleben.de/apex/f?p=103:16:::::P16_EURISCO_ACC_ID:425269
https://www.genesys-pgr.org/a/4156f2e9-7b36-4afe-8fba-affbe123489c
https://www.genesys-pgr.org/a/e8a41007-5739-4301-8781-a6eda61e6fe1
https://www.genesys-pgr.org/a/8bbe4958-c53d-4013-a24f-829c7be5c842
https://www.genesys-pgr.org/a/df35c068-87f4-4303-a323-a7f5f8c7ad6c
https://www.genesys-pgr.org/a/fe0df9d4-58a3-42ac-9c96-45fc043d5a6e
https://www.genesys-pgr.org/a/7dbc5877-7903-4959-9ce1-6428347930ca
https://www.genesys-pgr.org/a/79cf8a0b-af21-4385-a20a-0af5fb77e164
https://www.genesys-pgr.org/a/634cd393-292e-4152-ac2b-cf2b9cd3d1ad
https://www.genesys-pgr.org/a/00ffecd8-2b70-4655-b1fa-9d9a84d01ed2
https://www.genesys-pgr.org/a/98807b0e-f5ca-46fa-8ef8-3b5a5923ec93
https://www.genesys-pgr.org/a/550c8a64-7770-4cb8-9051-c3cf85d6be41
https://www.genesys-pgr.org/a/1192db68-278c-4033-aa43-d8f1e8351eaa
https://www.genesys-pgr.org/a/73f68421-d8fd-4571-9e7b-05002e7379d9
https://www.genesys-pgr.org/a/d6ac7ef1-84a9-4673-8c07-74155f9650ce
https://www.genesys-pgr.org/a/01350c6b-04b0-4603-8d9e-56acf90d7142
https://www.genesys-pgr.org/a/24ec99d8-4724-4554-bbfc-df0766cc51f5
https://www.genesys-pgr.org/a/4b49a737-80bb-4760-b47a-ee5380b9eb05
https://www.genesys-pgr.org/a/ebb47a6c-f06d-41ec-ae66-af58262293e0
https://www.genesys-pgr.org/a/4bad5d4c-b74e-4b67-8b6b-80c4da4e94b7
https://www.genesys-pgr.org/a/971ae4e1-7d19-4c5e-85ee-65fa2226b2b4
https://www.genesys-pgr.org/a/6a56fb48-2dcb-417e-b001-e43ff83d43cf
https://www.genesys-pgr.org/a/9514c75a-6fd8-4eee-8e83-725b1b1bff11
https://www.genesys-pgr.org/a/4261d38a-8f4e-4f73-890b-129286cc5560
https://www.genesys-pgr.org/a/c1e206f3-f130-47e7-a5dc-3efbf10cde35
https://www.genesys-pgr.org/a/cab67f3c-3d38-4cff-968d-73469604e8db
https://www.genesys-pgr.org/a/9c787299-8e8a-4cea-990a-664d4405398b
https://npgsweb.ars-grin.gov/gringlobal/accessiondetail?id=1541652
https://www.genesys-pgr.org/a/a66cf186-15e4-4de7-a339-e06383a83584
https://npgsweb.ars-grin.gov/gringlobal/accessiondetail?id=1854175
https://npgsweb.ars-grin.gov/gringlobal/accessiondetail?id=1854176
https://npgsweb.ars-grin.gov/gringlobal/accessiondetail?id=1854181
https://npgsweb.ars-grin.gov/gringlobal/accessiondetail?id=1854183
https://npgsweb.ars-grin.gov/gringlobal/accessiondetail?id=1854184
https://npgsweb.ars-grin.gov/gringlobal/accessiondetail?id=1854185
https://npgsweb.ars-grin.gov/gringlobal/accessiondetail?id=1854189
https://npgsweb.ars-grin.gov/gringlobal/accessiondetail?id=1854193
https://npgsweb.ars-grin.gov/gringlobal/accessiondetail?id=1854196

https://npgsweb.ars-grin.gov/gringlobal/accessiondetail?id=1854197
https://npgsweb.ars-grin.gov/gringlobal/accessiondetail?id=1854199
https://npgsweb.ars-grin.gov/gringlobal/accessiondetail?id=1854300
https://npgsweb.ars-grin.gov/gringlobal/accessiondetail?id=1145124
https://npgsweb.ars-grin.gov/gringlobal/accessiondetail?id=1146622
https://npgsweb.ars-grin.gov/gringlobal/accessiondetail?id=1155338
https://npgsweb.ars-grin.gov/gringlobal/accessiondetail?id=1184518
https://npgsweb.ars-grin.gov/gringlobal/accessiondetail?id=1186343
https://npgsweb.ars-grin.gov/gringlobal/accessiondetail?id=1188194
https://npgsweb.ars-grin.gov/gringlobal/accessiondetail?id=1188956
https://npgsweb.ars-grin.gov/gringlobal/accessiondetail?id=1193152
https://npgsweb.ars-grin.gov/gringlobal/accessiondetail?id=1193635
https://npgsweb.ars-grin.gov/gringlobal/accessiondetail?id=1194331
https://npgsweb.ars-grin.gov/gringlobal/accessiondetail?id=1195777
https://npgsweb.ars-grin.gov/gringlobal/accessiondetail?id=1195839
https://www.genesys-pgr.org/a/80666113-7acd-4927-8db3-75b900be8f09
https://www.genesys-pgr.org/a/ce44f80b-f158-4767-aa6e-15e9358a42cc
https://www.genesys-pgr.org/a/6d30f0fc-0426-42c1-abd8-ca3a909d82b7
https://www.genesys-pgr.org/a/d02b6ebe-d265-49c1-b39d-7ea975647710
https://npgsweb.ars-grin.gov/gringlobal/accessiondetail?id=1276492
https://npgsweb.ars-grin.gov/gringlobal/accessiondetail?id=1276495
https://npgsweb.ars-grin.gov/gringlobal/accessiondetail?id=1276496
https://npgsweb.ars-grin.gov/gringlobal/accessiondetail?id=1285865
https://www.genesys-pgr.org/a/47d4503d-0dc7-4110-a3e6-c1ac339c7002
https://www.genesys-pgr.org/a/a56fdd28-6638-4d02-b28d-2a0a9d2469d1
https://www.genesys-pgr.org/a/ff7cc0a2-8134-48e7-93d9-9498fb3d4602
https://npgsweb.ars-grin.gov/gringlobal/accessiondetail?id=1352857
https://npgsweb.ars-grin.gov/gringlobal/accessiondetail?id=1352859
https://npgsweb.ars-grin.gov/gringlobal/accessiondetail?id=1352860
https://npgsweb.ars-grin.gov/gringlobal/accessiondetail?id=1352862
https://npgsweb.ars-grin.gov/gringlobal/accessiondetail?id=1352863
https://npgsweb.ars-grin.gov/gringlobal/accessiondetail?id=1352864
https://npgsweb.ars-grin.gov/gringlobal/accessiondetail?id=1352865
https://npgsweb.ars-grin.gov/gringlobal/accessiondetail?id=1352866
https://npgsweb.ars-grin.gov/gringlobal/accessiondetail?id=1352867
https://npgsweb.ars-grin.gov/gringlobal/accessiondetail?id=1352868
https://npgsweb.ars-grin.gov/gringlobal/accessiondetail?id=1352869
https://npgsweb.ars-grin.gov/gringlobal/accessiondetail?id=1352870
https://npgsweb.ars-grin.gov/gringlobal/accessiondetail?id=1352871
https://npgsweb.ars-grin.gov/gringlobal/accessiondetail?id=1352872
https://npgsweb.ars-grin.gov/gringlobal/accessiondetail?id=1352873
https://npgsweb.ars-grin.gov/gringlobal/accessiondetail?id=1352874
https://npgsweb.ars-grin.gov/gringlobal/accessiondetail?id=1352875
https://npgsweb.ars-grin.gov/gringlobal/accessiondetail?id=1352876
https://www.genesys-pgr.org/a/93c8d6ec-211c-4b95-a64c-3938ce8e901c
https://www.genesys-pgr.org/a/a2840870-142e-438b-8515-a4a9cfa65410
https://www.genesys-pgr.org/a/8780898e-e5dc-435f-a326-72a94be6c873
https://www.genesys-pgr.org/a/e2aea3ff-6057-4235-bc38-bc650fe0e6a4
https://www.genesys-pgr.org/a/1d18cf5f-0ea0-4542-b12d-85dd788d25f8
https://www.genesys-pgr.org/a/02e9da3a-4dcc-45d8-9601-f7ebf406ac2d
https://www.genesys-pgr.org/a/c1d6f432-0ecb-45b7-a21f-21293f8c22ea
https://www.genesys-pgr.org/a/aa3be11a-95cd-4214-b4b4-c5b276b7fedc

https://www.genesys-pgr.org/a/b2585fe5-1107-4b55-a2dc-59b2008d0e11
https://www.genesys-pgr.org/a/615fc987-4172-41b9-81a0-392aa4583b68
https://www.genesys-pgr.org/a/893ddb23-2f36-4fe6-af6c-02985615edf0
https://www.genesys-pgr.org/a/9b7aa382-7827-49cc-ad33-e083e1270986
https://www.genesys-pgr.org/a/daa572a6-31e0-4fb6-8449-fa7609b2578b
https://www.genesys-pgr.org/a/4deda787-eb83-4a5d-bc94-22ce83a5039f
https://www.genesys-pgr.org/a/8f885d17-88ad-443e-89a8-2e6169a2ad97
https://www.genesys-pgr.org/a/6a593b34-69a5-47b2-98e1-0ddd7debdc7e
https://npgsweb.ars-grin.gov/gringlobal/accessiondetail?id=1352892
https://npgsweb.ars-grin.gov/gringlobal/accessiondetail?id=1352895
https://www.genesys-pgr.org/a/2780ed71-ec08-4521-9a4f-0e9d5a5f797f
https://www.genesys-pgr.org/a/1e3c4999-9c6c-4ef5-a327-0de00a41057e
https://www.genesys-pgr.org/a/a6937fd7-ea18-4a3e-949d-88ed21fc5853
https://www.genesys-pgr.org/a/5b5e6366-d720-48e5-9a8f-07517a36f88a
https://www.genesys-pgr.org/a/cb32f70f-c20b-4927-ab50-ff944a89786f
https://www.genesys-pgr.org/a/70a71e8b-5ef8-4f2b-82d5-1b4f5c80e1db
https://www.genesys-pgr.org/a/ffb86070-13c0-4f94-8f4e-e46d6f385ea7
https://www.genesys-pgr.org/a/02066ef7-d055-47c6-b9a7-27dbd12956c6
https://www.genesys-pgr.org/a/93328062-a343-427f-a470-28465fd55f3c
https://www.genesys-pgr.org/10.18730/XV4J8
https://www.genesys-pgr.org/a/67c00cce-4fa9-4aca-9b79-1f088d518ad8
https://www.genesys-pgr.org/a/de1d63c8-55b4-4261-820a-12fcddd5d875
https://www.genesys-pgr.org/a/9d511bab-412c-464d-a1f5-42c0527e330b
https://www.genesys-pgr.org/a/41885407-3698-4069-9c97-8683f8331fbb
https://www.genesys-pgr.org/a/c49f17ec-e45d-4994-8e4e-7a9a72945bdd
https://www.genesys-pgr.org/a/2ba7860c-a096-4908-a332-e5e089ba9b46
https://npgsweb.ars-grin.gov/gringlobal/accessiondetail?id=1376481
https://npgsweb.ars-grin.gov/gringlobal/accessiondetail?id=1376482
https://npgsweb.ars-grin.gov/gringlobal/accessiondetail?id=1376483
https://npgsweb.ars-grin.gov/gringlobal/accessiondetail?id=1376484
https://npgsweb.ars-grin.gov/gringlobal/accessiondetail?id=1376485
https://npgsweb.ars-grin.gov/gringlobal/accessiondetail?id=1376486
https://npgsweb.ars-grin.gov/gringlobal/accessiondetail?id=1376487
https://npgsweb.ars-grin.gov/gringlobal/accessiondetail?id=1376488
https://npgsweb.ars-grin.gov/gringlobal/accessiondetail?id=1376489
https://npgsweb.ars-grin.gov/gringlobal/accessiondetail?id=1376490
https://npgsweb.ars-grin.gov/gringlobal/accessiondetail?id=1376491
https://npgsweb.ars-grin.gov/gringlobal/accessiondetail?id=1376492
https://npgsweb.ars-grin.gov/gringlobal/accessiondetail?id=1376493
https://npgsweb.ars-grin.gov/gringlobal/accessiondetail?id=1376494
https://npgsweb.ars-grin.gov/gringlobal/accessiondetail?id=1376495
https://npgsweb.ars-grin.gov/gringlobal/accessiondetail?id=1376496
https://npgsweb.ars-grin.gov/gringlobal/accessiondetail?id=1378008
https://npgsweb.ars-grin.gov/gringlobal/accessiondetail?id=1378009
https://npgsweb.ars-grin.gov/gringlobal/accessiondetail?id=1378010
https://npgsweb.ars-grin.gov/gringlobal/accessiondetail?id=1378011
https://npgsweb.ars-grin.gov/gringlobal/accessiondetail?id=1382368
https://www.genesys-pgr.org/a/6d920c94-dc1a-41f2-b83f-98be6aca2725
https://www.genesys-pgr.org/a/d45f6078-b44d-47f4-b98f-6de282709738
https://www.genesys-pgr.org/a/8e19c19f-5b62-441e-b8a6-7d1207de75ff
https://npgsweb.ars-grin.gov/gringlobal/accessiondetail?id=1382373
https://npgsweb.ars-grin.gov/gringlobal/accessiondetail?id=1397584

https://www.genesys-pgr.org/a/e3f67dde-a43f-4da2-968d-473c62f24a5f
https://www.genesys-pgr.org/a/835a2ca7-6645-439d-95b1-2ce42d4a7e18
https://www.genesys-pgr.org/a/7b24e7c3-e0ac-47f3-8479-f309d217e1e6
https://npgsweb.ars-grin.gov/gringlobal/accessiondetail?id=1397588
https://npgsweb.ars-grin.gov/gringlobal/accessiondetail?id=1400780
https://npgsweb.ars-grin.gov/gringlobal/accessiondetail?id=1400781
https://npgsweb.ars-grin.gov/gringlobal/accessiondetail?id=1400782
https://npgsweb.ars-grin.gov/gringlobal/accessiondetail?id=1400783
https://npgsweb.ars-grin.gov/gringlobal/accessiondetail?id=1400784
https://www.genesys-pgr.org/a/645f81c4-ed0f-45a3-ae0f-fae3bcb8c858
https://npgsweb.ars-grin.gov/gringlobal/accessiondetail?id=1411558
https://npgsweb.ars-grin.gov/gringlobal/accessiondetail?id=1411559
https://npgsweb.ars-grin.gov/gringlobal/accessiondetail?id=1411560
https://npgsweb.ars-grin.gov/gringlobal/accessiondetail?id=1411561
https://npgsweb.ars-grin.gov/gringlobal/accessiondetail?id=1411562
https://npgsweb.ars-grin.gov/gringlobal/accessiondetail?id=1411563
https://npgsweb.ars-grin.gov/gringlobal/accessiondetail?id=1411565
https://npgsweb.ars-grin.gov/gringlobal/accessiondetail?id=1411566
https://npgsweb.ars-grin.gov/gringlobal/accessiondetail?id=1411575
https://npgsweb.ars-grin.gov/gringlobal/accessiondetail?id=1428630
https://npgsweb.ars-grin.gov/gringlobal/accessiondetail?id=1428631
https://npgsweb.ars-grin.gov/gringlobal/accessiondetail?id=1428632
https://npgsweb.ars-grin.gov/gringlobal/accessiondetail?id=1428633
https://npgsweb.ars-grin.gov/gringlobal/accessiondetail?id=1428634
https://npgsweb.ars-grin.gov/gringlobal/accessiondetail?id=1428635
https://npgsweb.ars-grin.gov/gringlobal/accessiondetail?id=1428636
https://npgsweb.ars-grin.gov/gringlobal/accessiondetail?id=1428637
https://npgsweb.ars-grin.gov/gringlobal/accessiondetail?id=1428638
https://npgsweb.ars-grin.gov/gringlobal/accessiondetail?id=1428639
https://npgsweb.ars-grin.gov/gringlobal/accessiondetail?id=1428640
https://npgsweb.ars-grin.gov/gringlobal/accessiondetail?id=1428641
https://npgsweb.ars-grin.gov/gringlobal/accessiondetail?id=1428642
https://npgsweb.ars-grin.gov/gringlobal/accessiondetail?id=1428643
https://npgsweb.ars-grin.gov/gringlobal/accessiondetail?id=1428650
https://npgsweb.ars-grin.gov/gringlobal/accessiondetail?id=1437949
https://npgsweb.ars-grin.gov/gringlobal/accessiondetail?id=1437950
https://npgsweb.ars-grin.gov/gringlobal/accessiondetail?id=1437951
https://npgsweb.ars-grin.gov/gringlobal/accessiondetail?id=1437952
https://npgsweb.ars-grin.gov/gringlobal/accessiondetail?id=1437954
https://npgsweb.ars-grin.gov/gringlobal/accessiondetail?id=1437955
https://npgsweb.ars-grin.gov/gringlobal/accessiondetail?id=1437956
https://npgsweb.ars-grin.gov/gringlobal/accessiondetail?id=1437958
https://npgsweb.ars-grin.gov/gringlobal/accessiondetail?id=1437959
https://npgsweb.ars-grin.gov/gringlobal/accessiondetail?id=1437960
https://npgsweb.ars-grin.gov/gringlobal/accessiondetail?id=1437962
https://npgsweb.ars-grin.gov/gringlobal/accessiondetail?id=1437963
https://npgsweb.ars-grin.gov/gringlobal/accessiondetail?id=1437964
https://www.genesys-pgr.org/a/960c19e1-1a77-4955-b56a-13cb8c40f118
https://www.genesys-pgr.org/a/6bef6588-7cc7-45cd-97c6-fd2678816cac
https://npgsweb.ars-grin.gov/gringlobal/accessiondetail?id=1111106
https://npgsweb.ars-grin.gov/gringlobal/accessiondetail?id=1048965
https://npgsweb.ars-grin.gov/gringlobal/accessiondetail?id=1048968

https://npgsweb.ars-grin.gov/gringlobal/accessiondetail?id=1052709
https://npgsweb.ars-grin.gov/gringlobal/accessiondetail?id=1052733
https://npgsweb.ars-grin.gov/gringlobal/accessiondetail?id=1111100
https://npgsweb.ars-grin.gov/gringlobal/accessiondetail?id=1111101
https://npgsweb.ars-grin.gov/gringlobal/accessiondetail?id=1111102
https://npgsweb.ars-grin.gov/gringlobal/accessiondetail?id=1111103
https://npgsweb.ars-grin.gov/gringlobal/accessiondetail?id=1111104
https://npgsweb.ars-grin.gov/gringlobal/accessiondetail?id=1111105
https://www.genesys-pgr.org/a/eabf9e39-bf3f-4f13-8dff-3b1eafc7c4c0
https://www.genesys-pgr.org/a/371b0664-99df-456d-b438-131a7513749a
https://www.genesys-pgr.org/a/0e3c12a3-fdb2-4db8-b434-b5f22ecedbec
https://www.genesys-pgr.org/a/40847b7b-63cb-49fb-9a63-fb8cbd4ed8eb
https://www.genesys-pgr.org/a/3f14c771-5151-4245-a70b-d0ce2f609c5f
https://www.genesys-pgr.org/a/2a7905bd-ee4a-42c7-81c6-c3867f394499
https://www.genesys-pgr.org/a/ed07e438-d805-4995-9b14-64f0ae391460
https://www.genesys-pgr.org/a/57839075-19d5-445f-a25c-ce9fee70ea58
https://www.genesys-pgr.org/a/64c925e1-4ff9-4fef-8562-fc1eff1df7b4
https://npgsweb.ars-grin.gov/gringlobal/accessiondetail?id=1668354
https://npgsweb.ars-grin.gov/gringlobal/accessiondetail?id=1681731
https://www.genesys-pgr.org/a/daa768a6-127d-460a-83cc-55d8f8970260
https://www.genesys-pgr.org/a/80be8b3e-391f-4843-82a5-cfcbb9d7cd5f
https://www.genesys-pgr.org/a/57e3b5b4-7294-41af-99a0-19b2f2a9f6ed
https://npgsweb.ars-grin.gov/gringlobal/accessiondetail?id=1072596
https://www.genesys-pgr.org/a/fbf1f5f1-a0ca-4a03-8bfd-357d1338bc48
https://npgsweb.ars-grin.gov/gringlobal/accessiondetail?id=1854172
https://npgsweb.ars-grin.gov/gringlobal/accessiondetail?id=1854173
https://npgsweb.ars-grin.gov/gringlobal/accessiondetail?id=1854174
https://npgsweb.ars-grin.gov/gringlobal/accessiondetail?id=1854178
https://npgsweb.ars-grin.gov/gringlobal/accessiondetail?id=1854179
https://npgsweb.ars-grin.gov/gringlobal/accessiondetail?id=1854180
https://npgsweb.ars-grin.gov/gringlobal/accessiondetail?id=1854186
https://npgsweb.ars-grin.gov/gringlobal/accessiondetail?id=1854188
https://npgsweb.ars-grin.gov/gringlobal/accessiondetail?id=1854190
https://npgsweb.ars-grin.gov/gringlobal/accessiondetail?id=1854191
https://npgsweb.ars-grin.gov/gringlobal/accessiondetail?id=1854192
https://npgsweb.ars-grin.gov/gringlobal/accessiondetail?id=1854194
https://npgsweb.ars-grin.gov/gringlobal/accessiondetail?id=1854195
https://www.genesys-pgr.org/a/f5e5f424-5fc8-4644-9ff6-b40e3776507f

https://www.genesys-pgr.org/a/d8ef94a9-4fe9-4ebf-bbbf-b693d1361657

https://www.ecpgr.cgiar.org/resources/germplasm-databases/list-of-germplasm-databases/crop-databases/crop-database-windows/lupin
Email from Massimo to Karolina_June_10_2021

Email from Massimo to Karolina_June_10_2022
Provided by Recanati 0

63
