## Supplementary material for "Genetic and phenotypic characterization of global *Lupinus albus* genetic resources for the development of a CORE collection": L.albus-manuscript_SupplementaryFigures

A

| Country | Number of Wild_Accessions |
|---|---|
| Algeria | 4 |
| Egypt | 2 |
| Ethiopia | 2 |
| France | 1 |
| Germany | 1 |
| Greece | 57 |
| Italy | 30 |
| Palestine | 5 |
| Poland | 1 |
| Portugal | 21 |
| Serbia and Montenegro | 1 |
| South Africa | 2 |
| Spain | 89 |
| Syria | 5 |
| Turkey | 5 |
| Ukraine | 1 |
| United Kingdom | 1 |
| United States of America | 2 |
| USSR | 2 |
| Venezuela | 1 |

C

| Country | Number of accessions (Non- wild) | Country | Number of accessions (Non-wild |
|---|---|---|---|
| Croatia | 2 | Greece | 95 |
| Czech Republic | 9 | Hungary | 28 |
| Denmark | 1 | India | 1 |
| Ethiopia | 25 | Morocco | 18 |
| Georgia | 2 | Netherlands | 4 |
| Germany | 109 | New Zealand | 2 |
| Italy | 98 | Poland | 66 |
| Jordan | 5 | Russia | 5 |
| Kenya | 4 | Serbia | 4 |
| Lebanon | 2 | Serbia and Montenegro | 8 |
| Lithuania | 1 | Slovakia | 1 |
| Palestine | 5 | South Africa | 7 |
| Portugal | 418 | Spain | 526 |
| Romania | 18 | Sudan | 17 |
| Algeria | 10 | Syria | 9 |
| Australia | 9 | Turkey | 9 |
| Belarus | 1 | Ukraine | 21 |
| Brazil | 3 | United Kingdom | 6 |
| Bulgaria | 5 | United States of America | 25 |
| Chile | 15 | Unknown | 187 |
| Egypt | 157 | USSR | 32 |
| France | 85 | | |

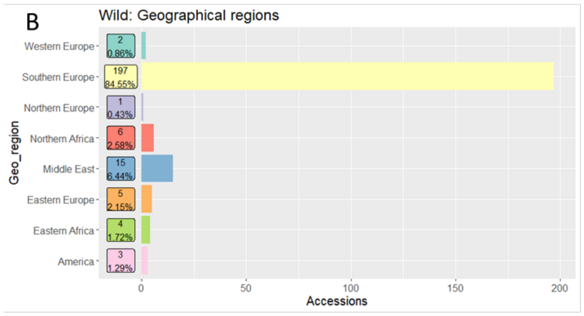


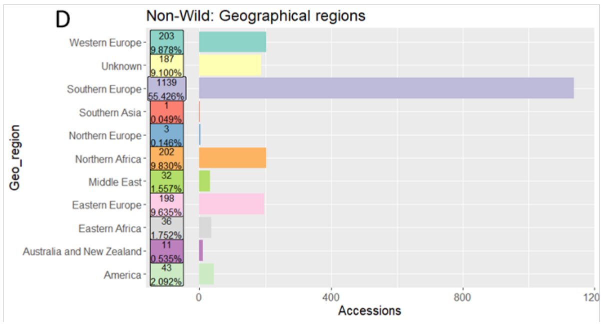


Figure S1: Country wise and geographical regions distribution of Wild accessions (A,B) and rest of all accessions (C,D) in *L. albus* R-CORE collection.

Cluster:     I    II    III
Accessions:182 1904   26

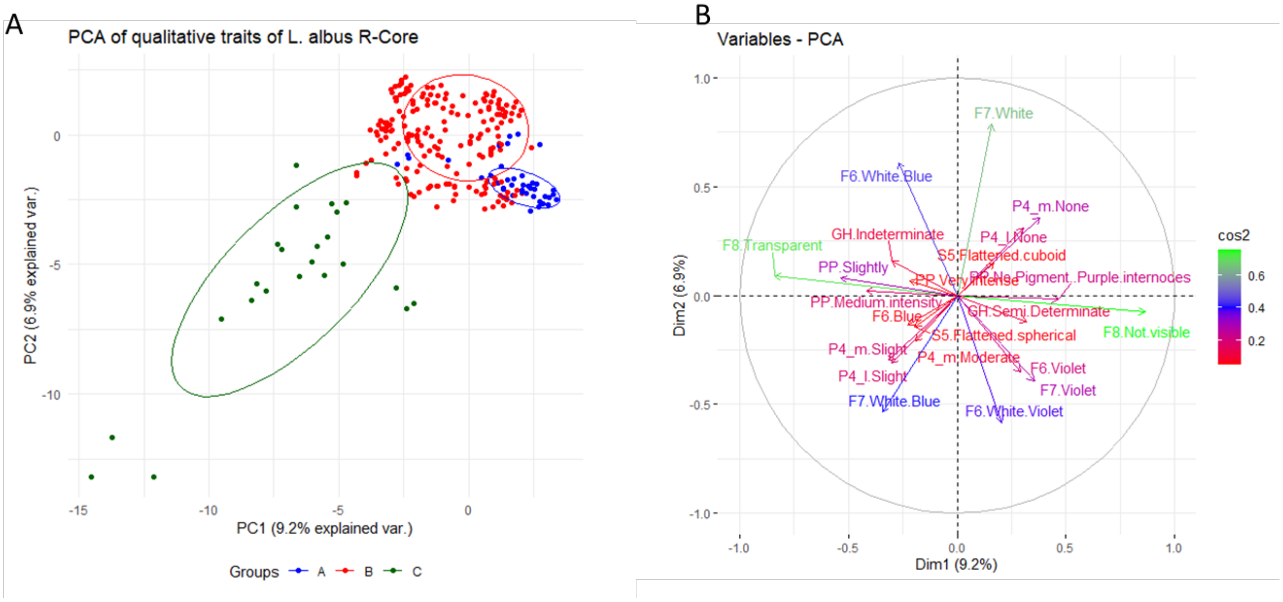


Figure S2: PCA analysis using all qualitative traits excluding seed ornamentation traits, and excluding the breeding/researchmaterial  (A) Biplot of R-CORE accessions of *L. albus* and (B) the qualitative traits resulted from the MCA analysis.

Cluster: I II III
Accesssions:1498 566 48

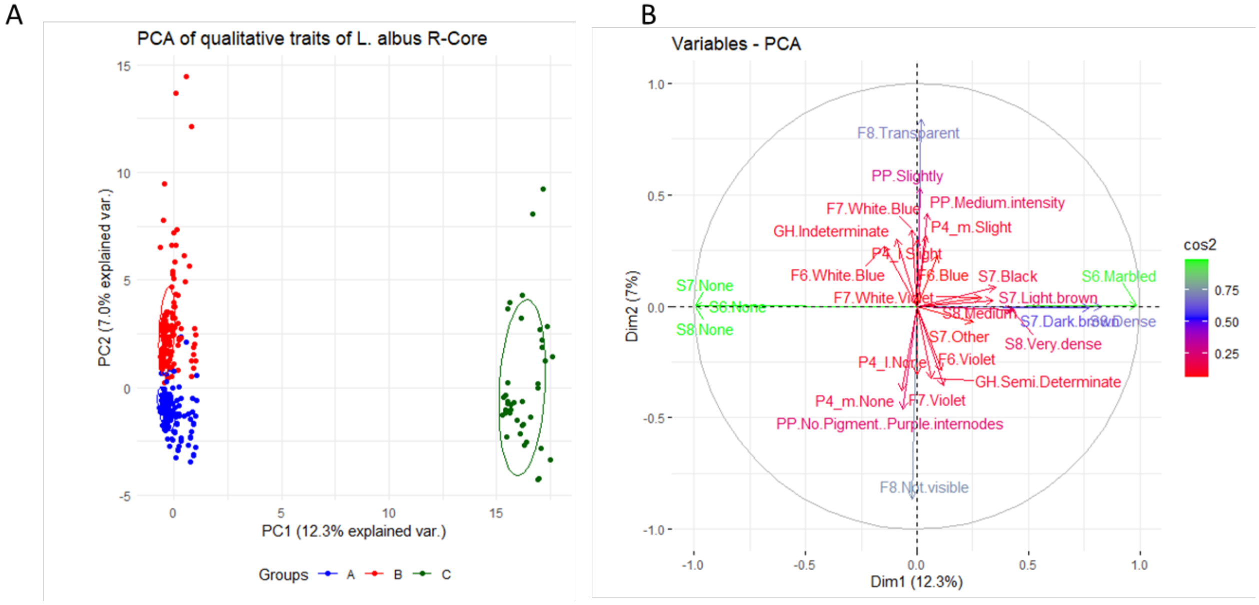


Figure S3: PCA analysis using all qualitative traits excluding the breeding/researchmaterial (A) Biplot of R-CORE accessions of *L. albus* and (B) the qualitative traits resulted from the MCA analysis.

```
Cluster:      I    II    III
Accesssions:190 2071  27
```

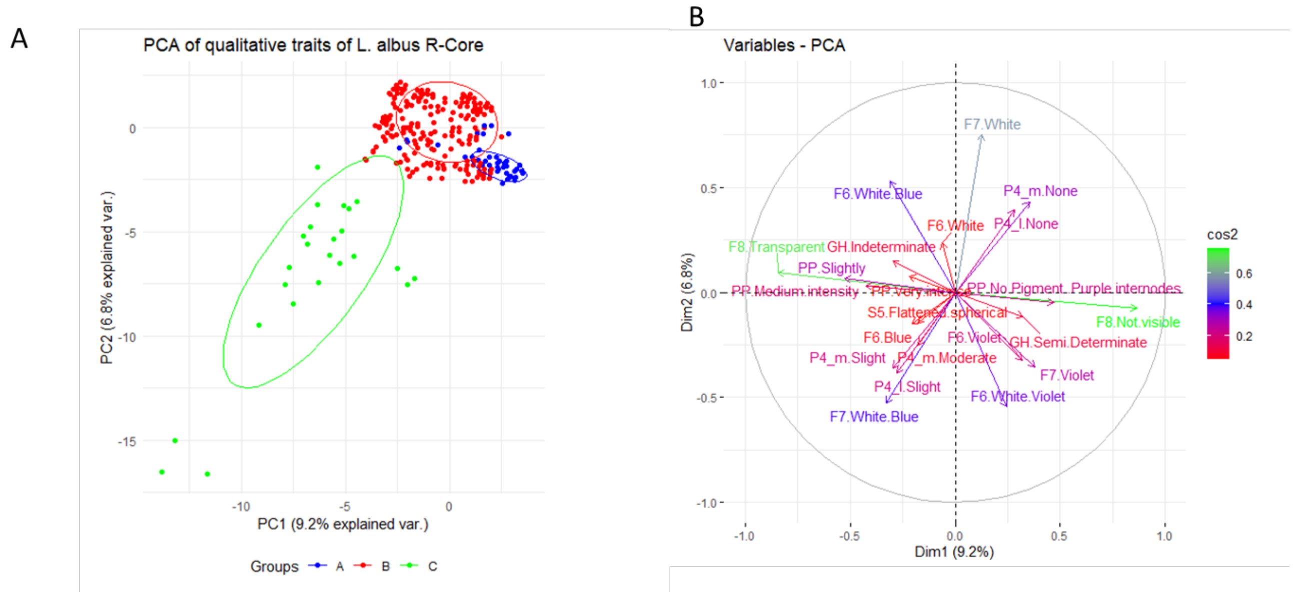


Figure S4: PCA analysis using qualitative traits excluding the seed ornamentation trait (A) Biplot of R-CORE accessions of *L. albus* and (B) the qualitative traits resulted from the MCA analysis.

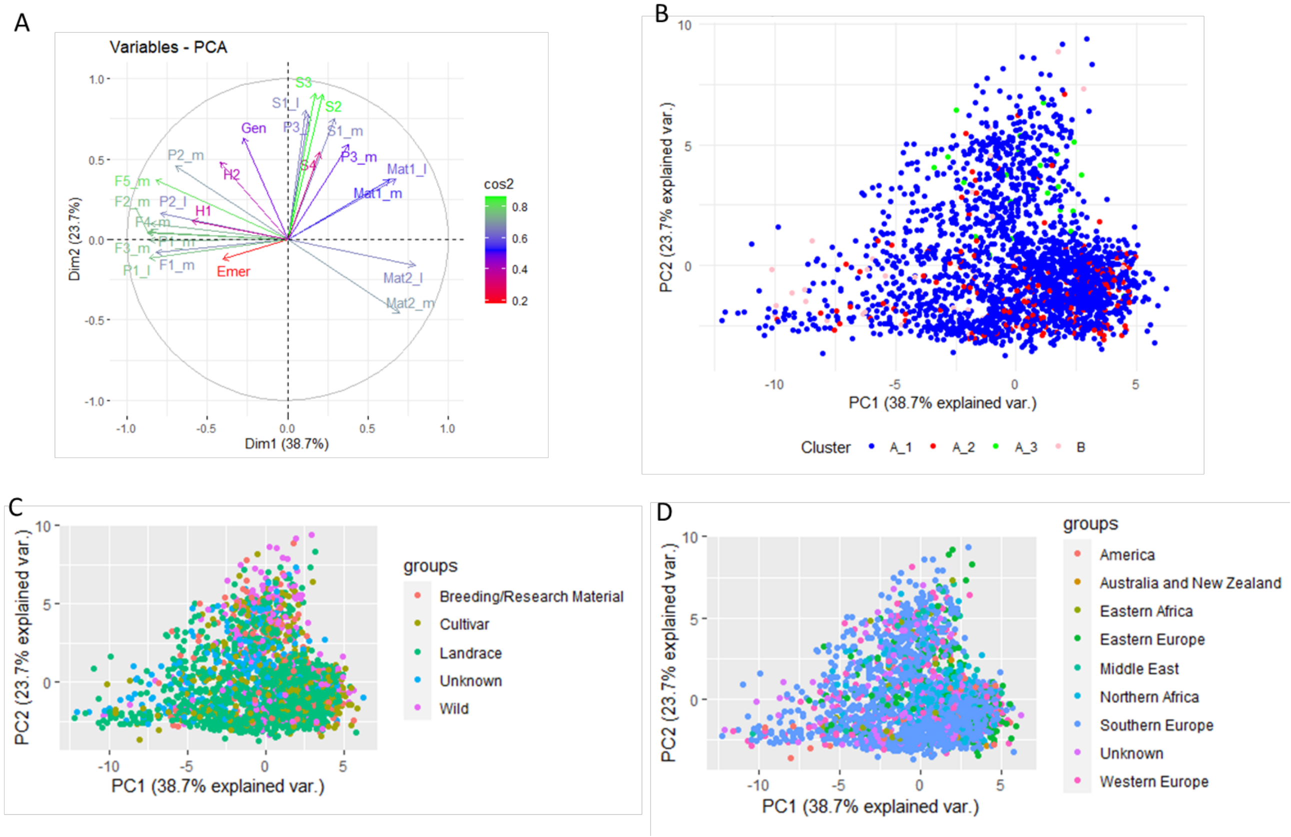


Figure S5: PC1- PC2 biplot resulted from the PCA analysis performed on the data of the 24 quantitative traits (A), and the structure of the phenotypic variation of R-CORE collection of *L. albus* with respect to the phenotypic clusters (B), biological status (C) and geographical regions (D).

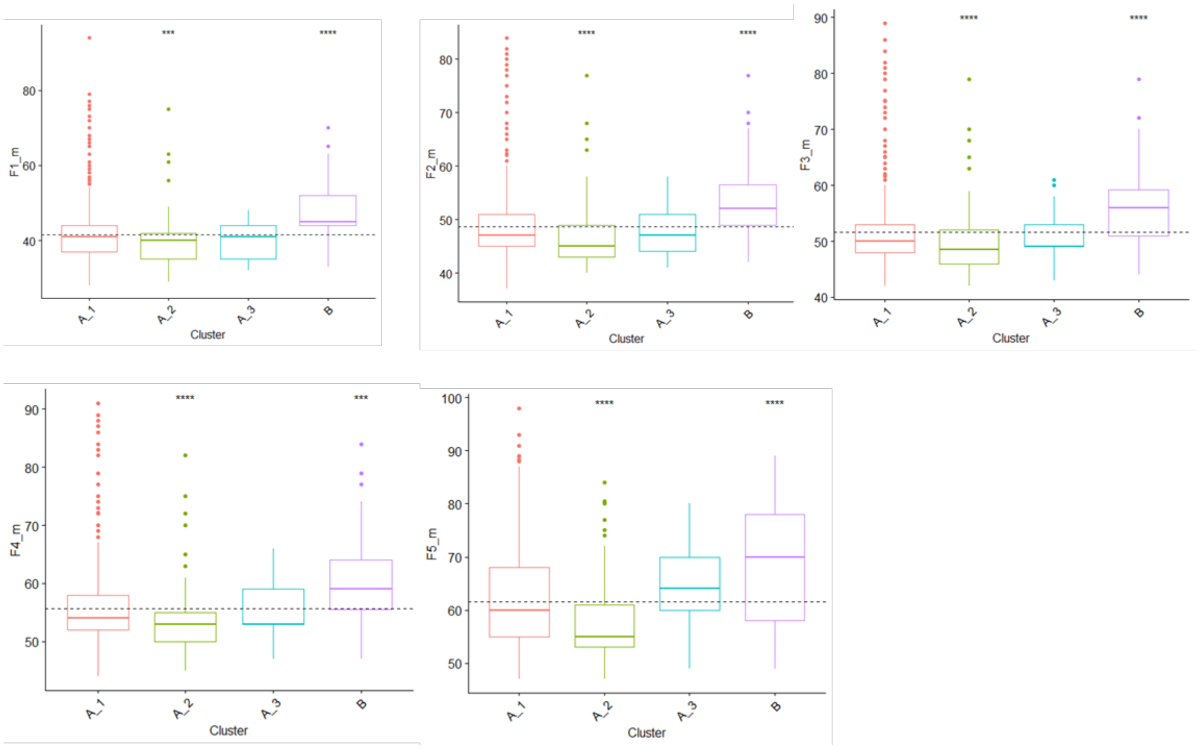


Figure S6: Box plots of flowering traits in phenotypic clusters of *L. albus* R-CORE collection.

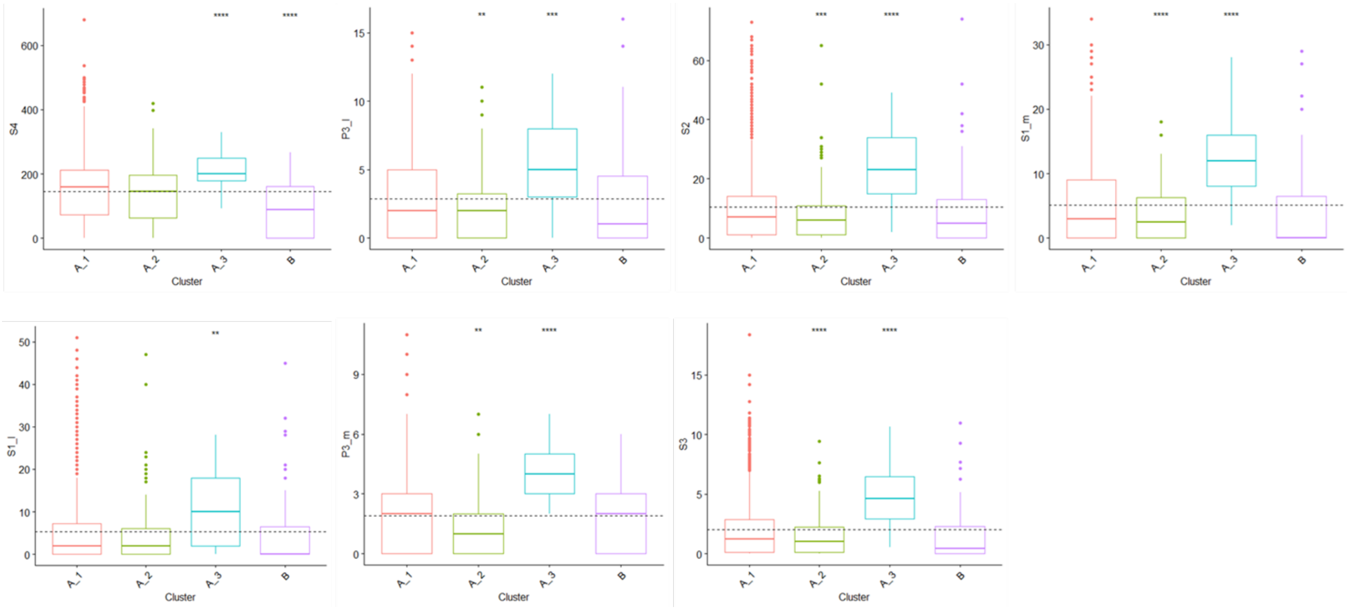


Figure S7: Box plots of seed related traits in phenotypic clusters of *L. albus* R-CORE collection.
